# Supercharging the calcium pump: Identification of an activation hotspot on SERCA by cryo-EM

**DOI:** 10.64898/2026.02.05.703879

**Authors:** Vinh H. Nguyen, Carlos Cruz-Cortés, Joseph O. Primeau, M. Joanne Lemieux, L. Michel Espinoza-Fonseca, Howard S. Young

## Abstract

The sarco-endoplasmic reticulum Ca²⁺-ATPase (SERCA) is a ubiquitous P-type ATPase that restores cytosolic Ca^2+^ to the sarco-endoplasmic reticulum. SERCA is essential for cardiac Ca^2+^ cycling and cellular energy metabolism. Several small molecules enhance SERCA function and show promise in models of metabolic and cardiovascular diseases. However, the structural basis for SERCA activation has remained unknown, hindering mechanism-driven lead optimization. Here we present cryo-EM structures of SERCA bound to two chemically distinct activators: the quinoline derivative CDN1163 (2.6 Å resolution) and a benzofuran derivative UM-52 (3.1 Å resolution). Biochemical assays show that both compounds stimulate Ca^2+^-dependent ATPase activity of SERCA without altering the apparent Ca²⁺ affinity. The structures reveal a previously unrecognized “activation hotspot” in the transmembrane domain, a shallow groove formed by helices M3 and M4 and capped by M1. Despite low chemical similarity, both activators occupy the same pocket and share conserved interactions with Ser^265^, Trp^272^, and Phe^296^. These residues are unique to SERCA and help explain selectivity relative to other P-type ATPases. Activator binding stabilizes a catalytically competent conformation, shifting SERCA toward an E1-like state poised for ATP binding and coordinated movements of the M1-M4 bundle and the cytosolic domains. Notably, density consistent with a detergent acyl chain bridges an otherwise open cavity adjacent to the compound, suggesting that altered protein-lipid interactions may contribute to activation. Together, these findings define a structural framework for SERCA activation and provide a blueprint for rational design of next-generation SERCA activators.

**SIGNIFICANCE STATEMENT:** SERCA pumps Ca²⁺ into the sarco-endoplasmic reticulum, enabling muscle relaxation and shaping calcium signals across tissues. Small-molecule SERCA activators improve cardiac and metabolic phenotypes in animal models, but drug development has been limited by the absence of a defined binding site and activation mechanism. We determined cryo-EM structures of SERCA bound to two distinct activators, CDN1163 and UM-52. Both compounds occupy a groove formed by transmembrane segments M3-M4, anchored by a hydrogen bond to the SERCA-specific Ser^265^ and aromatic contacts near Trp^272^ and Phe^296^. An acyl chain bridges a gap in the binding pocket toward M1, suggesting that protein-lipid coupling may play a role in SERCA activation. These results directly enable structure-mechanism guided design of next-generation selective SERCA activators.

## INTRODUCTION

The sarco-endoplasmic reticulum calcium ATPase (SERCA, EC 7.2.2.10) is an integral membrane protein that is responsible for pumping calcium back into the sarco- or endoplasmic reticulum (SR/ER) from the cytosol. SERCA is found in all eukaryotic cells, where it plays a major role in intracellular calcium homeostasis. Due to its prevalence in the ER of all cell types and the SR of cardiac and skeletal muscles, dysregulation of SERCA-dependent calcium homeostasis leads to a variety of disease states including metabolic disorders, type 2 diabetes, cardiomyopathies, and muscle-wasting diseases. Recent efforts to improve SERCA-dependent calcium homeostasis have focused on small molecules that increase SERCA’s activity. CDN1163 is the most extensively studied small molecule activator of SERCA, demonstrating positive outcomes on metabolic efficiency including reduced ER stress, blood glucose levels, and hepatic lipid accumulation, as well as improved mitochondrial function (1). Additional efforts have identified small molecule activators that increase SERCA activity as much as ∼1.6 fold (2–4). Recently, small-molecule activators, including compound UM-52, were shown to significantly reverse methylglyoxal-induced inhibition of SERCA, revealing a novel therapeutic approach to preserve ER calcium homeostasis in diabetes (5). Despite these advances, there are no structures of SERCA bound to small molecule activators, limiting our understanding of how small molecules increase SERCA’s activity and hindering lead compound optimization.

## RESULTS

We determined cryo-EM structures of SERCA bound to two chemically distinct activators, CDN1163 bound at 2.6 Å resolution and a recently developed small-molecule activator UM-52 (3) bound at 3.1 Å resolution (Table 1). The two structures reveal a novel binding site on SERCA that appears to be unique among the P-Type ATPase family, thereby explaining their selectivity for SERCA. Upon activator binding, SERCA adopts a conformation that is poised for ATP binding, likely shifting the conformational equilibrium in favor of the calcium- and ATP-bound E1 state of SERCA.

**Table 1:** Cryo-EM data collection, refinement and validation statistics.

|  | SERCA+CDN1163<br>(EMDB-73823)<br>(PDB 9Z5S) | SERCA+UM-52<br>(bound)<br>(EMDB-73820)<br>(PDB 9Z5O) | SERCA+UM-52<br>(apo)<br>(EMDB-73822)<br>(PDB 9Z5R) |
| --- | --- | --- | --- |
| **Data collection and processing** |  |  |  |
| Microscope | Titan Krios G3i | Titan Krios G3i | Titan Krios G3i |
| Detector | Falcon 4i | Falcon 4i | Falcon 4i |
| Energy Filter | Selectris X | Selectris X | Selectris X |
| Magnification | 165,000x | 165,000x | 165,000x |
| Voltage (keV) | 300 | 300 | 300 |
| Electron exposure (e-/Å<sup>2</sup>) | 50 | 50 | 50 |
| Defocus range (µm) | -0.6 to -1.8 | -0.6 to -1.8 | -0.6 to -1.8 |
| Pixel size (Å) | 0.743 | 0.743 | 0.743 |
| Symmetry imposed | C2 | C1 | C1 |
| Initial particle images (no.) | 2,513,942 | 2,573,740 | 2,573,740 |
| Final particle images (no.) | 104,006 | 143,855 | 83,166 |
| Map resolution (Å) | 2.5 | 3.1 | 3.8 |
| FSC threshold | 0.143 | 0.143 | 0.143 |
| Map resolution range (Å) | 2.5 – 39.2 | 3.1 – 46.9 | 3.8 – 60.9 |
| **Refinement** |  |  |  |
| Initial model used (PDB code) | 2ZBD | 2ZBD | 2ZBD |
| Model resolution (Å) | 2.8 | 3.2 | 4.1 |
| FSC threshold | 0.5 | 0.5 | 0.5 |
| Map sharpening *B* factor (Å<sup>2</sup>) | -58.9 | -86.2 | -116.6 |
| **Model composition** |  |  |  |
| Non-hydrogen atoms | 15,614 | 7717 | 7668 |
| Protein residues | 1986 | 993 | 993 |
| Ligands | Ca<sup>2+</sup>: 4<br>CDN1163: 2<br>C<sub>12</sub>E<sub>8</sub>: 2 | Ca<sup>2+</sup>: 2<br>UM-52: 1 | Ca<sup>2+</sup>: 2 |
| ***B* factors (Å<sup>2</sup>)** |  |  |  |
| Protein | 158.54 | 54.89 | 183.53 |
| Ligand | 90.43 | 35.85 | 122.34 |
| **R.m.s. deviations** |  |  |  |
| Bond lengths (Å) (no. >4σ) | 0.002(0) | 0.002(0) | 0.003(0) |
| Bond angles (°) (no. >4σ) | 0.480(0) | 0.393(0) | 0.647(0) |
| **Validation** |  |  |  |
| MolProbity score | 2.14 | 2.23 | 2.26 |
| Clashscore | 28.55 | 18.06 | 13.35 |
| Poor rotamers (%) | 0.95 | 2.50 | 2.86 |
| **Ramachandran plot** |  |  |  |
| Favored (%) | 96.77 | 96.97 | 95.96 |
| Allowed (%) | 3.13 | 3.03 | 3.94 |
| Disallowed (%) | 0.10 | 0.00 | 0.10 |
| **Correlation model versus data** |  |  |  |
| CC (mask, volume) | 0.87(0.85) | 0.88(0.87) | 0.85(0.84) |
| CC (ligands) | 0.85 | 0.80 | 0.90 |

CDN1163 is a quinoline derivative first reported in 2016 as a small-molecule activator of SERCA (Figure 1A). ATPase assays of SERCA proteoliposomes showed that CDN1163 increased calcium-dependent ATPase activity by approximately 1.5-fold (Supplementary Data Figure 1). Similarly, surface-electrogenic event reader (SEER) experiments showed that CDN1163 also increased calcium transport by SERCA (6). In this study, single particle cryo-EM resolved two different populations for SERCA in the presence of CDN1163 (Supplementary Data Figure 2), a monomer and an antiparallel dimer (cytoplasmic domains oriented on opposite sides of the micelle). The presence of SERCA monomers and dimers was also observed by mass photometry (Supplementary Data Figure 3). Interestingly, density corresponding to CDN1163 was only found at the dimerization interface between SERCA molecules in the antiparallel dimer (Figure 1A). We hypothesized that the antiparallel dimer stabilized CDN1163-bound SERCA, preventing its dissociation before freezing for cryo-EM single-particle analysis. We also assumed that activator binding must be inherently distinct from SERCA inhibitors such as thapsigargin (7) and cyclopiazonic acid (8, 9), which occupy relatively deep binding pockets and lock SERCA in a calcium-free state that prevents progression through the calcium transport cycle (Supplementary Data Figure 4).

**Figure 1:**
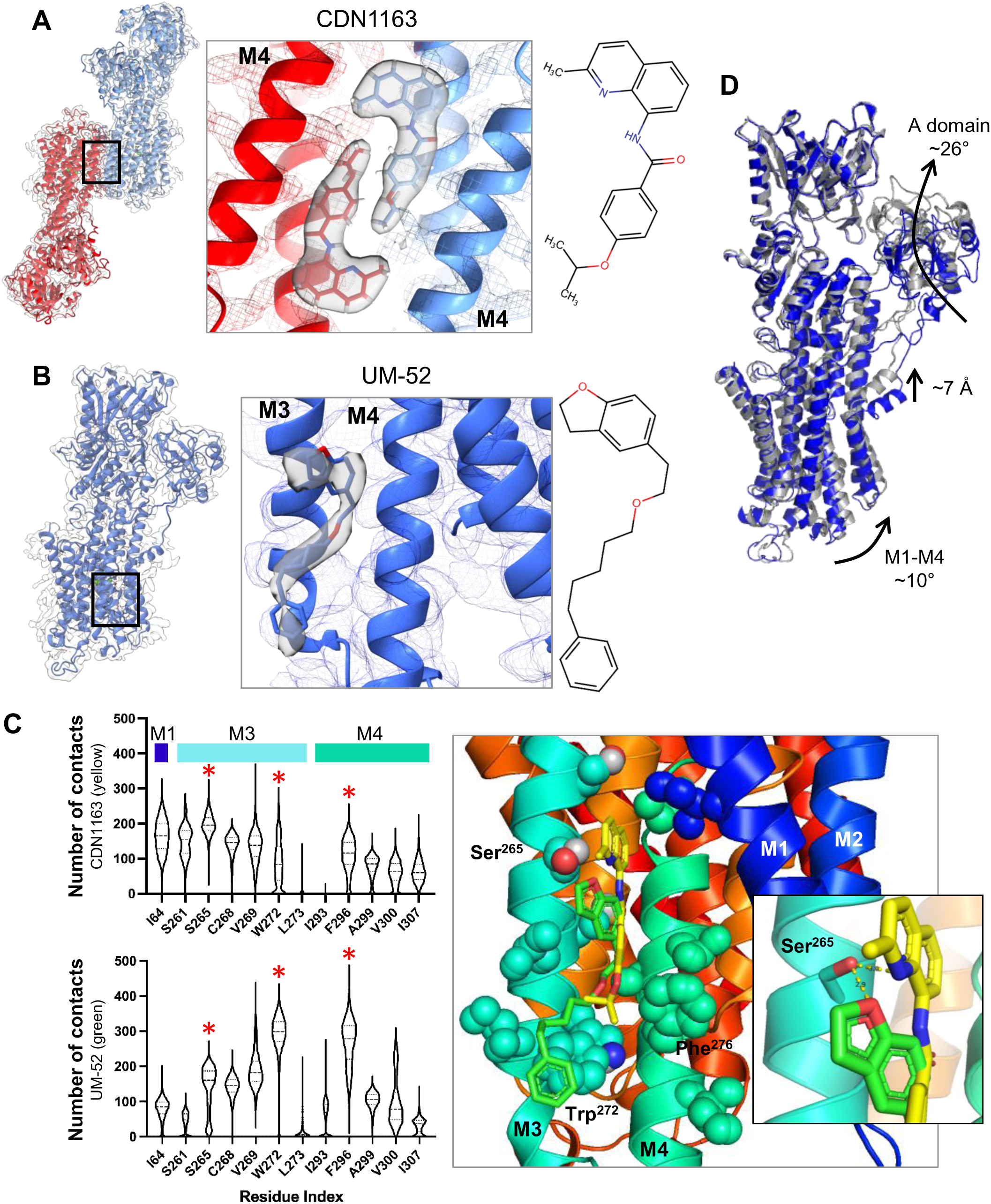
(**A**) Cryo-EM density map (mesh) of SERCA (blue and red ribbon diagrams) in the presence of CDN1163 adopted an antiparallel dimer configuration. Density for SERCA was represented as a mesh, whereas activator density was represented as a grey volume. The chemical structure of CDN1163 is shown. (**B**) Cryo-EM density map (mesh) of SERCA (blue ribbon diagram) in the presence of UM-52 was predominantly monomeric, with UM-52 binding at the same site as CDN1163. Density for SERCA was represented as mesh, whereas UM-52 density was represented as a grey volume. The chemical structure of UM-52 is shown. (**C**) Number of contacts between CDN1163 and UM-52 and specific residues in M1, M3, and M4, obtained from molecular dynamics simulations and represented as violin plots. The residues indicated by asterisks are shown as spheres on the ribbon diagram of SERCA with CDN1163 (yellow) and UM-52 (green). Inset shows the hydrogen bond distances between Ser^265^ and the quinolone nitrogen of CDN1163 (3.0 Å) and the benzofuran oxygen of UM-52 (2.9 Å). (**D**) Superposition of apo SERCA (blue ribbon diagram) versus CDN1163- and UM-52-bound SERCA (grey ribbon diagrams) indicating the domain movements upon CDN1163 and UM-52 binding.

The structure of CDN1163 bound to SERCA suggested two possible interpretations. CDN1163 binding to an activation site on SERCA may have provided a complementary interface for antiparallel dimer formation. Alternatively, a preformed antiparallel dimer of SERCA may have provided an interface for CDN1163 binding that was not relevant to the mechanism of activation. To discriminate between these two possibilities, we determined the structure of SERCA in the presence of a recently developed SERCA activator, UM-52, a potent activator with low chemical resemblance to CDN1163 (Figure 1B; ∼24% RDKit Fingerprint and Tanimoto similarity coefficient). ATPase assays of SERCA proteoliposomes showed that UM-52 increased calcium-dependent ATPase activity by approximately 1.6-fold (Supplementary Data Figure 1). Interestingly, cryo-EM images of SERCA in the presence of UM-52 appeared mainly monomeric, which could be further separated into apo and bound populations (Supplementary Data Figures 5 & 6). The activator UM-52 was found to bind to a similar site to CDN1163 involving transmembrane segments M3 and M4 of SERCA. While UM-52 has a 2,3-dihydrobenzofuran group instead of the quinolone derivative in CDN1163, the benzofuran oxygen of UM-52 and the quinolone nitrogen of CDN1163 form a hydrogen bond with Ser^265^ on M3 (Figure 1C). Both UM-52 and CDN1163 cause SERCA to adopt the same conformation (RMSD 0.3 Å), suggesting that their modes of activation are similar (Figure 1D). We were able to resolve a SERCA monomer bound to the activator, UM-52, suggesting that it may form a more stable interaction with SERCA compared to CDN1163. This may be due to the interaction of a phenyl ring of UM-52 with Trp^272^ of SERCA (Figure 1C). Despite these differences, the observation that two chemically distinct activators occupy the same groove on SERCA supports the existence of a genuine activator site and indicates that the antiparallel SERCA dimer did not force CDN1163 into an artificial binding mode.

The activators bind along transmembrane segment M3 of SERCA in a shallow groove lined by M4. M1, M3, and M4 play different roles in the formation of the activator binding site. M4 runs parallel to the activators and small hydrophobic residues (e.g. Ala^303^, Ala^299^) allow formation of a cavity that accommodates the quinoline and benzofuran groups of CDN1163 and UM-52, respectively (Figures 1C & 2A; Supplementary Data Figure 7). M3 also runs parallel and lines one face of the activator binding site, making the closest contacts among the three helices and contributing van der Waals interactions via several small non-polar residues to stabilize the activators (e.g. Cys^266^, Val^269^). M3 contains two key residues that are unique to SERCA that are important for positioning the activators in the binding pocket. Trp^272^ stabilizes one end of the activators through CH-π interactions with the isopropyl-group of CDN1163 and π-stacking with the phenyl-group of UM-52 (Figure 1C). Phe^296^ in M4 is near Trp^272^ and could also play a role in forming an aromatic cleft and π-network to stabilize the activators (both residues are ∼4 Å away from the isopropyl group). Importantly, hydrophobicity in this region of the pocket appears to be essential for stabilizing activator binding, as polar substitutions on the aryl-ring of UM-52 derivatives yield inactive compounds (3). As mentioned above, Ser^265^ is an important residue unique to SERCA that forms a hydrogen bond with the activators, aiding in their positioning in the binding site. The specificity of the activators for SERCA is likely due to Ser^265^ as other P-type ATPases do not have a residue that can act as a hydrogen-bond donor at this position (Figure 2 & Supplementary Data Figure 8). Ser^265^ appears to play an important role in activator positioning because UM-52 sits lower in the binding pocket compared to CDN1163 to allow formation of a hydrogen bond between the 2,3-dihydrobenzofuran group and the serine (Figure 1C). Recent SERCA activator development has noted the importance of a hydrogen-bond capable atom (with hydrogen-bond accepting oxygen producing higher stimulation than hydrogen-bond donor nitrogen) in the fused-ring of the activators, supporting the role of Ser^265^ in facilitating hydrogen-bond interactions (3). Importantly, the structures support the notion that balanced polarity within the fused ring is essential for maintaining activity, consistent with experimental data showing that even modest changes in polarity convert SERCA activators into inhibitors (3). Thus, M3 interactions appear necessary for activator binding, while M4 provides a sufficient cavity for insertion of the quinoline and benzofuran ‘head groups’ of the activators.

**Figure 2:**
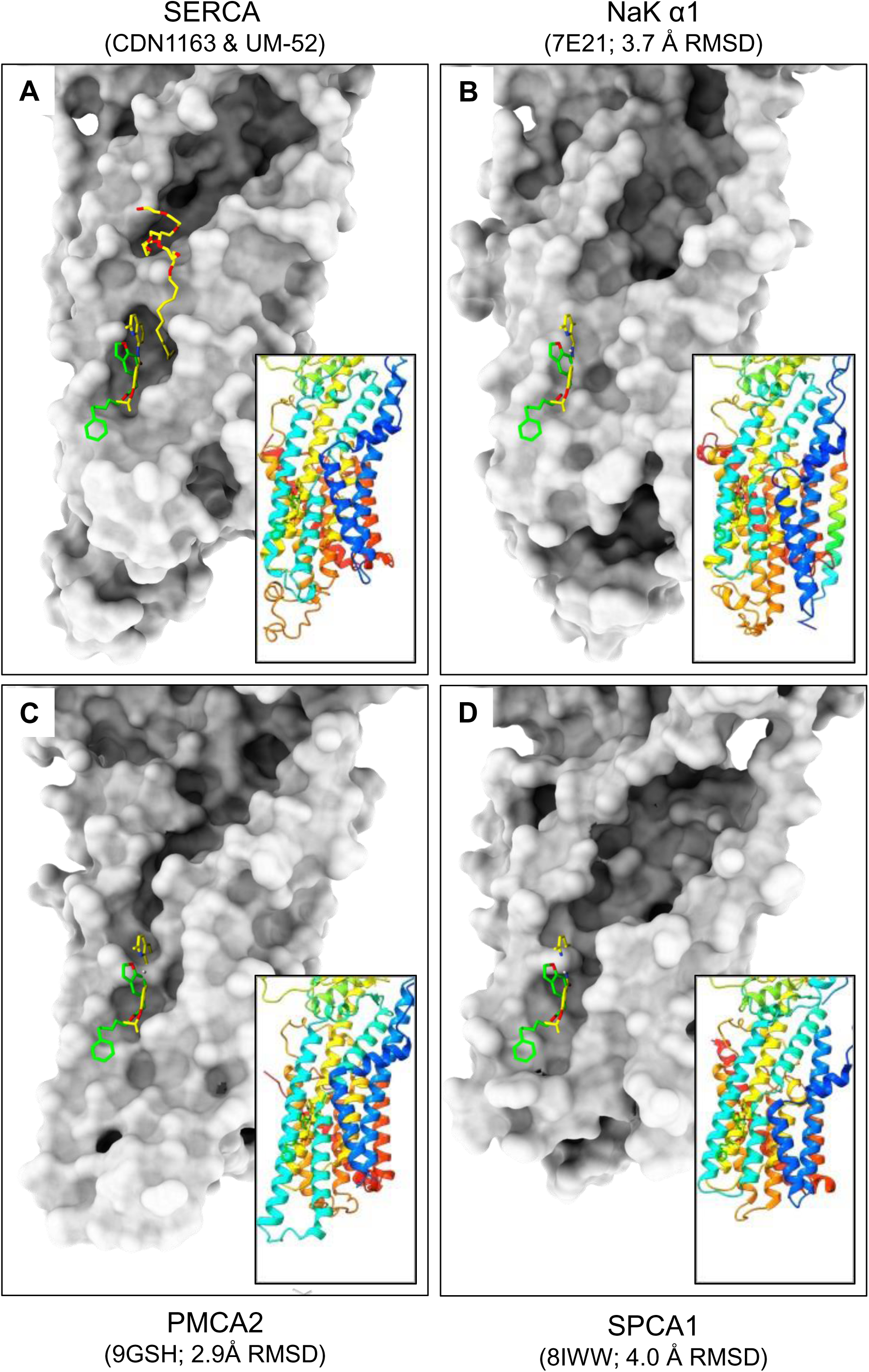
(**A**) Surface representation of SERCA with CDN1163, UM-52, and a C_12_E_8_ molecule represented as sticks within the binding site of SERCA. A ribbon diagram is also shown as an inset. (**B**) Surface representation and ribbon diagram of the Na^+^,K^+^-ATPase aligned with SERCA (RMSD 3.7 Å; PDB code 7E21) indicating the relative positions of the activators. Surface representations and ribbon diagrams of the (**C**) plasma membrane Ca^2+^-ATPase aligned with SERCA (RMSD 2.9 Å; PDB code 9GSH) and (**D**) secretory pathway Ca^2+^-ATPase aligned with SERCA (RMSD 4.0 Å; PDB code 8IWW) indicating the relative positions of the activators. Steric clashes with the other P-type ATPases likely prevent binding of the SERCA activators.

M1 appears to play a relatively minor role, with Leu^60^ and Ile^64^ forming the cap of the activator head group binding site (Supplementary Data Figure 7). Interestingly, there is a large gap in the binding cavity (∼10 Å distance) between the activator head groups and M1, where van der Waals interactions were absent. However, there was a long density that could be fit by a C_12_E_8_ detergent molecule in the SERCA-CDN1163 map that tunneled from the aqueous environment into the space between CDN1163 and M1 (Figure 2A). This suggests that an acyl chain of a detergent or lipid may stabilize CDN1163 and provide a mobile element that links activator binding to M1. A lipid acyl chain occupying the space between activator and M1 may allow the movements of M1, M2 and the A domain that facilitate ATP-hydrolysis and calcium transport while still activator-bound. The presence of an acyl chain could explain why small-molecule activators have planar head groups, as the M1 gap and lipid binding may be necessary for activation. We hypothesize that a potential activator having a chemical moiety that fills this gap could immobilize M1 and lead to SERCA inhibition, effectively turning the activator into an inhibitor.

## DISCUSSION

Several potential binding sites have been proposed for activator binding, including both the cytoplasmic and transmembrane domains of SERCA (4), but these studies did not predict the binding site presented here. Interestingly, the activator binding site coincides with an alternative binding mode of phospholamban (PLN) and sarcolipin (SLN) to SERCA (10, 11). PLN and SLN were found to interact with transmembrane segment M3 and modulate the maximal activity of SERCA, which is consistent with the binding of CDN1163 and UM-52 to M3 of SERCA. Among the P-type ATPases, SERCA possesses a novel binding site for small-molecule activators in the groove formed by M1, M3, and M4 (Figure 2A). Surface representation of other P-type ATPases whose structures are known, such as the plasma membrane calcium ATPase, the Na^+^, K^+^ ATPase, and the secretory pathway calcium ATPase, reveal that these family members lack a cavity along M3 and M4 that could accommodate CDN1163 or UM-52 (Figure 2). While there are cavities in the M3/M4 region of PMCA2 and SPCA1 that may be suitable for structure-guided drug development (Figure 2C, D), none of the other P-type ATPases can accommodate the chemical structures of the SERCA activators, suggesting that this effector site is unique to SERCA.

The structure of SERCA in the bound complexes is similar for both CDN1163 and UM-52 (RMSD 0.3 Å). When compared to the apo (unbound) structure of SERCA in the presence of UM-52, they reveal the mechanism behind SERCA activation that has remained elusive, where both small molecules cause SERCA to adopt a conformation that closely resembles the E1, calcium-and ATP-bound state of SERCA. The activator-bound structures of SERCA are most similar to SERCA2b in the E1·2Ca state (RMSD 1.3 Å; PDB code 7E7S (12)), indicating that CDN1163 and UM-52 promote a conformation of SERCA with bound calcium that is poised to bind ATP without significant changes in the cytoplasmic domains. We suggest that activator binding in the presence of calcium causes SERCA to adopt a catalytically competent conformation that will readily hydrolyze ATP, thus shifting the equilibrium towards E1∼P·ADP·2Ca conformation. In comparison to the unbound apo structure of SERCA (Figure 1D), M1-M4 undergo a ∼10° rotation toward the cytoplasmic domains and the A domain undergoes a ∼26° upward rotation and contacts the N-domain to poise SERCA for ATP binding. With movement of the A domain, the cytoplasmic domains are in a more compact conformation compared to the E1·2Ca conformation, which aligns with previous TCSPC studies showing that CDN1163 induced closure of the cytoplasmic headpiece (13). The apo structure of SERCA is most similar to the E1-like state in the presence of SLN (14, 15) and PLN (16) (e.g. RMSD 2.5 Å; PDB code 3W5B), though calcium is bound in this case rather than magnesium (14, 15).

Overall, the structures presented here suggest a novel mechanism where chemically distinct small-molecule activators bind to a similar site in the transmembrane domain of SERCA and promote the transition from the E1-like state to the catalytically competent E1·2Ca·ATP state of the calcium pump. The question remains as to how the small molecules promote this transition. The location of the binding site and the presence of an acyl chain in the SERCA-CDN1163 complex (Figure 2A) suggest that activator binding may alter lipid interactions as a mechanism for increasing SERCA activity.

## METHODS

### Purification of SERCA from rabbit hind leg muscle

Purification of SERCA was carried out as previously described (17, 18). Briefly, rabbit hind leg, fast-twitch muscle was harvested, homogenized, and sarcoplasmic reticulum (SR) membranes were collected by differential centrifugation. SR membranes were solubilized in 1% C_12_E_8_ (Sigma-Aldrich) and SERCA was purified by affinity chromatography using Reactive Green-19 cross-linked agarose (Sigma-Aldrich). Fractions containing SERCA were concentrated using a 100 kDa MW cut-off Amicon centrifugal filter to approximately 30 µL and incubated with 10 µM CDN1163 (MedChemExpress). Sample was loaded onto a Superdex 200 column pre-equilibrated with 50 mM MOPS pH 7.0, 5% glycerol, 100 mM KCl, 1 mM CaCl_2_, 10 µM CDN1163, 1 mM DTT, and 0.006% C_12_E_8_. Fractions were pooled and concentrated to approximately 20 µL, flash frozen in liquid N_2_ and stored at −80°C. Concentrated SERCA samples were analyzed by SDS-PAGE (19) to assess purity. Sample preparation for SERCA-UM-52 complex followed the same procedure.

### ATPase activity assays

C_12_E_8_-solubilized SERCA was reconstituted into proteoliposomes as previously described (20). Briefly, SERCA was added to a thin-lipid film of egg yolk phosphatidylcholine and egg yolk phosphatidic acid (EYPC & EYPA; Avanti Polar Lipids) and Biobeads SM-2 (Bio-Rad Laboratories) were slowly added over a four-hour time course to promote proteoliposome formation through detergent adsorption. Proteoliposomes were isolated by ultracentrifugation at 137,000 RCF with a 20/50 % sucrose step-gradient. Proteoliposomes were collected from the sucrose step-gradient interface, flash frozen in liquid N_2_ and stored at −80°C.

Calcium-dependent ATPase activity assays were performed as previously described using a coupled enzyme assay adapted to a 96-well format (21, 22). A Biotek Epoch2 (Agilent) plate reader was used to measure the decrease in absorbance at 340 nm due to NADH depletion coupled to ATP hydrolysis over a range of calcium concentrations in the presence or absence of 100 µM CDN1163 or UM-52. The purity of the activators was >95% by HPLC. Specific activities of SERCA were calculated at each calcium concentration range, and SigmaPlot (Grafiti LLC) was used to fit the data to the Hill equation.

### Sample quality control by mass photometry

Mass photometry (TwoMP, Refeyn Ltd. (23)) was used to monitor sample quality for cryo-EM by determining the oligomerization behavior and relative detergent micelle concentrations of candidate samples following gel-filtration chromatography and sample concentration. Mass calibrations were conducted using bovine serum albumin and apoferritin as standards in PBS buffer. For SERCA-containing samples, buffer (20 mM MOPS pH 7.0, 5% glycerol, 100 mM KCl, 1 mM CaCl_2_, 1 mM DTT, 0.004% C_12_E_8_) was used to focus the TwoMP (20 µL) followed by the addition of 0.2 µL of SERCA sample. One-minute recordings were collected and processed using the DiscoverMP software. Samples that generated distinct SERCA peaks with relatively small peaks for free detergent micelles were selected for preparation of frozen-hydrated grids, cryo-EM, and single particle analysis.

### Cryo-EM grid preparation

Quantifoil® Cu 300 R 2/1 grids were glow-discharged in a Pelco easiGlow™ for 30 seconds at 15 mV. SERCA (3 µL at 10 mg/mL) in the presence of activators (10 µM) was applied to the grids, blotted for 7-10 s with zero blot force, then plunge-frozen in liquid ethane using a FEI Vitrobot mark IV (Thermo Fisher Scientific) with 100% humidity at 4°C. Frozen grids were screened at the Stanford-SLAC cryo-EM centre (S^2^C^2^) on a Titan Krios operated at 300 kV equipped with a Selectris X energy filter and a Falcon 4i detector (Thermo Fisher Scientific). Data was collected using EPU in fast acquisition mode on candidate grids at a nominal magnification of 165k with a pixel size of 0.743 Å. Images were collected with a 6.2 s exposure over 43 frames, resulting in 1.16 e^-^ /Å^2^ per frame with a total dose of 50 e^-^. The defocus range of the data was −0.6 µm to −1.8 µm.

### Data Processing and Model Building

Micrographs were processed using CryoSPARC (Structura Biotechnology Inc. (24)), with workflow summarized in the Extended Data Figures. Density maps for a SERCA monomer and antiparallel dimer in the presence of CDN1163 were determined at 3 Å and 2.6 Å, respectively. Unsharpened maps were used for subsequent model building. Density maps for unbound (apo) and bound SERCA monomers in the presence of UM-52 were determined at 3.8 Å and 3.1 Å, respectively. The atomic coordinates for SERCA were obtained from a previously published structure (PDB ID: 2ZBD), with individual domains of SERCA (A-, N-, P- and TM-domains) isolated as separate PDB files, and each domain was sequentially fitted into the cryo-EM density maps using Phenix dock in map program (25). Manual model-building along with ligand placement was performed in COOT (26), with eLBOW (27) used to generate CDN1163 and UM-52 restraints from the SMILES input. Models were refined in Phenix using real-space refinement after each manual model-building stage, with Molprobity (28) used for model validation. Model statistics can be found in Extended Data Table 1. Figures were created using USCF Chimera (29) and ChimeraX (30).

### Molecular dynamics simulations

The structures of SERCA bound to CDN1163 or UM-52 were inserted in a pre-equilibrated 120×120 Å bilayer that contained POPC and POPE (2:1 ratio) lipids to mimic the lipid composition of the SR. For lipids, we used the LIPID21 force field (31) and a cutoff distance of 10 Å for non-bonded interactions. We used the replacement method to generate lipid packing around the protein-ligand complex. We solvated each system using the OPC water model with a minimum margin of 20 Å between the protein and the z-axis edges. K^+^ and Cl^-^ ions were added to neutralize the system and to produce a KCl concentration of 100 mM. The systems were prepared using the CHARMM-GUI web server (32, 33). Energy minimization and equilibration were performed as follows: two 25-ps restrained canonical ensemble (NVT) simulations, one 25-ps restrained isothermal-isobaric ensemble (NPT) simulation, three 250-ps restrained NPT simulations, and a single unrestrained NPT simulation for 5 ns. The Langevin thermostat was used to keep the temperature at 25°C and the Monte Carlo barostat to maintain a constant pressure of 1.0 bar. Bonds involving hydrogen atoms were constrained using the SHAKE algorithm. We performed three independent 1-µs MD simulations of the complexes using AMBER24 on Tesla V100 GPUs (34) and the AMBER ff14SB force field (35). Data analysis was performed using VMD 2.0 (36).

## Acknowledgements

Some of this work was performed at the Stanford-SLAC Cryo-EM Center (S^2^C^2^), which is supported by the National Institute of General Medical Sciences (1R24GM154186). The content is solely the responsibility of the authors and does not necessarily represent the official views of the National Institutes of Health. The authors would also like to thank the following S^2^C^2^ personnel for their invaluable support and assistance: Grace Nye, Alexandre Cassago, Yan Liu, and Nathan D. Burrows. This work was funded by the Heart and Stroke Foundation of Canada (to HSY), the Canadian Institutes of Health Research (PJT-180387 to HSY), the National Institutes of Health (R01GM120142 and R01HL148068 to LMEF), and the Natural Sciences and Engineering Research Council of Canada (549297-2019 to MJL). This research was supported in part through computational resources and services provided by Advanced Research Computing at the University of Michigan, Ann Arbor, Michigan.

## Author Contributions

VHN, HSY and MJL conceptualized the study; HSY and VHN designed the experiments; LMEF and CCC provided a small-molecule activator; VHN purified proteins, performed biochemical measurements, data analyses, and cryo-EM sample preparation; VHN and JOP performed data collection and data processing; VHN generated atomic models on the basis of cryo-EM maps; VHN and HSY wrote the paper with comments from all authors.

## Data Availability

Cryo-EM maps and atomic coordinates were deposited into the EMDB and RCSB PDB, respectively. SERCA-CDN1163 bound maps and coordinates were assigned EMD-73823 and PDB 9Z5S. SERCA-UM52 bound maps and coordinates were assigned EMD-73820 and PDB 9Z5O, whereas the apo SERCA was assigned EMD-73822 and PDB 9Z5R. The coordinates for the initial models used are available in the PDB under accession codes 2ZBD (SERCA E1 Ca^2+^ ADP-AlF4).

## Supplementary Data Figure Legends

**Supplementary Data Figure 1:**
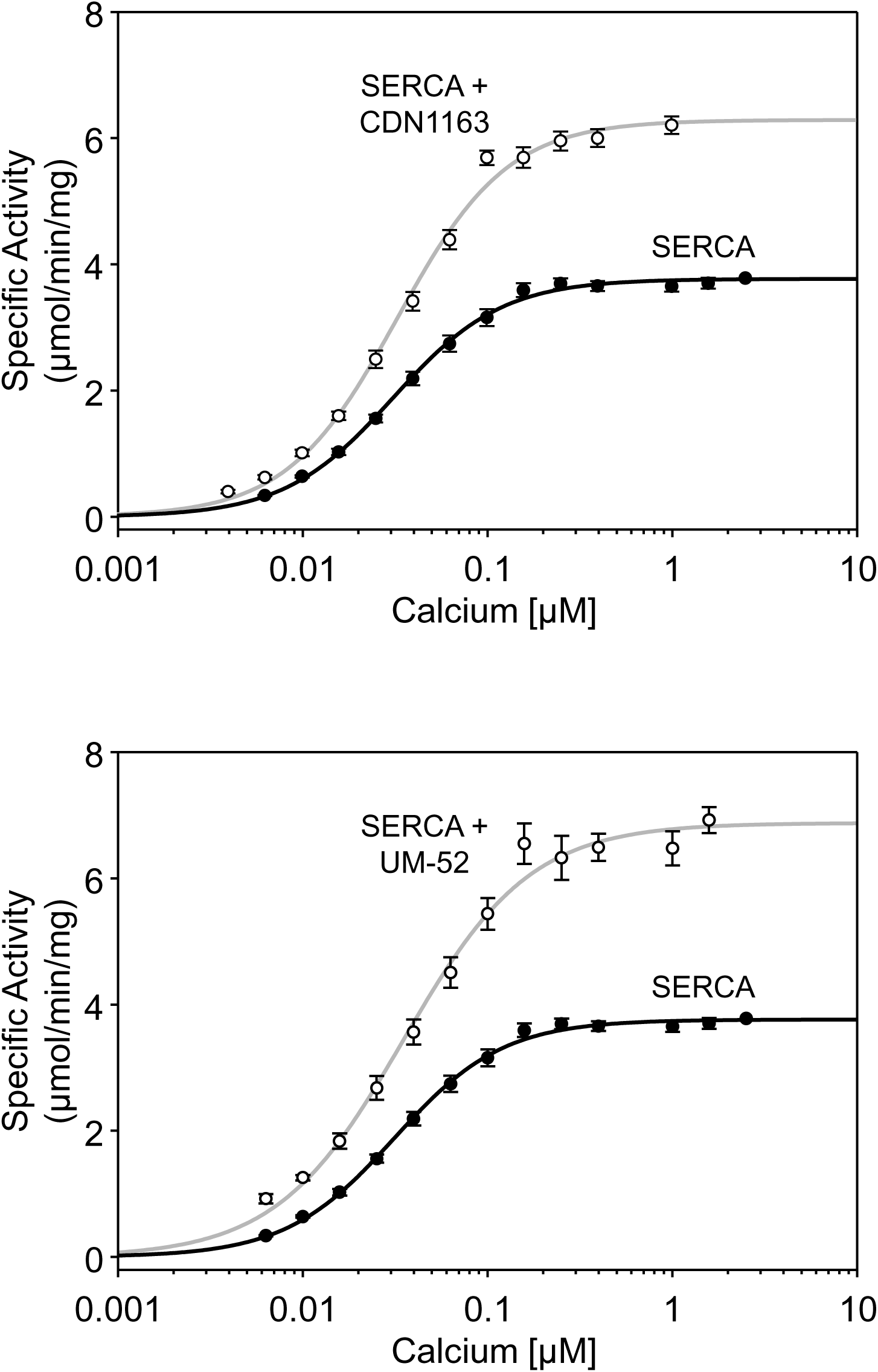
Calcium-dependent ATPase activity data for SERCA in the absence (filled circles) and presence (open circles) of CDN1163 (upper panel) or UM-52 (lower panel). Stimulation of SERCA maximal activity (V_max_) was observed upon addition of either CDN1163 or UM-52 (1.5-fold and 1.6-fold, respectively), without altering the apparent calcium affinity of SERCA (K_Ca_). Each data point corresponds to a biological replicate of three independent proteoliposome preparations (± SEM). Curves were generated by fitting the data to the Hill equation (SigmaPlot Software, Grafiti LLC).

**Supplementary Data Figure 2:**
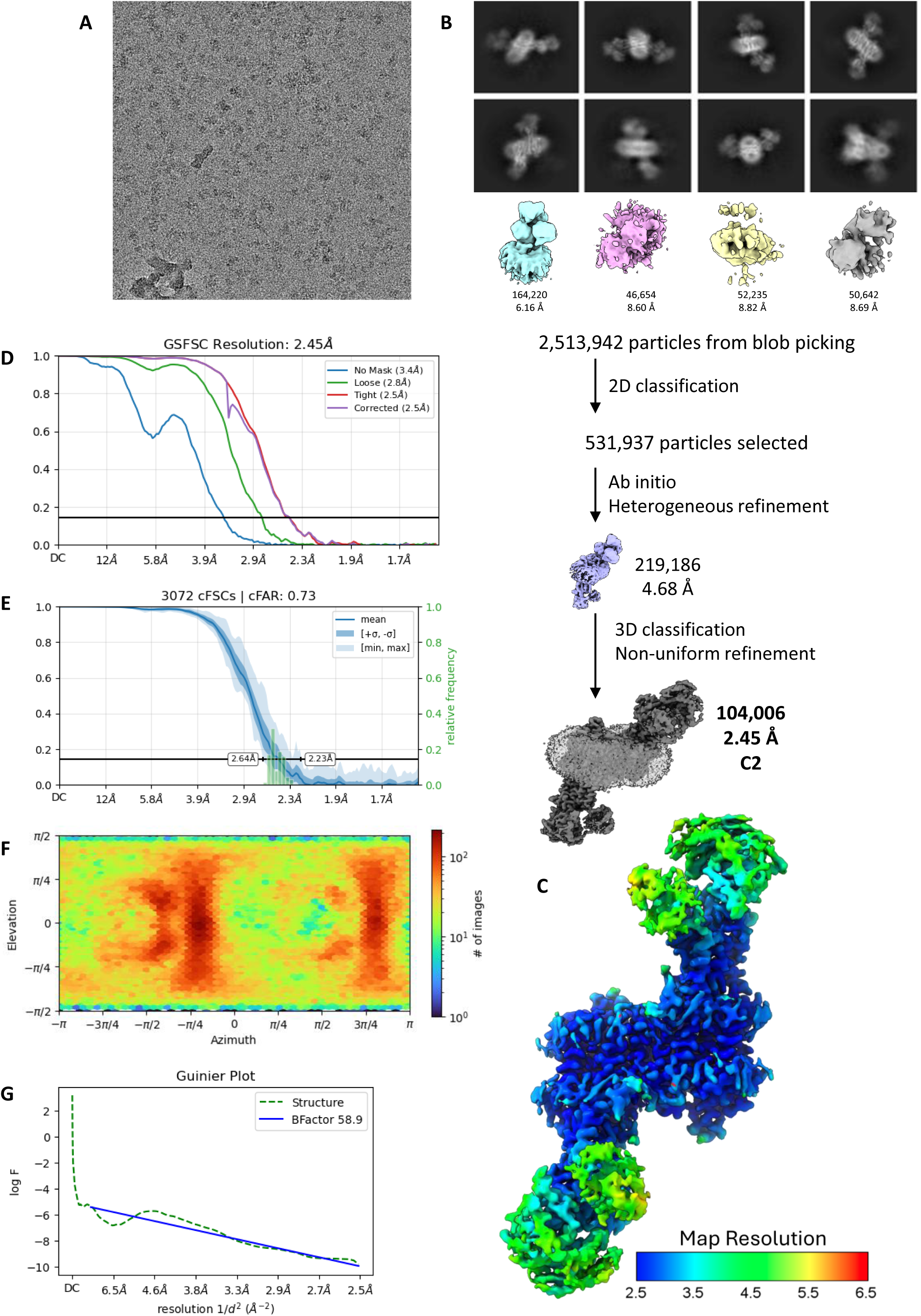
(**A**) Representative micrograph of the SERCA + CDN1163 sample, low-pass filtered to 3 Å. (**B**) Summary of cryo-EM data processing, showing representative 2D classes, ab initio volumes, and the final reconstruction of an antiparallel SERCA dimer. (**C**) Local resolution estimation of the antiparallel SERCA dimer showing high resolution for the transmembrane domain and lower resolution in the cytoplasmic domains. (**D**) Gold Standard Fourier Shell Correlation (GSFSC) as a function of resolution for the map shown in (C). (**E**) Conical FSC area ratio (cFAR) indicates the absence of orientation bias. (**F**) Viewing direction distribution plot for the shown map. (**G**) Guinier plot for the shown map showing the approximate B-factor.

**Supplementary Data Figure 3:**
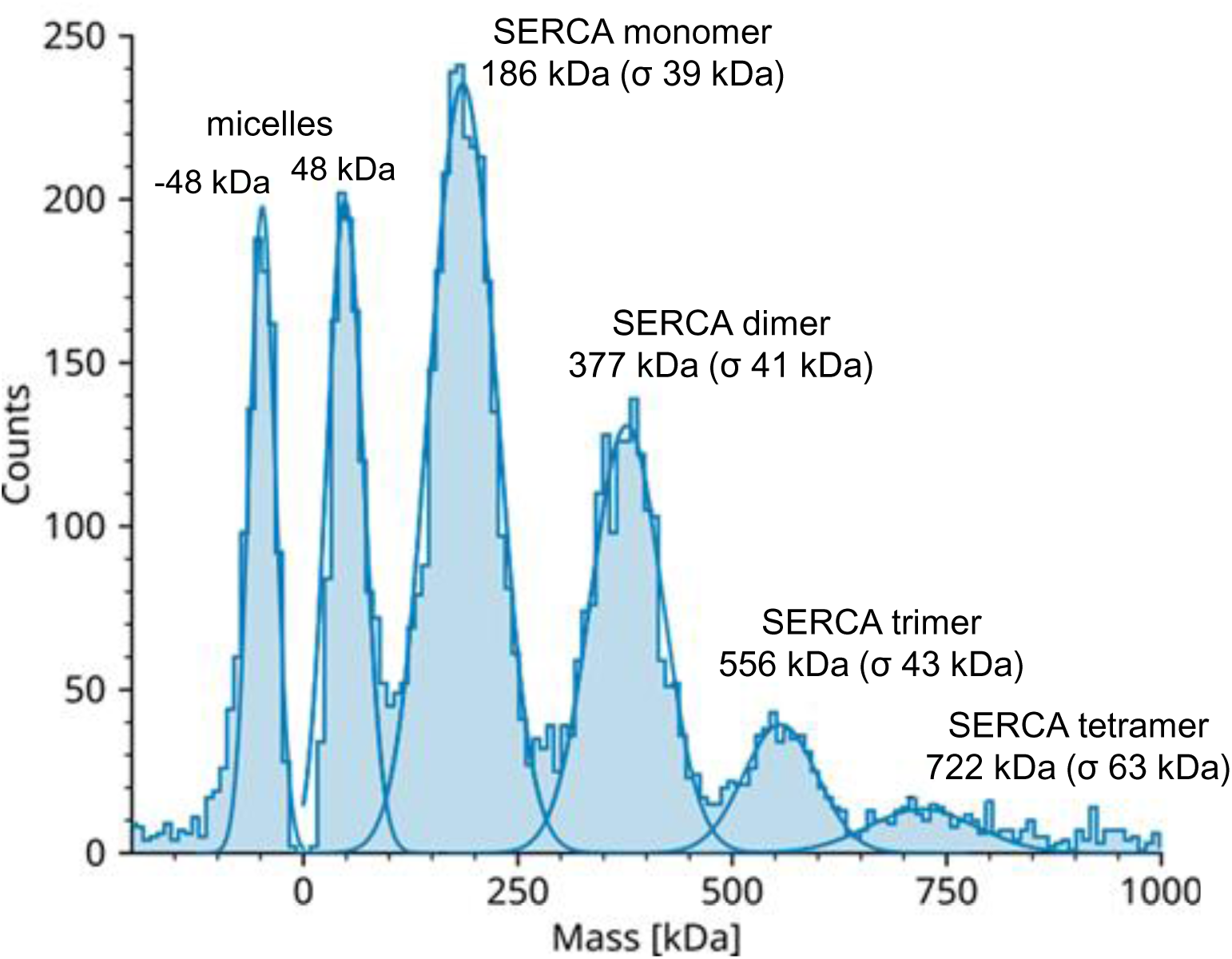
Evaluating sample quality. Histogram of mass photometry counts of SERCA in the presence of CDN1163 shows clear peaks corresponding to detergent micelle, along with SERCA in various oligomeric states. 20 µL buffer was used to focus the TwoMP optics, 0.2 µL of SERCA was added, and data was recorded for 60 seconds. Bovine serum albumin and apoferritin were used as molecular weight standards in PBS buffer.

**Supplementary Data Figure 4:**
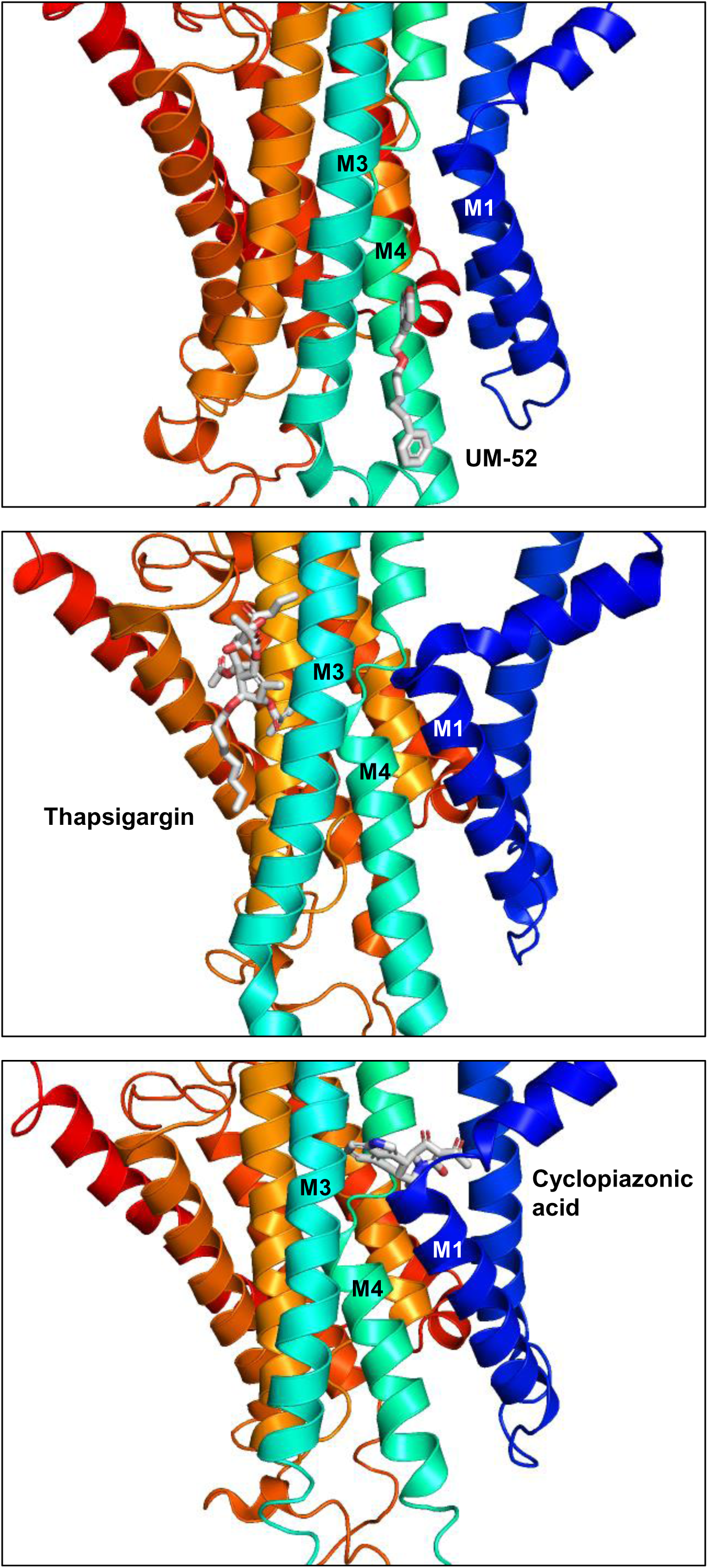
Distinct binding sites for SERCA activators and inhibitors. Ribbon diagrams of SERCA with M1, M3, and M4 labeled for SERCA in the presence of the activator UM-52 (upper panel), the inhibitor thapsigargin (middle panel; PDB code 3AR4), and the inhibitor cyclopiazonic acid (lower panel; PDB code 3FPB).

**Supplementary Data Figure 5:**
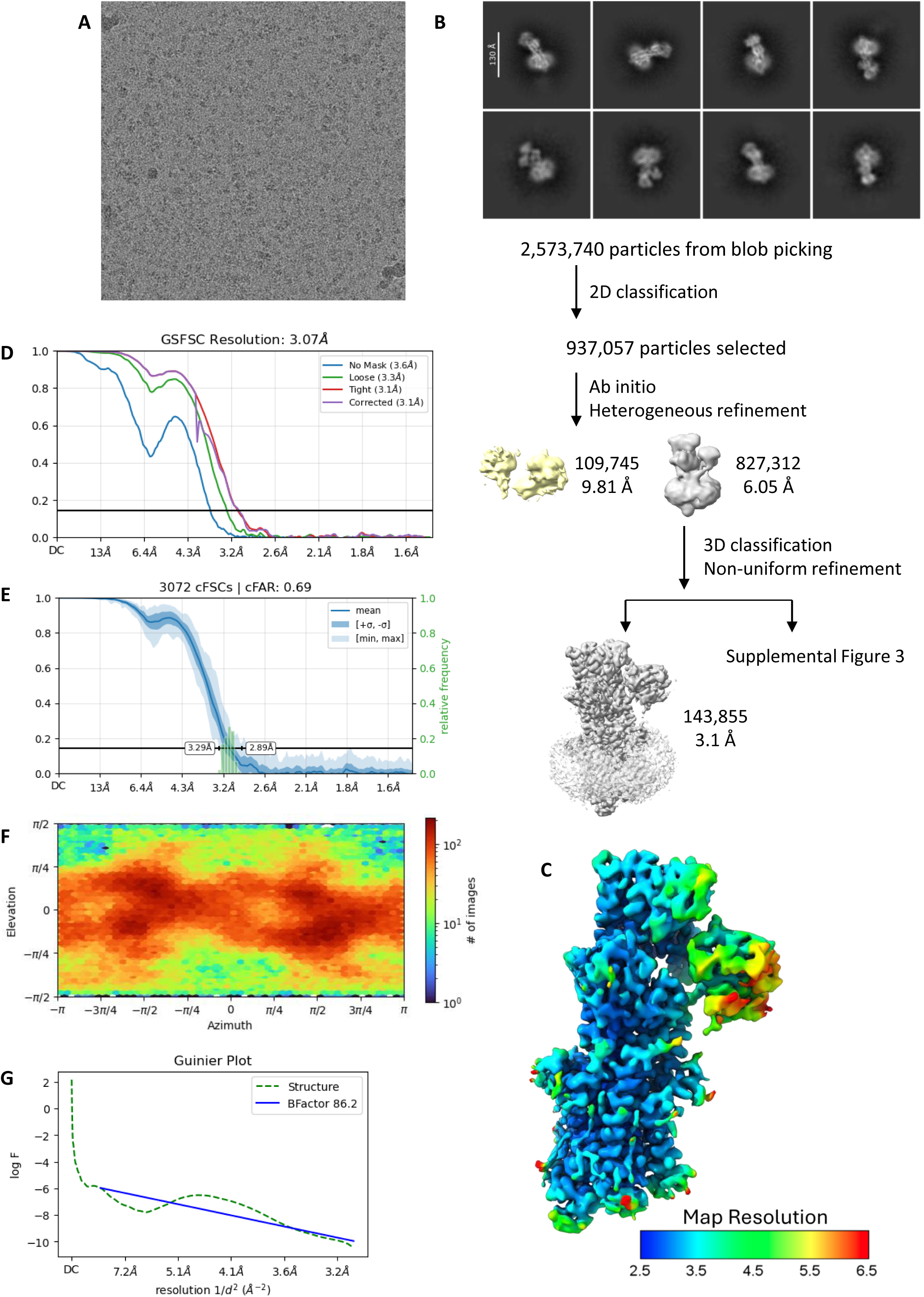
Cryo-EM workflow for SERCA bound to UM-52. (**A**) Representative micrograph of the SERCA + UM-52 sample, low-pass filtered to 3 Å. (**B**) Summary of cryo-EM data processing, showing representative 2D classes, ab initio volumes, and the final reconstruction of a bound SERCA monomer. (**C**) Local resolution estimation of the SERCA monomer showing high resolution for the transmembrane domain and lower resolution in the actuator domain. (**D**) Gold Standard Fourier Shell Correlation (GSFSC) as a function of resolution for the map shown in (C). (**E**) Conical FSC area ratio (cFAR) indicates the absence of orientation bias. (**F**) Viewing direction distribution plot for the shown map. (**G**) Guinier plot for the shown map showing the approximate B-factor.

**Supplementary Data Figure 6:**
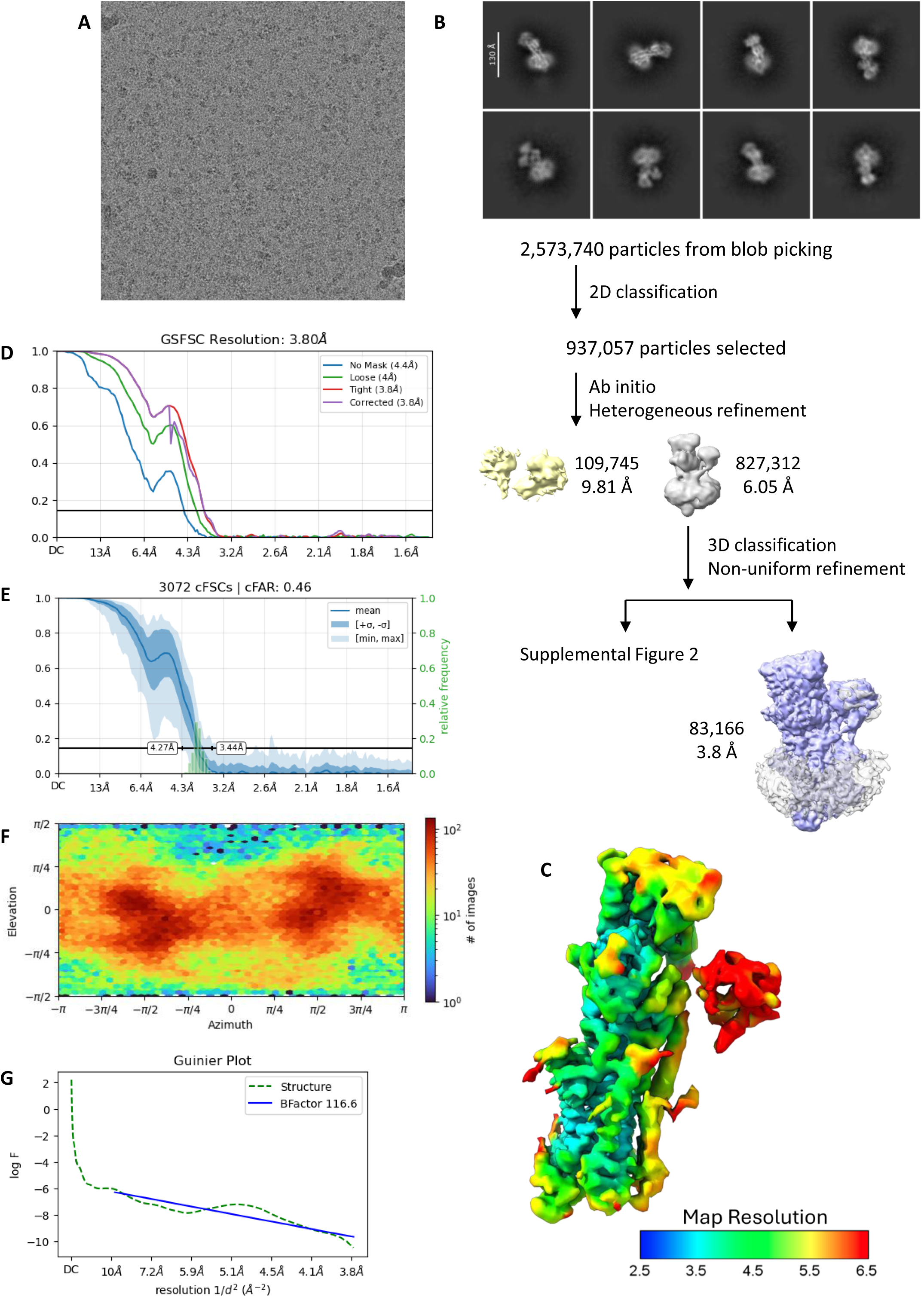
Cryo-EM workflow for apo (unbound) SERCA in the presence of UM-52. (**A**) Representative micrograph of the SERCA + UM-52 sample, low-pass filtered to 3 Å. (**B**) Summary of cryo-EM data processing, showing representative 2D classes, ab initio volumes, and the final reconstruction of a bound SERCA monomer. (**C**) Local resolution estimation of the SERCA monomer showing reasonable resolution for the transmembrane domain and lower resolution in the cytoplasmic domains. (**D**) Gold Standard Fourier Shell Correlation (GSFSC) as a function of resolution for the map shown in (C). (**E**) Conical FSC area ratio (cFAR) indicates the absence of orientation bias. (**F**) Viewing direction distribution plot for the shown map. (**G**) Guinier plot for the shown map showing the approximate B-factor.

**Supplementary Data Figure 7:**
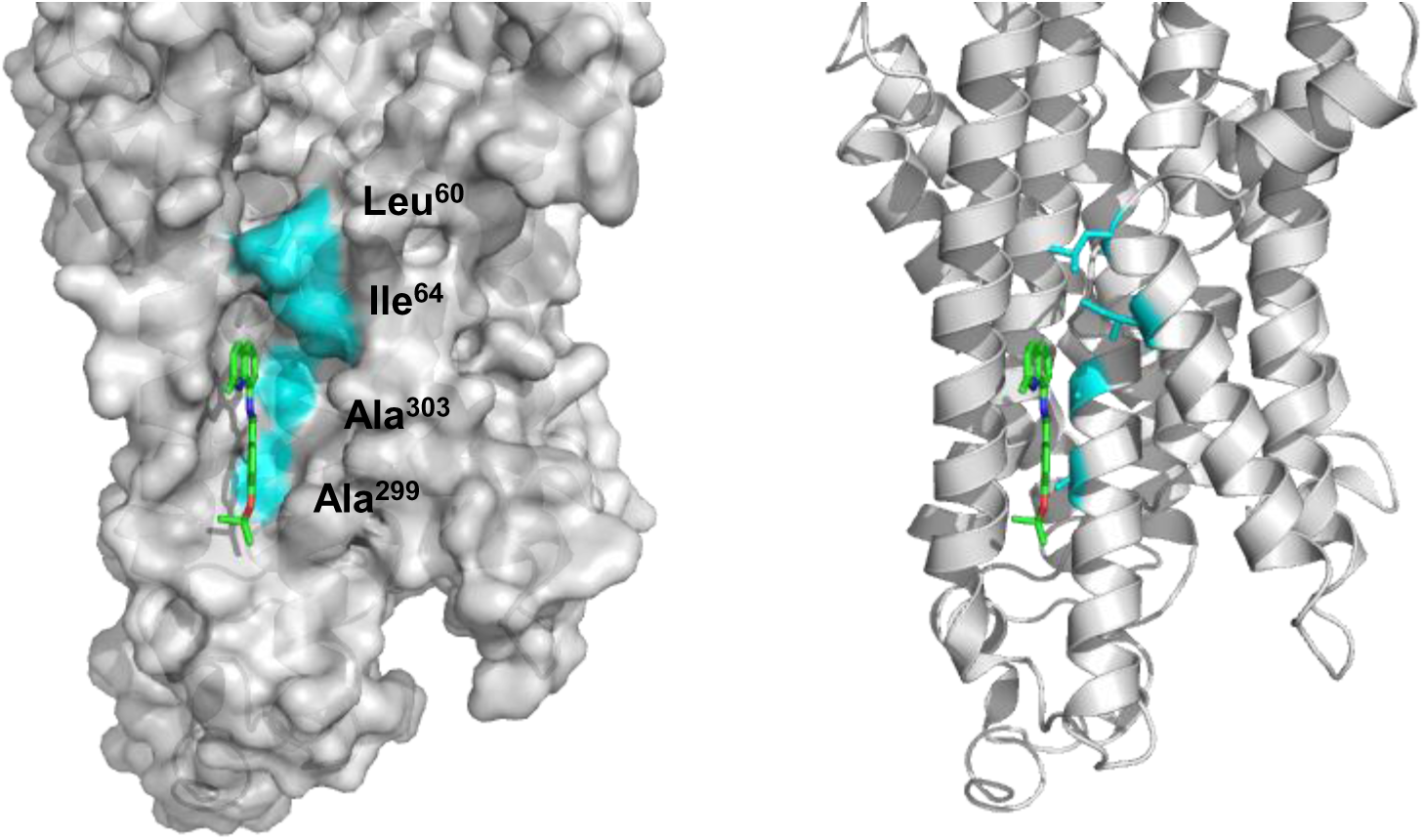
Surface and ribbon representations of SERCA bound to CDN1163 (stick model). Residues that are important for the shape of the binding cavity are indicated, including Ala^299^ and Ala^303^ that form the base of the cavity and Leu^60^ and Ile^64^ that cap the binding cavity.

**Supplementary Data Figure 8:**
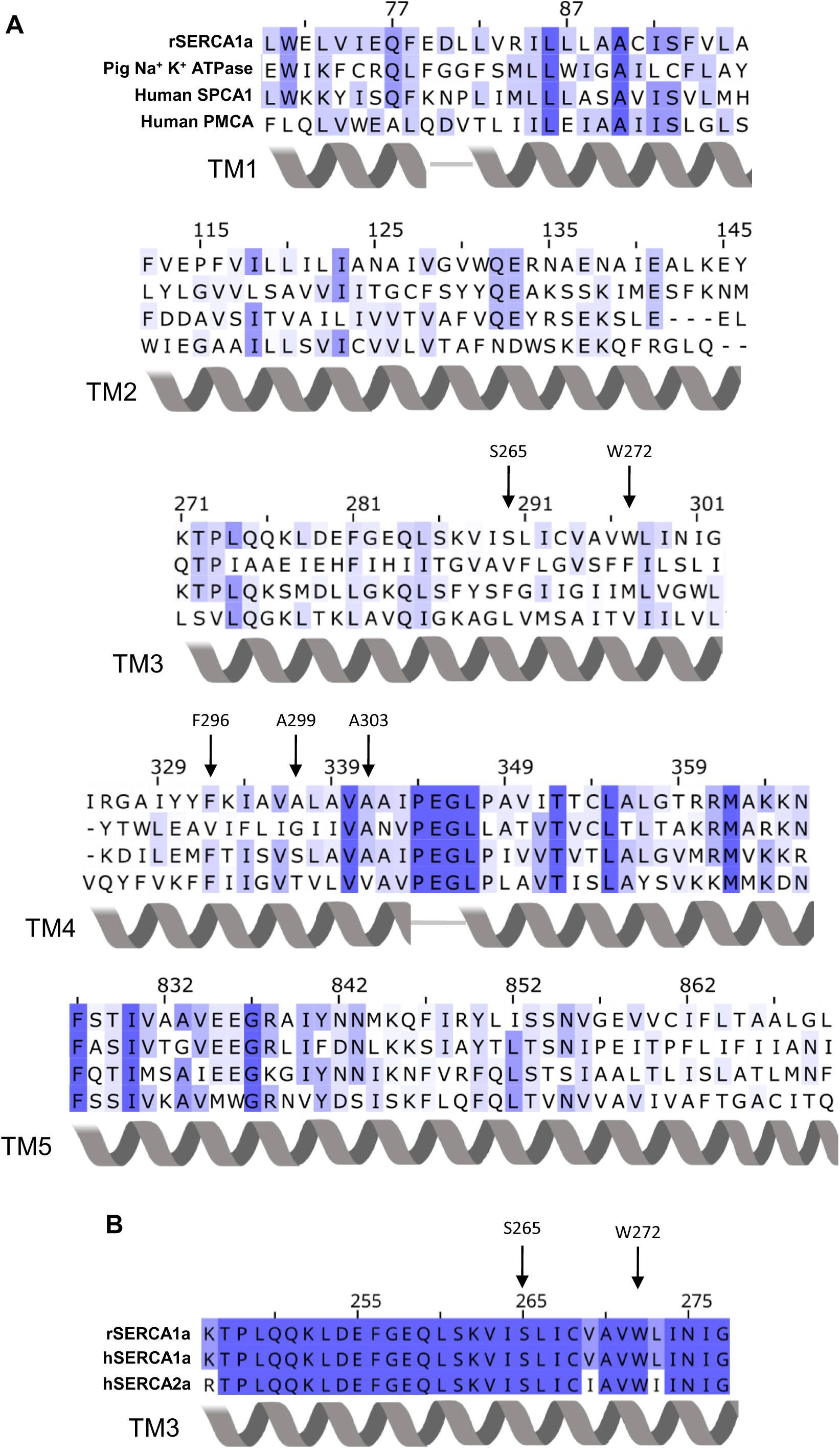
(**A**) Sequence alignments for the transmembrane helices of SERCA, NaK, PMCA2, and SPCA1 shown in Figure 2. Residues that are important for activator binding are labeled and indicated by arrows. (**B**) Sequence comparison between rabbit SERCA1, human SERCA1, and human SERCA2 for transmembrane segment M3 containing Ser^265^ and Trp^272^.

---

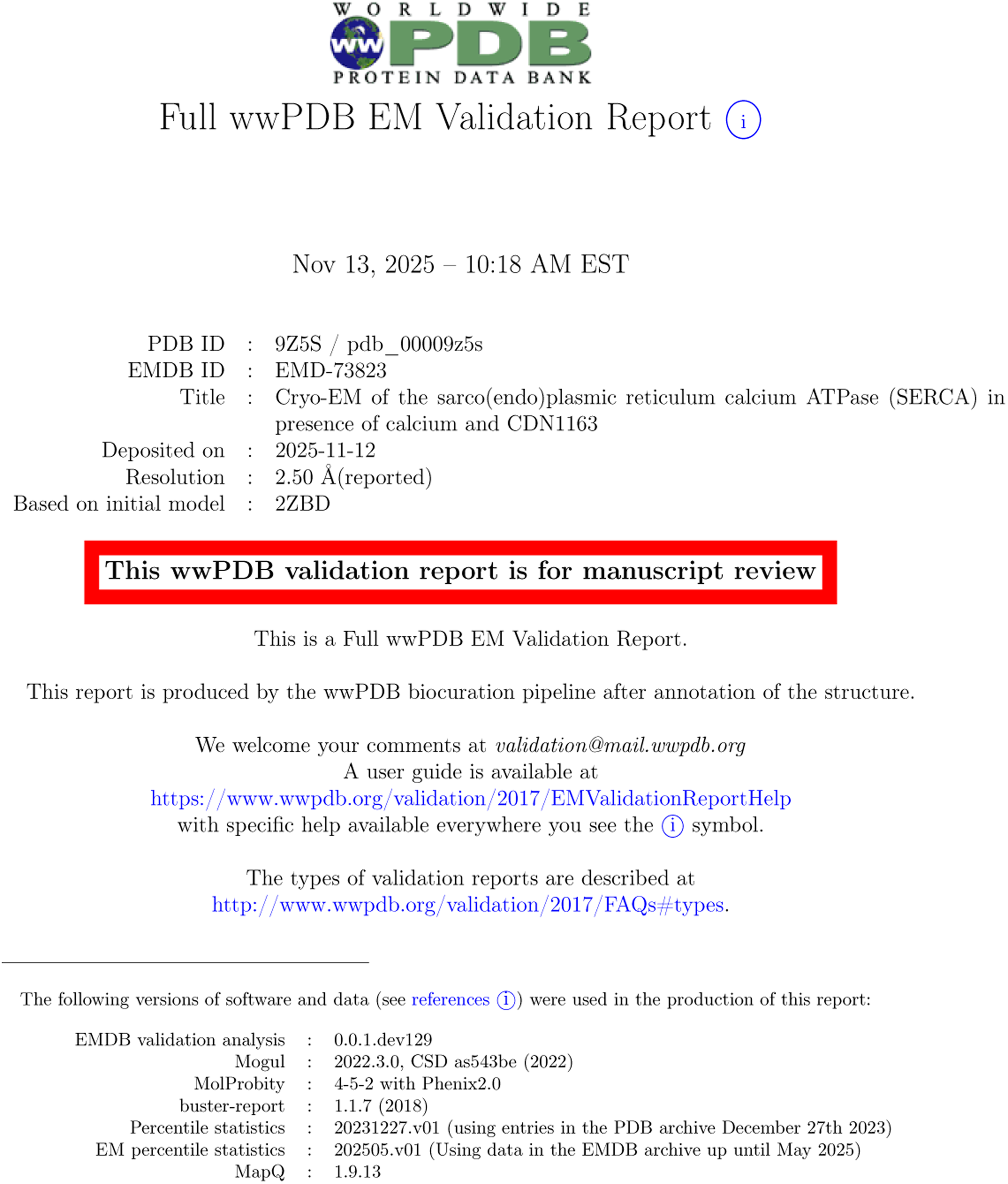

## 1 Overall quality at a glance

The following experimental techniques were used to determine the structure: *ELECTRON MICROSCOPY*

The reported resolution of this entry is 2.50 Å.

Percentile scores (ranging between 0-100) for global validation metrics of the entry are shown in the following graphic. The table shows the number of entries on which the scores are based.

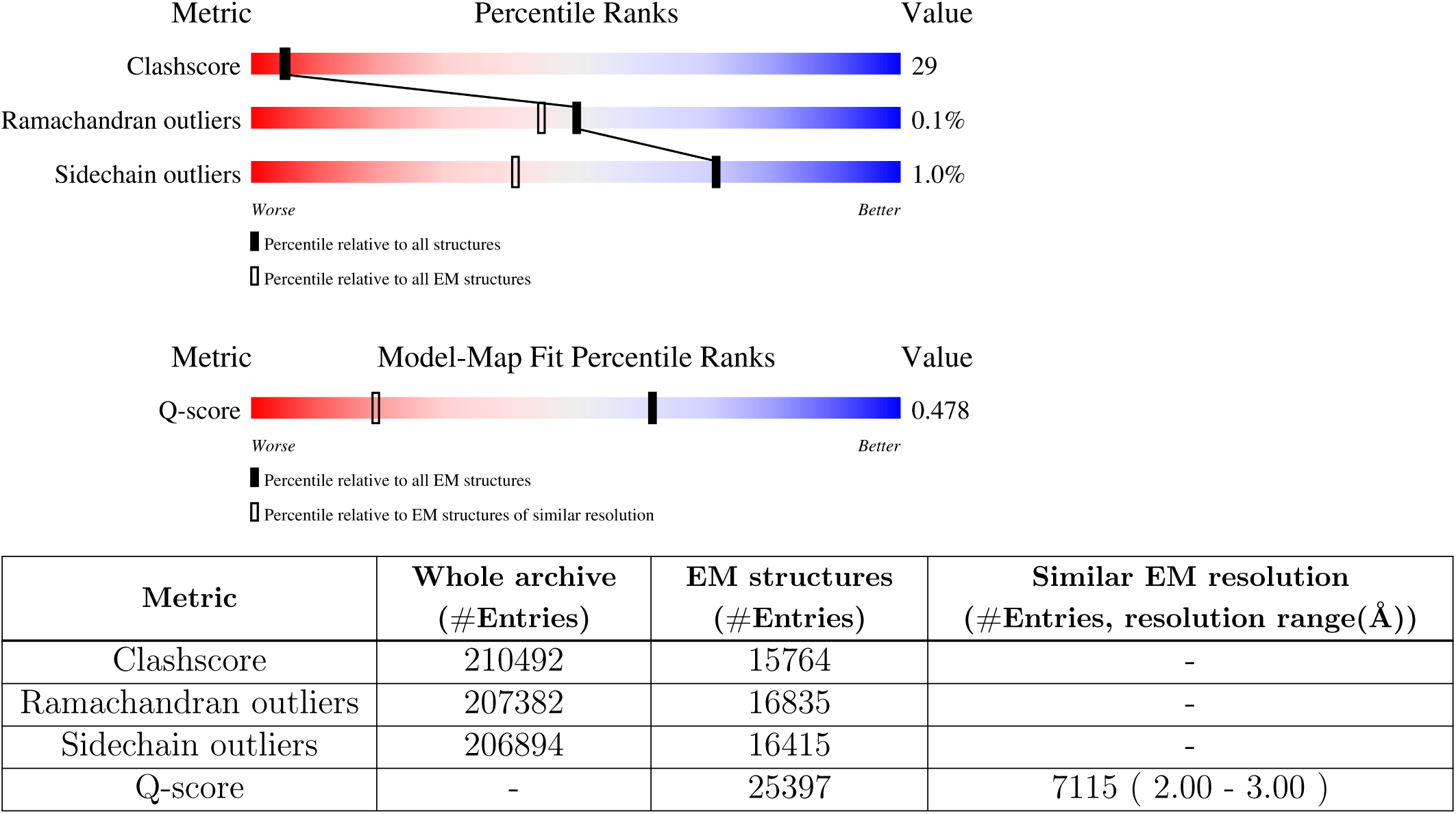

The table below summarises the geometric issues observed across the polymeric chains and their fit to the map. The red, orange, yellow and green segments of the bar indicate the fraction of residues that contain outliers for *>*=3, 2, 1 and 0 types of geometric quality criteria respectively. A grey segment represents the fraction of residues that are not modelled. The numeric value for each fraction is indicated below the corresponding segment, with a dot representing fractions *<*=5% The upper red bar (where present) indicates the fraction of residues that have poor fit to the EM map (all-atom inclusion *<* 40%). The numeric value is given above the bar.

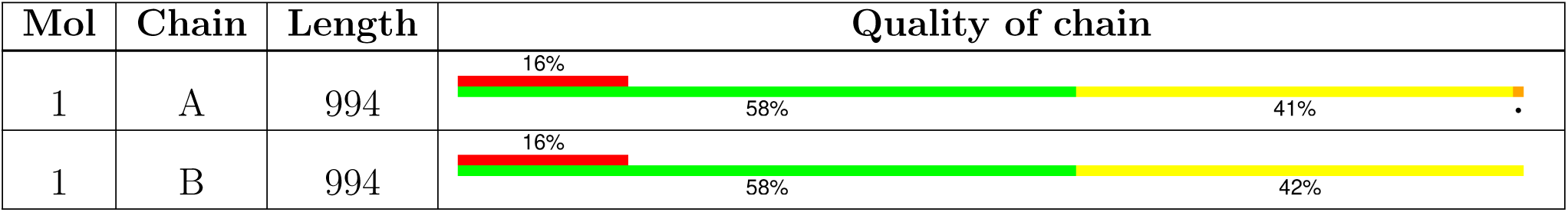

## 2 Entry composition

There are 4 unique types of molecules in this entry. The entry contains 15614 atoms, of which 156 are hydrogens and 0 are deuteriums.

In the tables below, the AltConf column contains the number of residues with at least one atom in alternate conformation and the Trace column contains the number of residues modelled with at most 2 atoms.

- Molecule 1 is a protein called Sarcoplasmic/endoplasmic reticulum calcium ATPase 1.

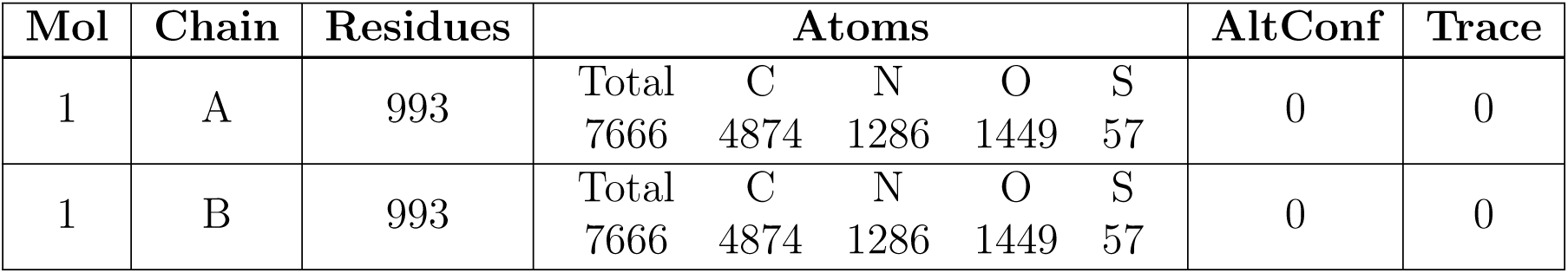

- Molecule 2 is N-(2-methylquinolin-8-yl)-4-[(propan-2-yl)oxy]benzamide (CCD ID: 9HM) (formula: C_20_H_20_N_2_O_2_) (labeled as “Ligand of Interest” by depositor).

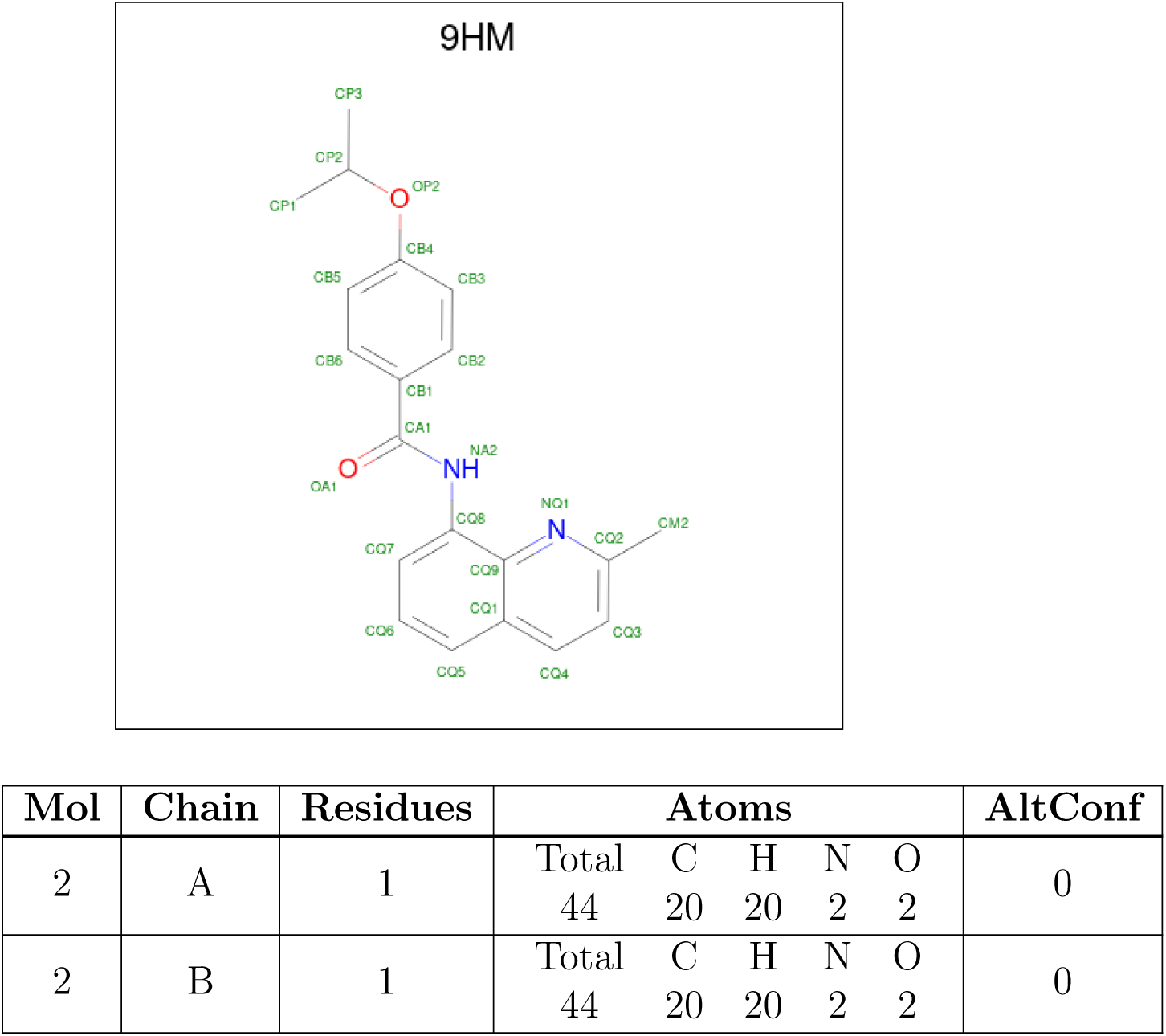

- Molecule 3 is O-DODECANYL OCTAETHYLENE GLYCOL (CCD ID: CE1) (formula: C_28_H_58_O_9_) (labeled as “Ligand of Interest” by depositor).

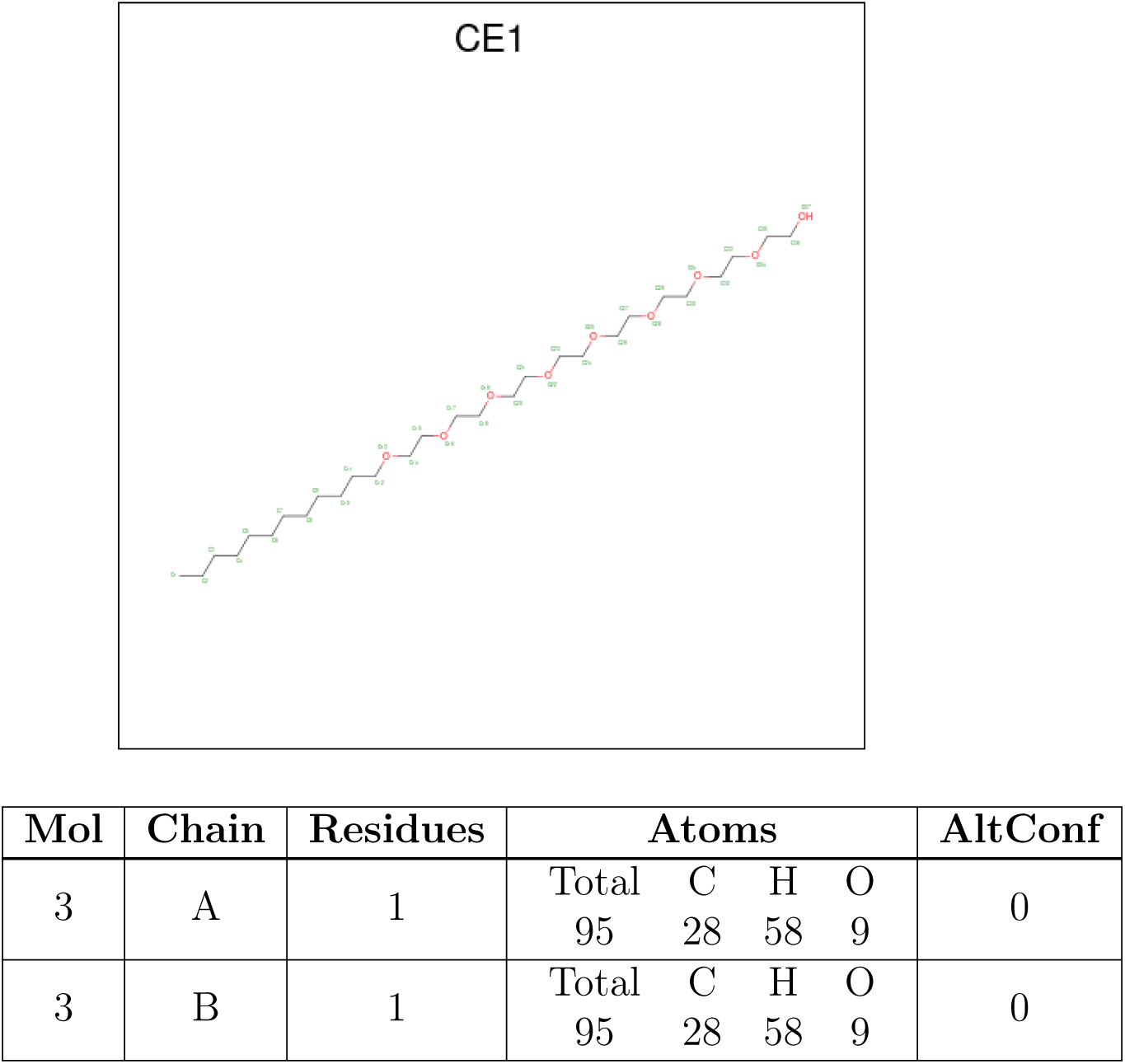

- Molecule 4 is CALCIUM ION (CCD ID: CA) (formula: Ca) (labeled as “Ligand of Interest” by depositor).

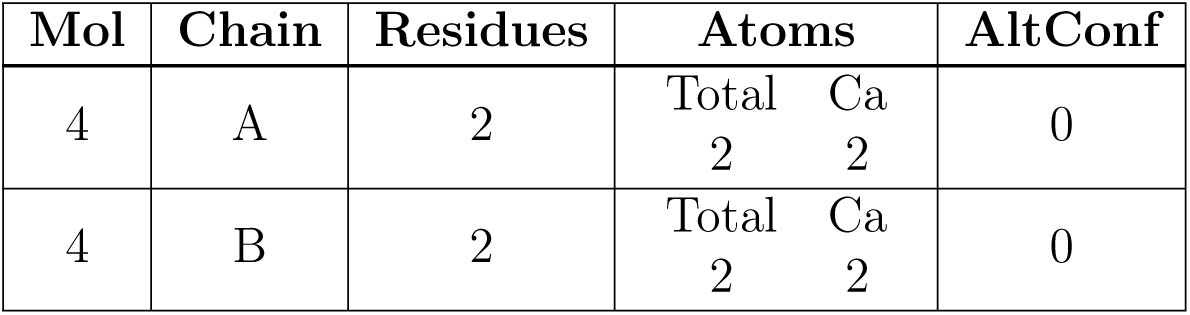

## 3 Residue-property plots

These plots are drawn for all protein, RNA, DNA and oligosaccharide chains in the entry. The first graphic for a chain summarises the proportions of the various outlier classes displayed in the second graphic. The second graphic shows the sequence view annotated by issues in geometry and atom inclusion in map density. Residues are color-coded according to the number of geometric quality criteria for which they contain at least one outlier: green = 0, yellow = 1, orange = 2 and red = 3 or more. A red diamond above a residue indicates a poor fit to the EM map for this residue (all-atom inclusion < 40%). Stretches of 2 or more consecutive residues without any outlier are shown as a green connector. Residues present in the sample, but not in the model, are shown in grey.

- Molecule 1: Sarcoplasmic/endoplasmic reticulum calcium ATPase 1

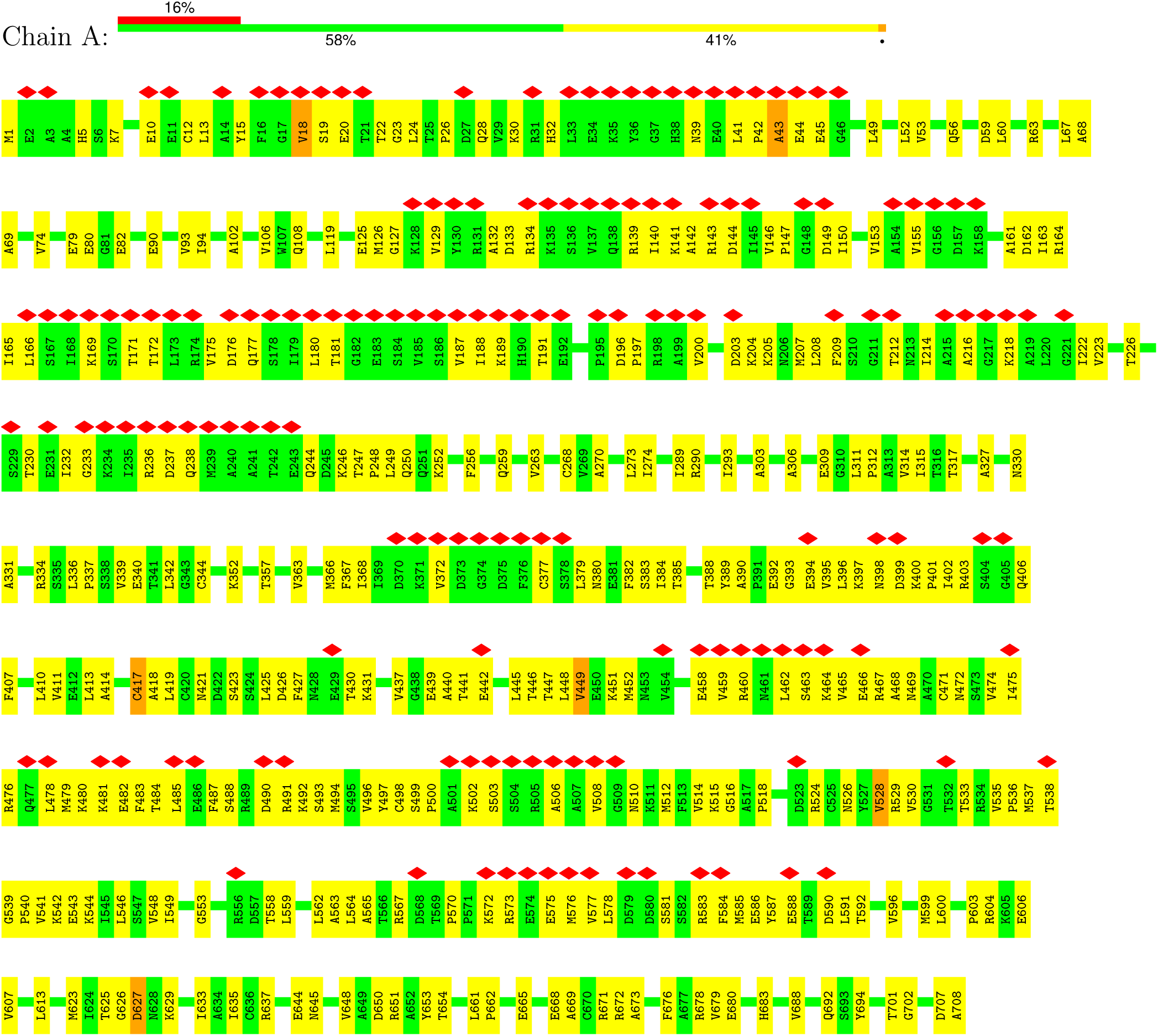
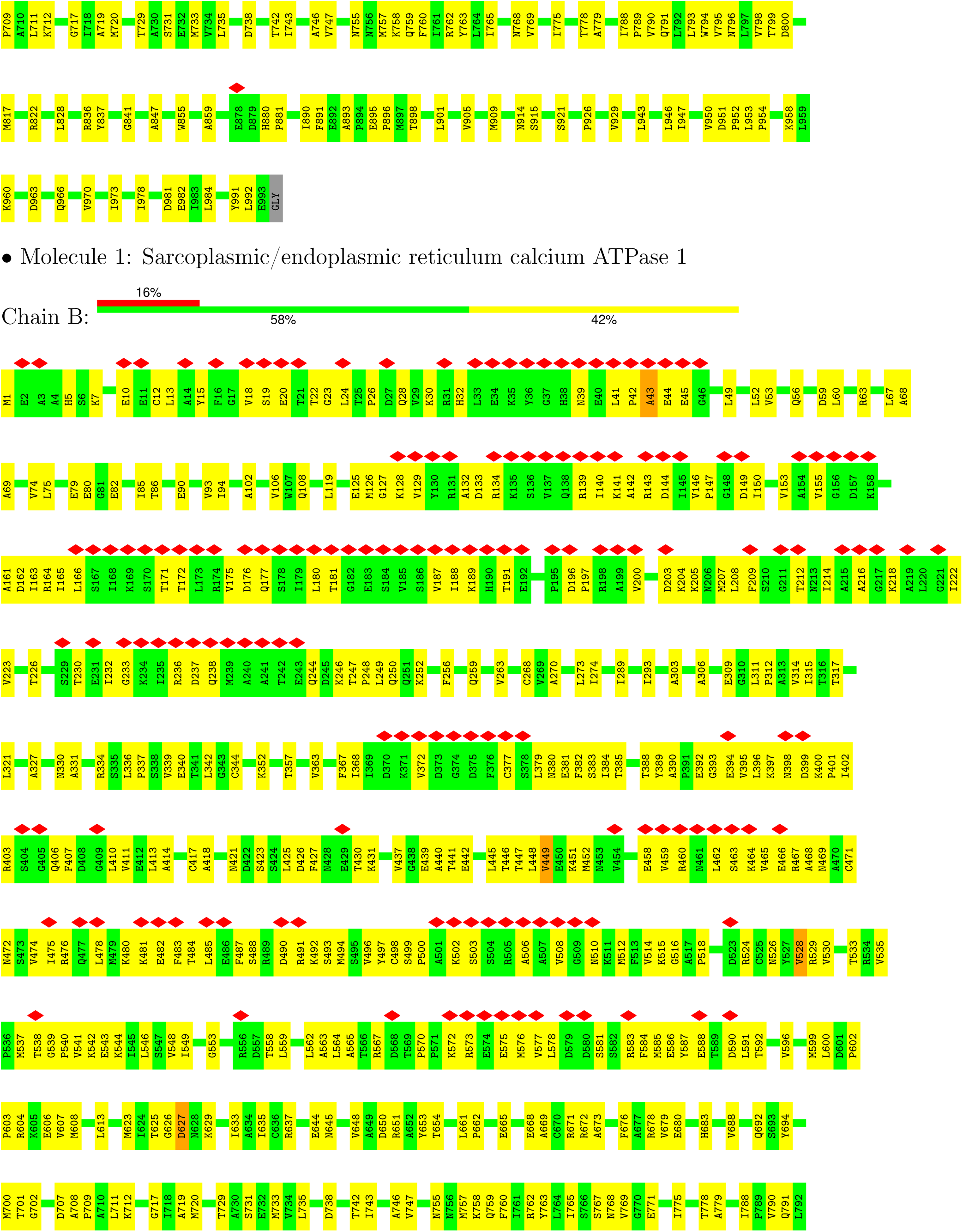
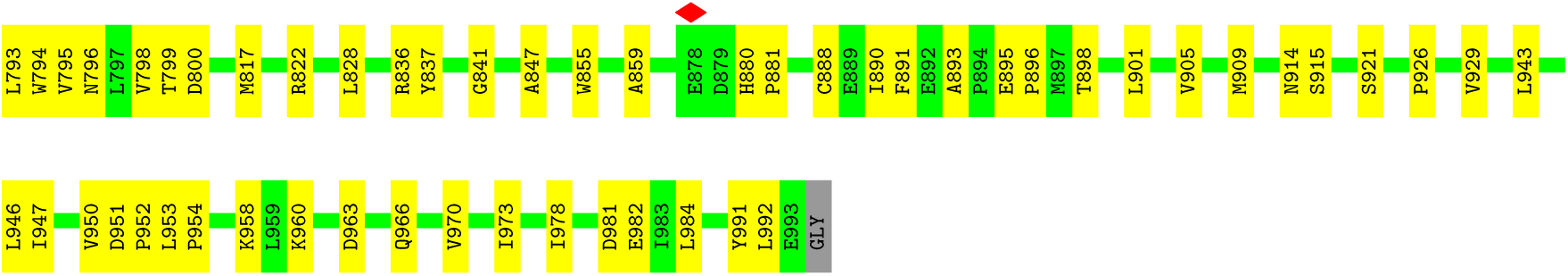

## Experimental information

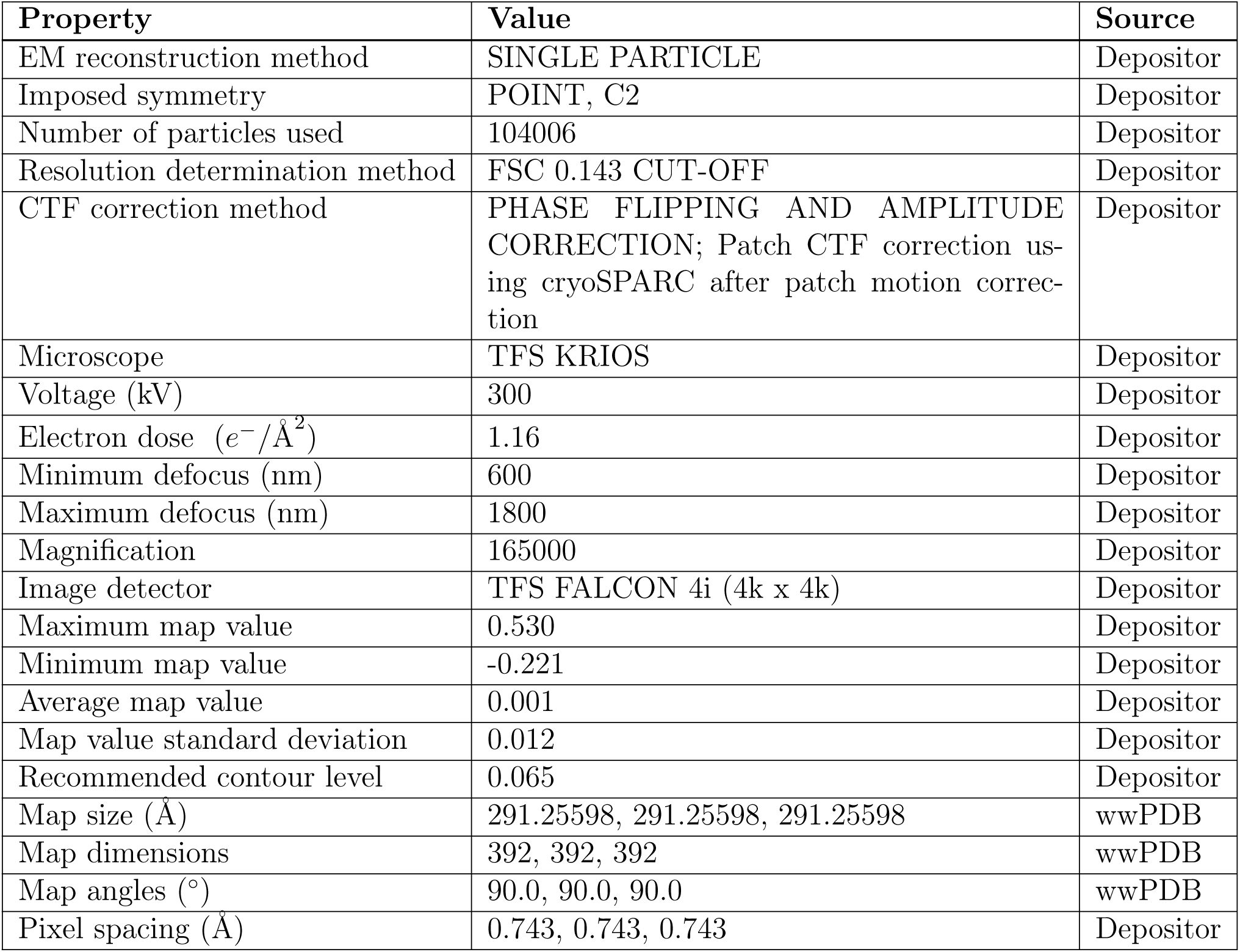

## 5 Model quality

### 5.1 Standard geometry

Bond lengths and bond angles in the following residue types are not validated in this section: CA, CE1, 9HM

The Z score for a bond length (or angle) is the number of standard deviations the observed value is removed from the expected value. A bond length (or angle) with *|Z| >* 5 is considered an outlier worth inspection. RMSZ is the root-mean-square of all Z scores of the bond lengths (or angles).

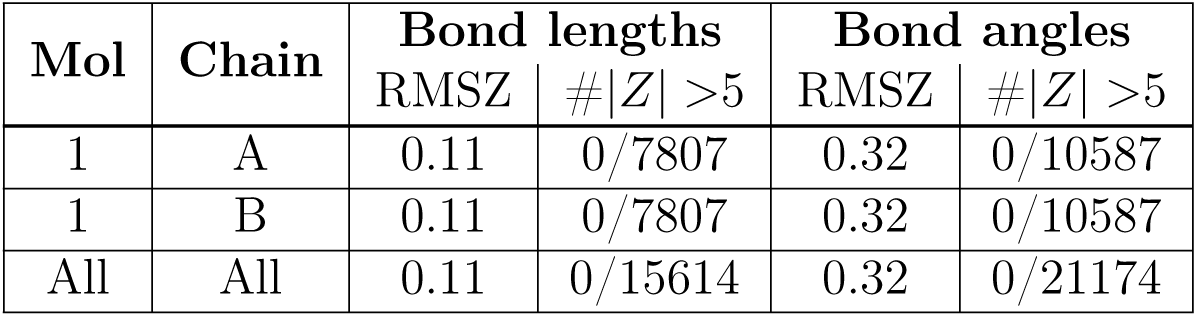

There are no bond length outliers.

There are no bond angle outliers.

There are no chirality outliers.

There are no planarity outliers.

### 5.2 Too-close contacts

In the following table, the Non-H and H(model) columns list the number of non-hydrogen atoms and hydrogen atoms in the chain respectively. The H(added) column lists the number of hydrogen atoms added and optimized by MolProbity. The Clashes column lists the number of clashes within the asymmetric unit, whereas Symm-Clashes lists symmetry-related clashes.

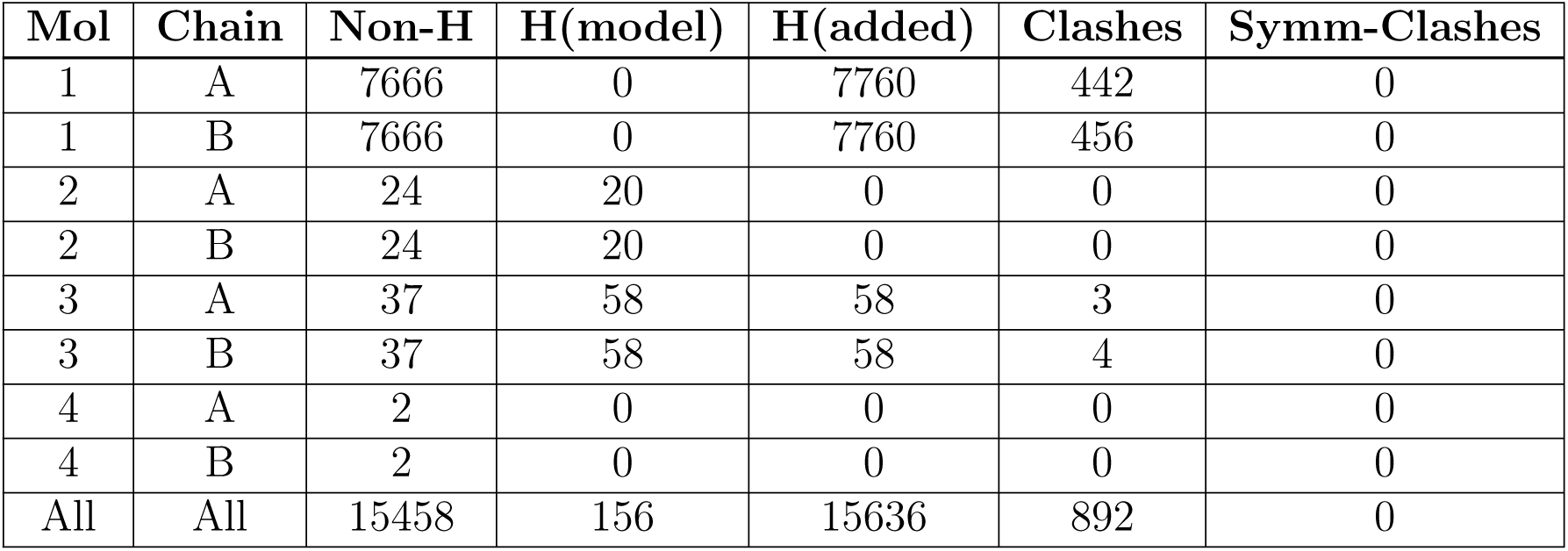

The all-atom clashscore is defined as the number of clashes found per 1000 atoms (including hydrogen atoms). The all-atom clashscore for this structure is 29.

All (892) close contacts within the same asymmetric unit are listed below, sorted by their clash magnitude.

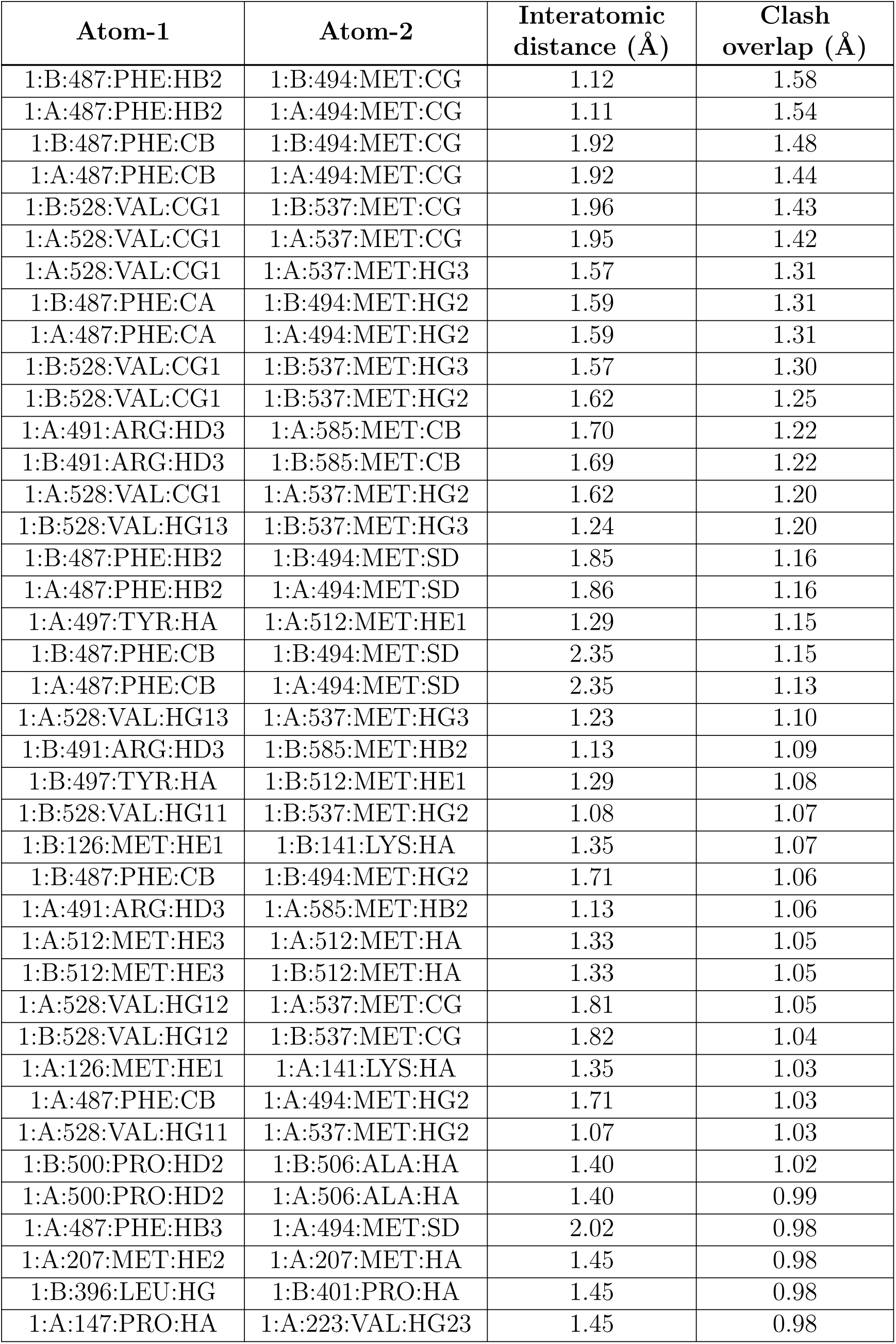
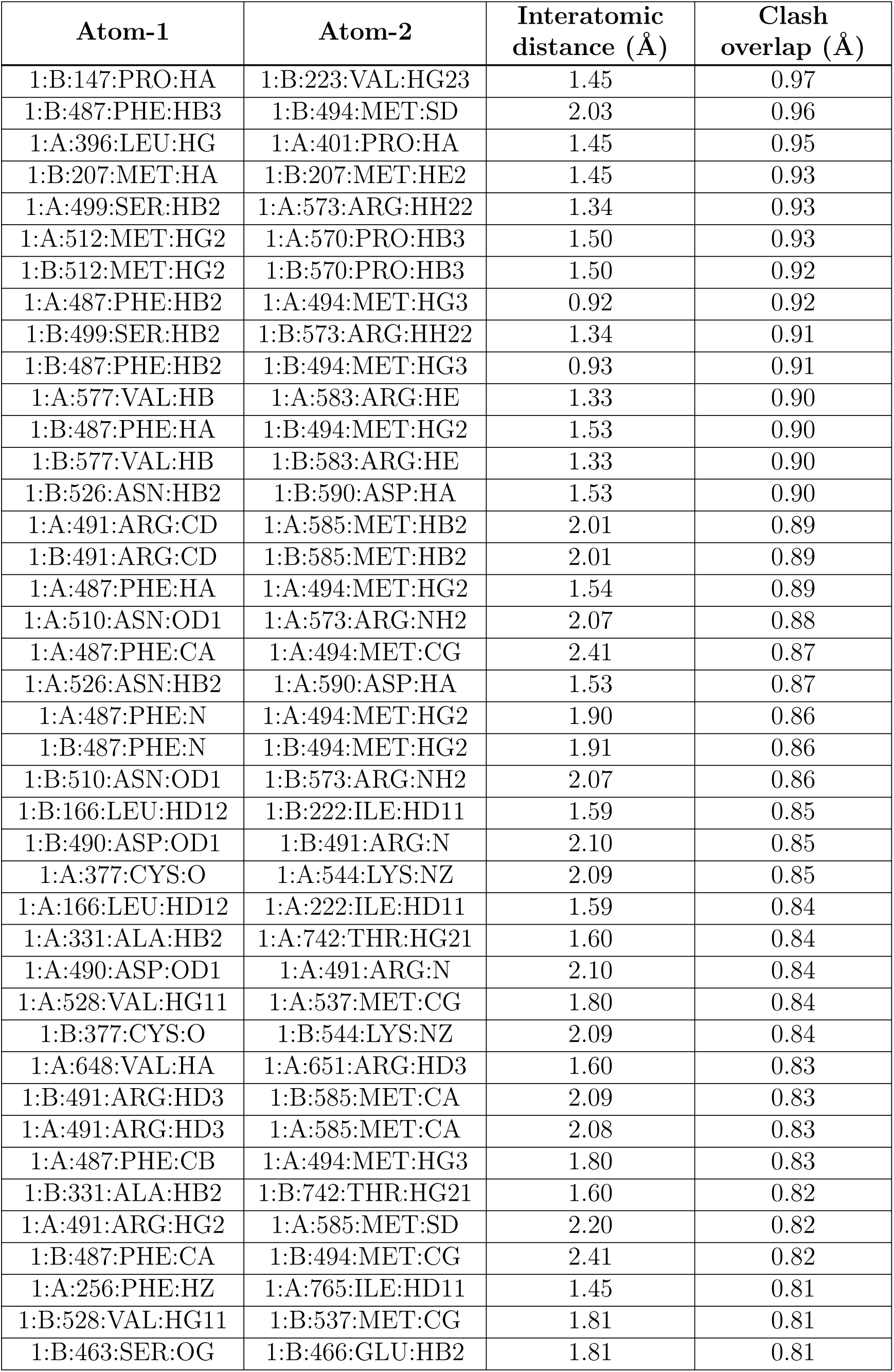
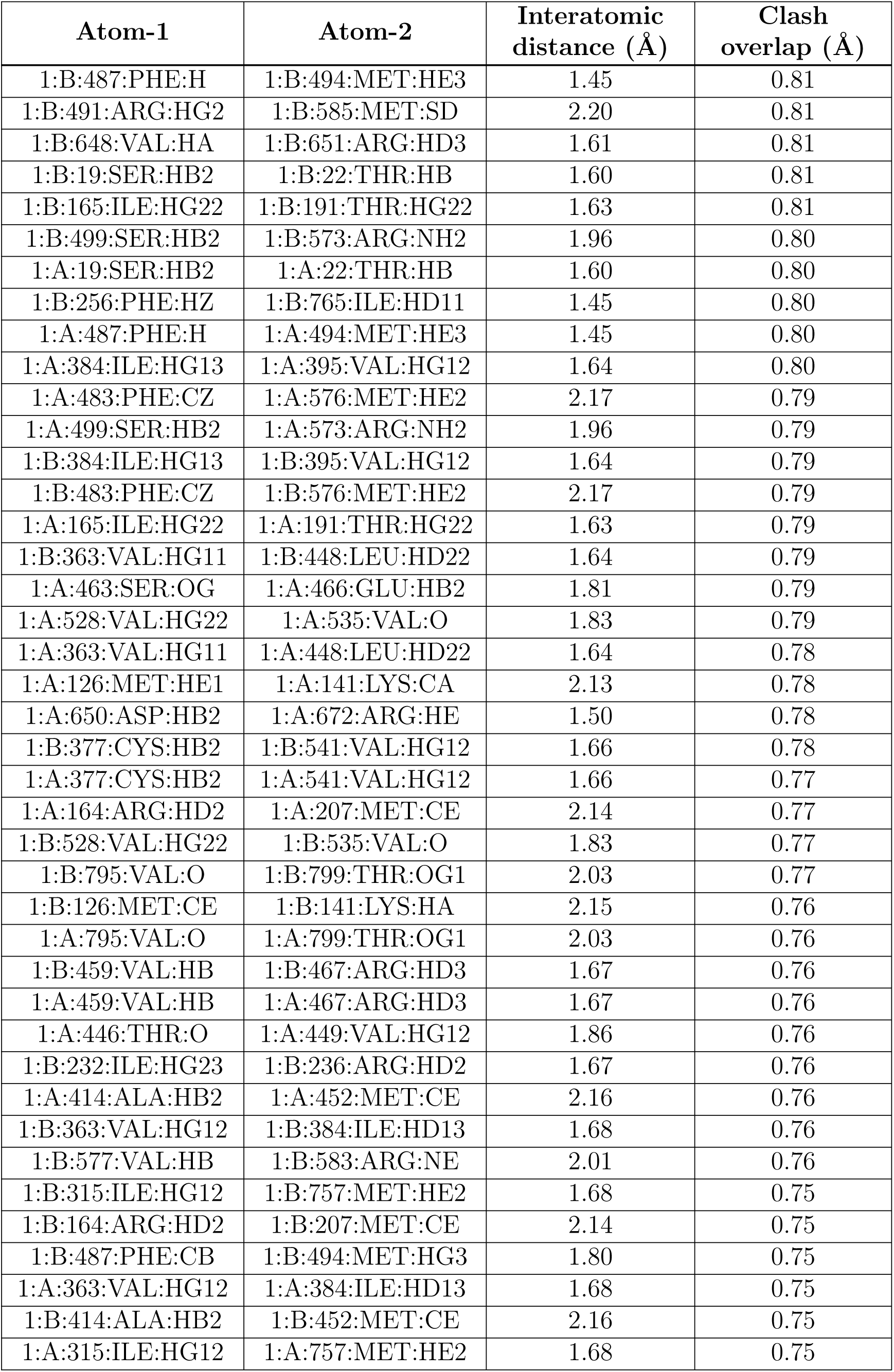
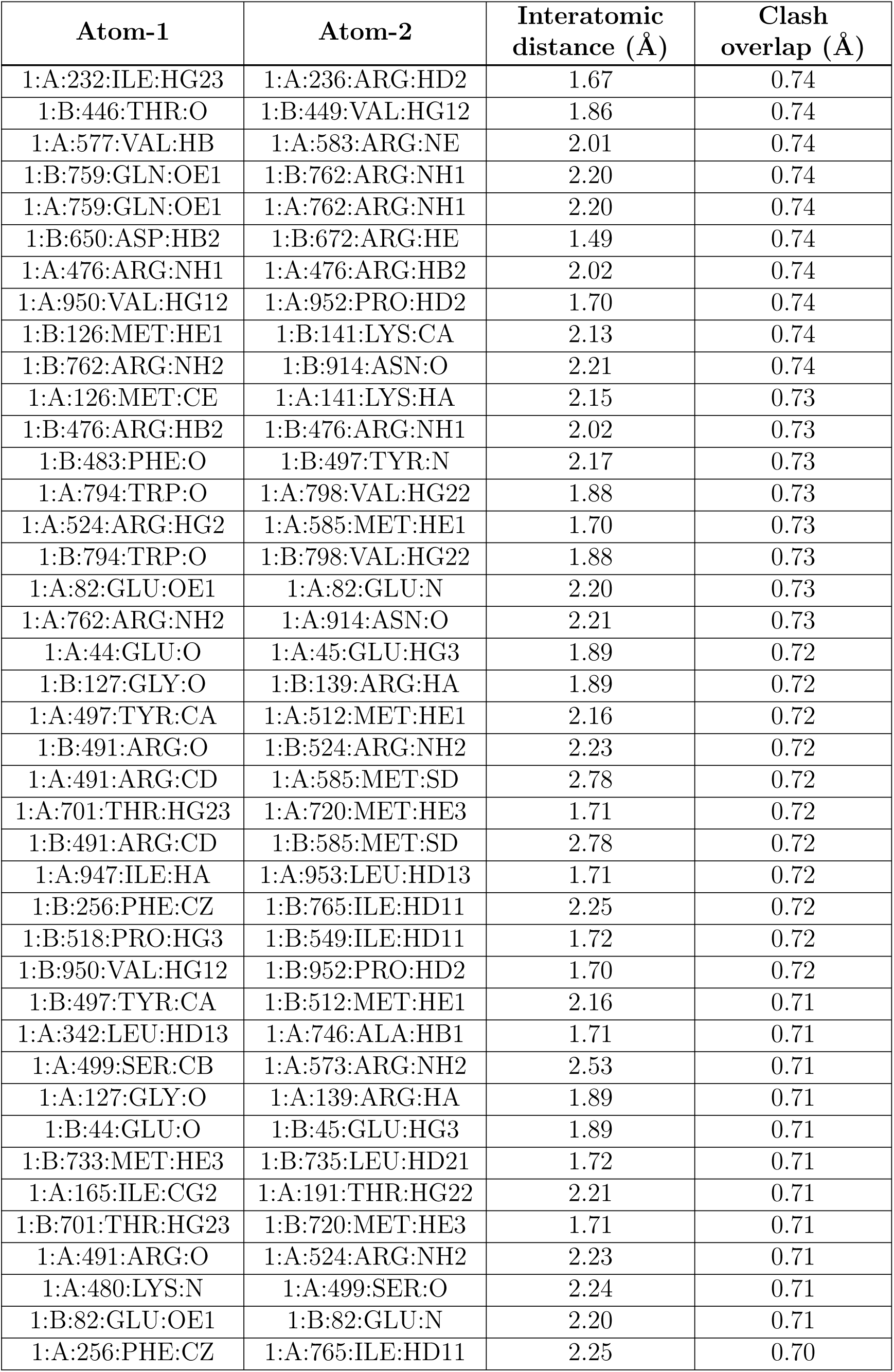
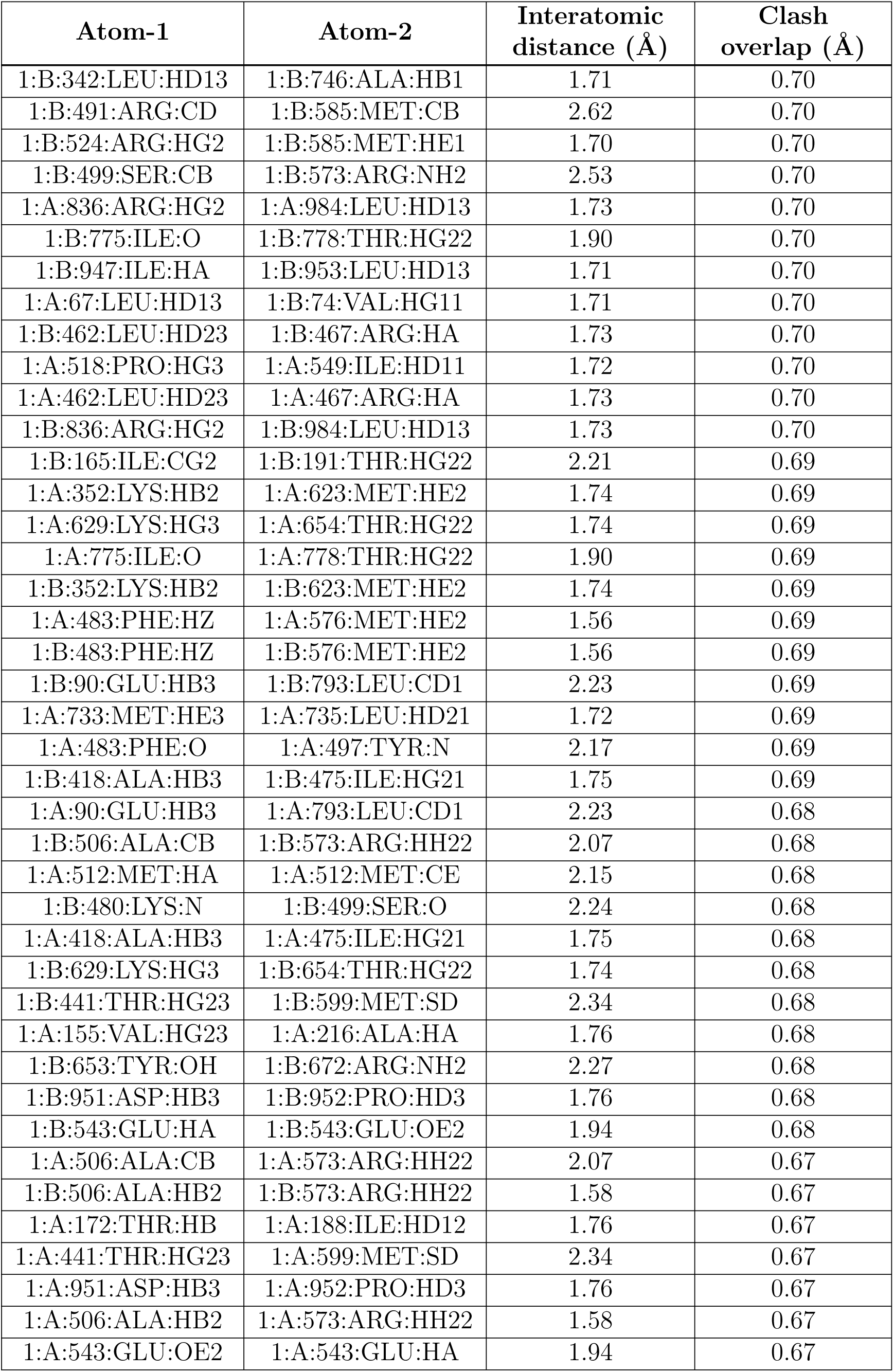
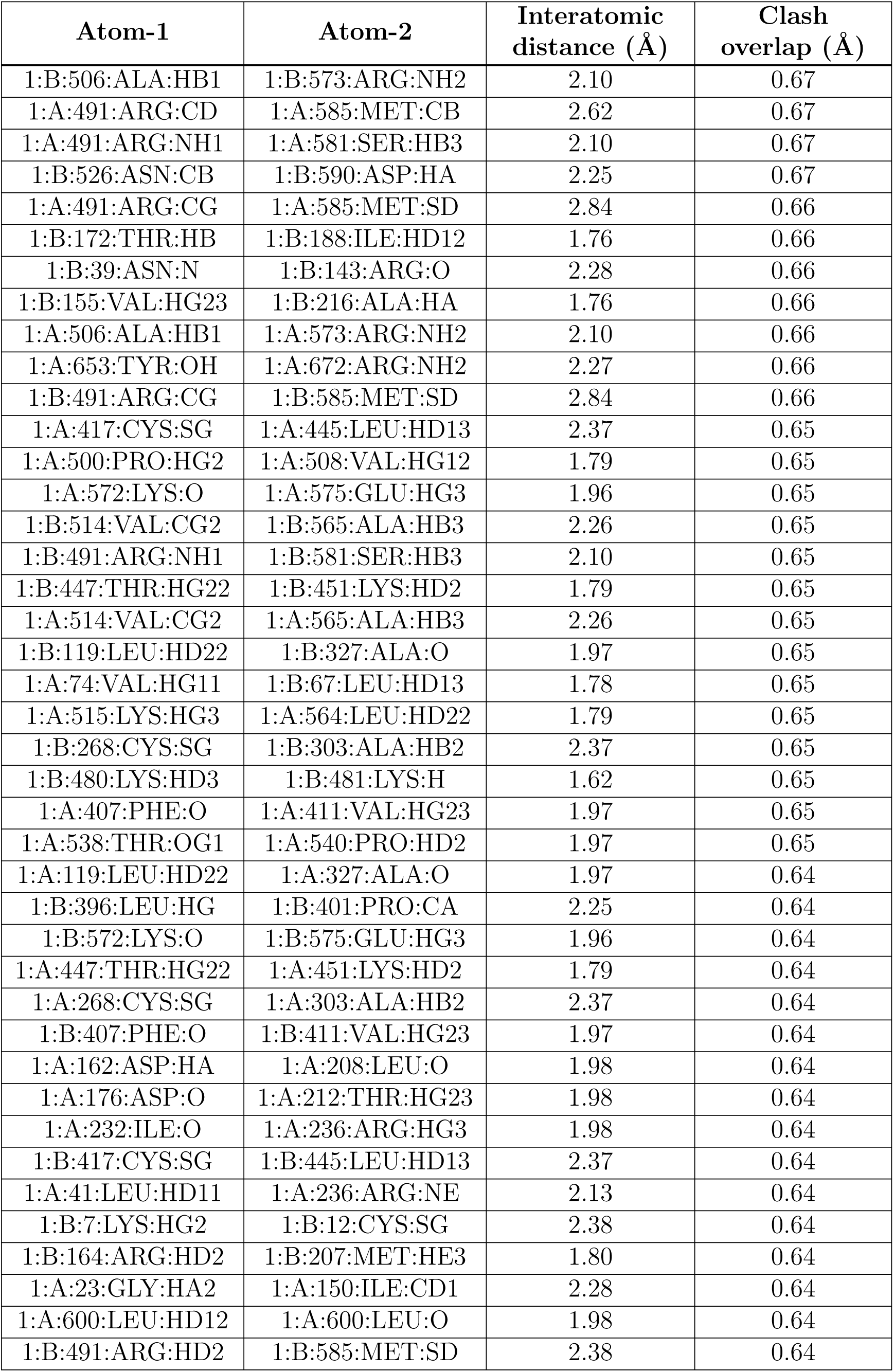
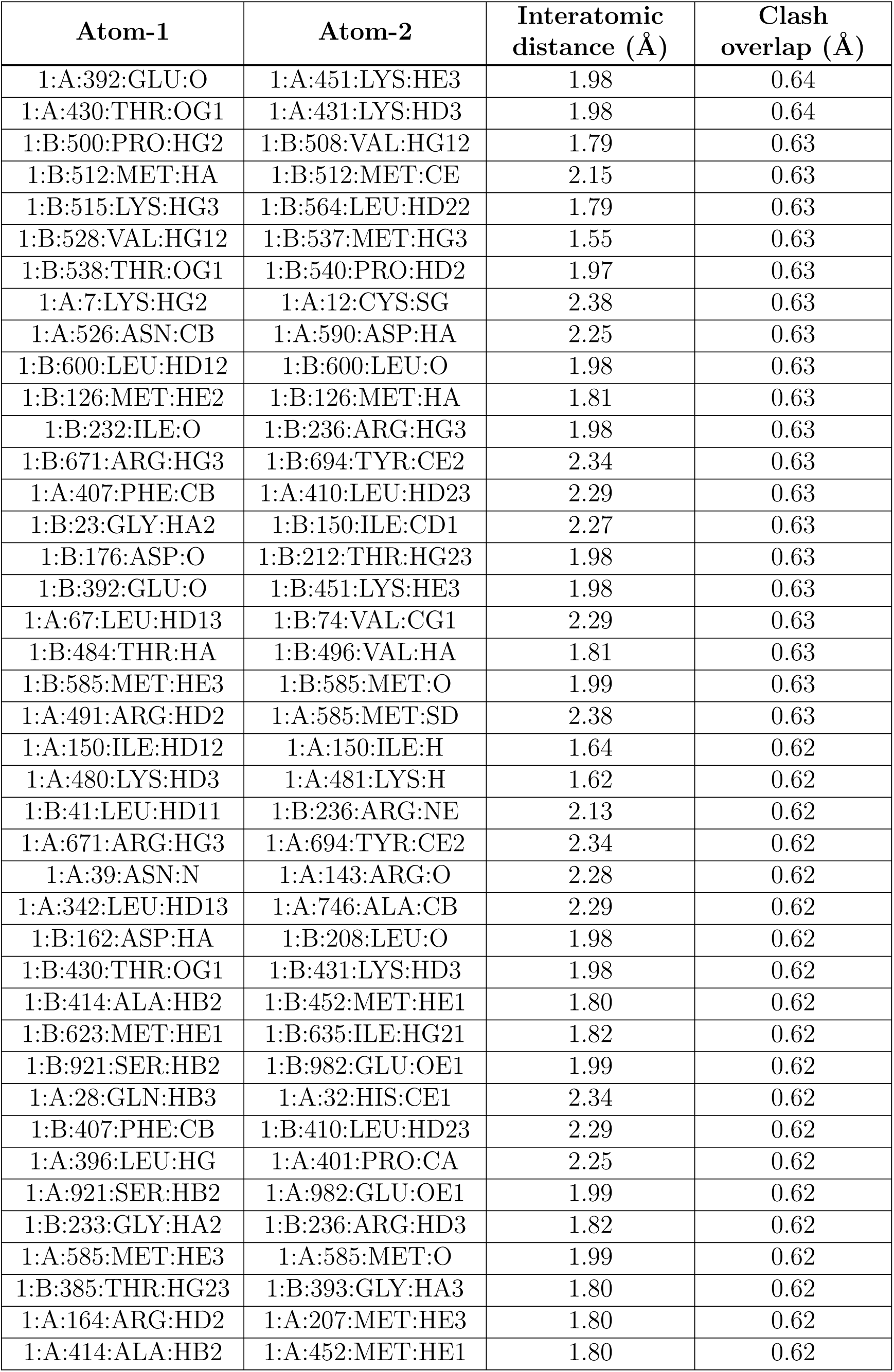
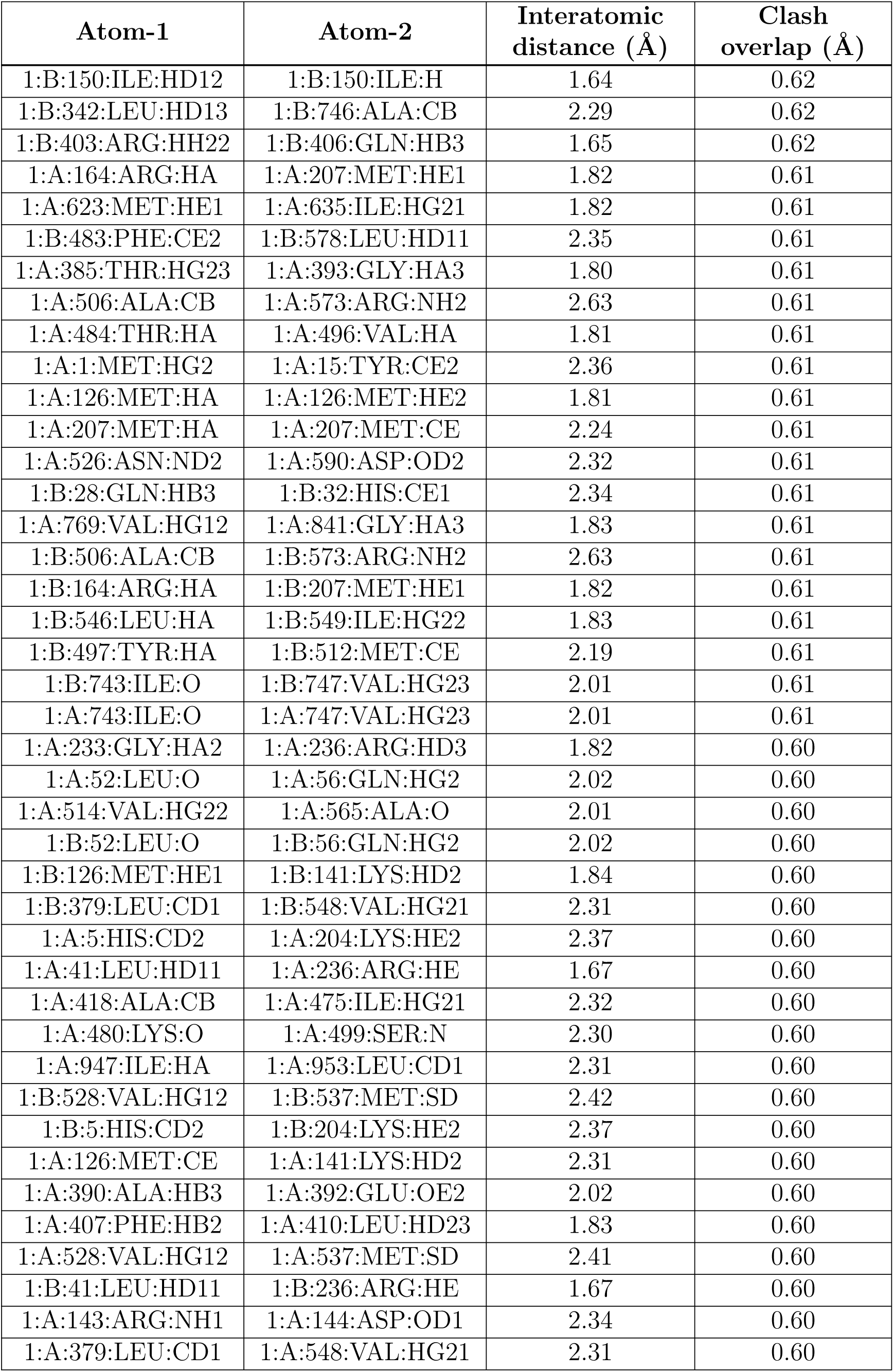
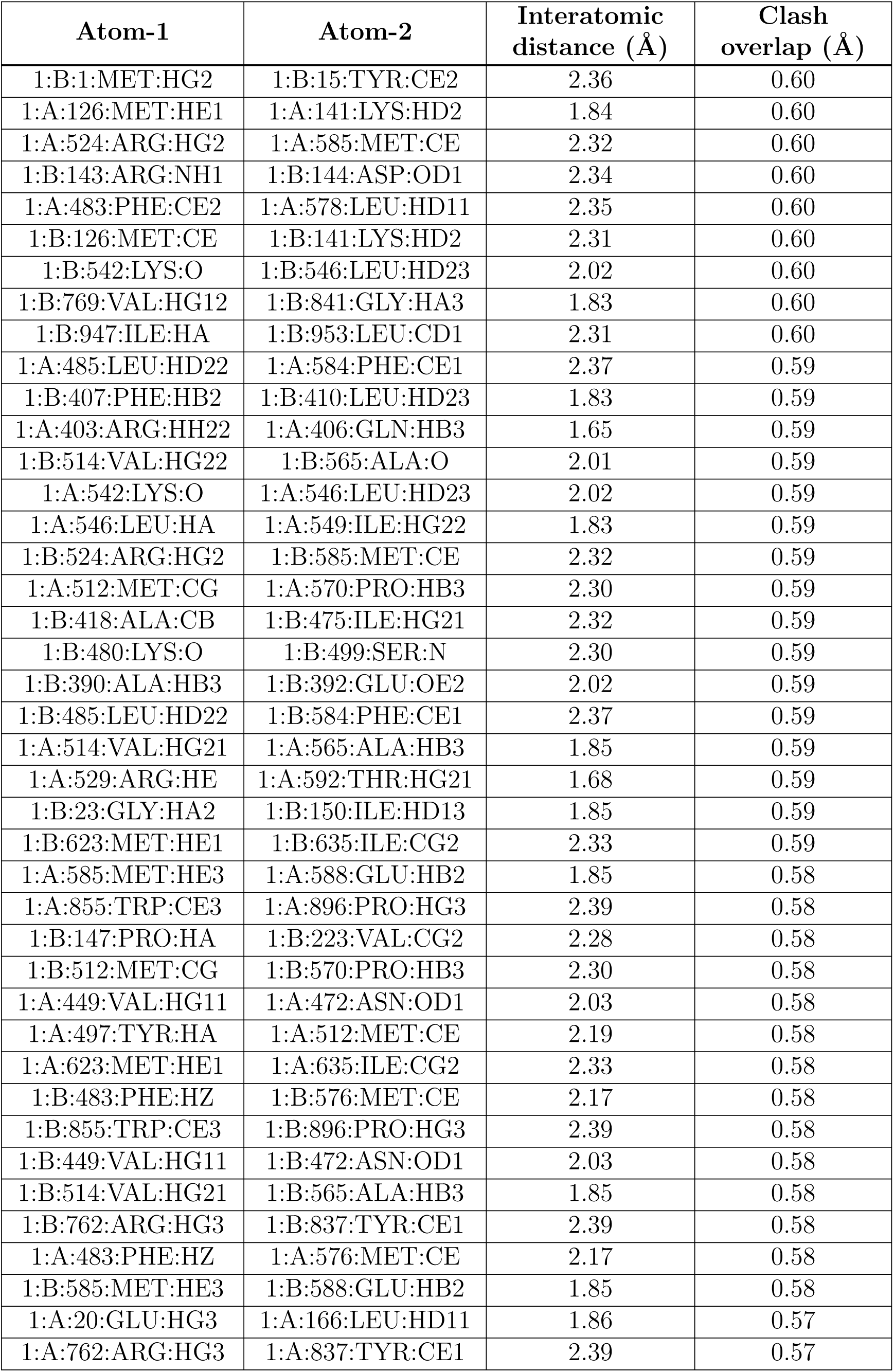
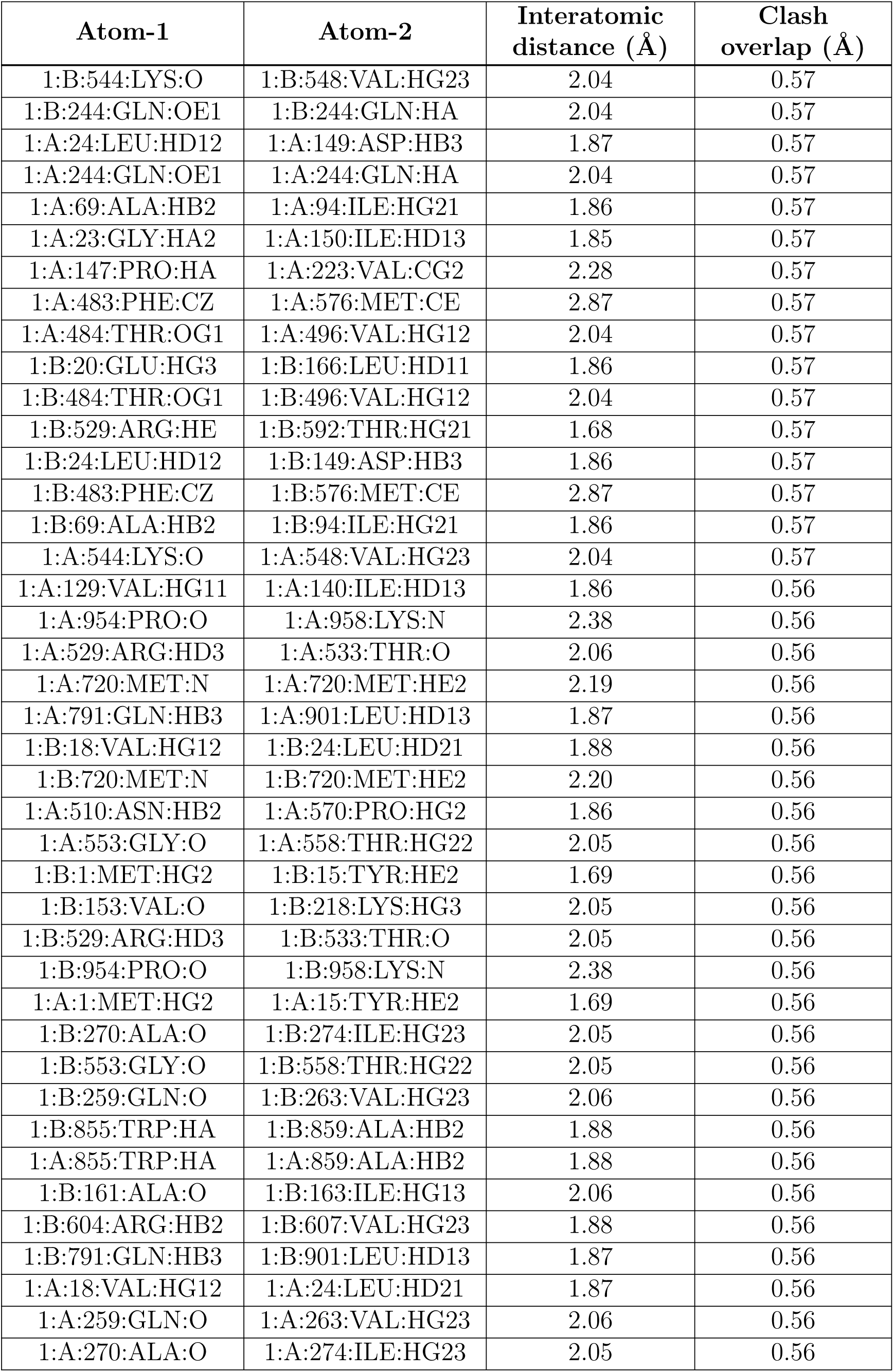
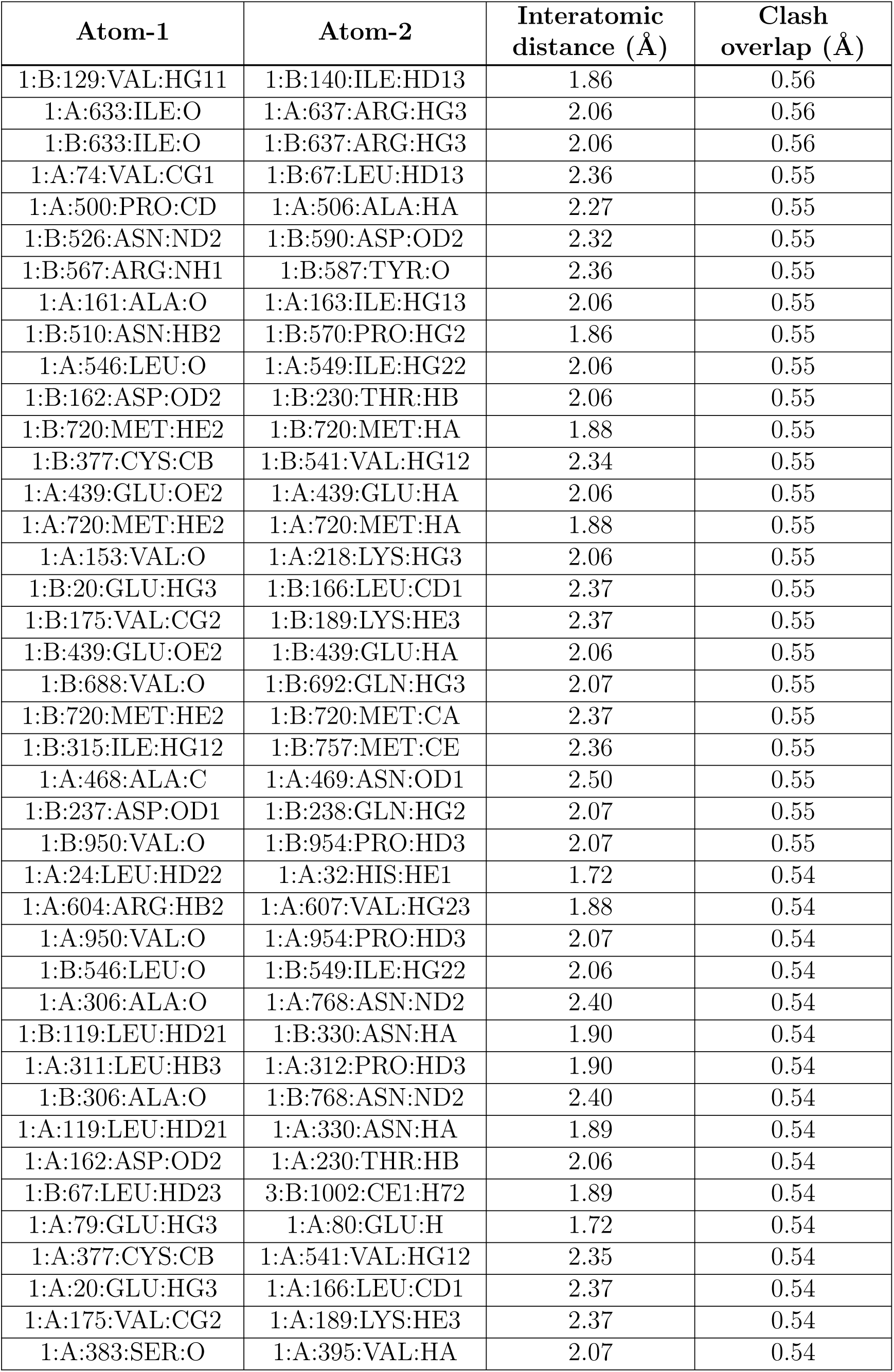
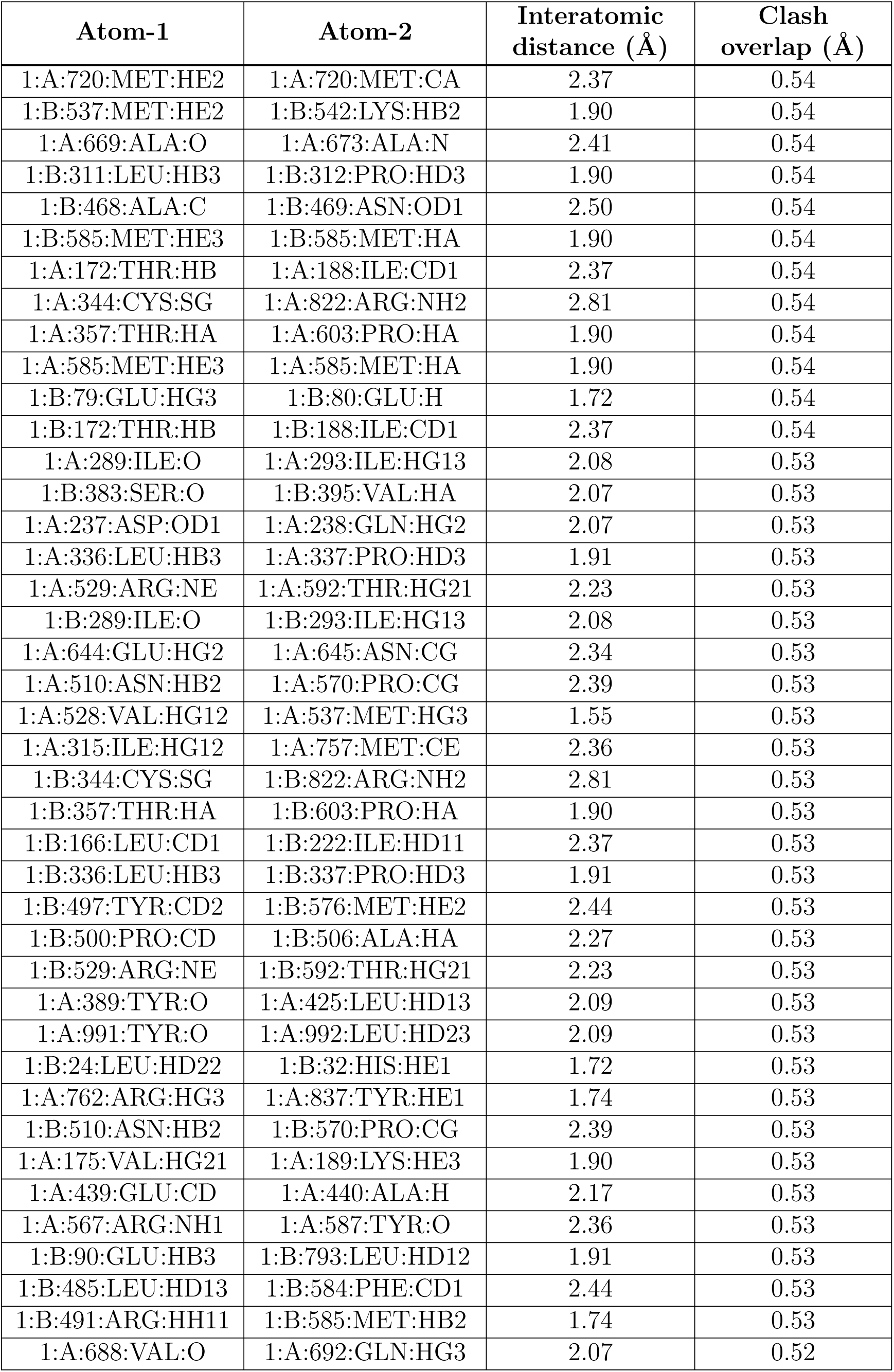
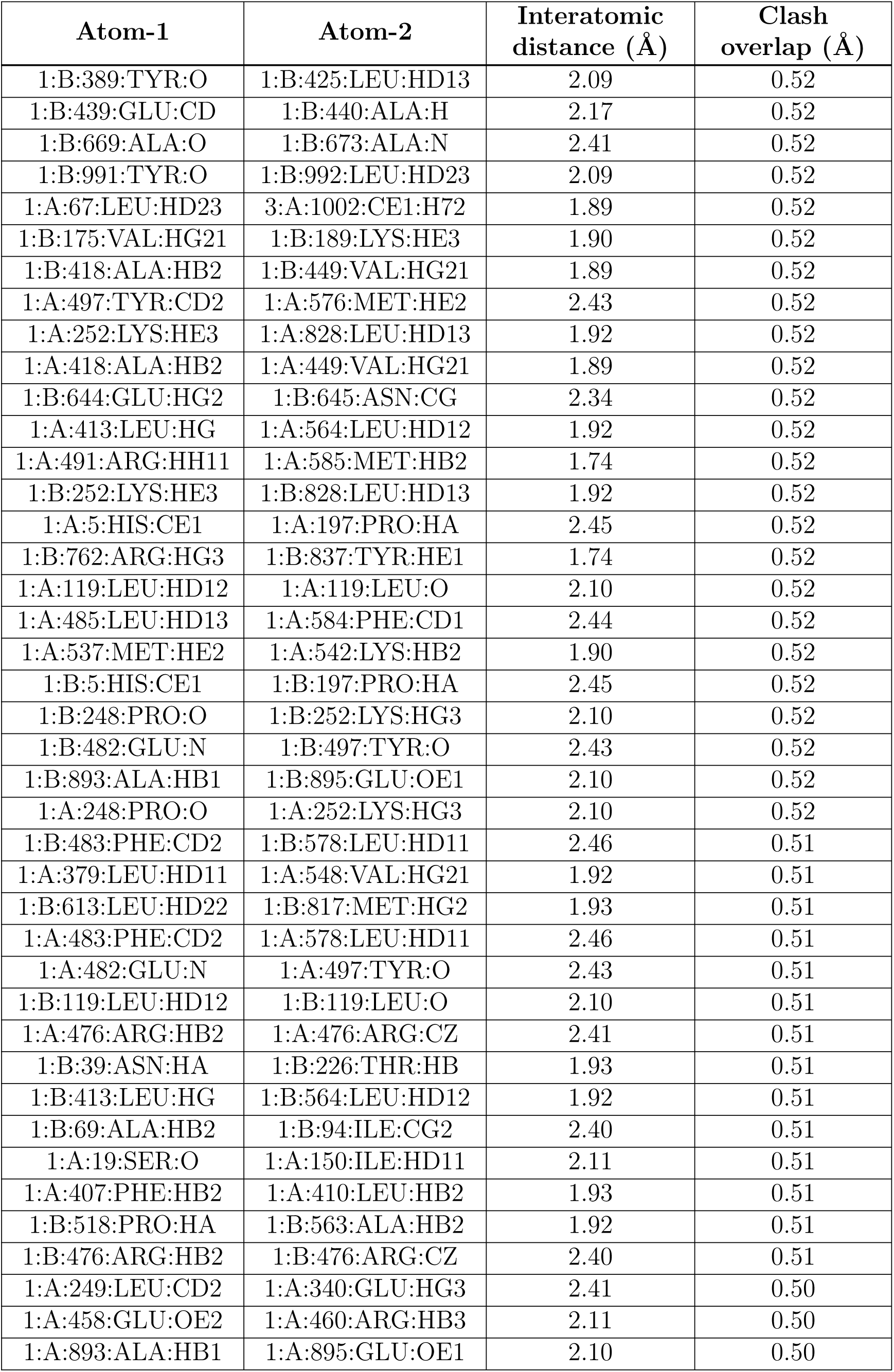
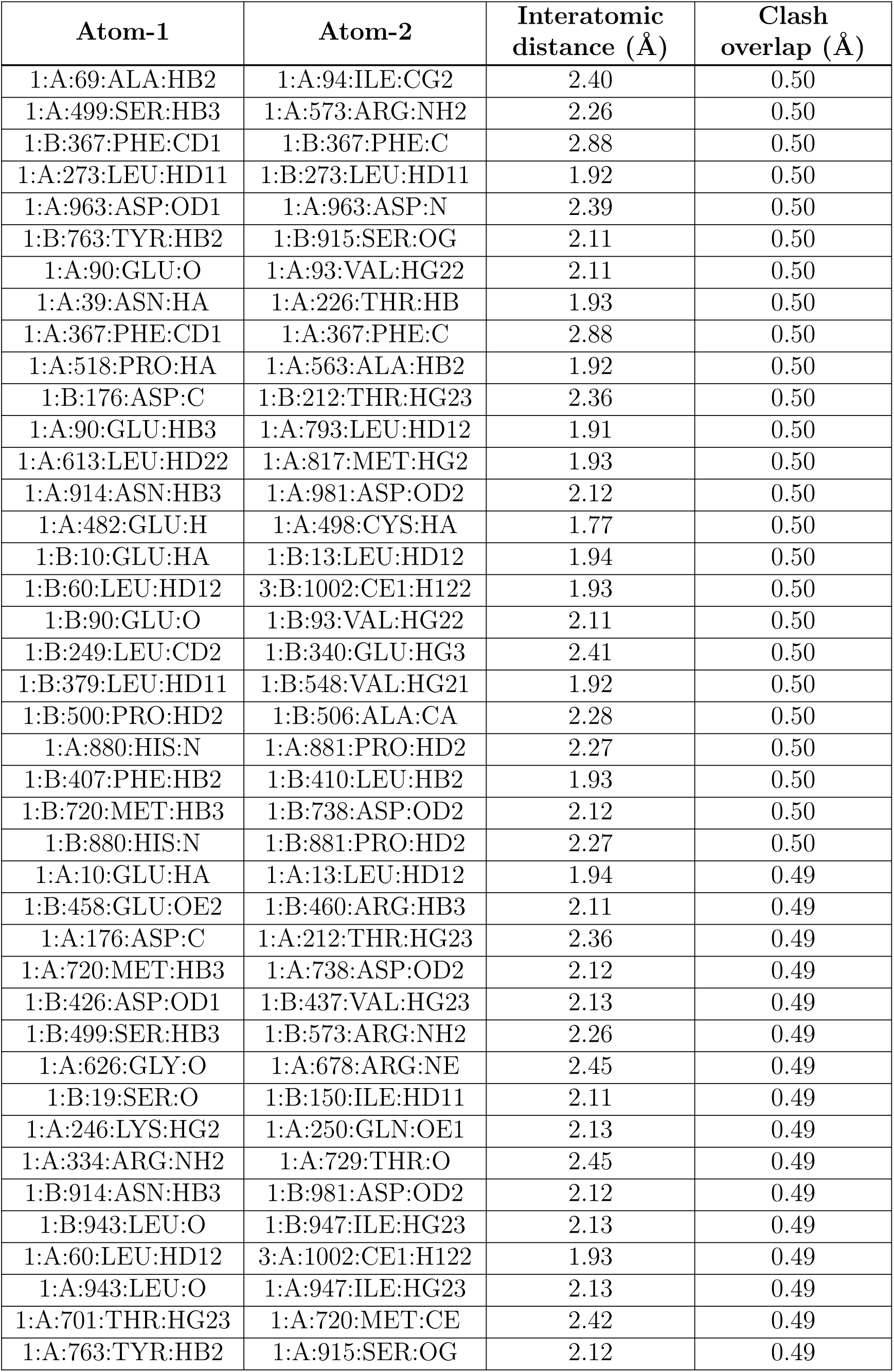
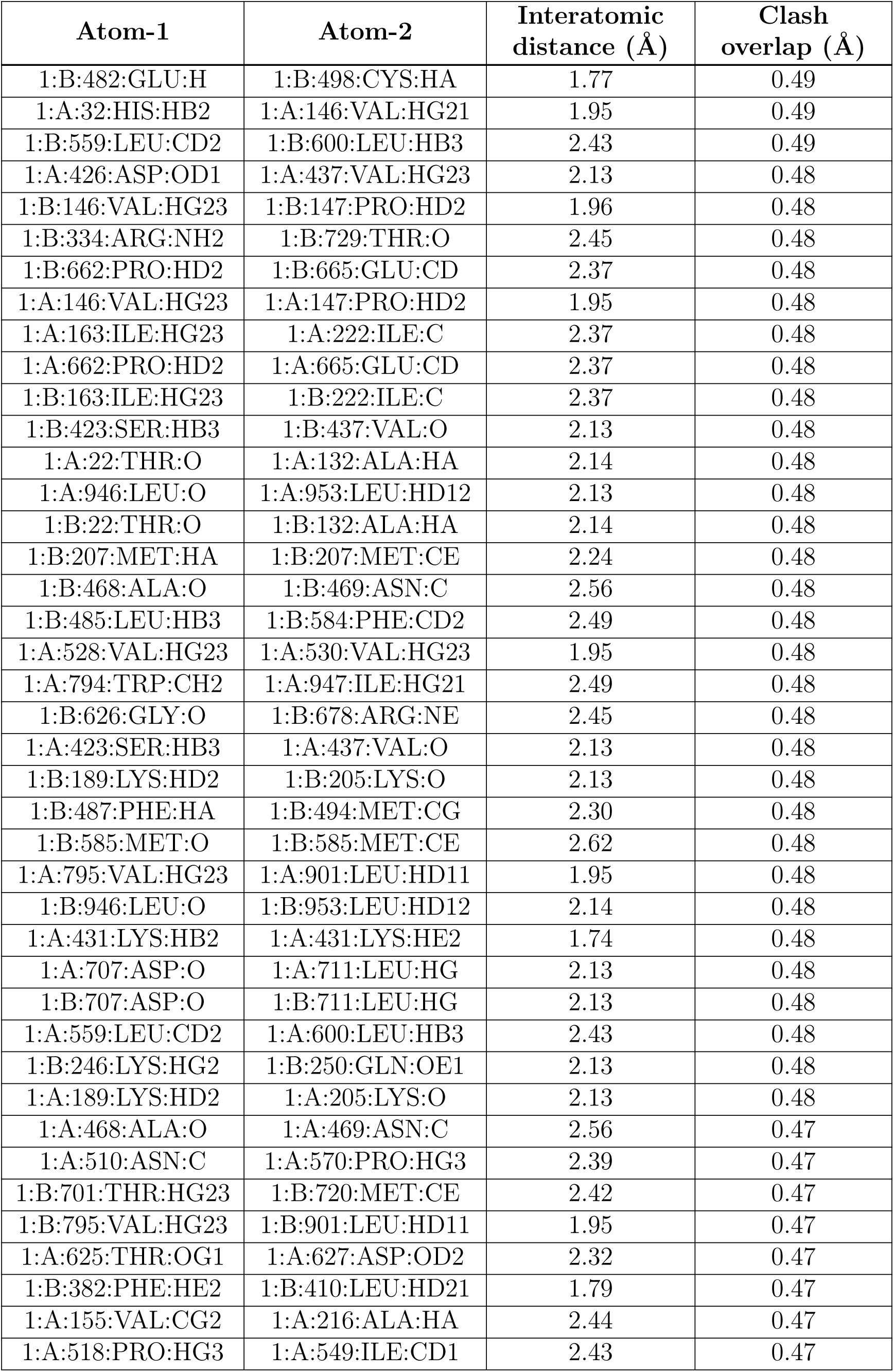
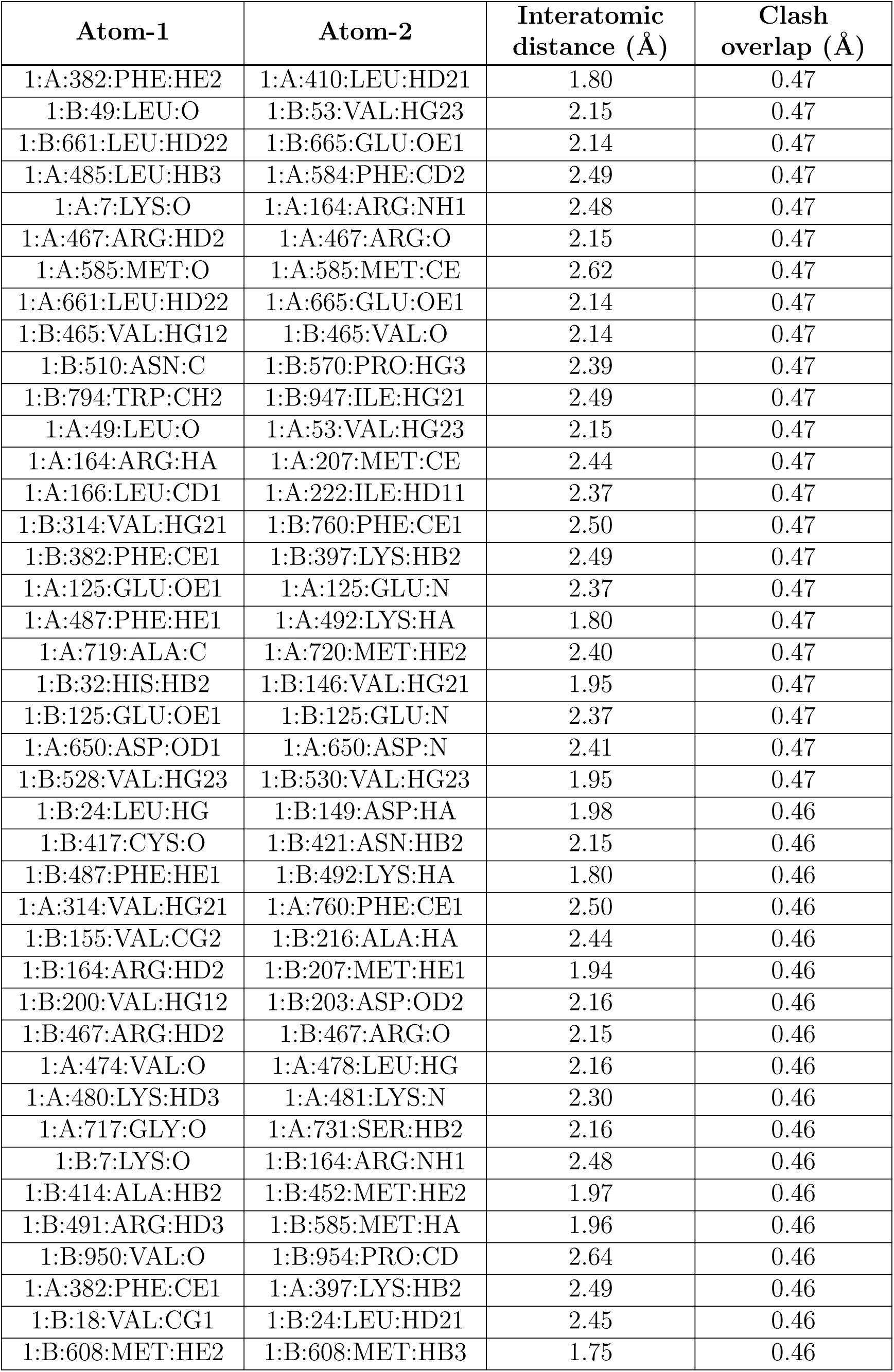
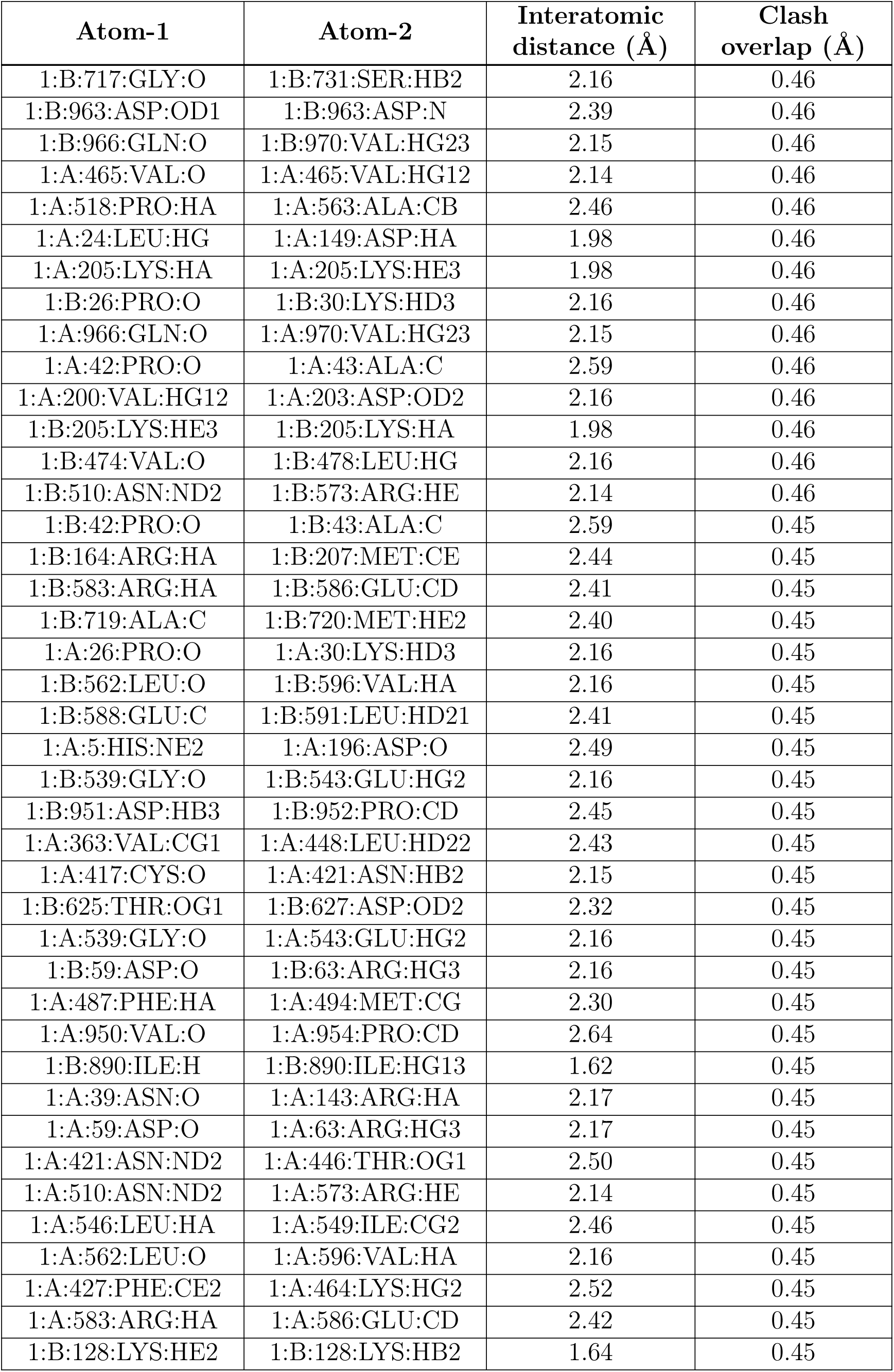
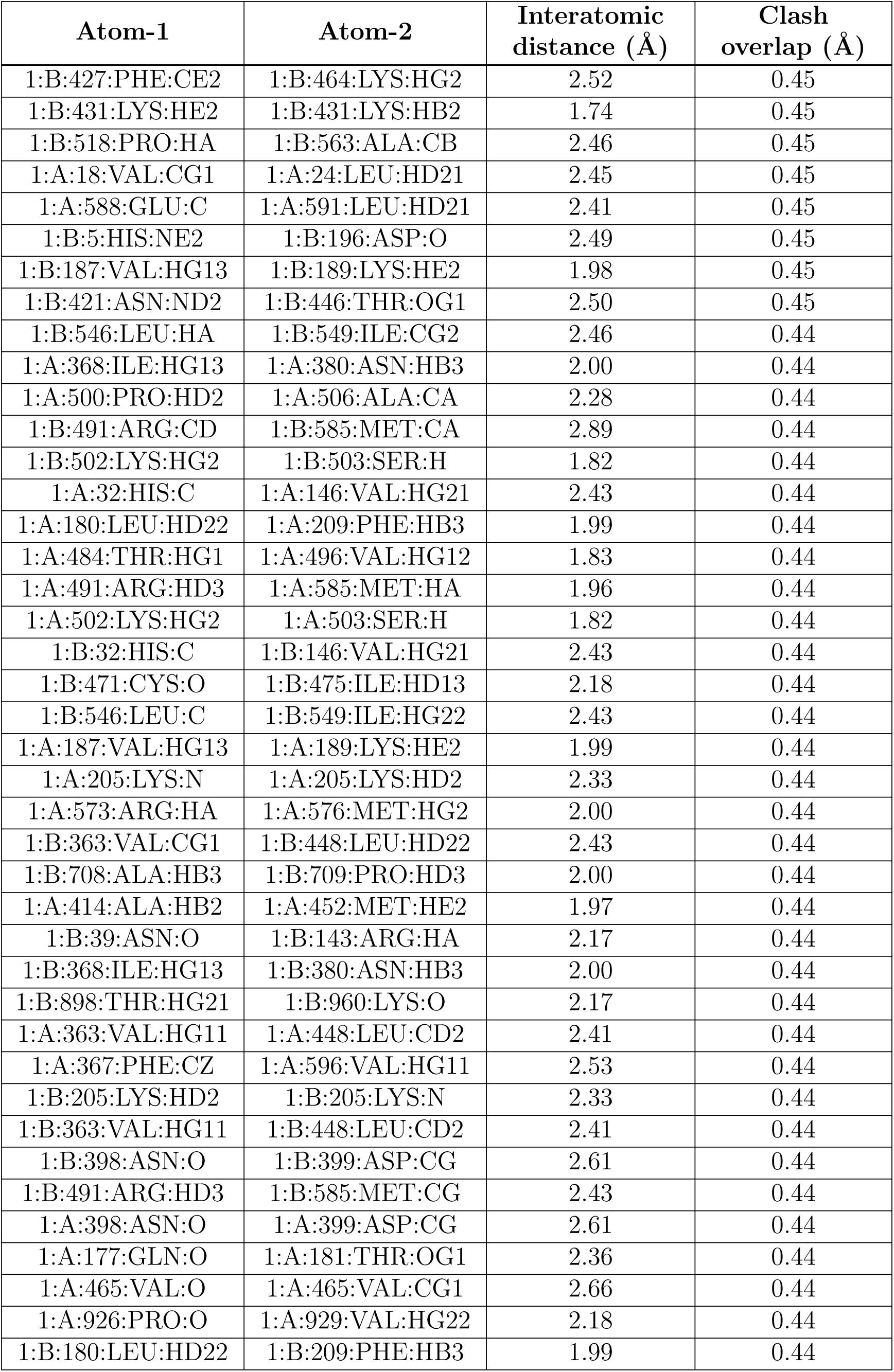
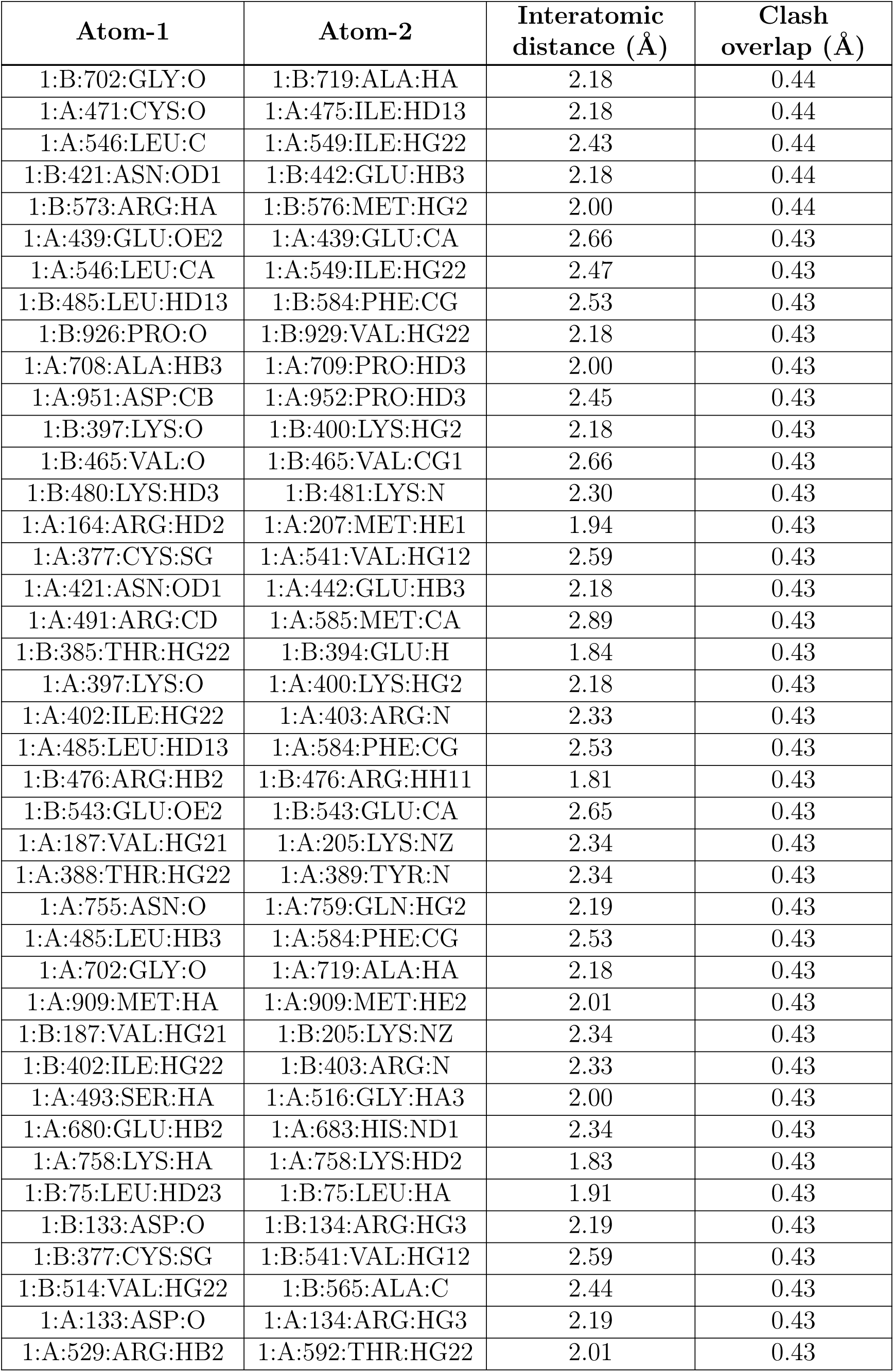
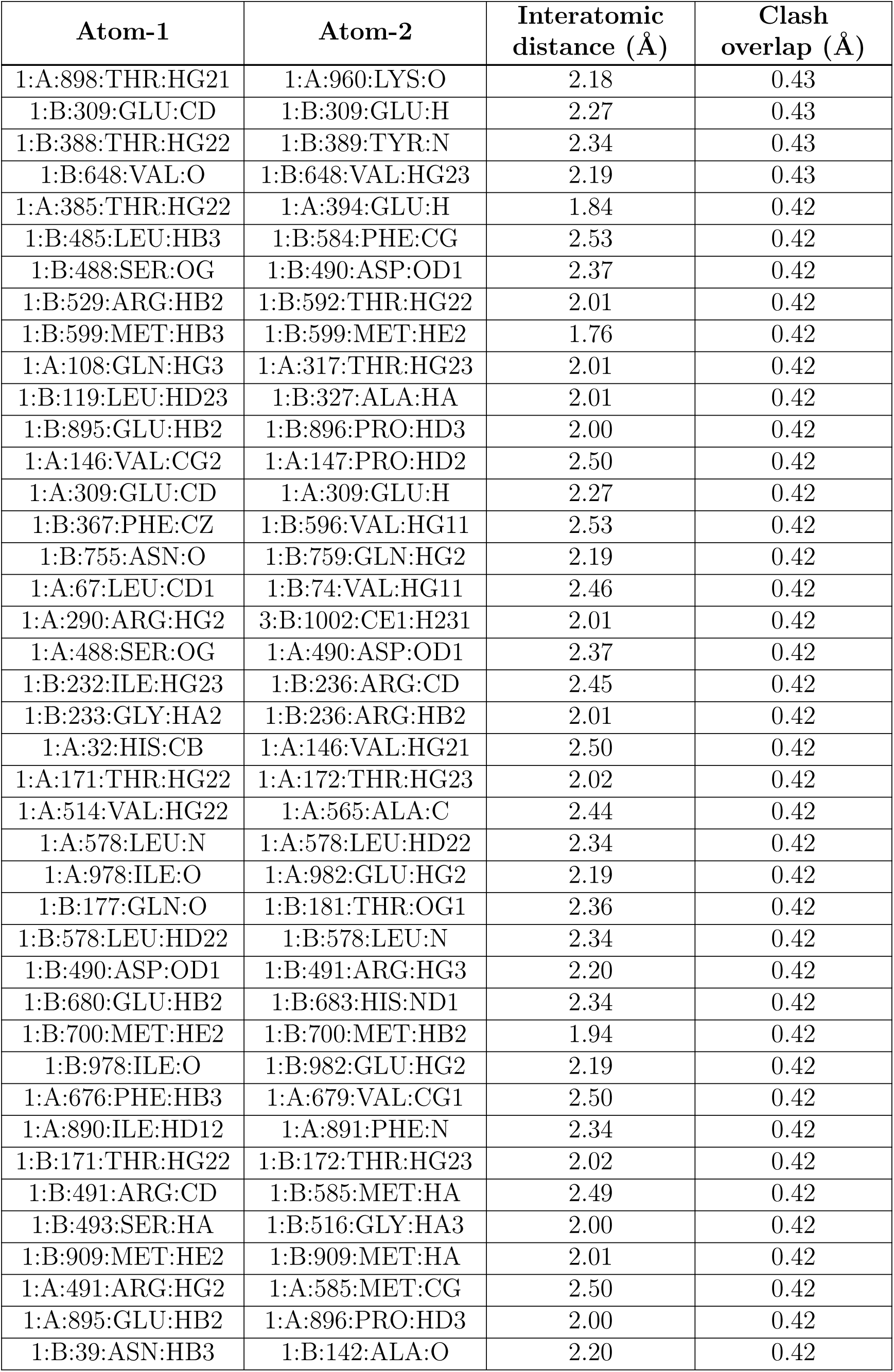
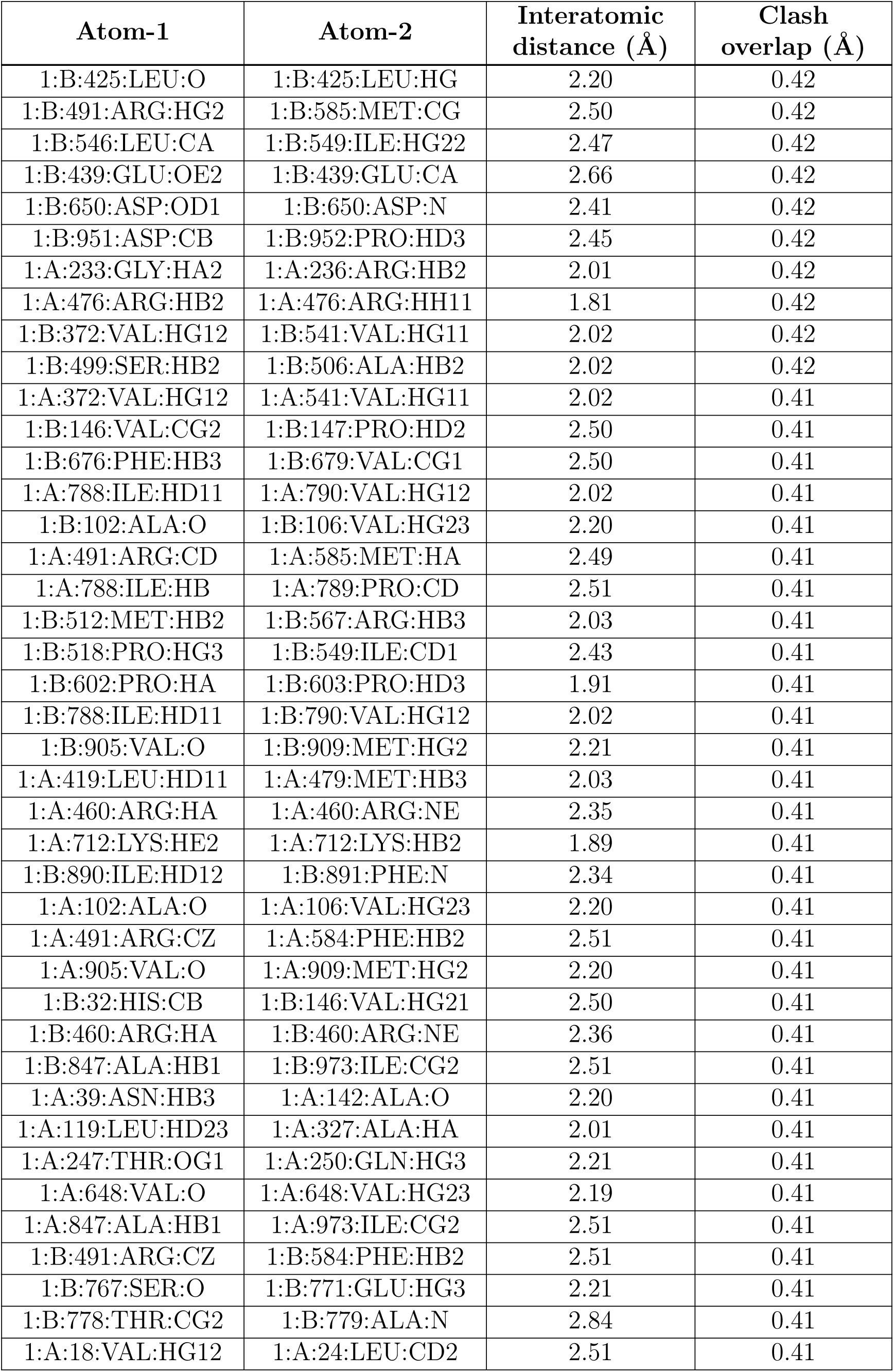
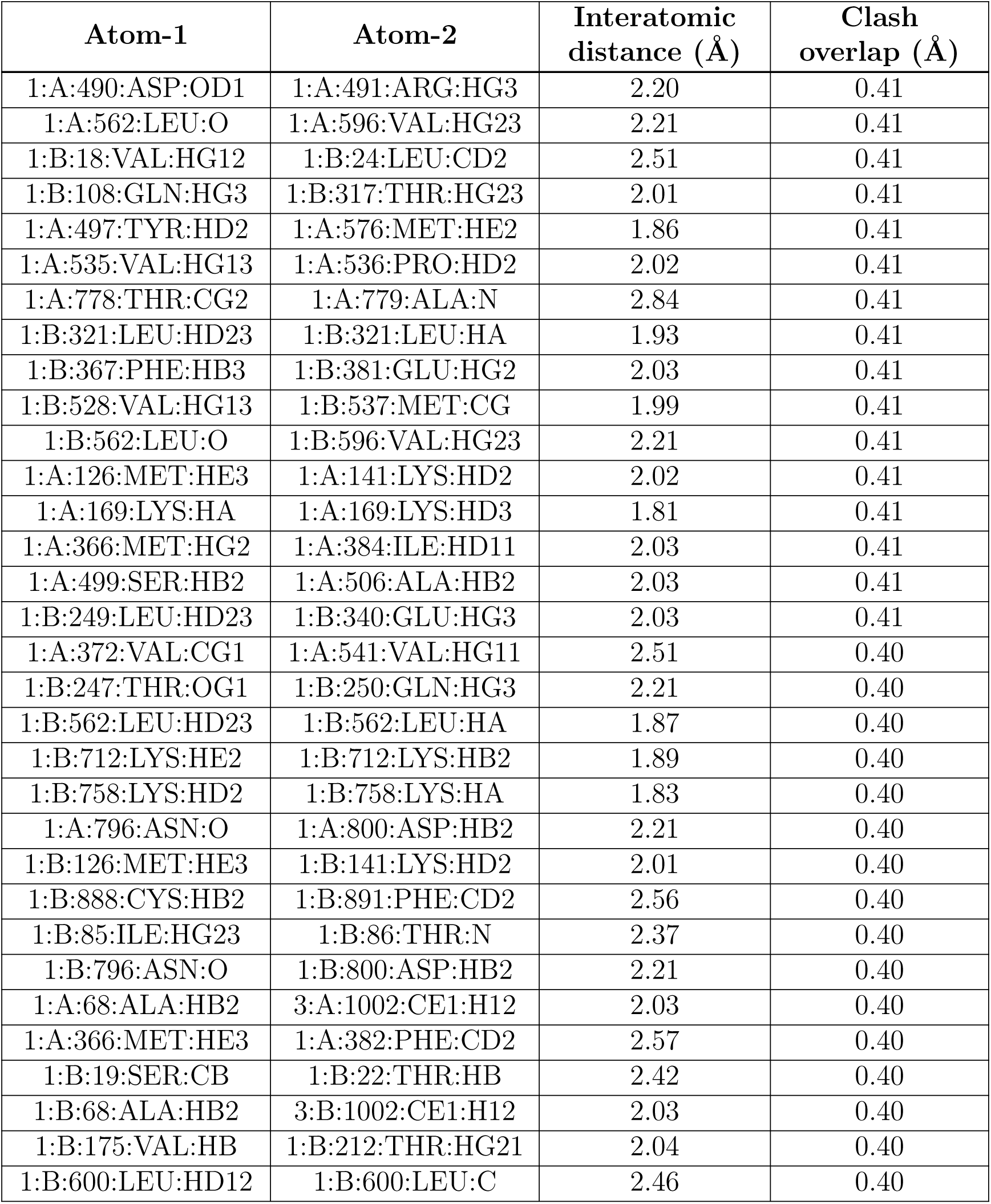

There are no symmetry-related clashes.

### 5.3 Torsion angles

#### 5.3.1 Protein backbone

In the following table, the Percentiles column shows the percent Ramachandran outliers of the chain as a percentile score with respect to all PDB entries followed by that with respect to all EM entries.

The Analysed column shows the number of residues for which the backbone conformation was analysed, and the total number of residues.

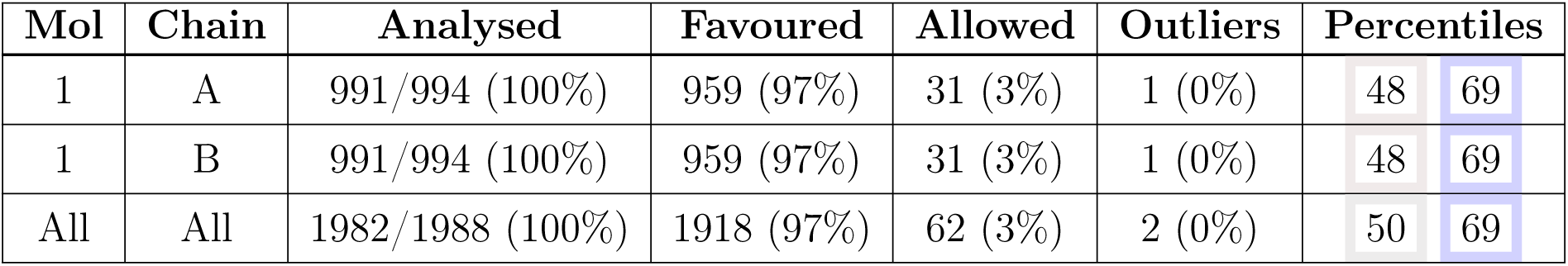

All (2) Ramachandran outliers are listed below:

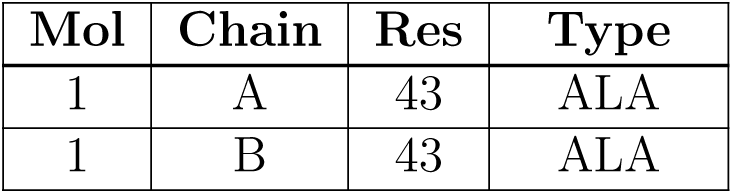

#### 5.3.2 Protein sidechains

In the following table, the Percentiles column shows the percent sidechain outliers of the chain as a percentile score with respect to all PDB entries followed by that with respect to all EM entries.

The Analysed column shows the number of residues for which the sidechain conformation was analysed, and the total number of residues.

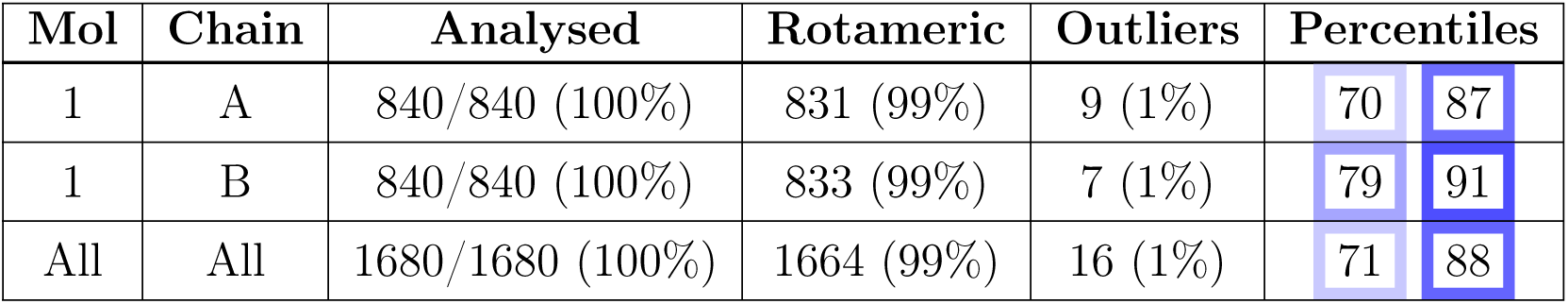

All (16) residues with a non-rotameric sidechain are listed below:

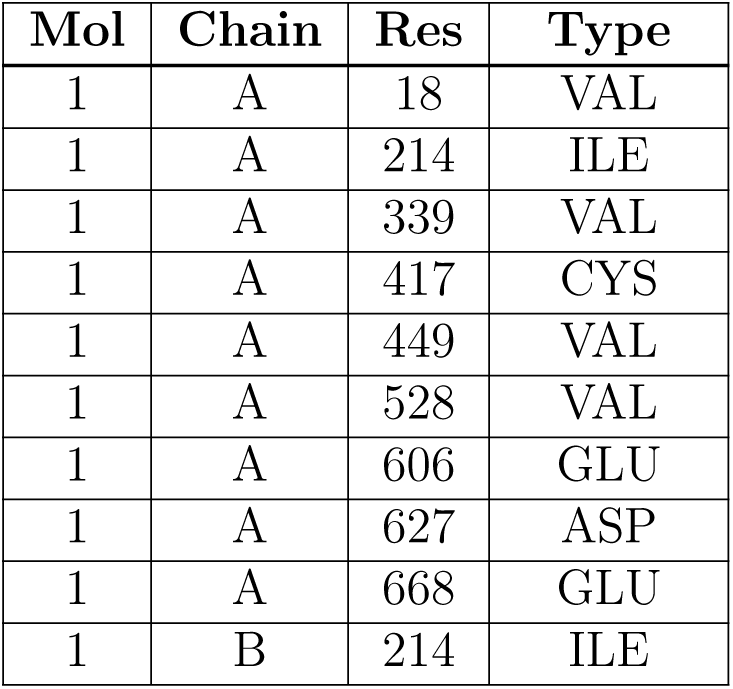
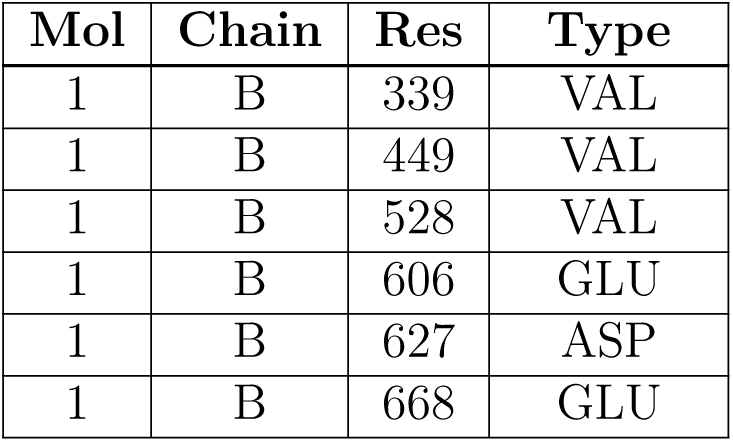

Sometimes sidechains can be flipped to improve hydrogen bonding and reduce clashes. All (9) such sidechains are listed below:

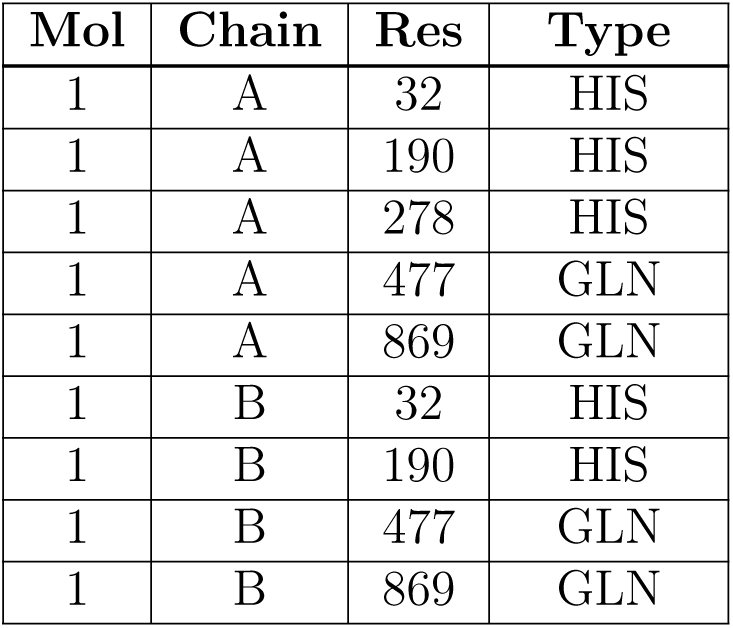

#### 5.3.3 RNA

There are no RNA molecules in this entry.

### 5.4 Non-standard residues in protein, DNA, RNA chains

There are no non-standard protein/DNA/RNA residues in this entry.

### 5.5 Carbohydrates

There are no oligosaccharides in this entry.

### 5.6 Ligand geometry

Of 8 ligands modelled in this entry, 4 are monoatomic - leaving 4 for Mogul analysis.

In the following table, the Counts columns list the number of bonds (or angles) for which Mogul statistics could be retrieved, the number of bonds (or angles) that are observed in the model and the number of bonds (or angles) that are defined in the Chemical Component Dictionary. The Link column lists molecule types, if any, to which the group is linked. The Z score for a bond length (or angle) is the number of standard deviations the observed value is removed from the expected value. A bond length (or angle) with *|Z| >* 2 is considered an outlier worth inspection. RMSZ is the root-mean-square of all Z scores of the bond lengths (or angles).

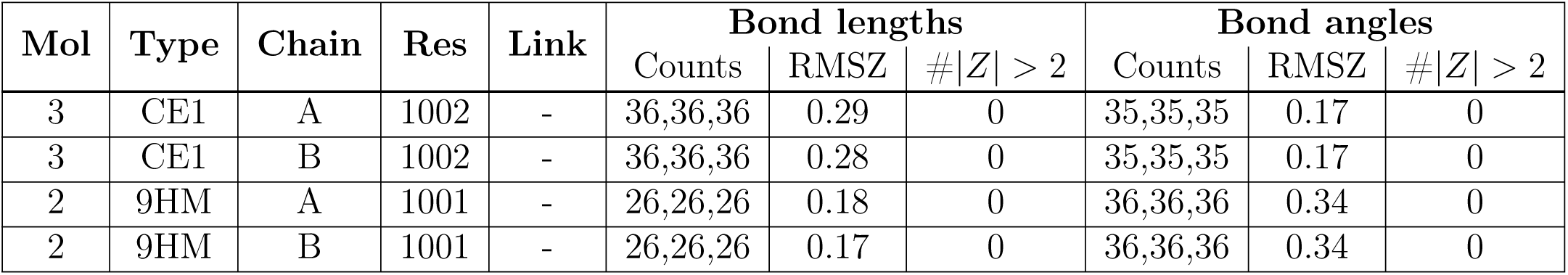

In the following table, the Chirals column lists the number of chiral outliers, the number of chiral centers analysed, the number of these observed in the model and the number defined in the Chemical Component Dictionary. Similar counts are reported in the Torsion and Rings columns.’-’ means no outliers of that kind were identified.

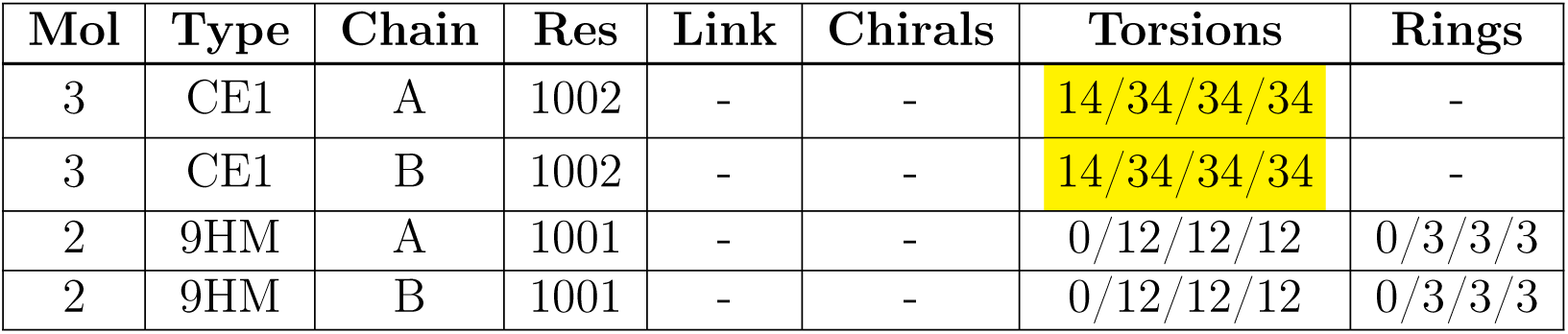

There are no bond length outliers.

There are no bond angle outliers.

There are no chirality outliers.

All (28) torsion outliers are listed below:

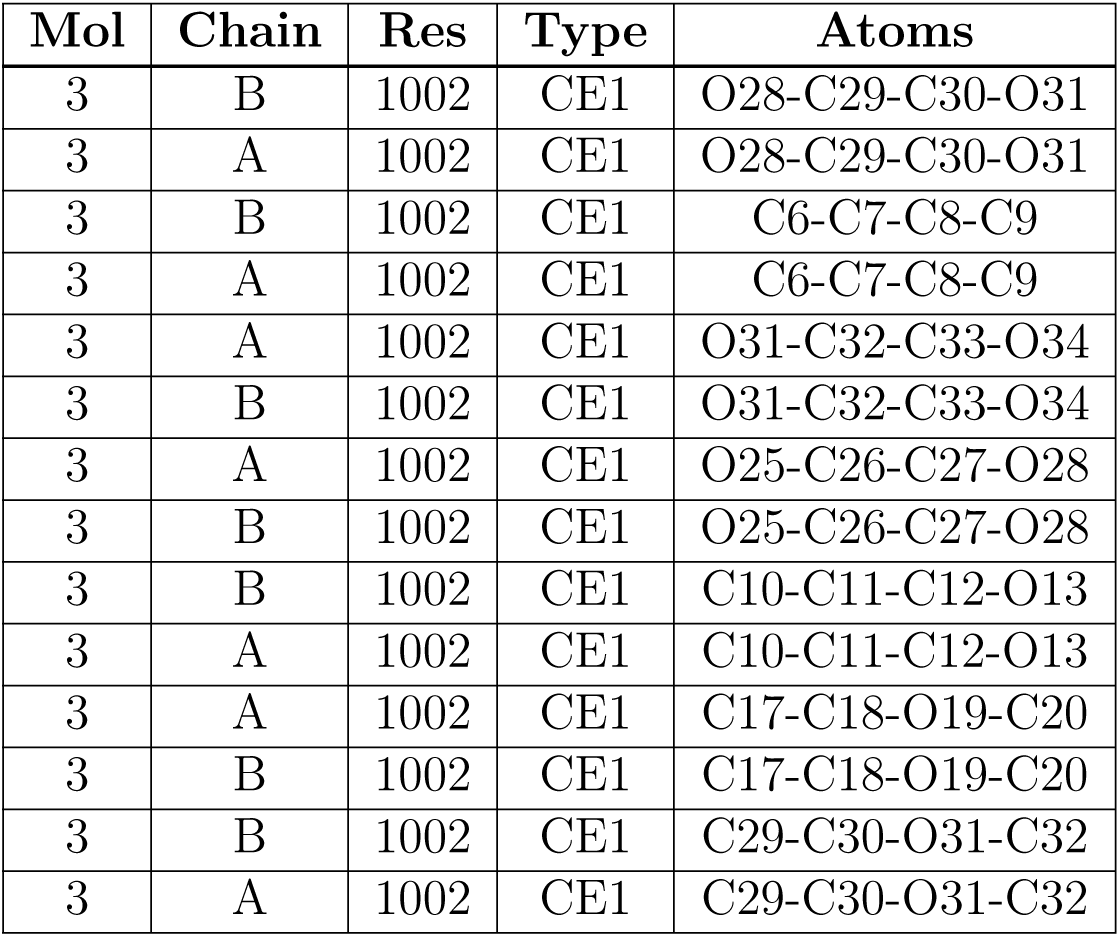
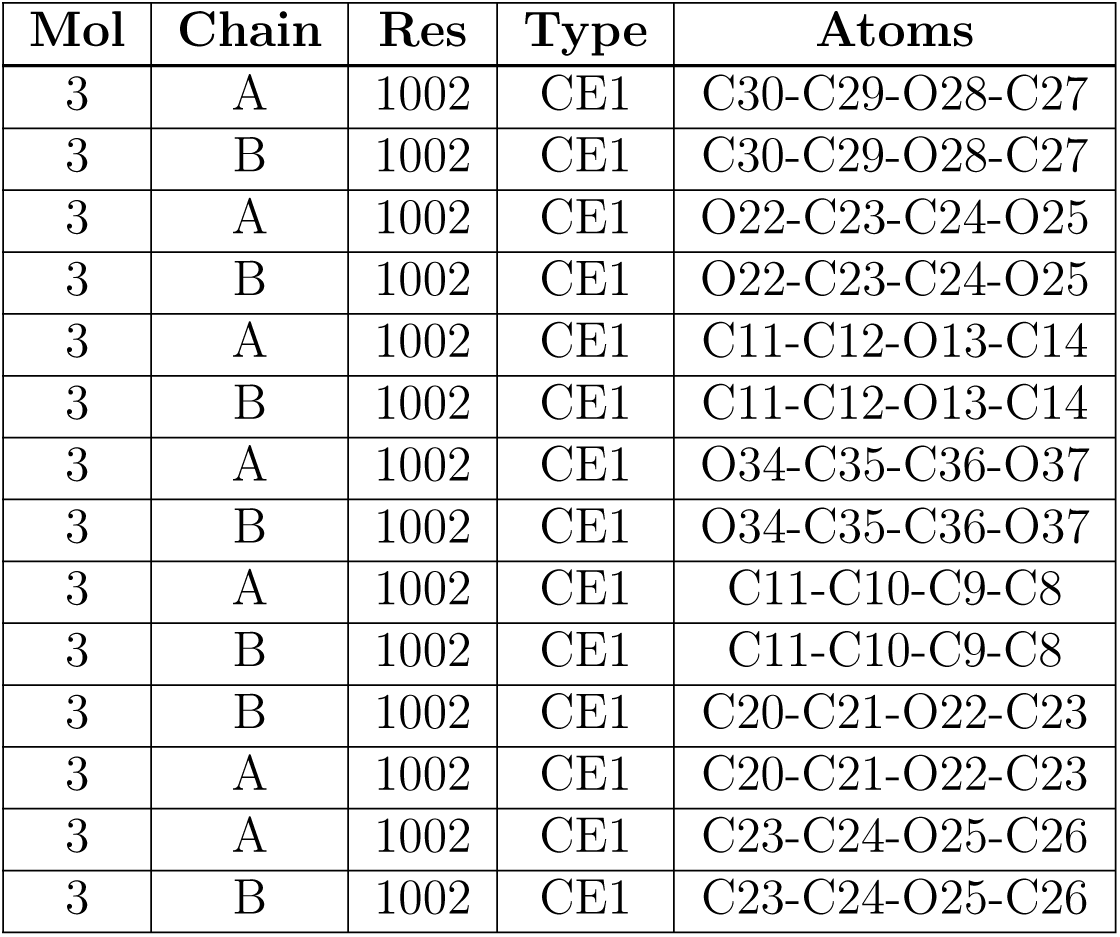

There are no ring outliers.

2 monomers are involved in 7 short contacts:

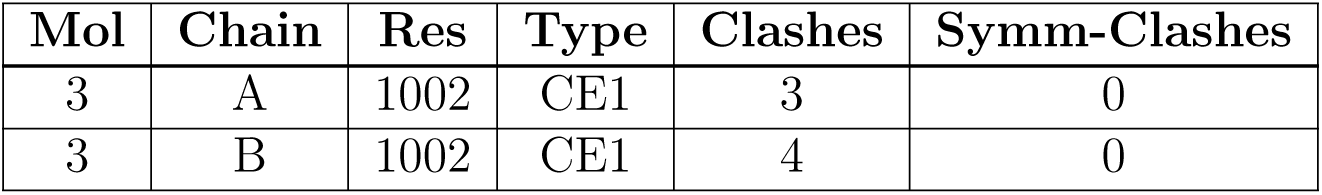

The following is a two-dimensional graphical depiction of Mogul quality analysis of bond lengths, bond angles, torsion angles, and ring geometry for all instances of the Ligand of Interest. In addition, ligands with molecular weight *>* 250 and outliers as shown on the validation Tables will also be included. For torsion angles, if less then 5% of the Mogul distribution of torsion angles is within 10 degrees of the torsion angle in question, then that torsion angle is considered an outlier. Any bond that is central to one or more torsion angles identified as an outlier by Mogul will be highlighted in the graph. For rings, the root-mean-square deviation (RMSD) between the ring in question and similar rings identified by Mogul is calculated over all ring torsion angles. If the average RMSD is greater than 60 degrees and the minimal RMSD between the ring in question and any Mogul-identified rings is also greater than 60 degrees, then that ring is considered an outlier. The outliers are highlighted in purple. The color gray indicates Mogul did not find sufficient equivalents in the CSD to analyse the geometry.

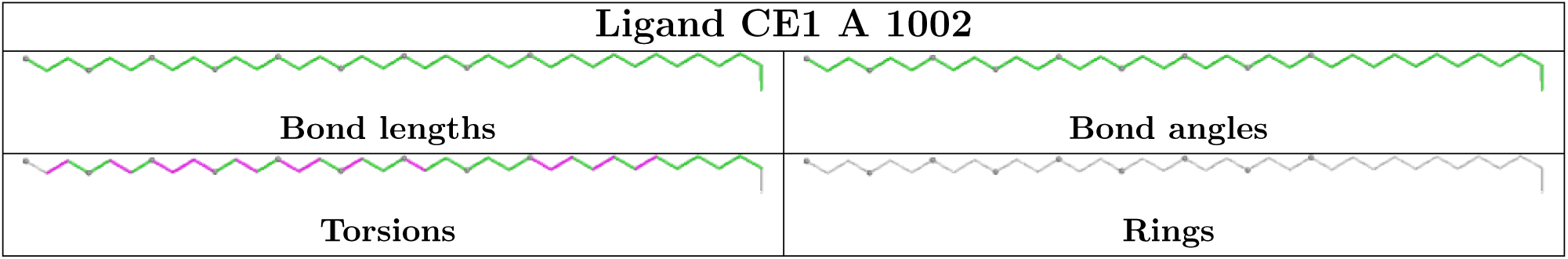
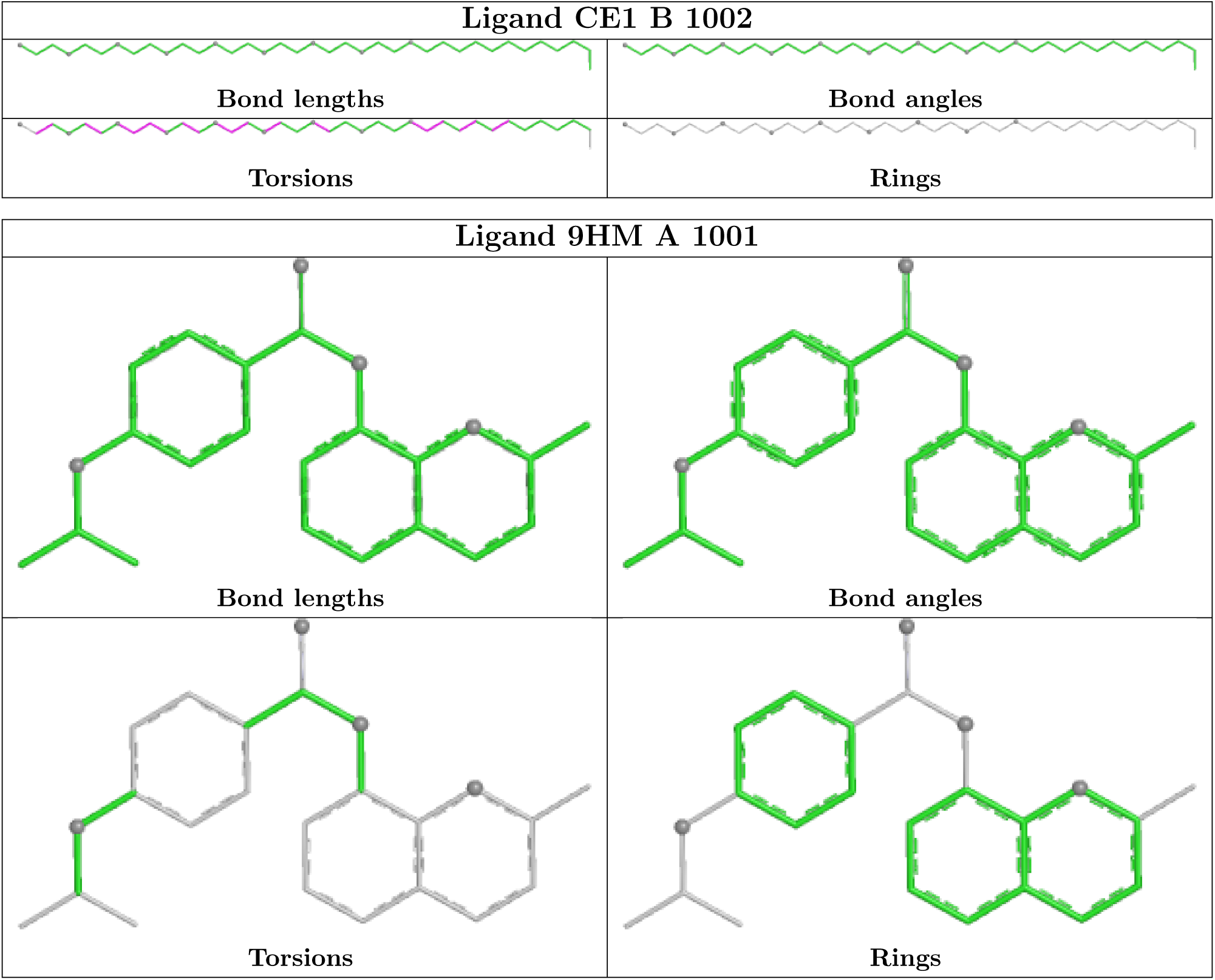
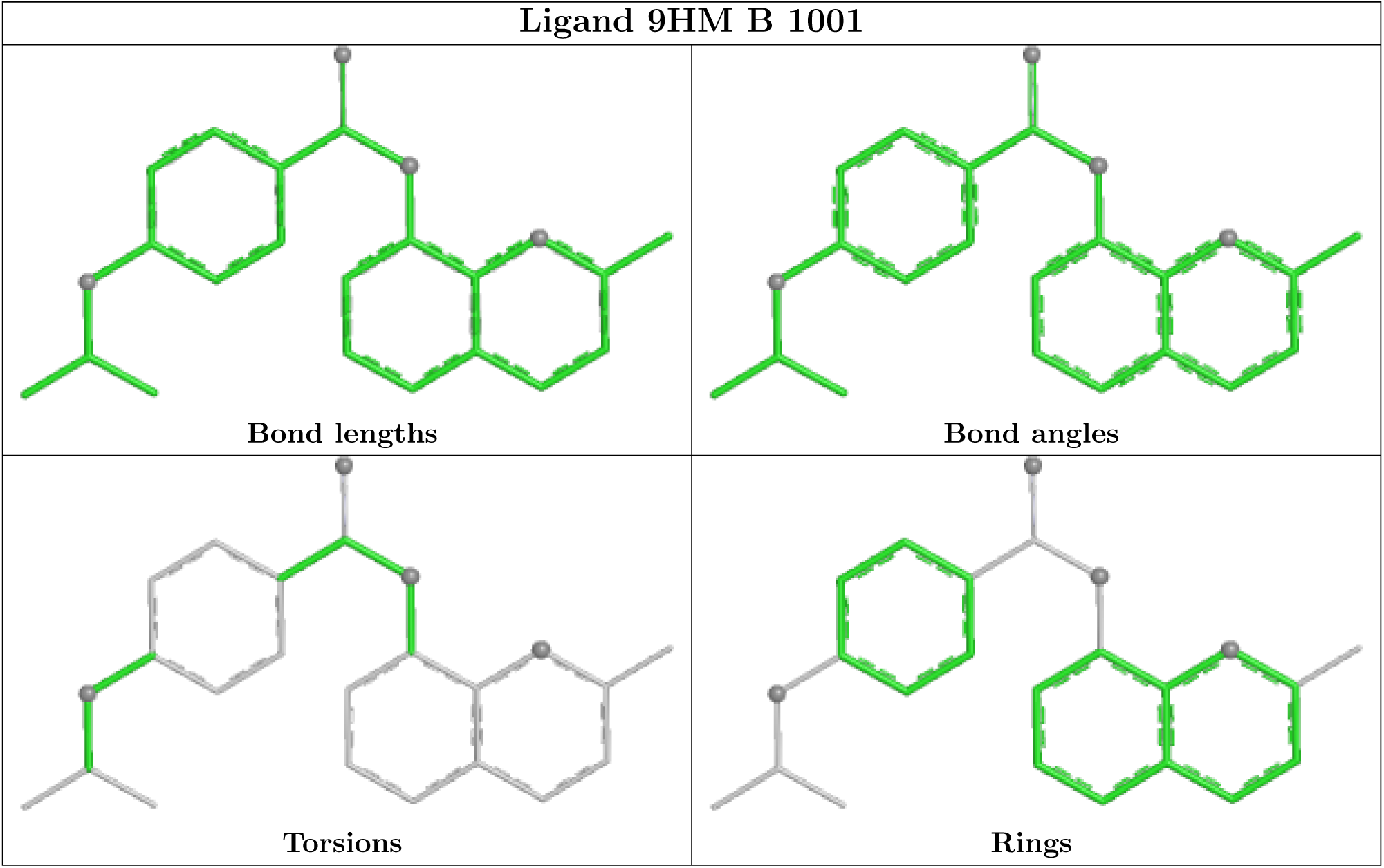

### 5.7 Other polymers

There are no such residues in this entry.

### 5.8 Polymer linkage issues

There are no chain breaks in this entry.

## 6 Map visualisation

This section contains visualisations of the EMDB entry EMD-73823. These allow visual inspection of the internal detail of the map and identification of artifacts.

Images derived from a raw map, generated by summing the deposited half-maps, are presented below the corresponding image components of the primary map to allow further visual inspection and comparison with those of the primary map.

### 6.1 Orthogonal projections

#### 6.1.1 Primary map

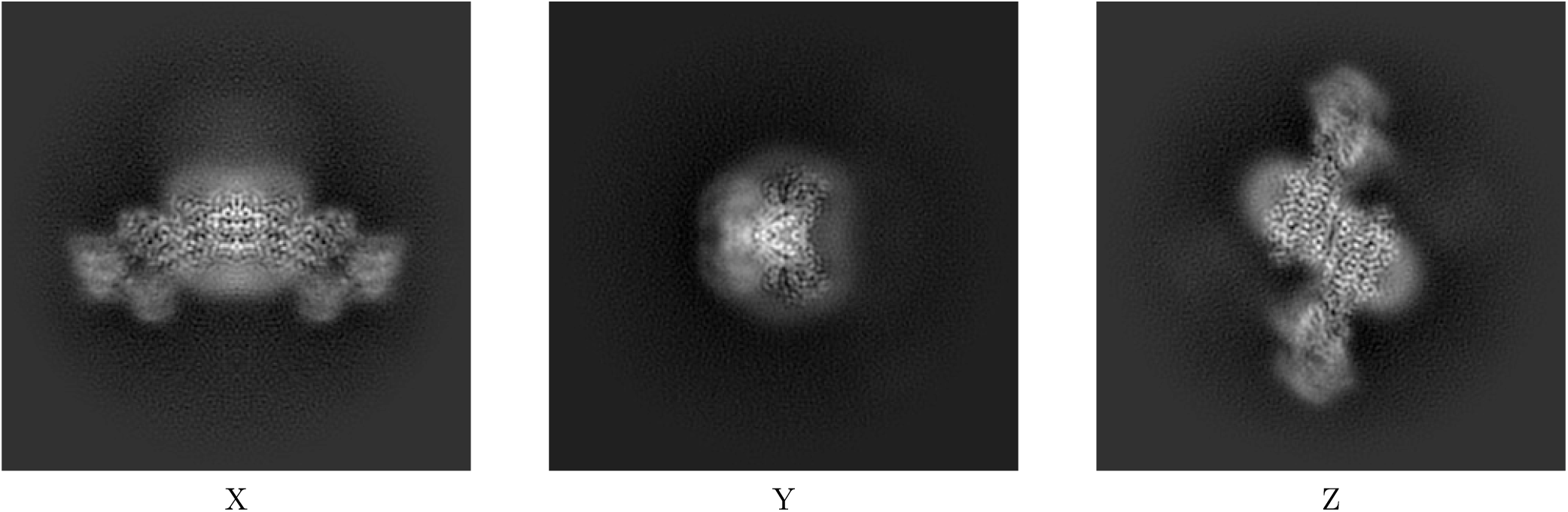

#### 6.1.2 Raw map

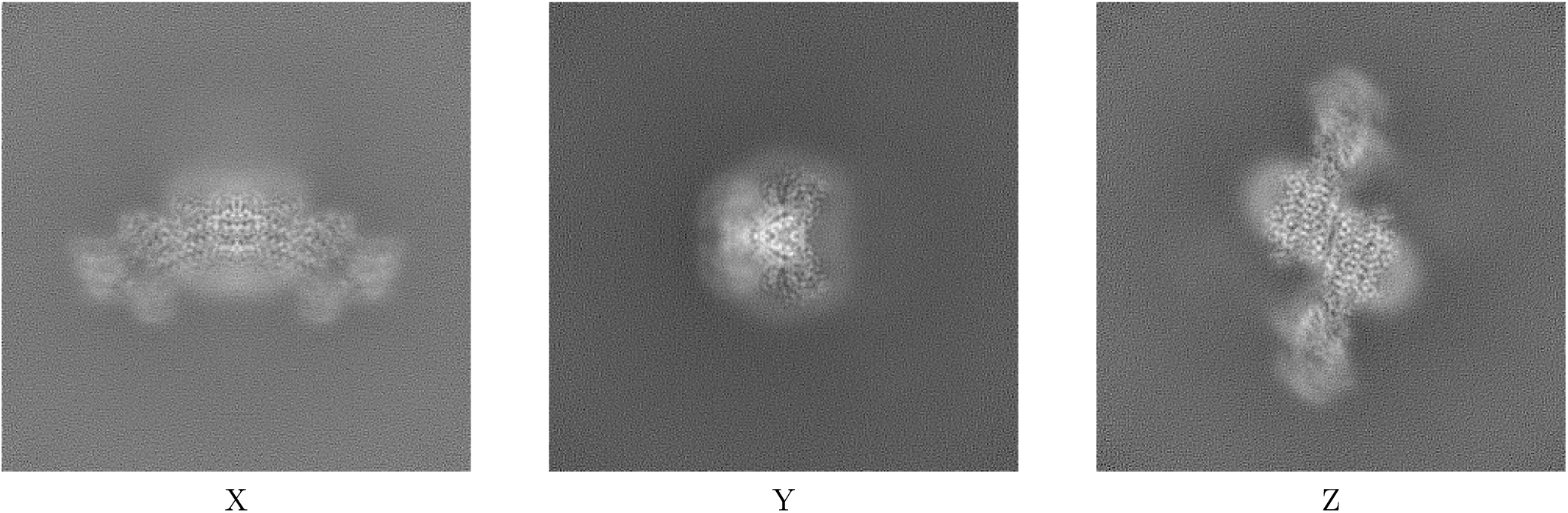

The images above show the map projected in three orthogonal directions.

### 6.2 Central slices

#### 6.2.1 Primary map

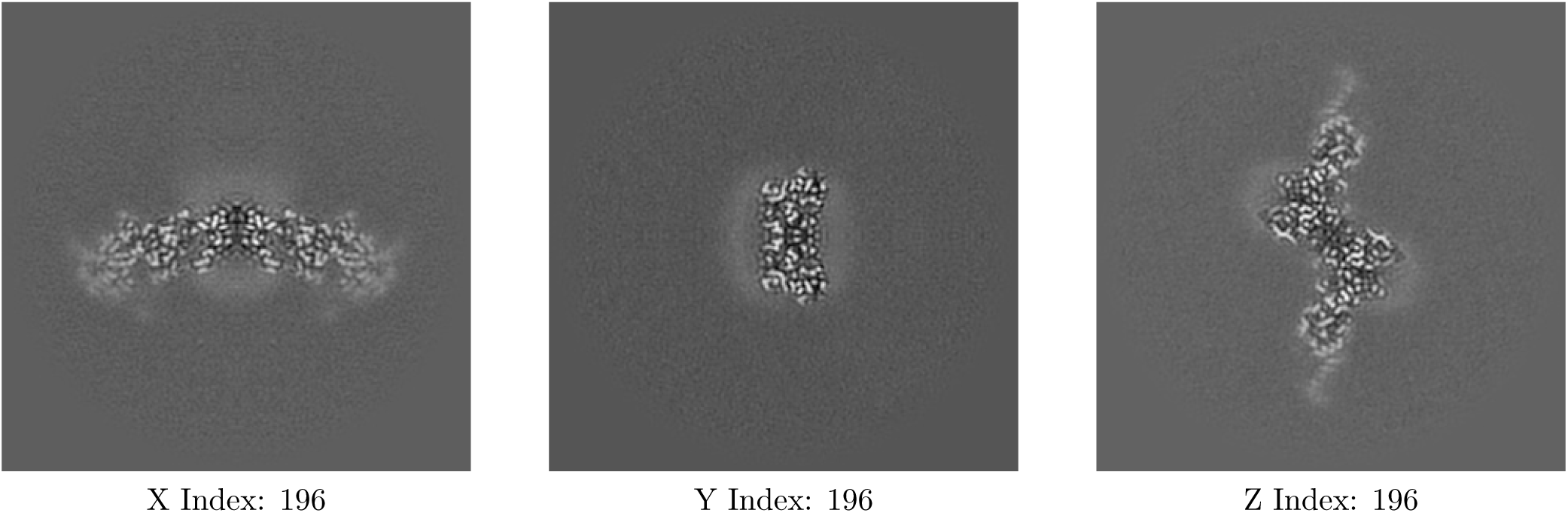

#### 6.2.2 Raw map

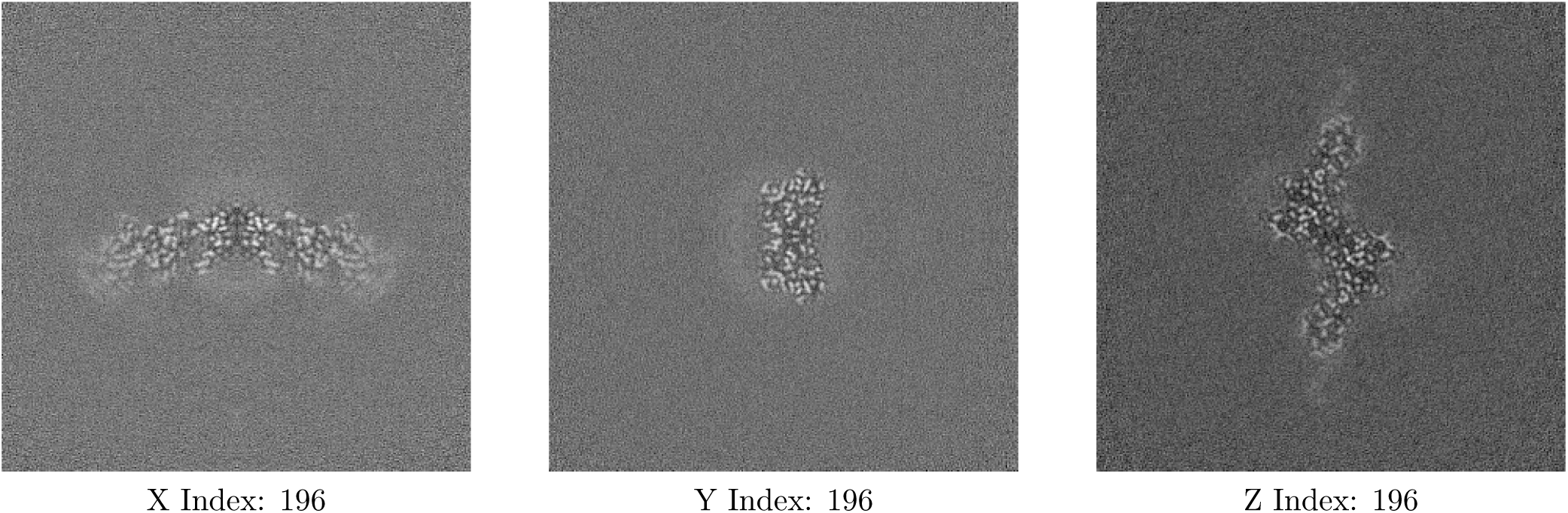

The images above show central slices of the map in three orthogonal directions.

### 6.3 Largest variance slices

#### 6.3.1 Primary map

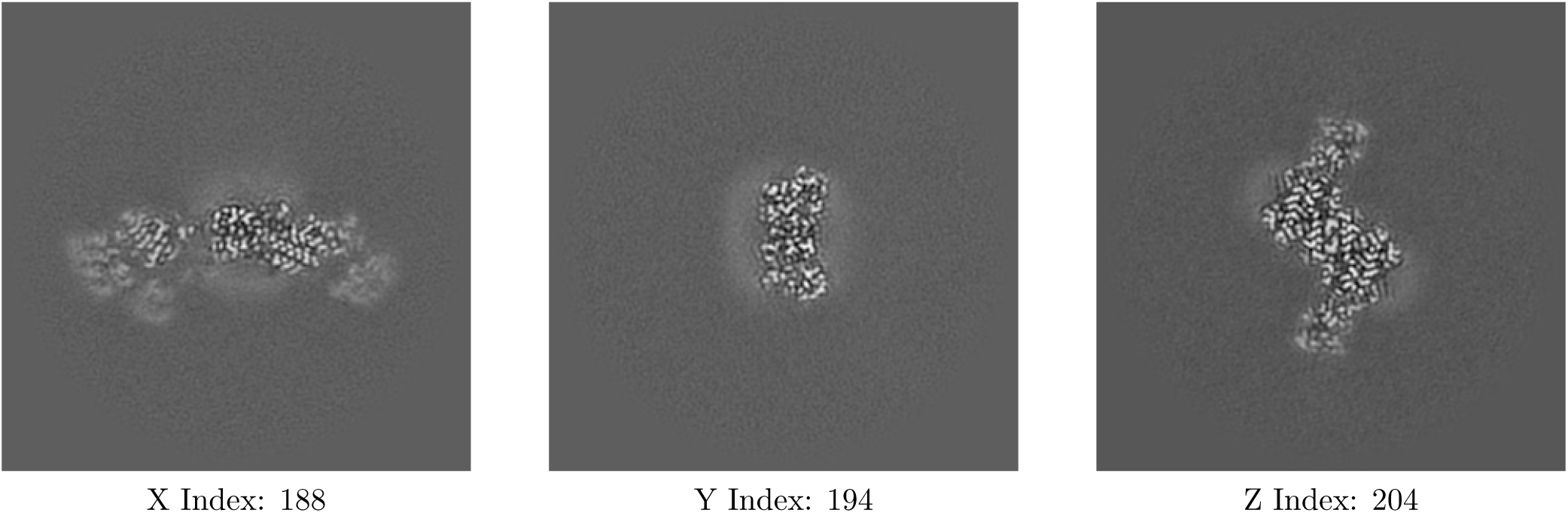

#### 6.3.2 Raw map

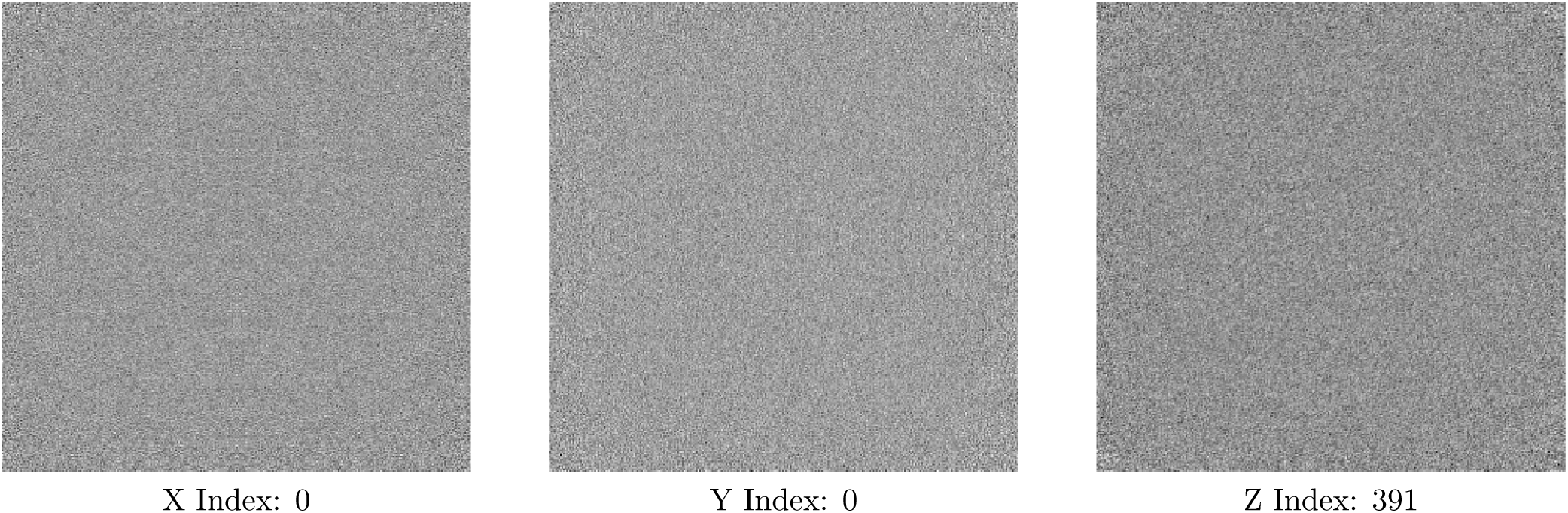

The images above show the largest variance slices of the map in three orthogonal directions.

### 6.4 Orthogonal standard-deviation projections (False-color)

#### 6.4.1 Primary map

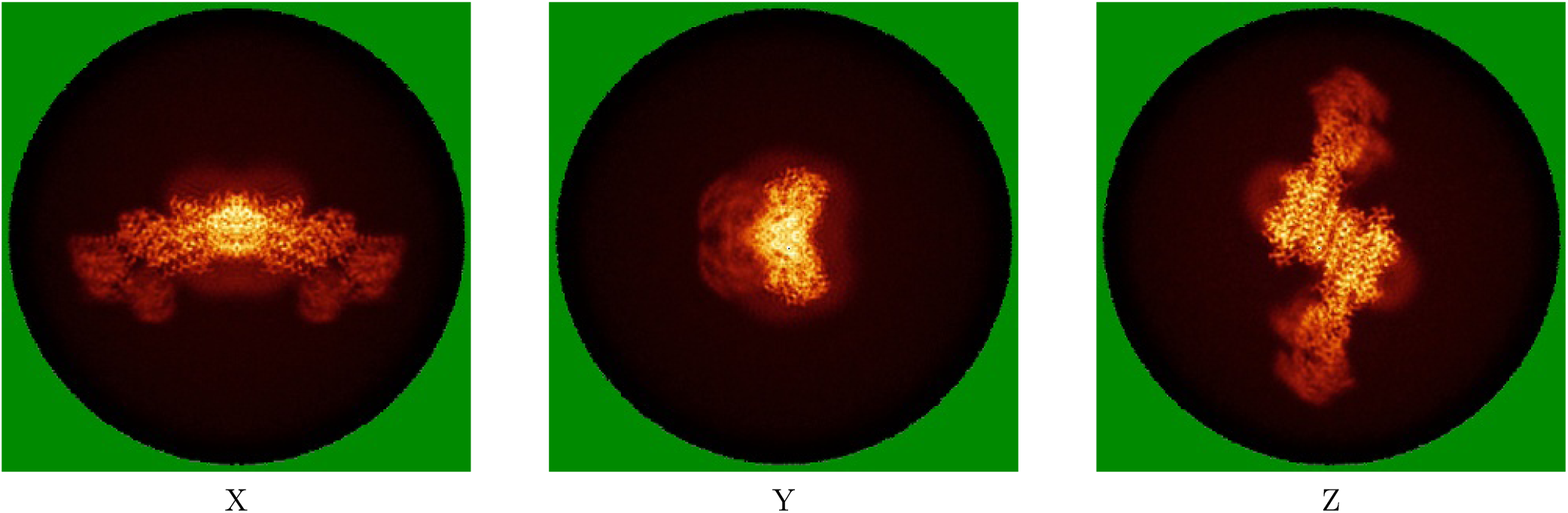

#### 6.4.2 Raw map

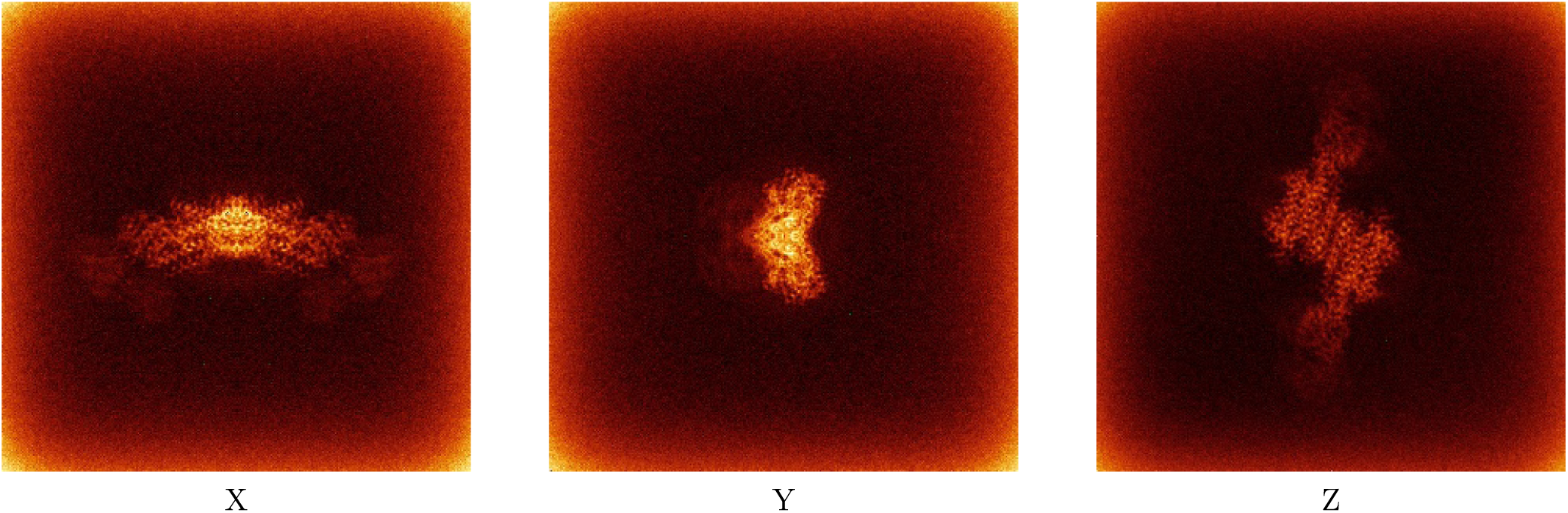

The images above show the map standard deviation projections with false color in three orthogonal directions. Minimum values are shown in green, max in blue, and dark to light orange shades represent small to large values respectively.

### 6.5 Orthogonal surface views

#### 6.5.1 Primary map

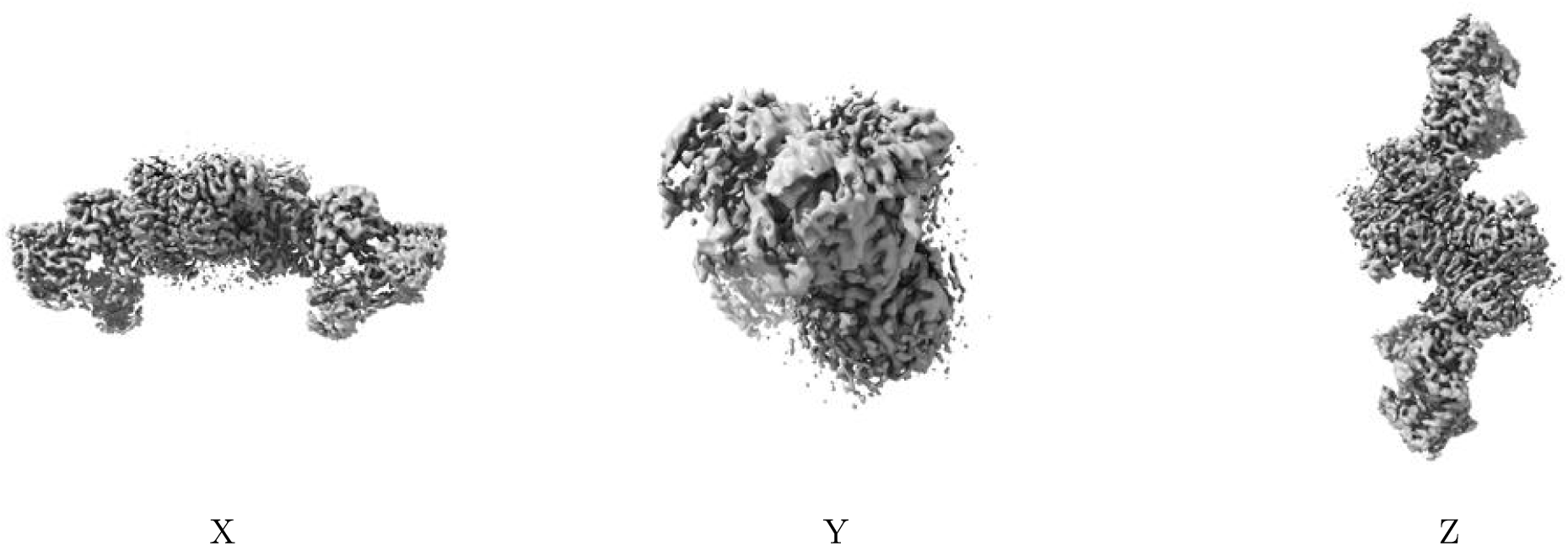

The images above show the 3D surface view of the map at the recommended contour level 0.065. These images, in conjunction with the slice images, may facilitate assessment of whether an appropriate contour level has been provided.

#### 6.5.2 Raw map

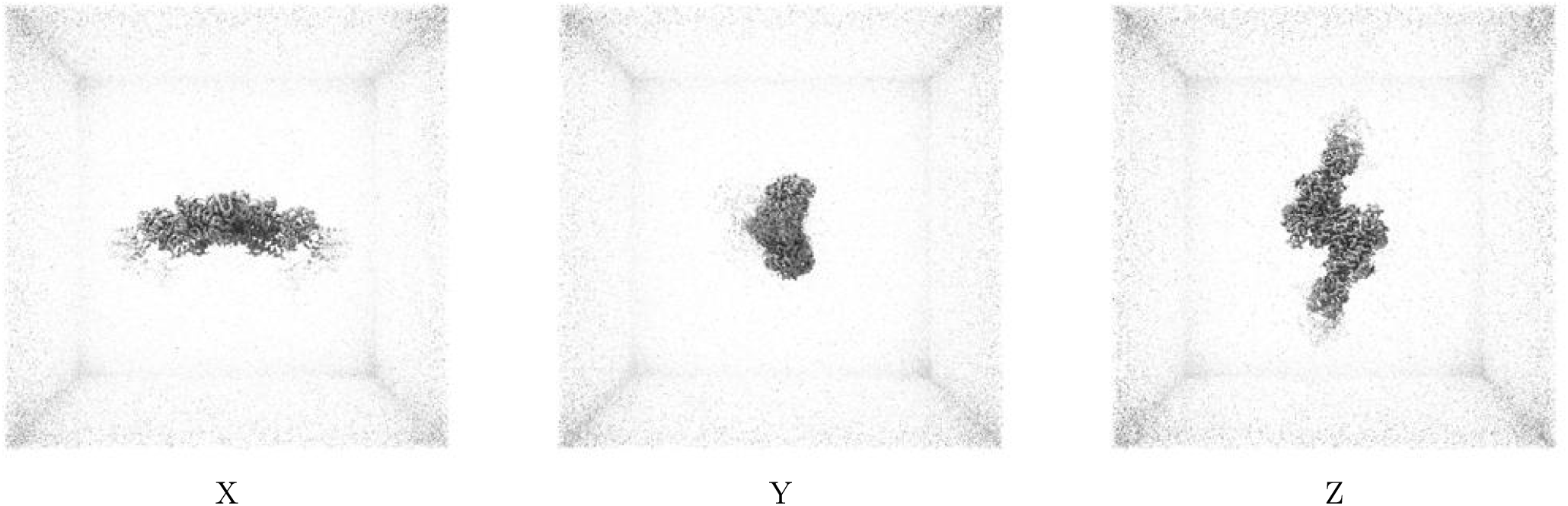

These images show the 3D surface of the raw map. The raw map’s contour level was selected so that its surface encloses the same volume as the primary map does at its recommended contour level.

### 6.6 Mask visualisation

This section was not generated. No masks/segmentation were deposited.

## 7 Map analysis

This section contains the results of statistical analysis of the map.

### 7.1 Map-value distribution

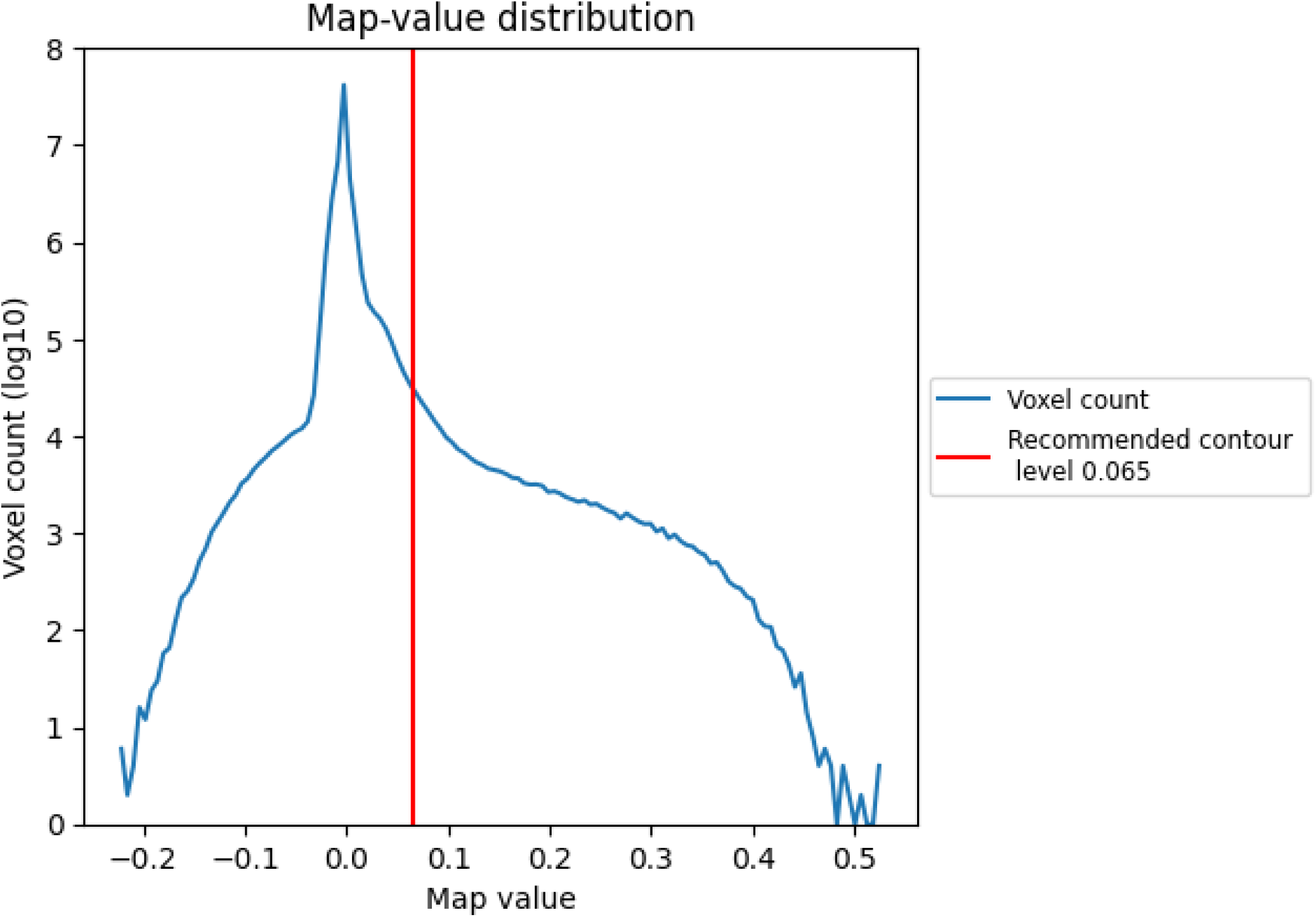

The map-value distribution is plotted in 128 intervals along the x-axis. The y-axis is logarithmic. A spike in this graph at zero usually indicates that the volume has been masked.

### 7.2 Volume estimate

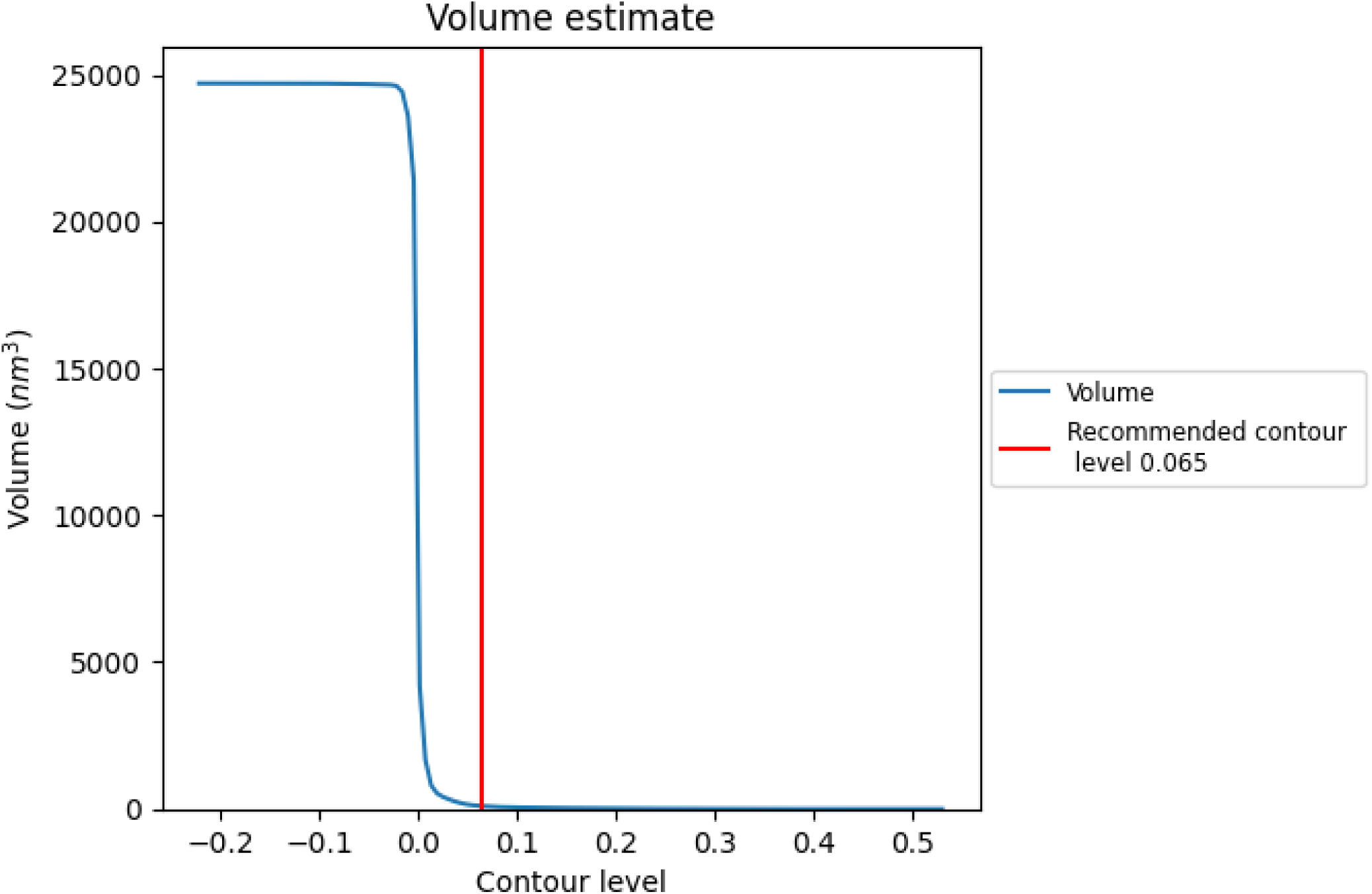

The volume at the recommended contour level is 101 nm^3^; this corresponds to an approximate mass of 91 kDa.

The volume estimate graph shows how the enclosed volume varies with the contour level. The recommended contour level is shown as a vertical line and the intersection between the line and the curve gives the volume of the enclosed surface at the given level.

### 7.3 Rotationally averaged power spectrum

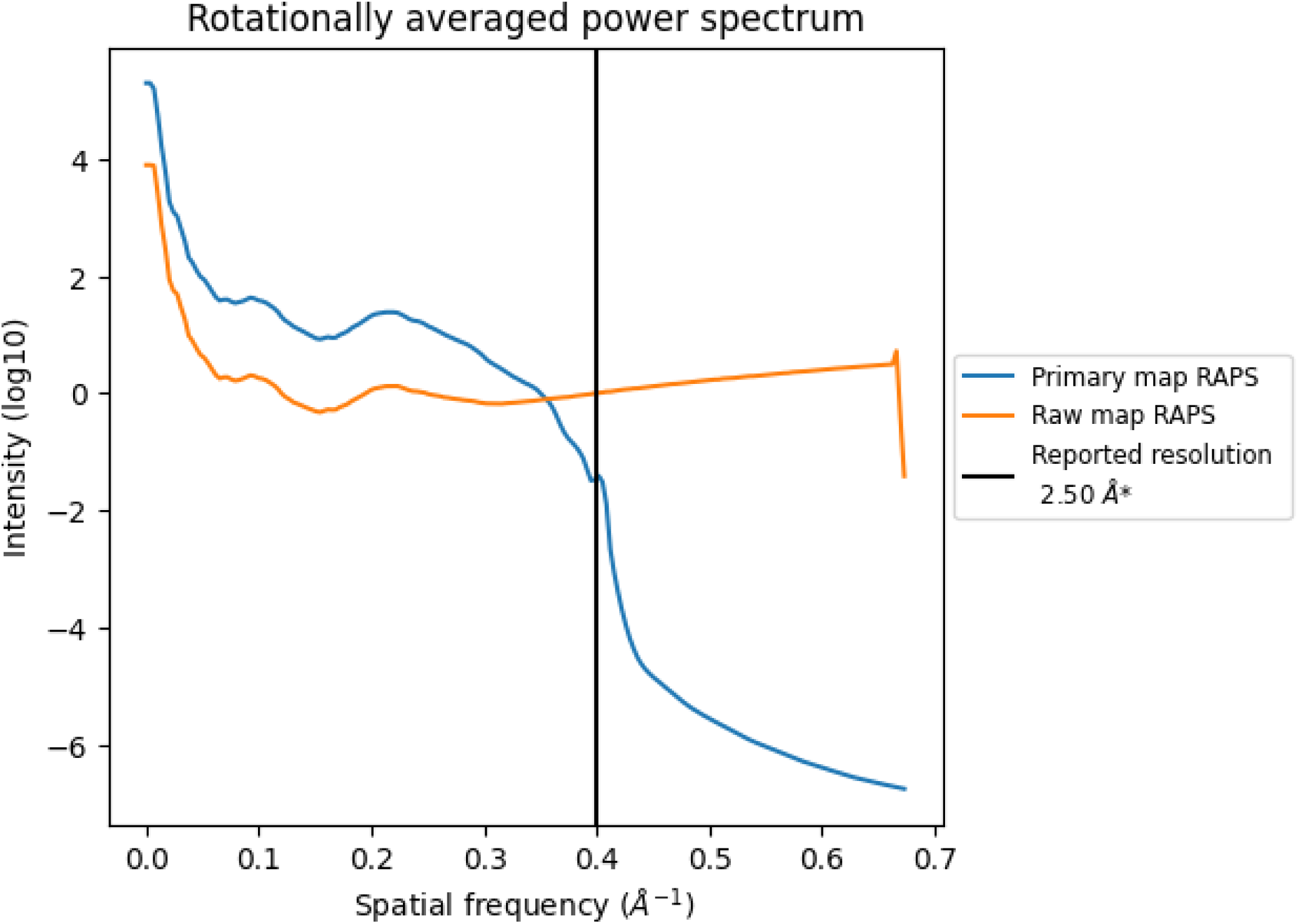

*Reported resolution corresponds to spatial frequency of 0.400 Å*^−^*^1^

## 8 Fourier-Shell correlation

Fourier-Shell Correlation (FSC) is the most commonly used method to estimate the resolution of single-particle and subtomogram-averaged maps. The shape of the curve depends on the imposed symmetry, mask and whether or not the two 3D reconstructions used were processed from a common reference. The reported resolution is shown as a black line. A curve is displayed for the half-bit criterion in addition to lines showing the 0.143 gold standard cut-off and 0.5 cut-off.

### 8.1 FSC

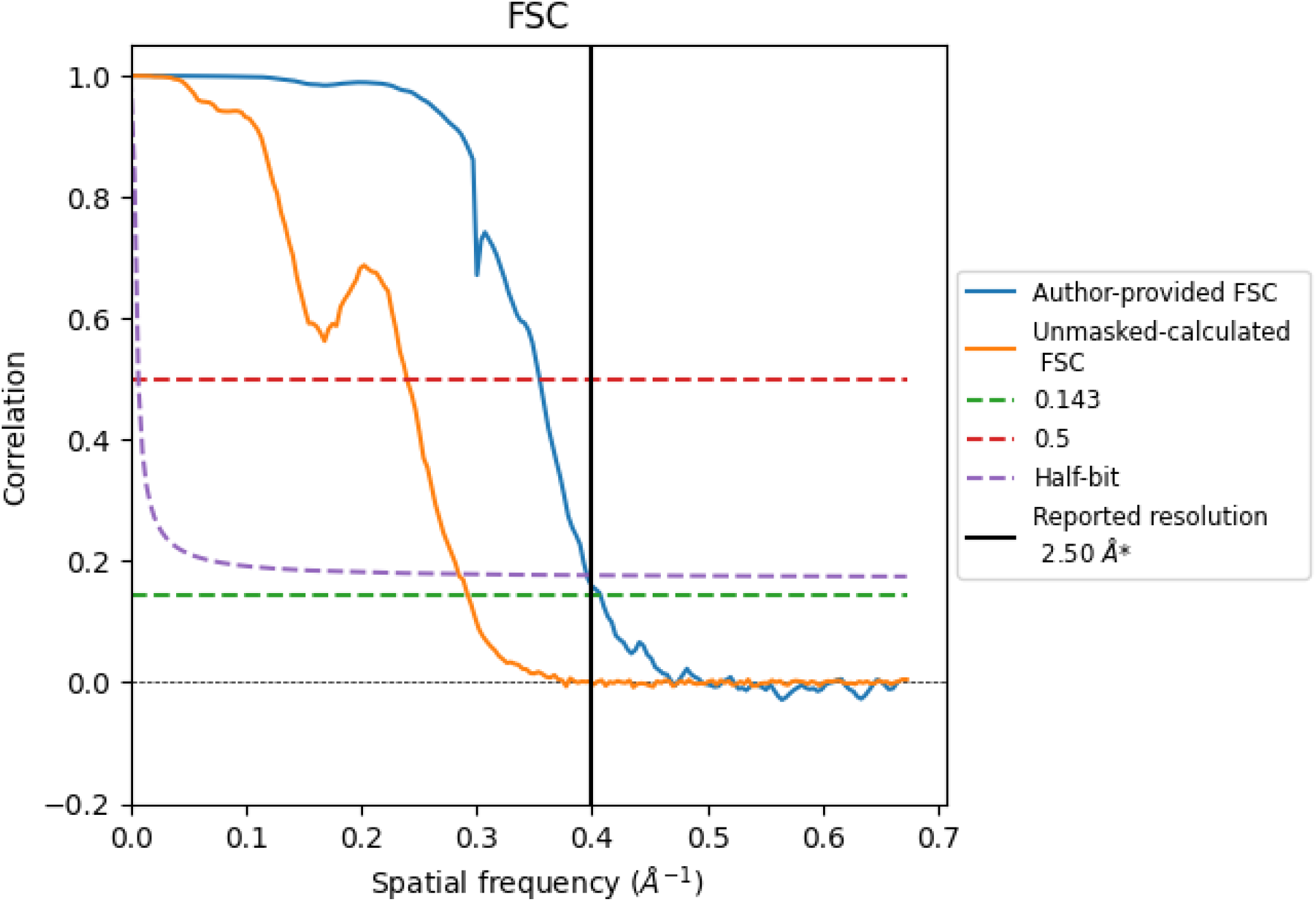

*Reported resolution corresponds to spatial frequency of 0.400 Å^-1^

### 8.2 Resolution estimates

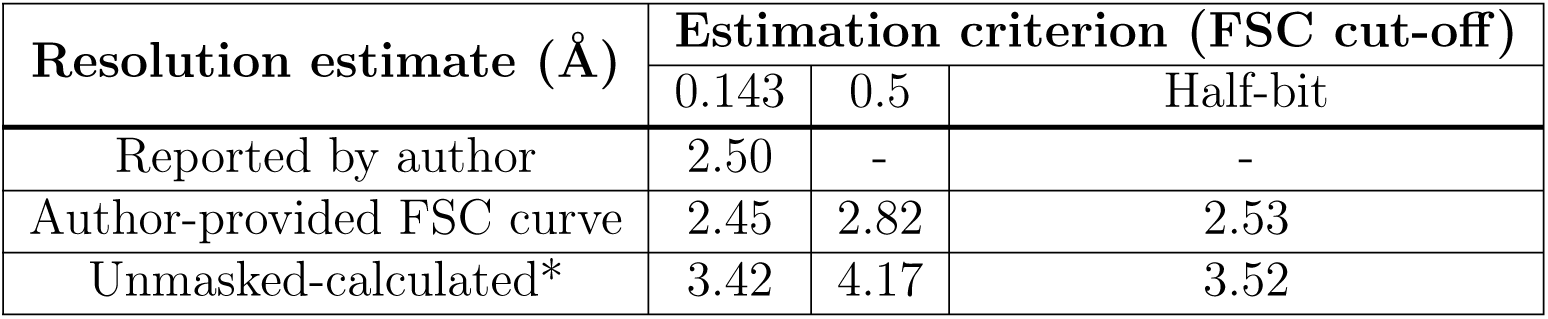

*Resolution estimate based on FSC curve calculated by comparison of deposited half-maps. The value from deposited half-maps intersecting FSC 0.143 CUT-OFF 3.42 differs from the reported value 2.5 by more than 10 %

## 9 Map-model fit

This section contains information regarding the fit between EMDB map EMD-73823 and PDB model 9Z5S. Per-residue inclusion information can be found in section 3 on page 5.

### 9.1 Map-model overlay

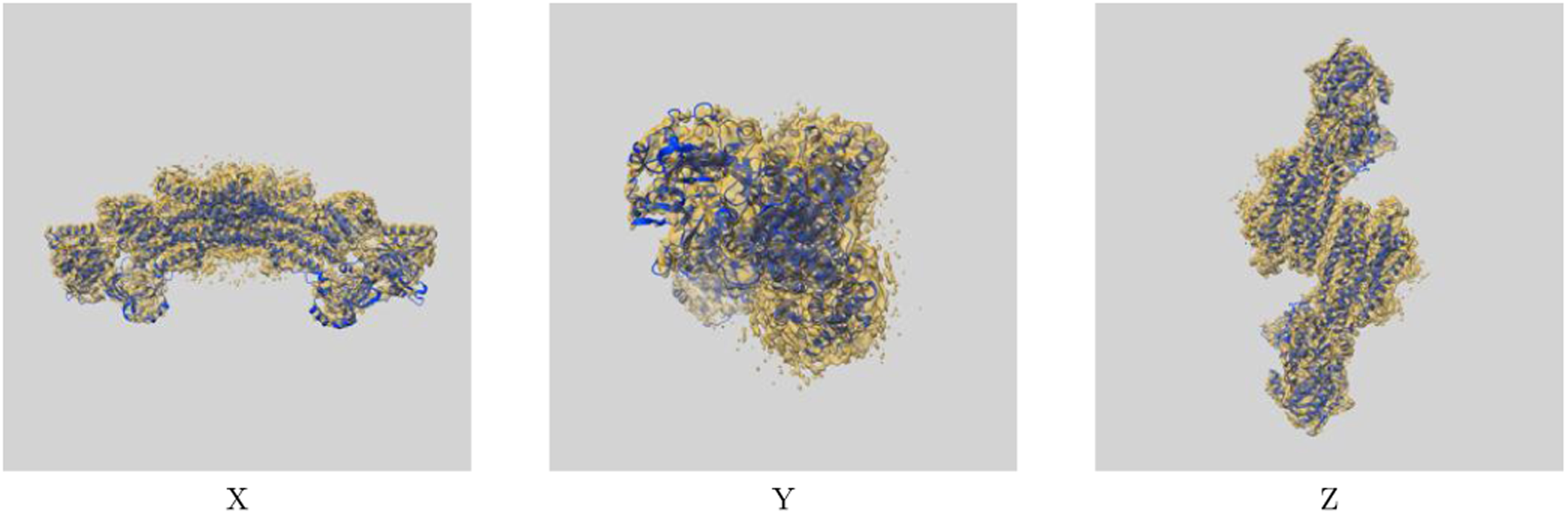

The images above show the 3D surface view of the map at the recommended contour level 0.065 at 50% transparency in yellow overlaid with a ribbon representation of the model coloured in blue. These images allow for the visual assessment of the quality of fit between the atomic model and the map.

### 9.2 Q-score mapped to coordinate model

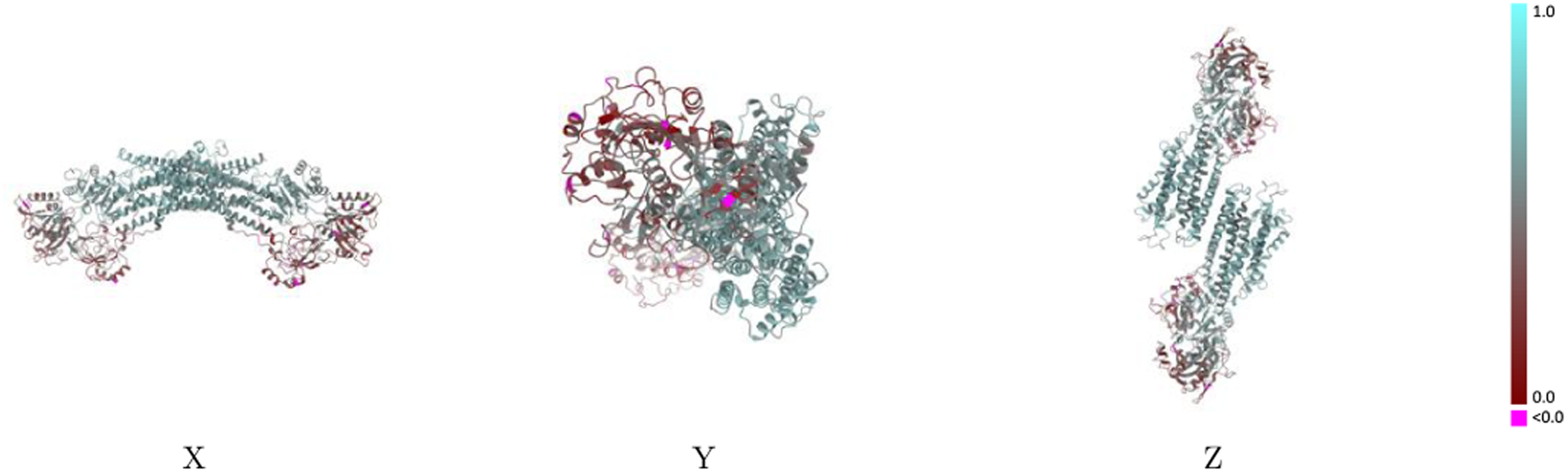

The images above show the model with each residue coloured according its Q-score. This shows their resolvability in the map with higher Q-score values reflecting better resolvability. Please note: Q-score is calculating the resolvability of atoms, and thus high values are only expected at resolutions at which atoms can be resolved. Low Q-score values may therefore be expected for many entries.

### 9.3 Atom inclusion mapped to coordinate model

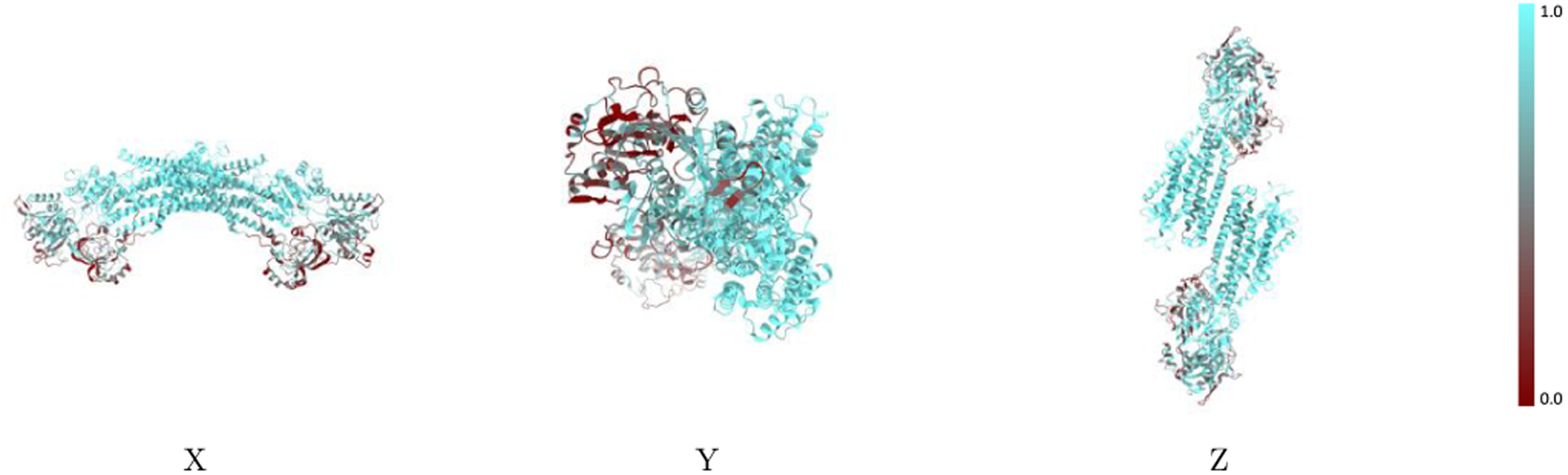

The images above show the model with each residue coloured according to its atom inclusion. This shows to what extent they are inside the map at the recommended contour level (0.065).

### 9.4 Atom inclusion

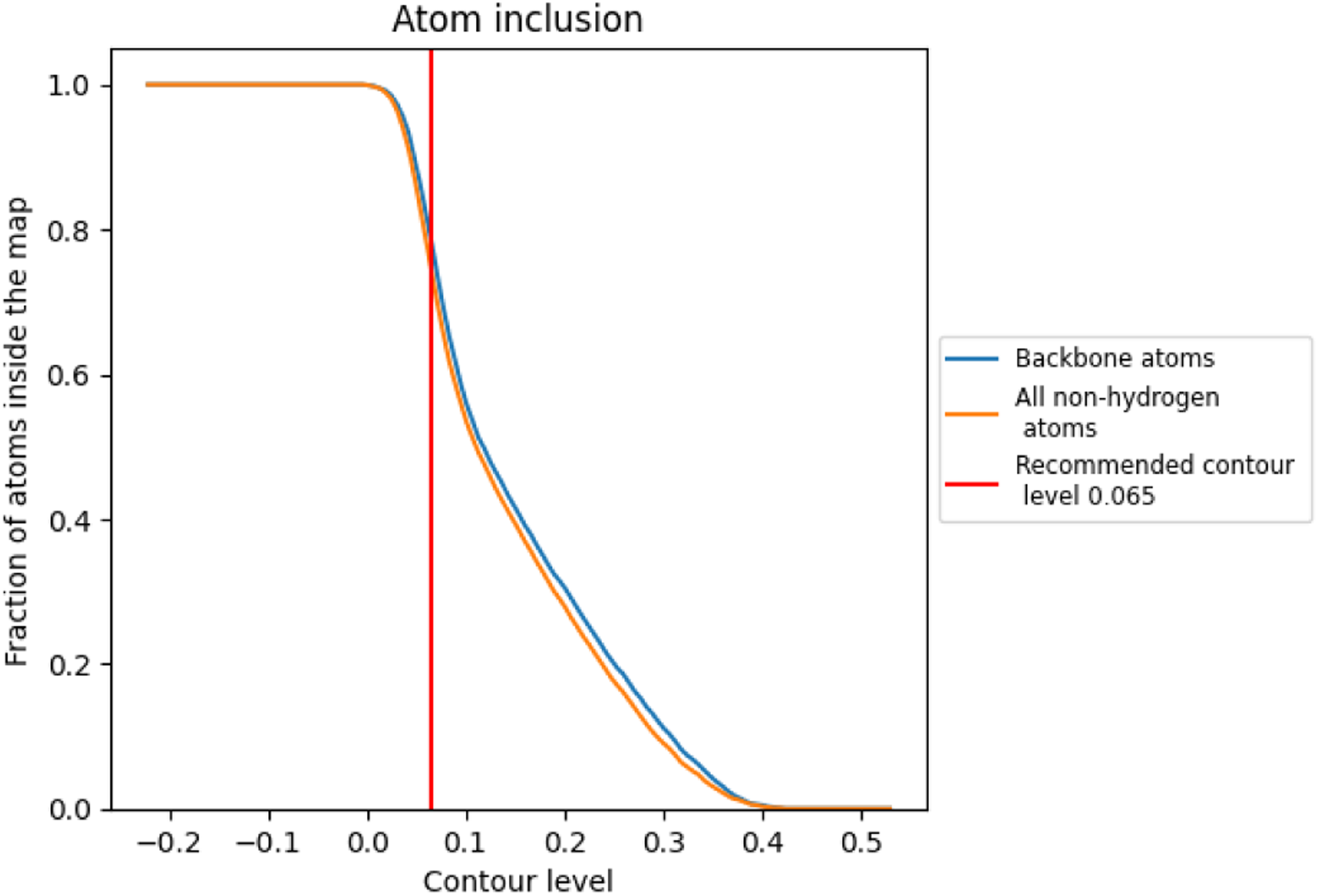

At the recommended contour level, 79% of all backbone atoms, 75% of all non-hydrogen atoms, are inside the map.

### 9.5 Map-model fit summary

The table lists the average atom inclusion at the recommended contour level (0.065) and Q-score for the entire model and for each chain.

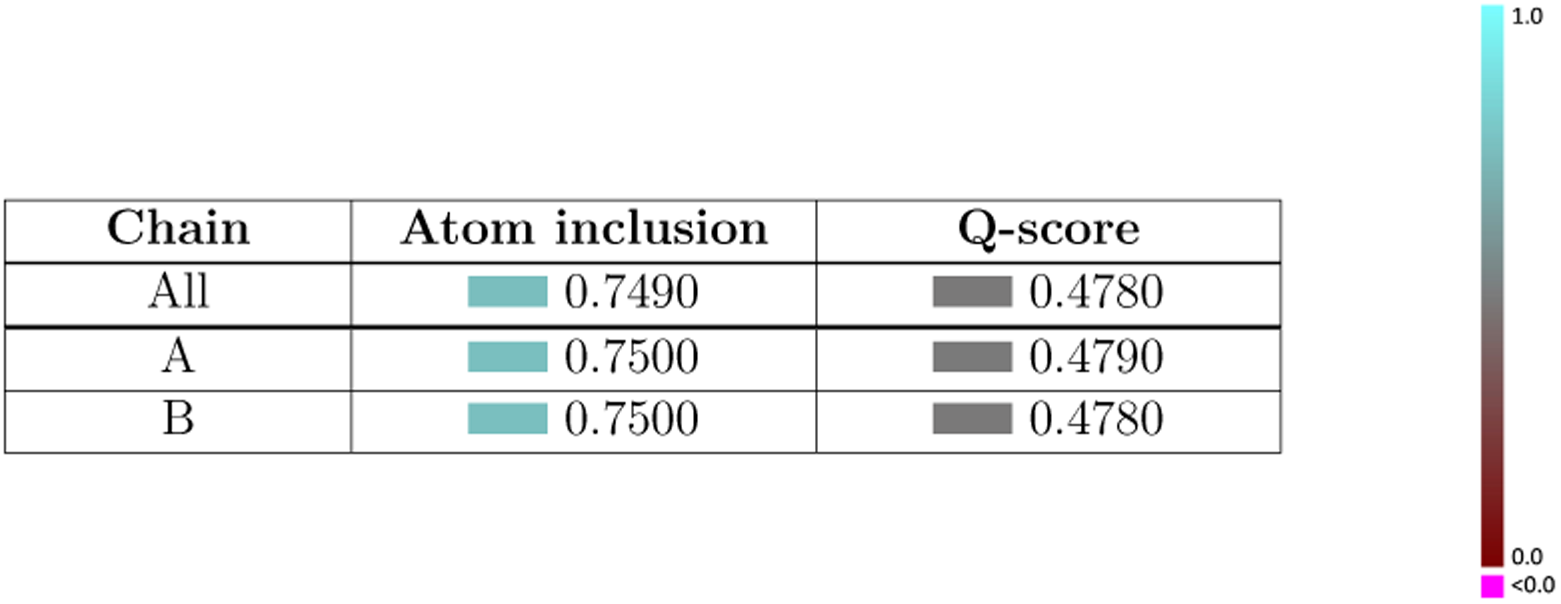

---

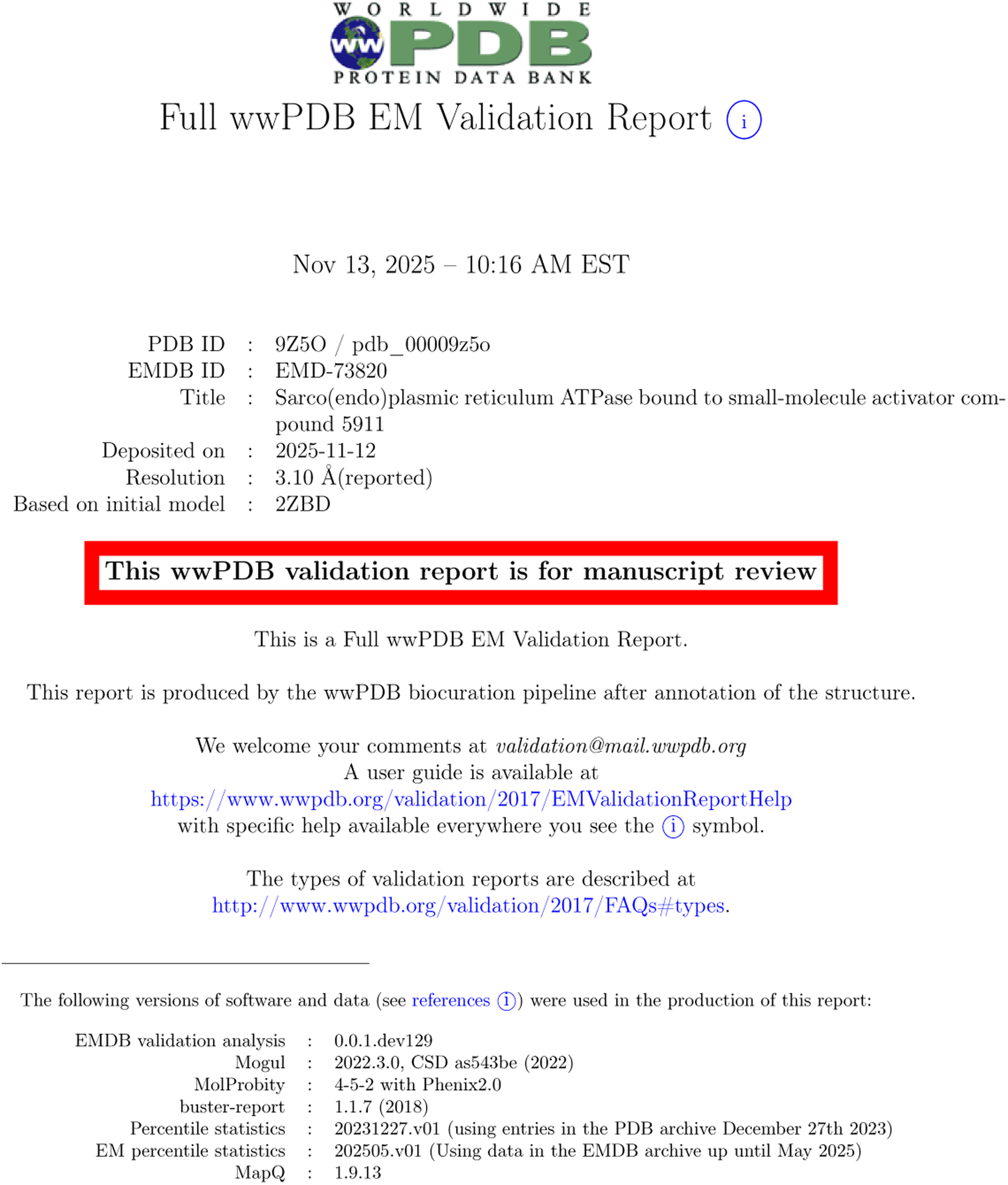

## 1 Overall quality at a glance

The following experimental techniques were used to determine the structure:

*ELECTRON MICROSCOPY*

The reported resolution of this entry is 3.10 Å.

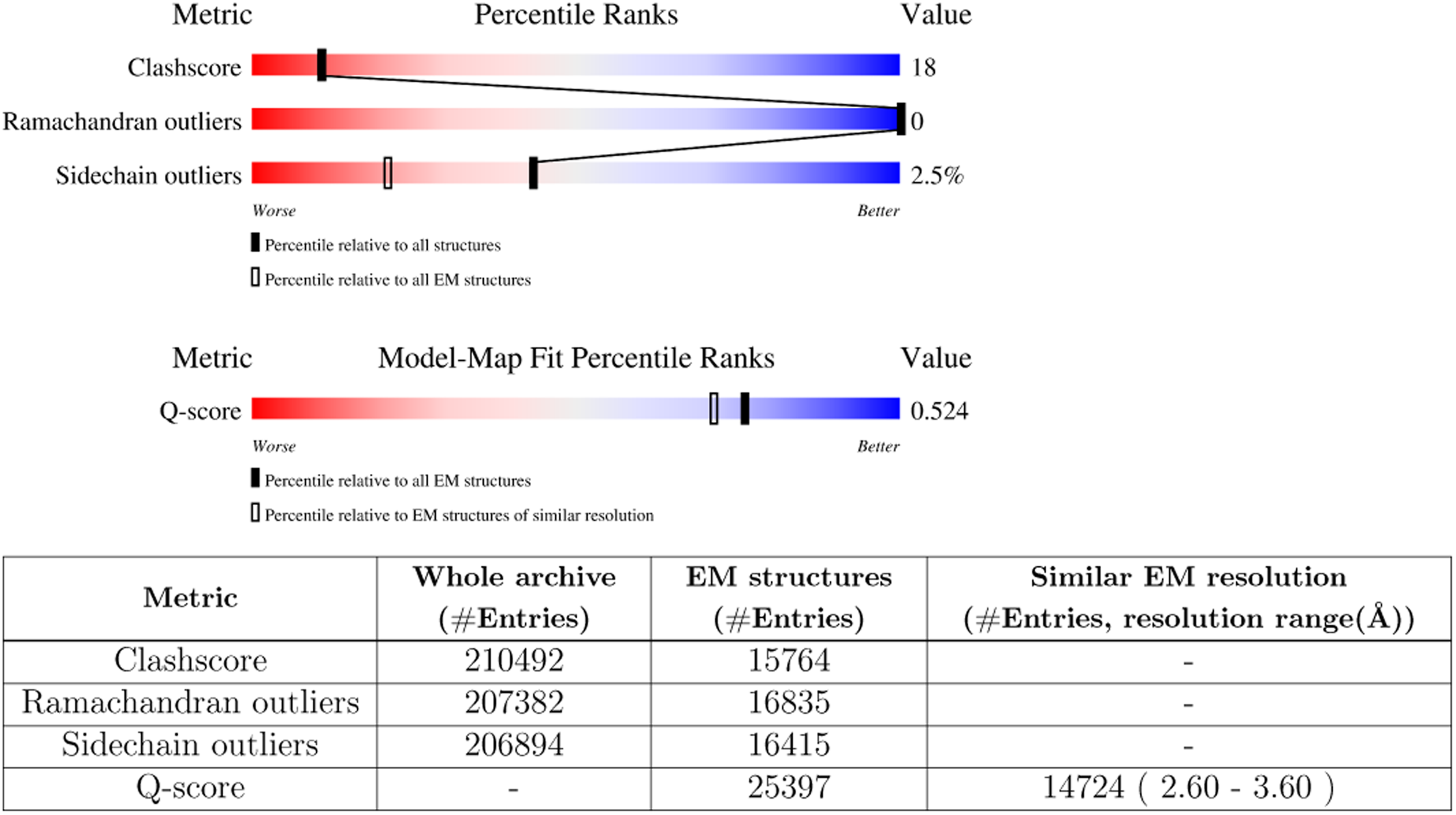

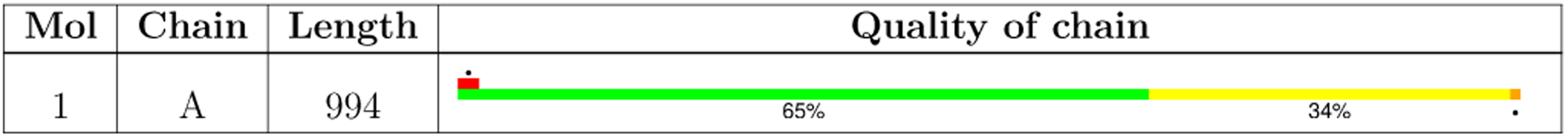

## 2 Entry composition

There are 3 unique types of molecules in this entry. The entry contains 7717 atoms, of which 26 are hydrogens and 0 are deuteriums.

- Molecule 1 is a protein called Sarcoplasmic/endoplasmic reticulum calcium ATPase 1.

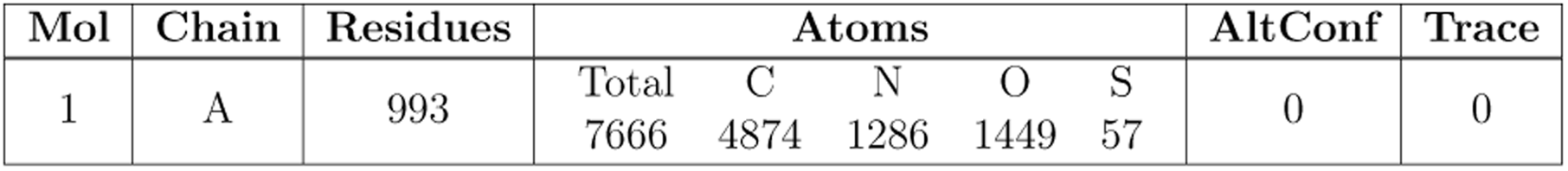

- Molecule 2 is CALCIUM ION (CCD ID: CA) (formula: Ca) (labeled as “Ligand of Interest” by depositor).

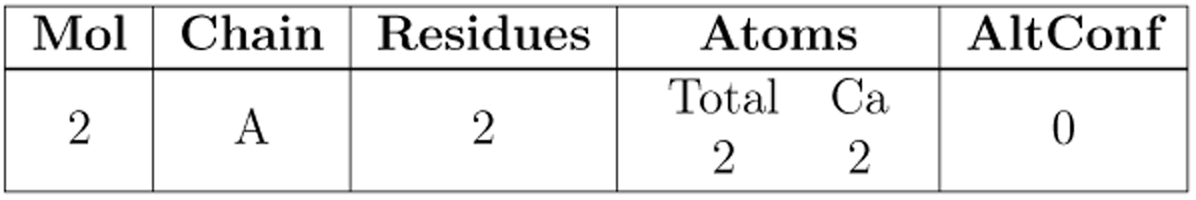

- Molecule 3 is 5-{2-[(5-phenylpentyl)oxy]ethyl}-2,3-dihydro-1-benzofuran (CCD ID: A1C01) (formula: C_21_H_26_O_2_) (labeled as “Ligand of Interest” by depositor).

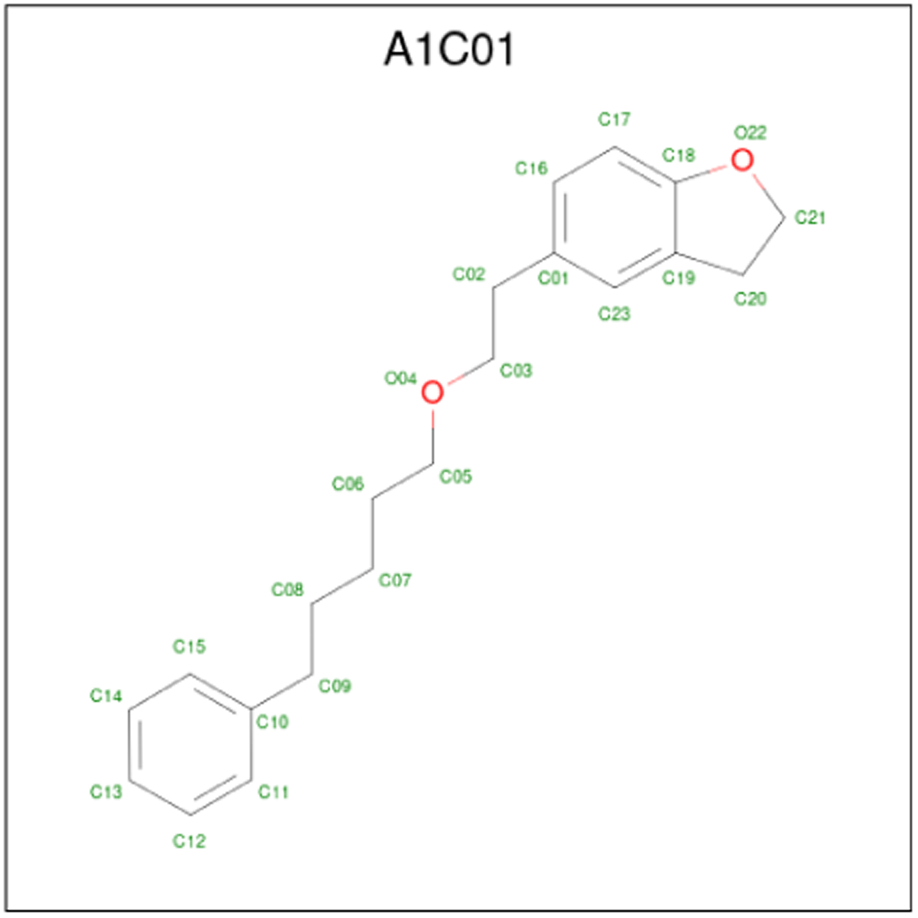

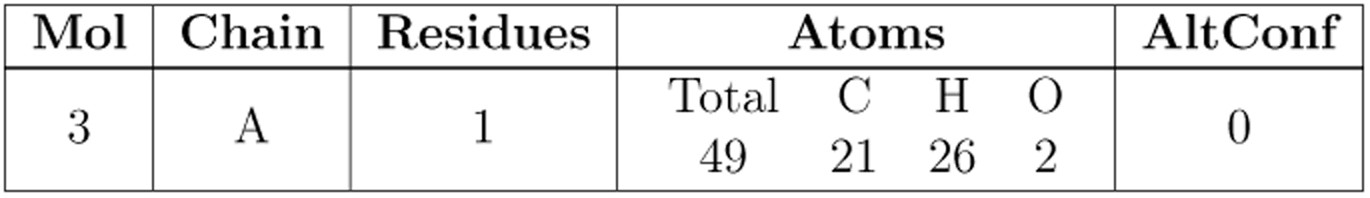

## 3 Residue-property plots

- Molecule 1: Sarcoplasmic/endoplasmic reticulum calcium ATPase 1

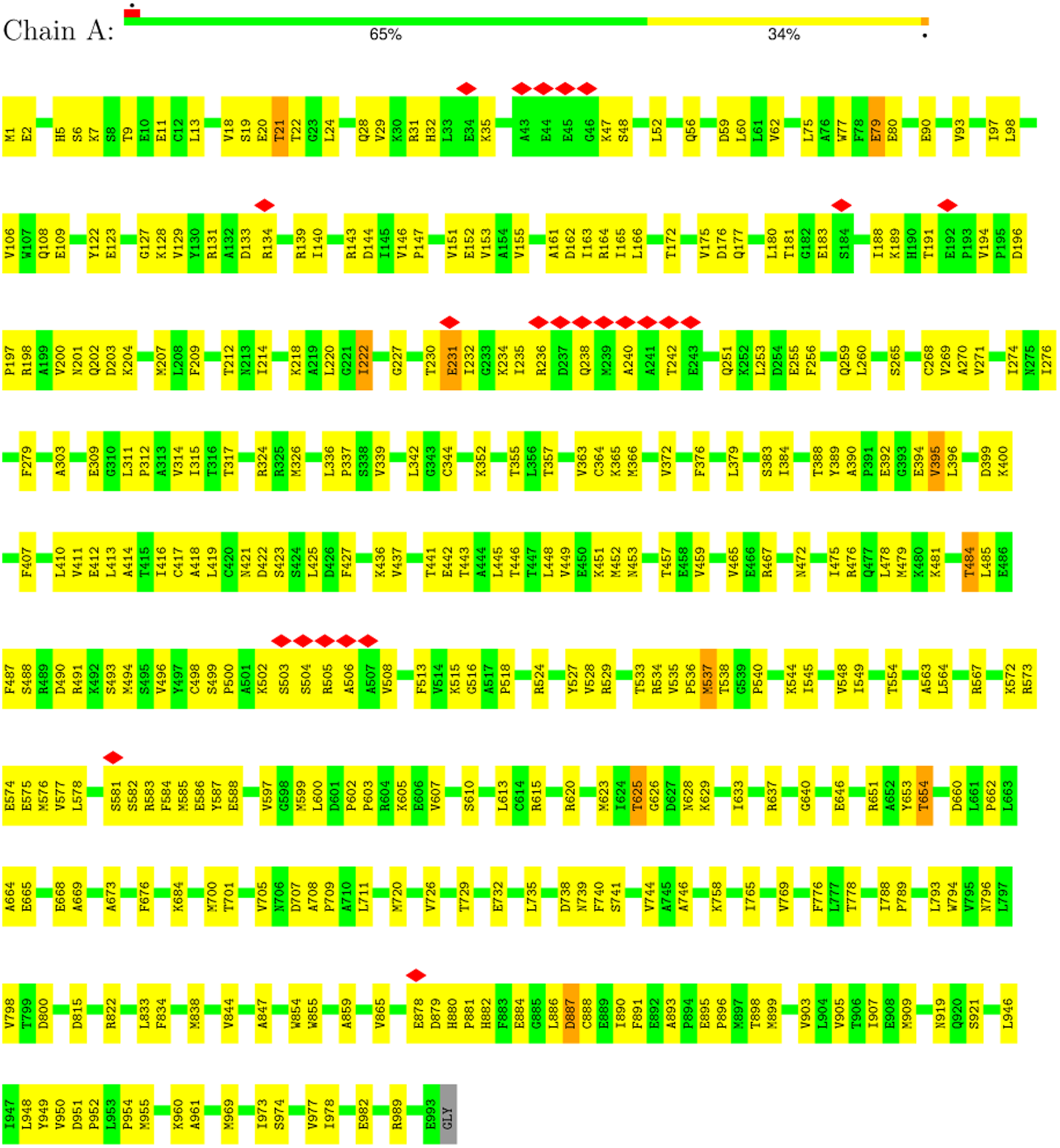

## 4 Experimental information

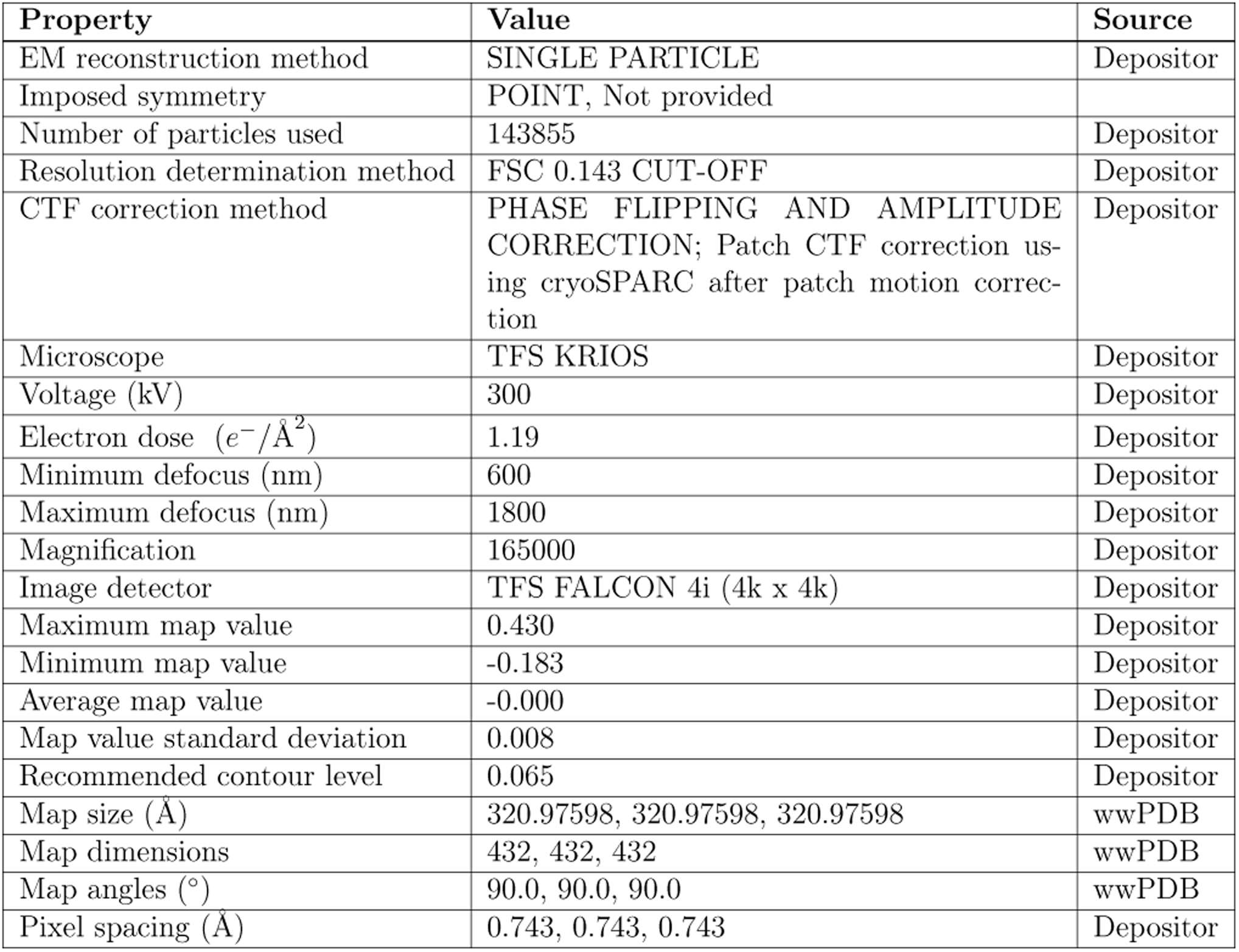

## 5 Model quality

### 5.1 Standard geometry

Bond lengths and bond angles in the following residue types are not validated in this section: A1C01, CA

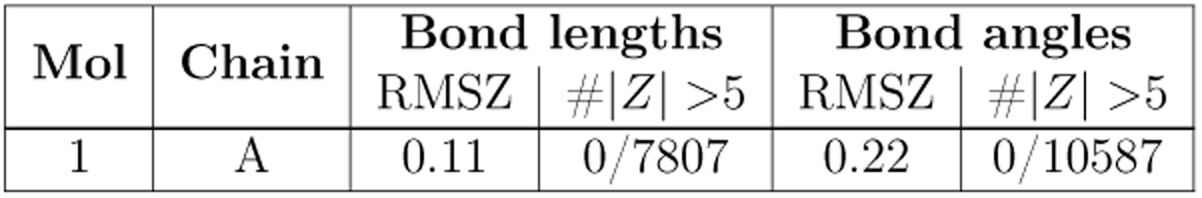

There are no bond length outliers.

There are no bond angle outliers.

There are no chirality outliers.

There are no planarity outliers.

### 5.2 Too-close contacts

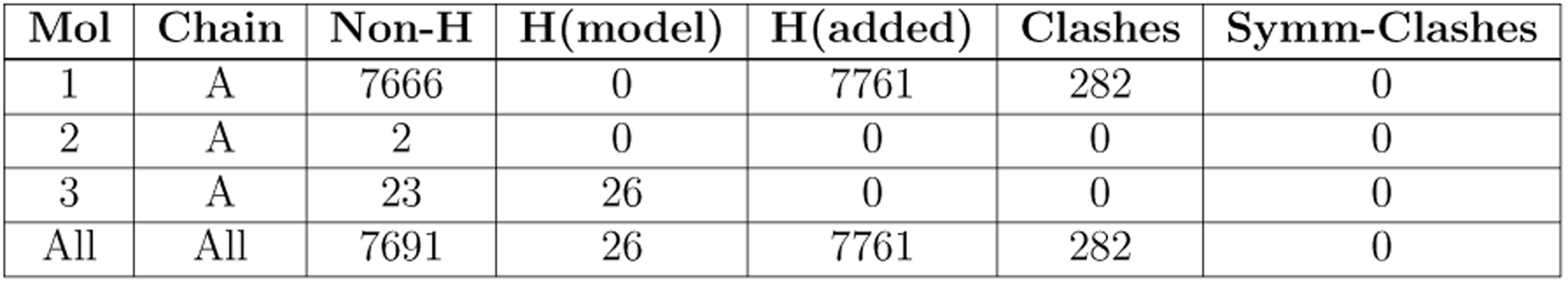

The all-atom clashscore is defined as the number of clashes found per 1000 atoms (including hydrogen atoms). The all-atom clashscore for this structure is 18.

All (282) close contacts within the same asymmetric unit are listed below, sorted by their clash magnitude.

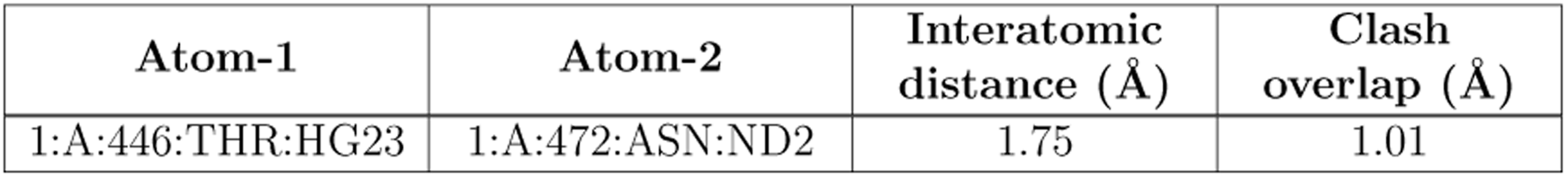
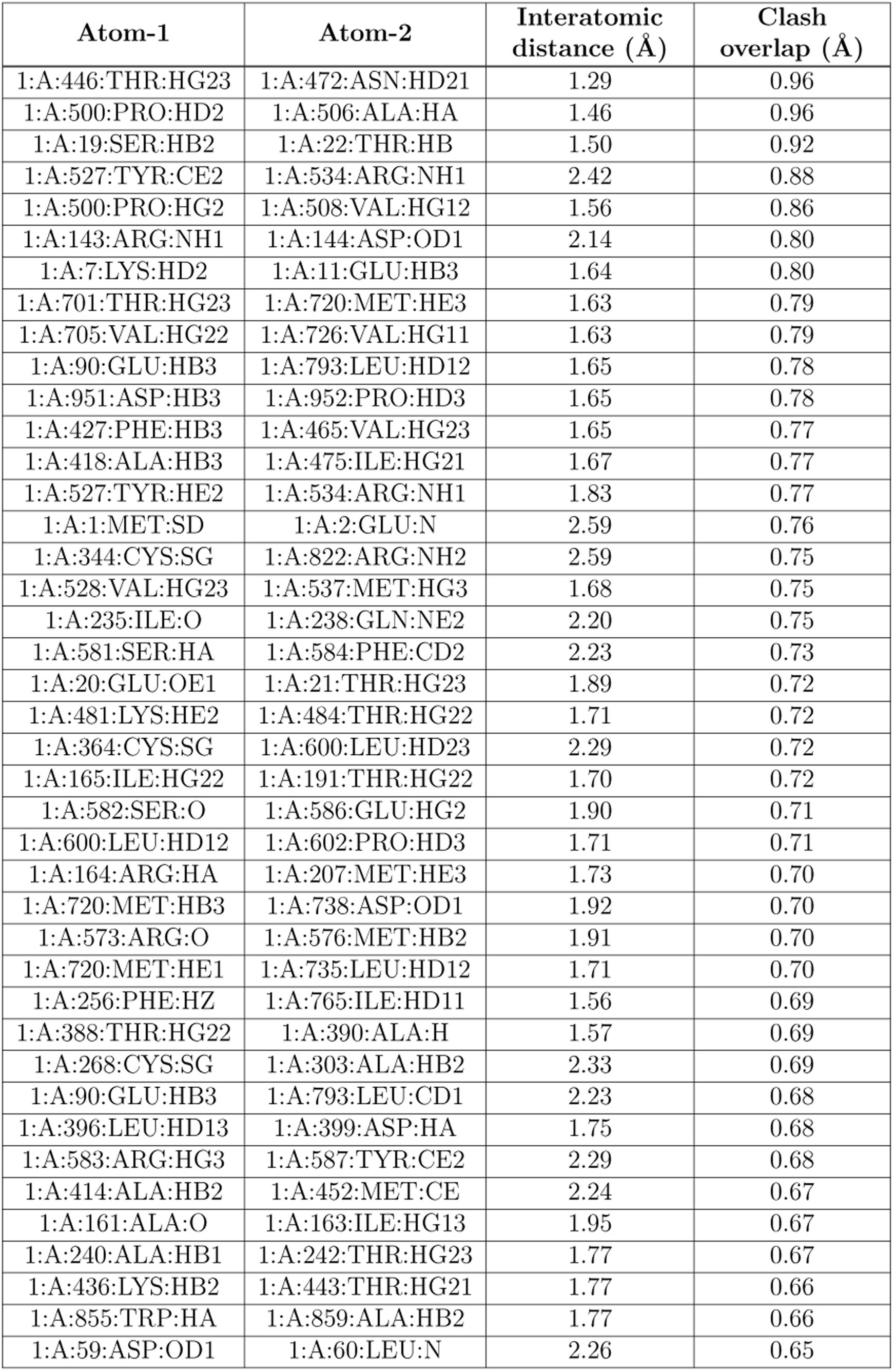
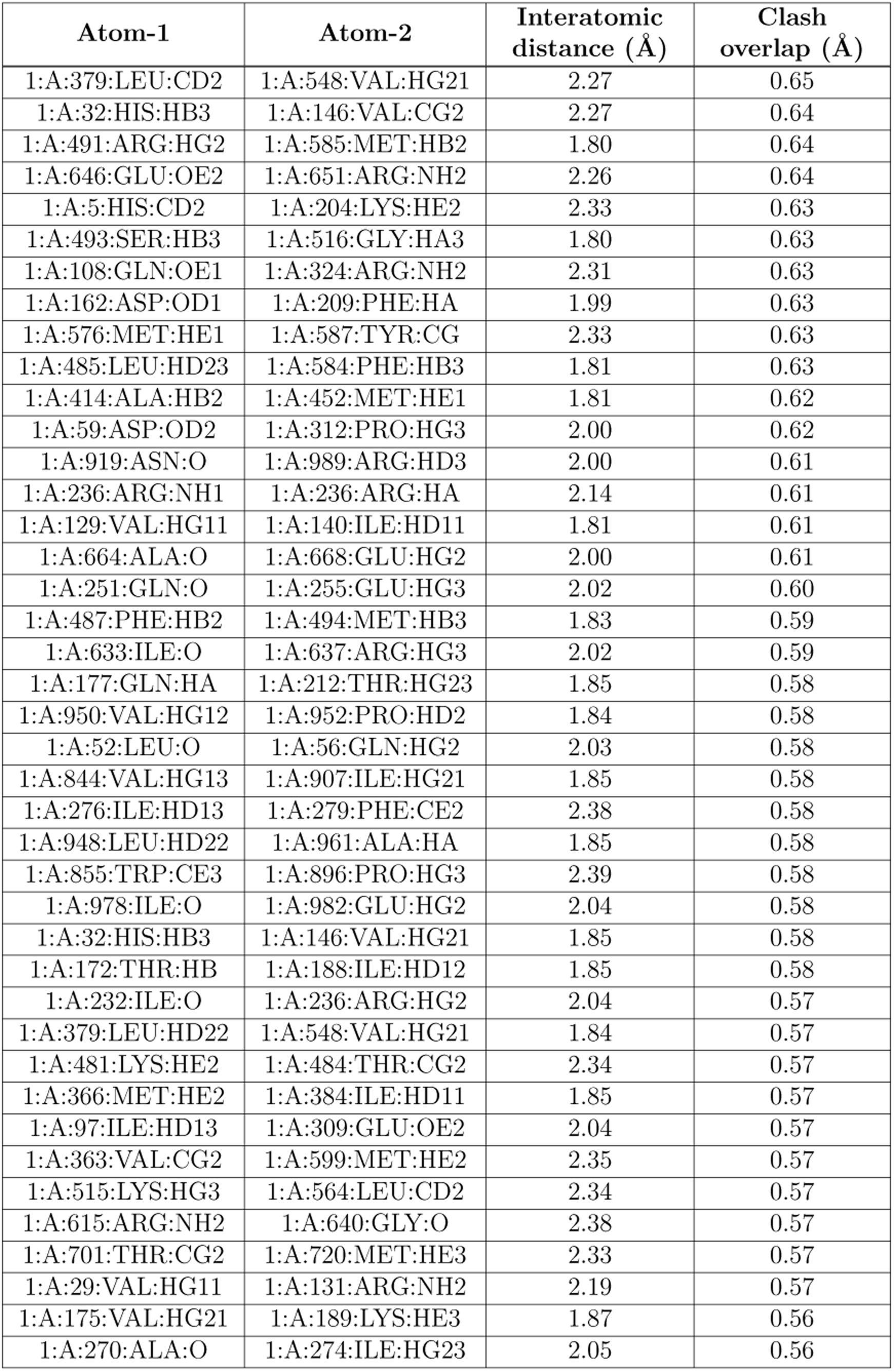
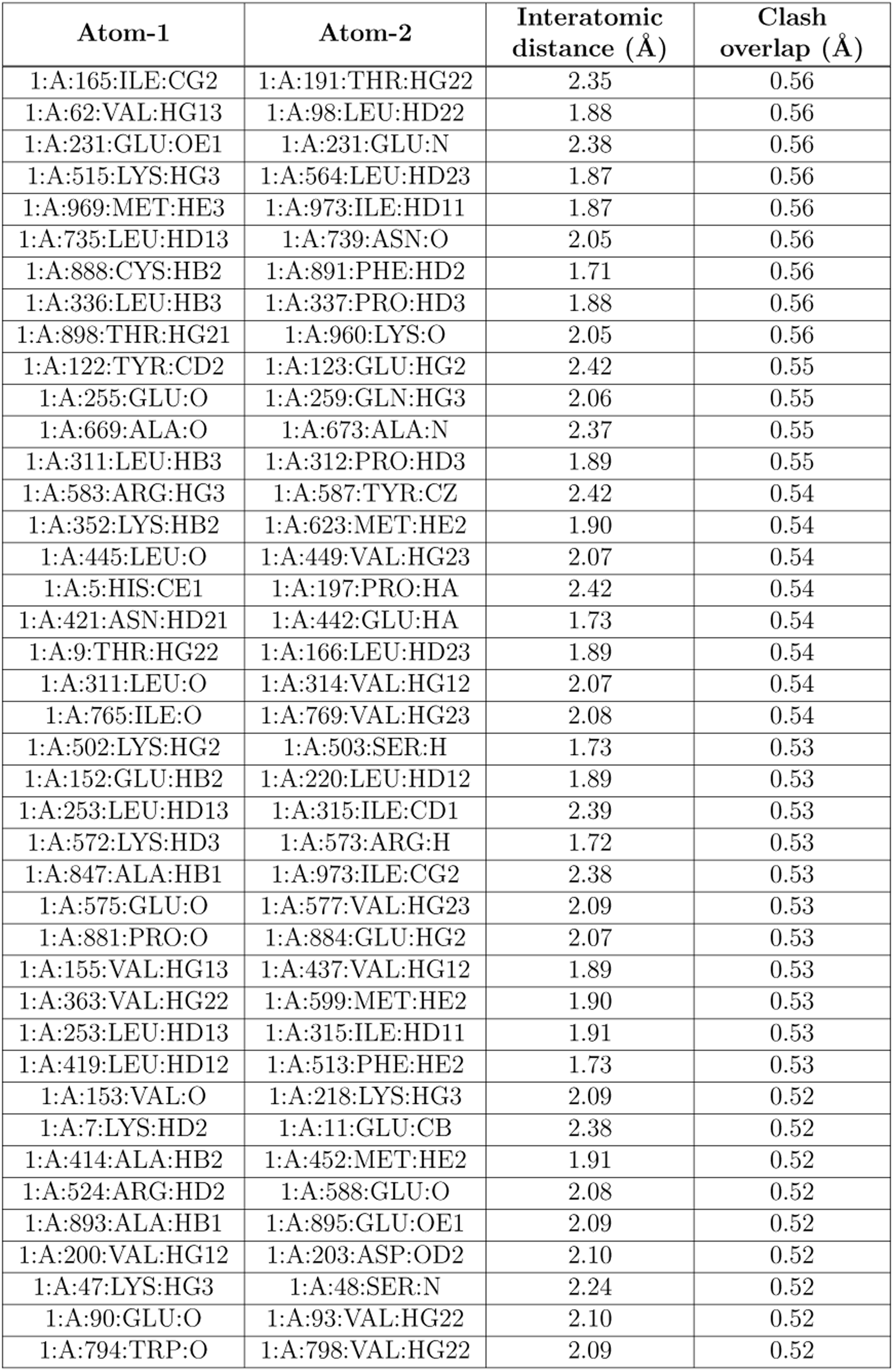
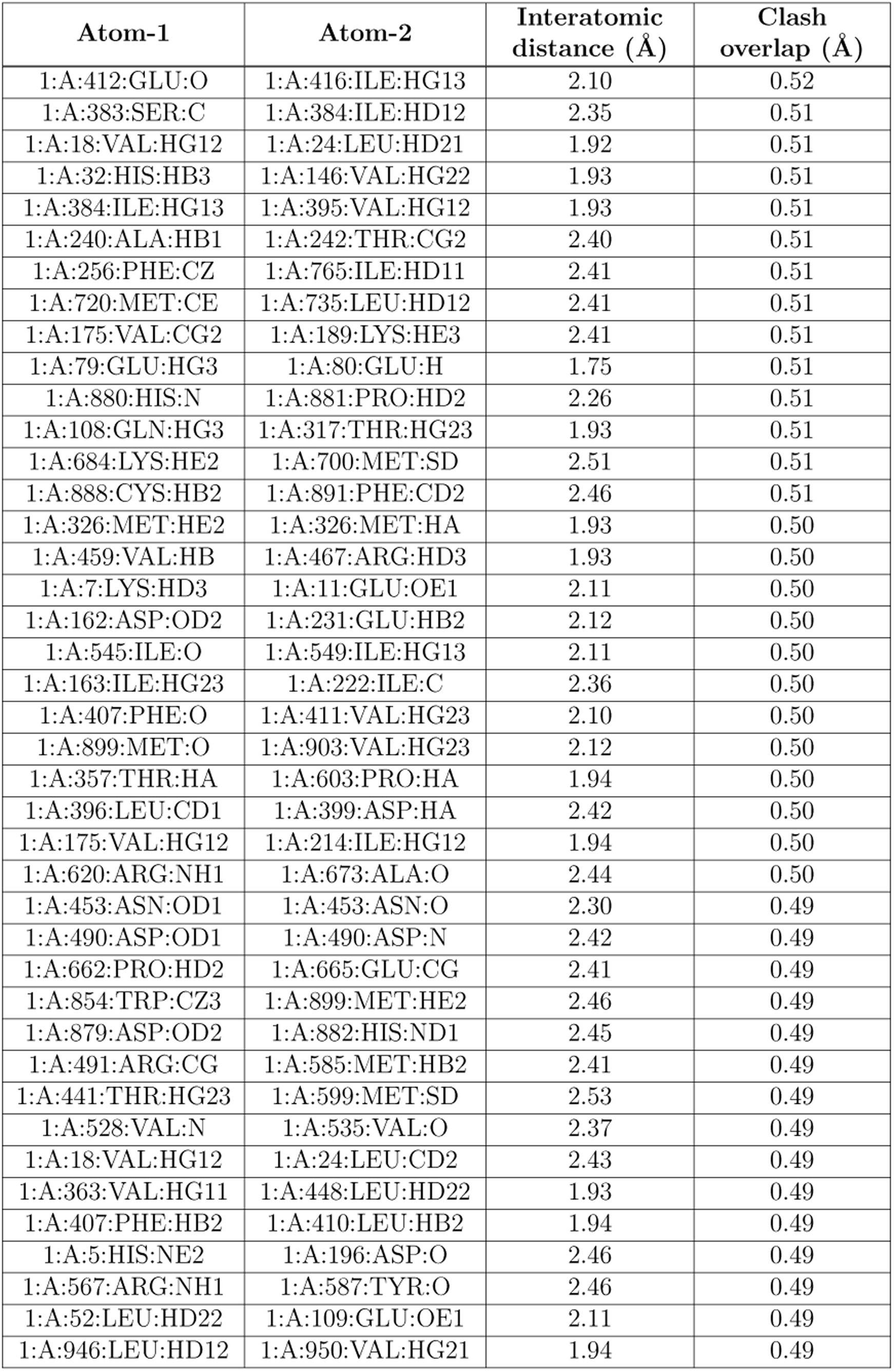
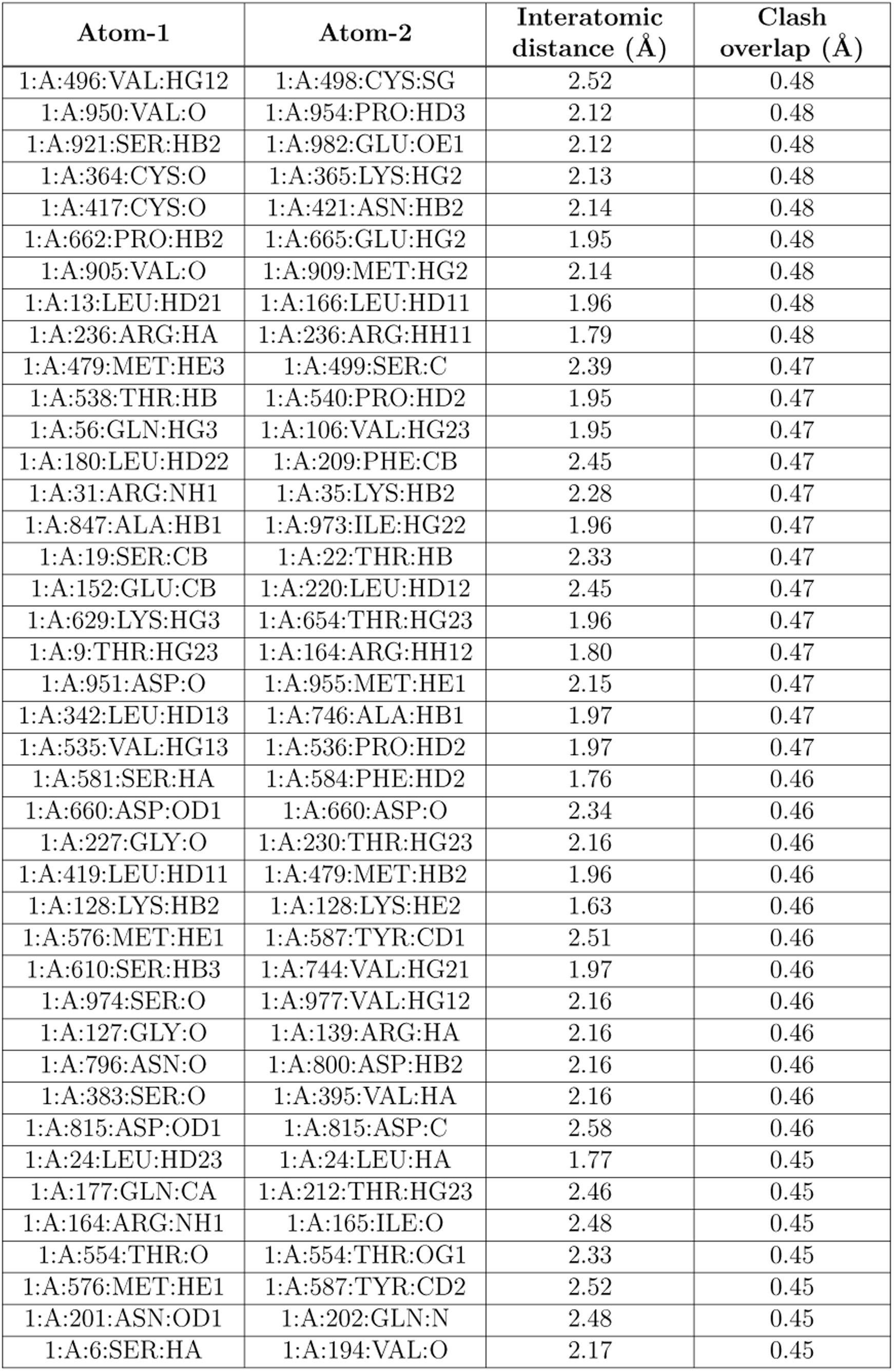
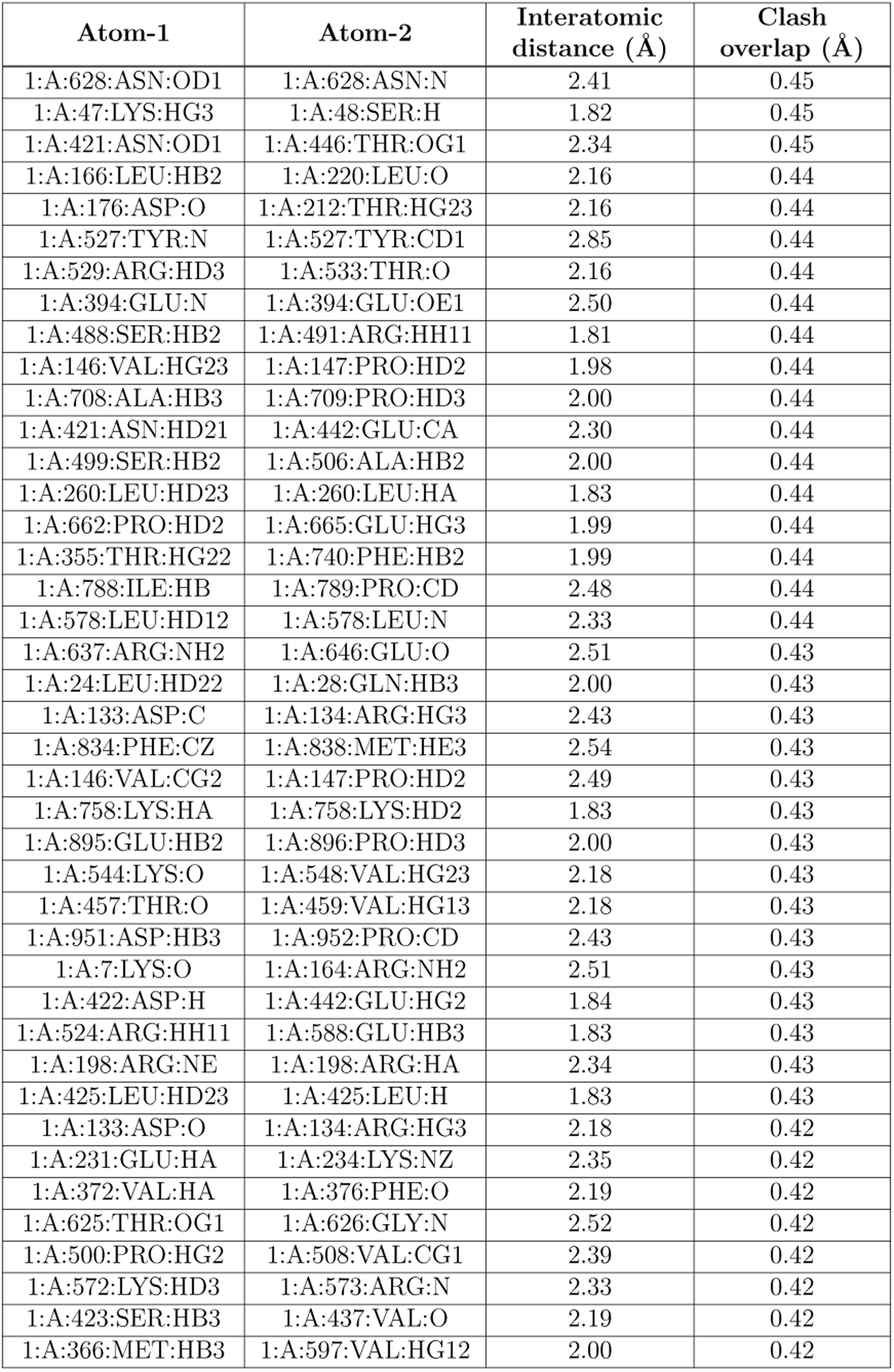
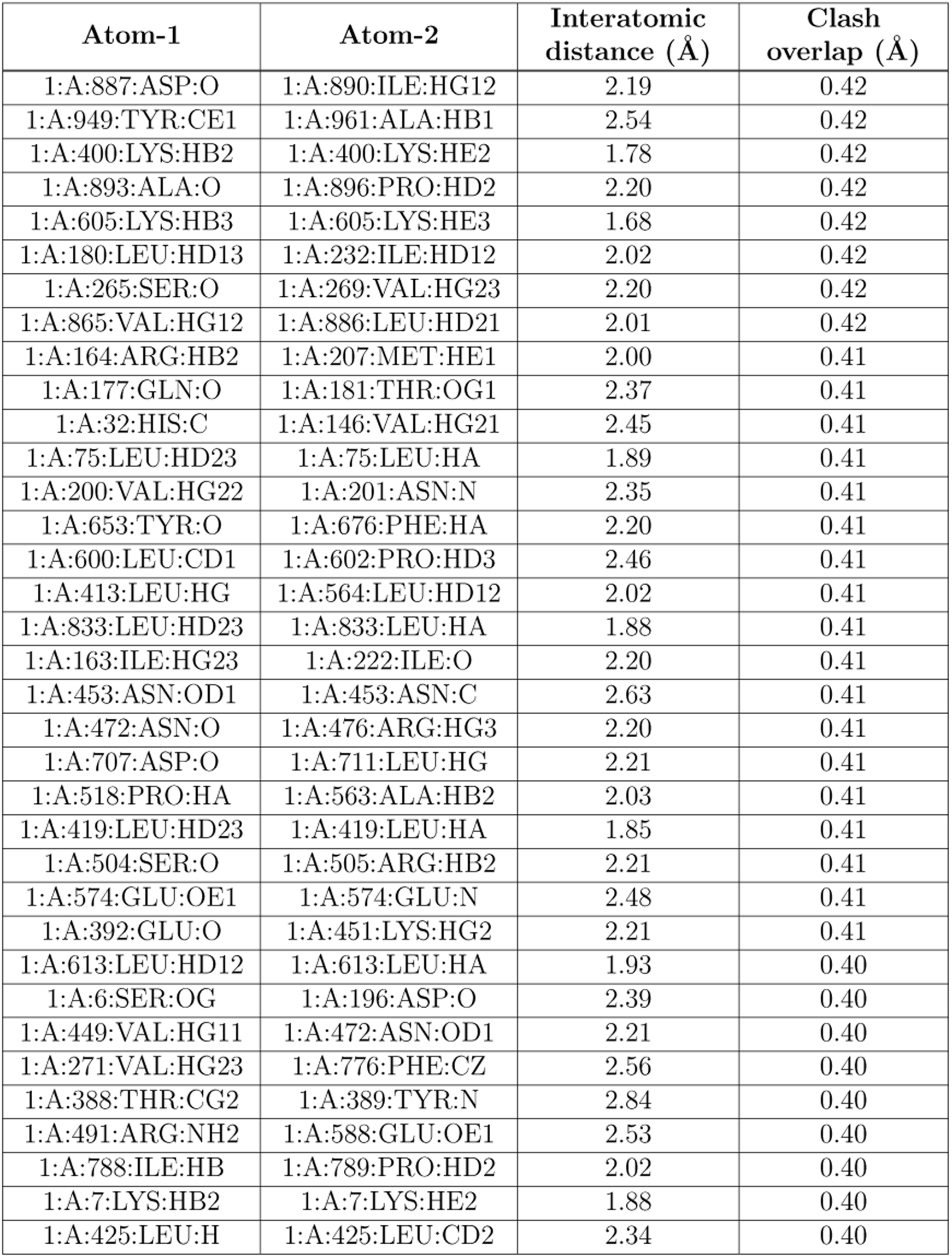

### 5.3 Torsion angles

#### 5.3.1 Protein backbone

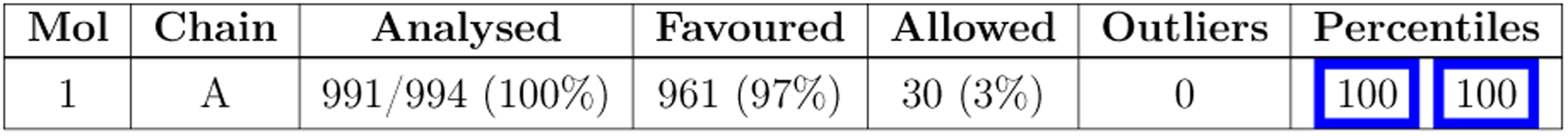

There are no Ramachandran outliers to report.

#### 5.3.2 Protein sidechains

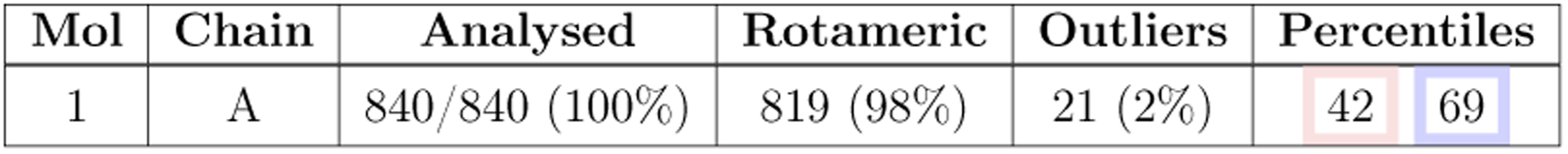

All (21) residues with a non-rotameric sidechain are listed below:

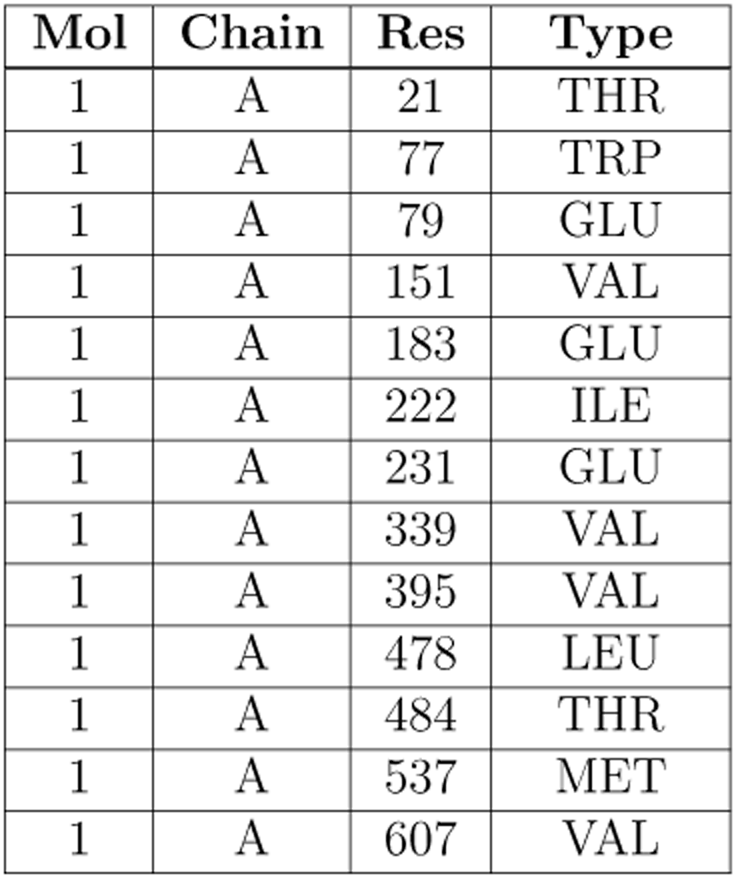
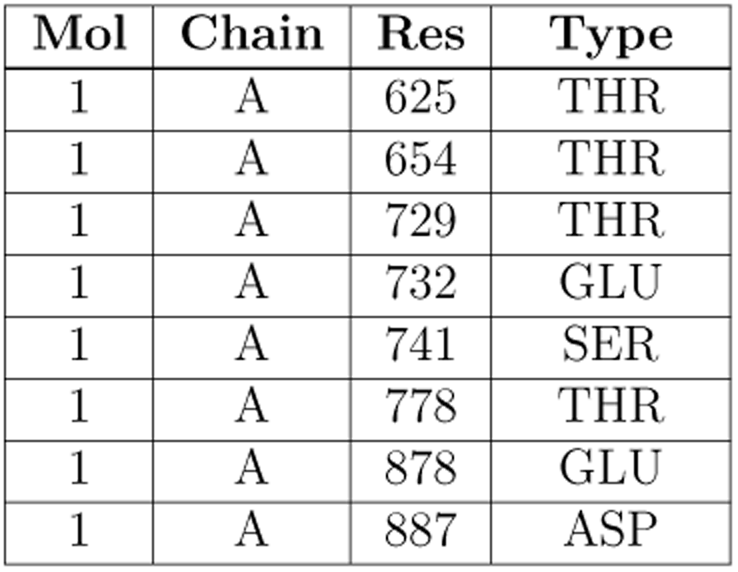

Sometimes sidechains can be flipped to improve hydrogen bonding and reduce clashes. All (2) such sidechains are listed below:

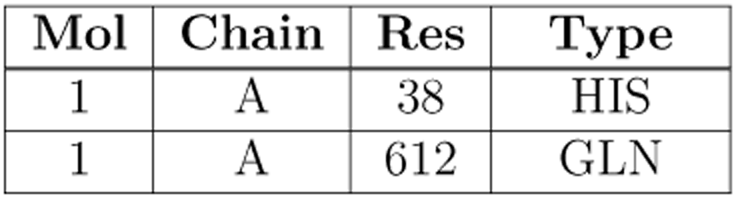

#### 5.3.3 RNA

There are no RNA molecules in this entry.

### 5.4 Non-standard residues in protein, DNA, RNA chains

There are no non-standard protein/DNA/RNA residues in this entry.

### 5.5 Carbohydrates

There are no oligosaccharides in this entry.

### 5.6 Ligand geometry

Of 3 ligands modelled in this entry, 2 are monoatomic - leaving 1 for Mogul analysis.

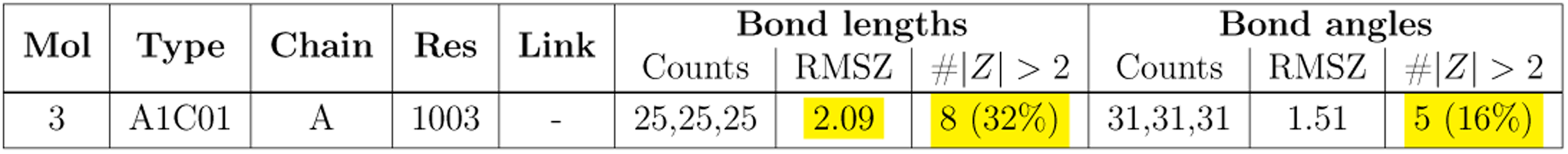

In the following table, the Chirals column lists the number of chiral outliers, the number of chiral centers analysed, the number of these observed in the model and the number defined in the Chemical Component Dictionary. Similar counts are reported in the Torsion and Rings columns. ‘-’ means no outliers of that kind were identified.

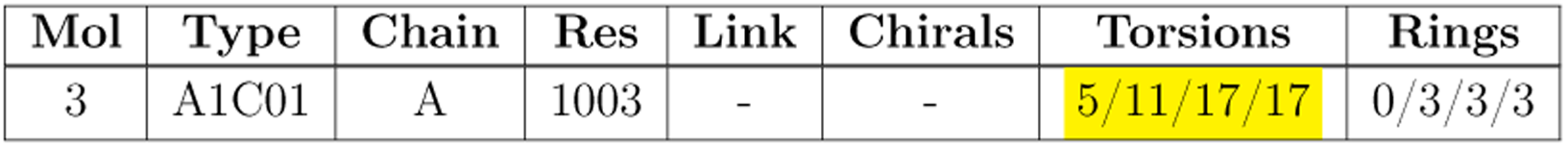

All (8) bond length outliers are listed below:

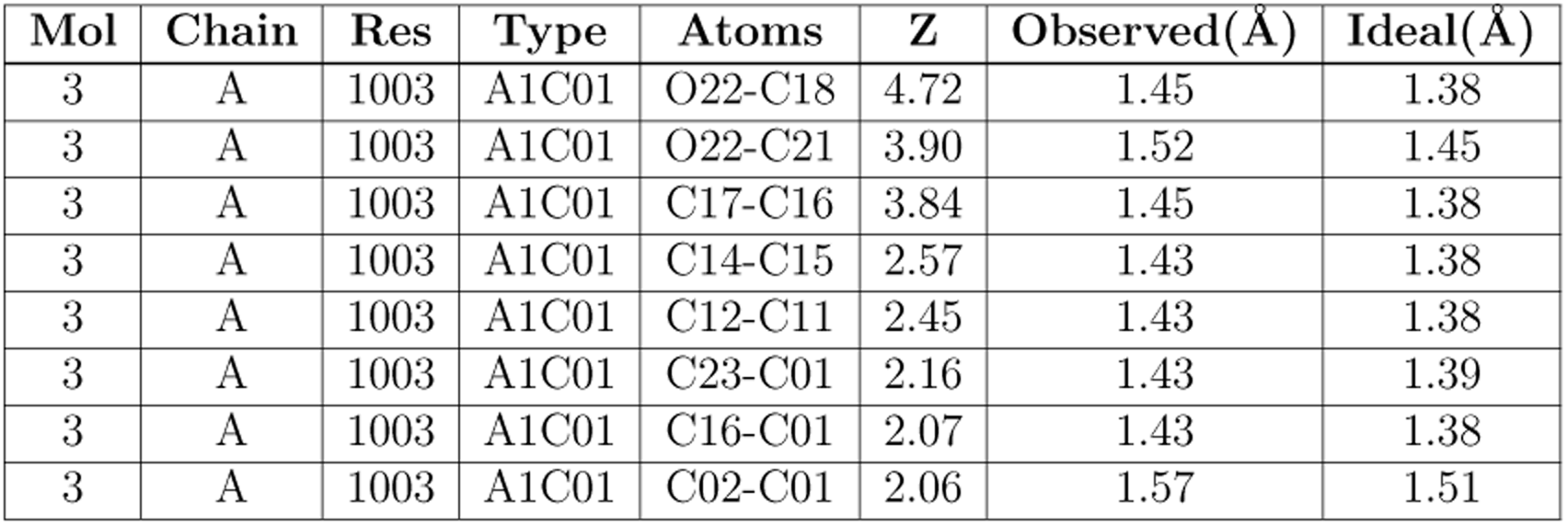

All (5) bond angle outliers are listed below:

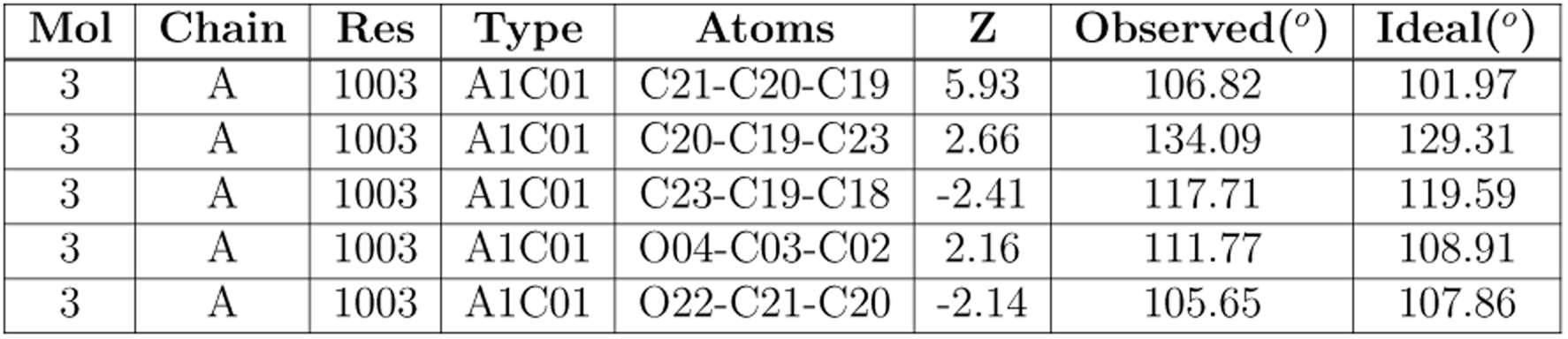

There are no chirality outliers.

All (5) torsion outliers are listed below:

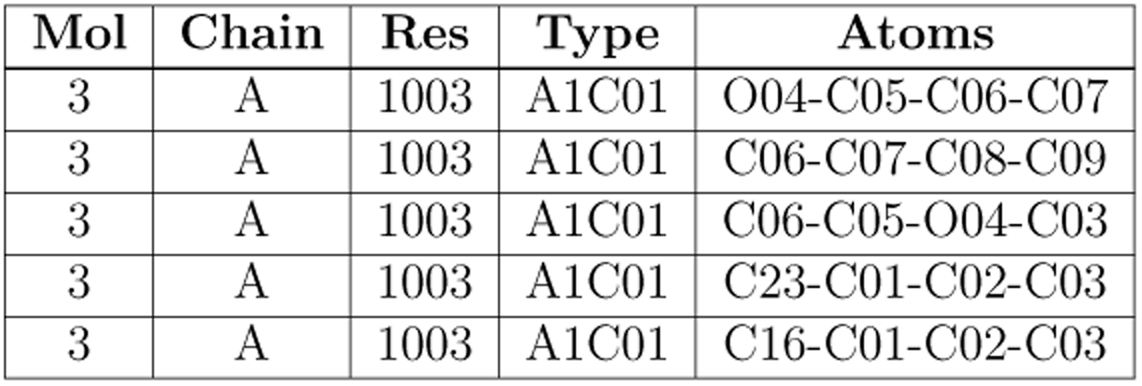

There are no ring outliers.

No monomer is involved in short contacts.

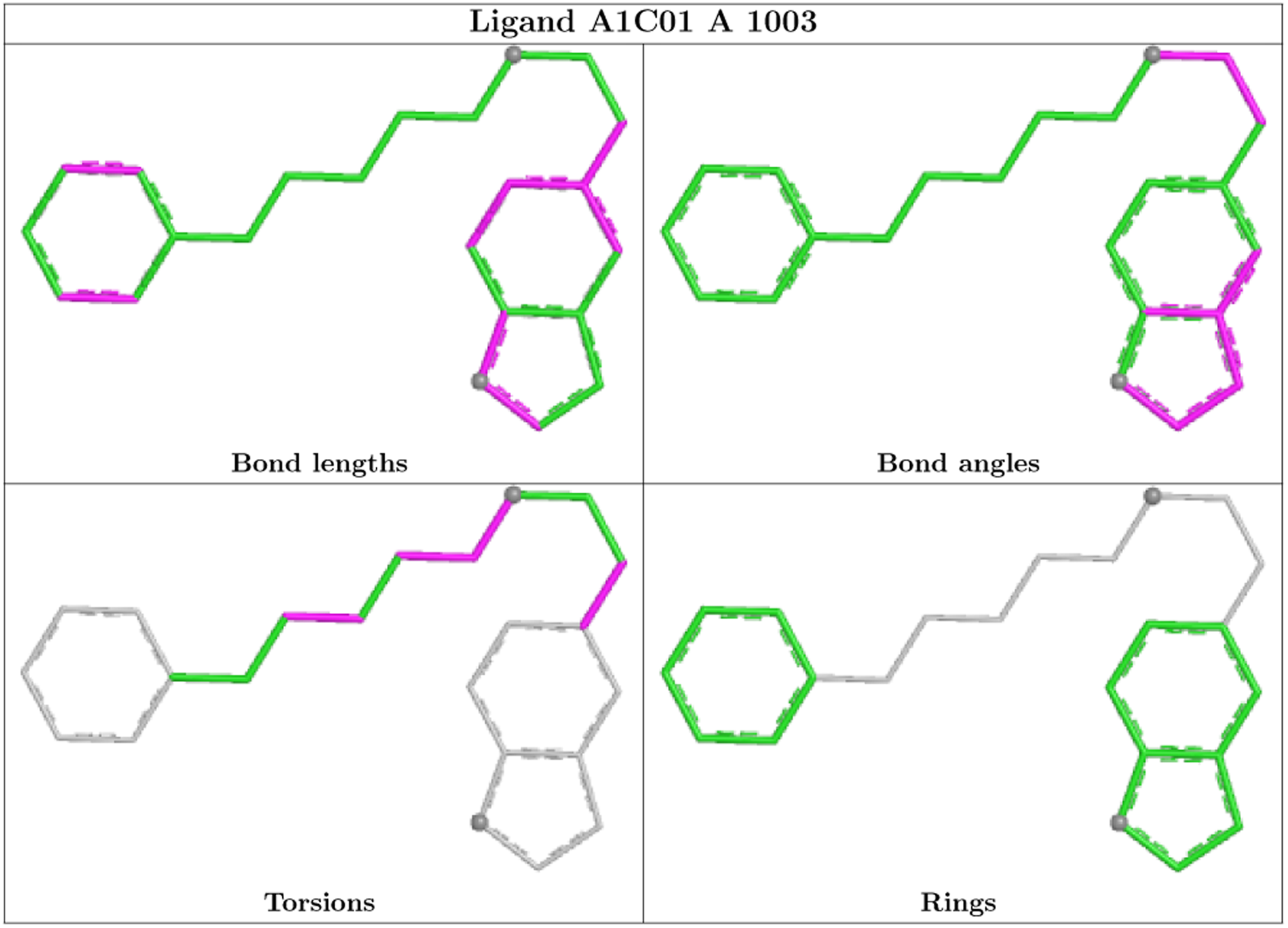

### 5.7 Other polymers

There are no such residues in this entry.

### 5.8 Polymer linkage issues

There are no chain breaks in this entry.

## 6 Map visualisation

This section contains visualisations of the EMDB entry EMD-73820. These allow visual inspection of the internal detail of the map and identification of artifacts.

### 6.1 Orthogonal projections

#### 6.1.1 Primary map

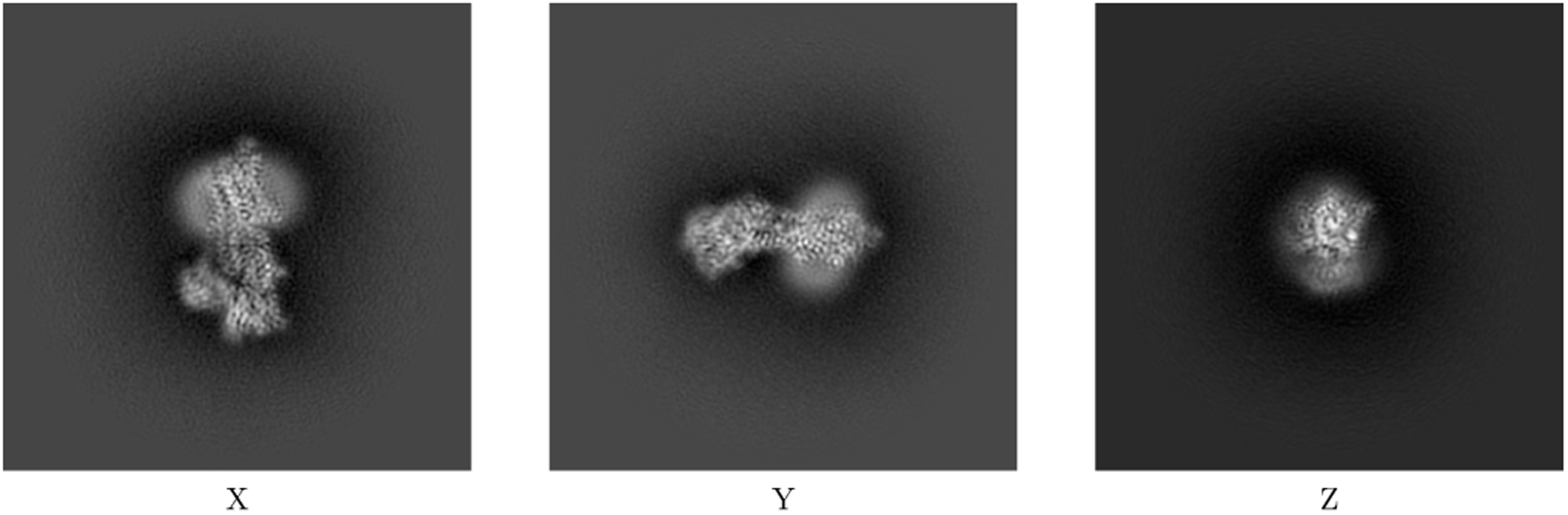

#### 6.1.2 Raw map

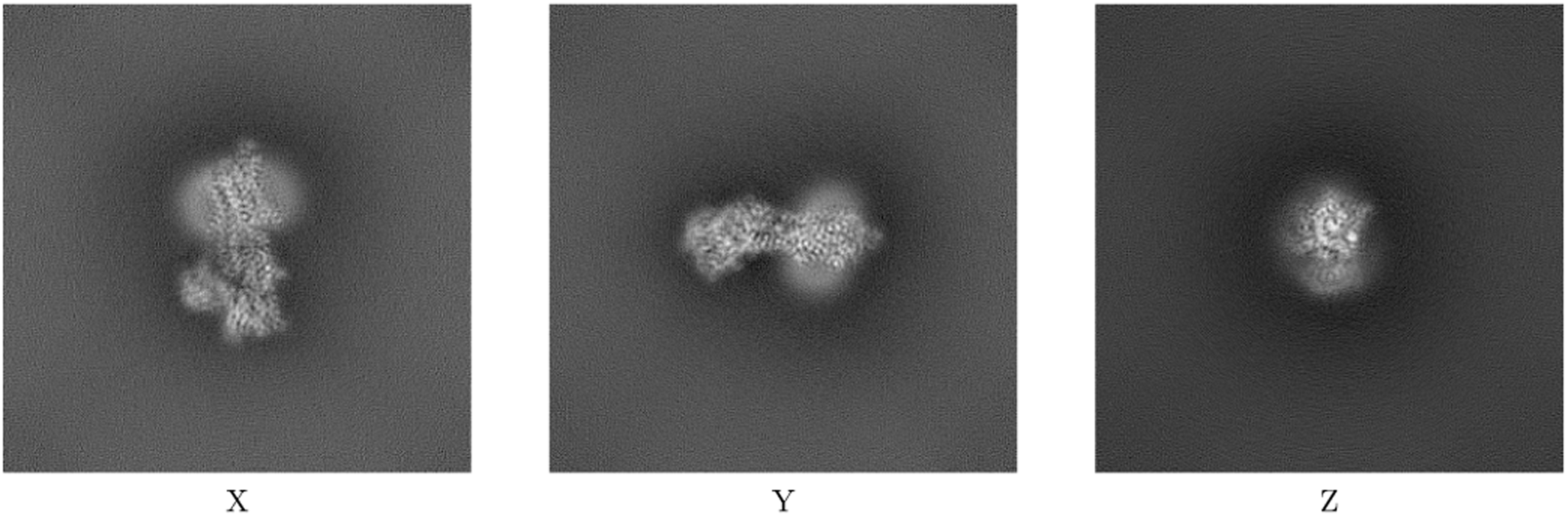

The images above show the map projected in three orthogonal directions.

### 6.2 Central slices

#### 6.2.1 Primary map

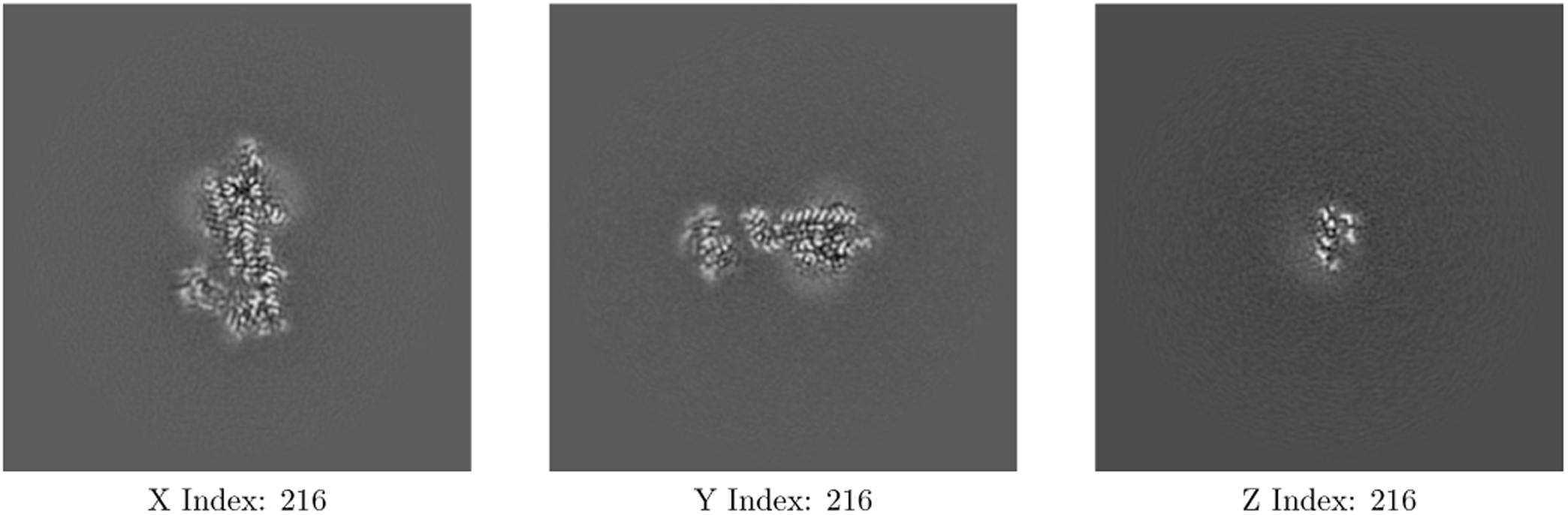

#### 6.2.2 Raw map

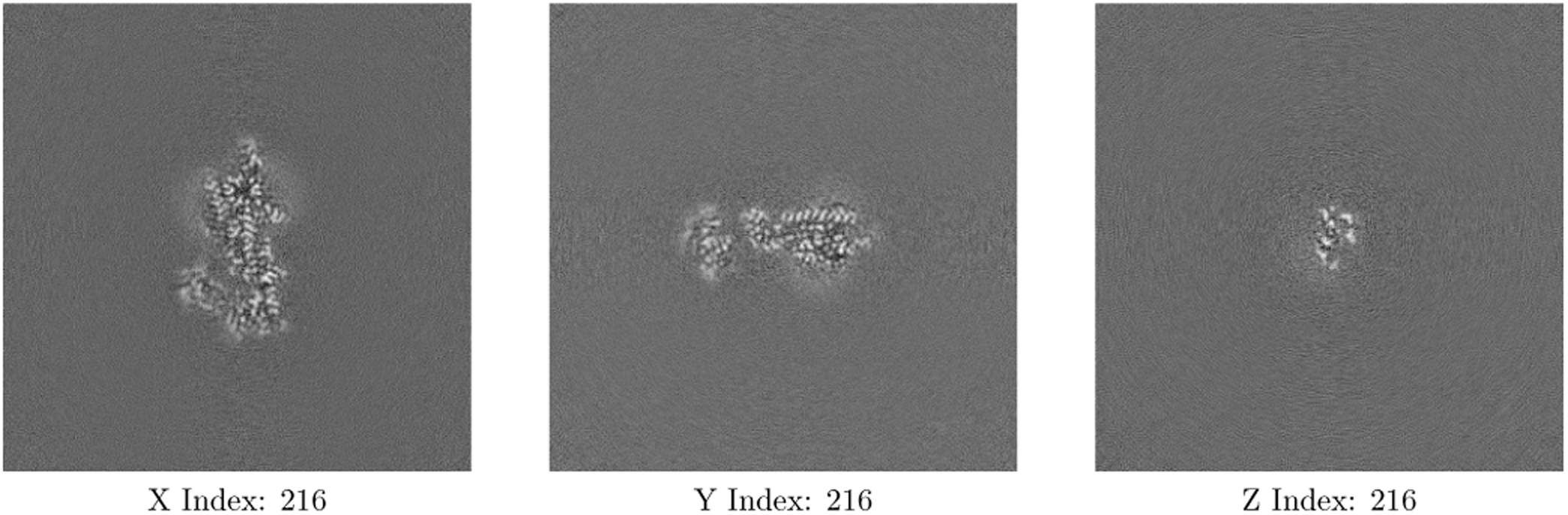

The images above show central slices of the map in three orthogonal directions.

### 6.3 Largest variance slices

#### 6.3.1 Primary map

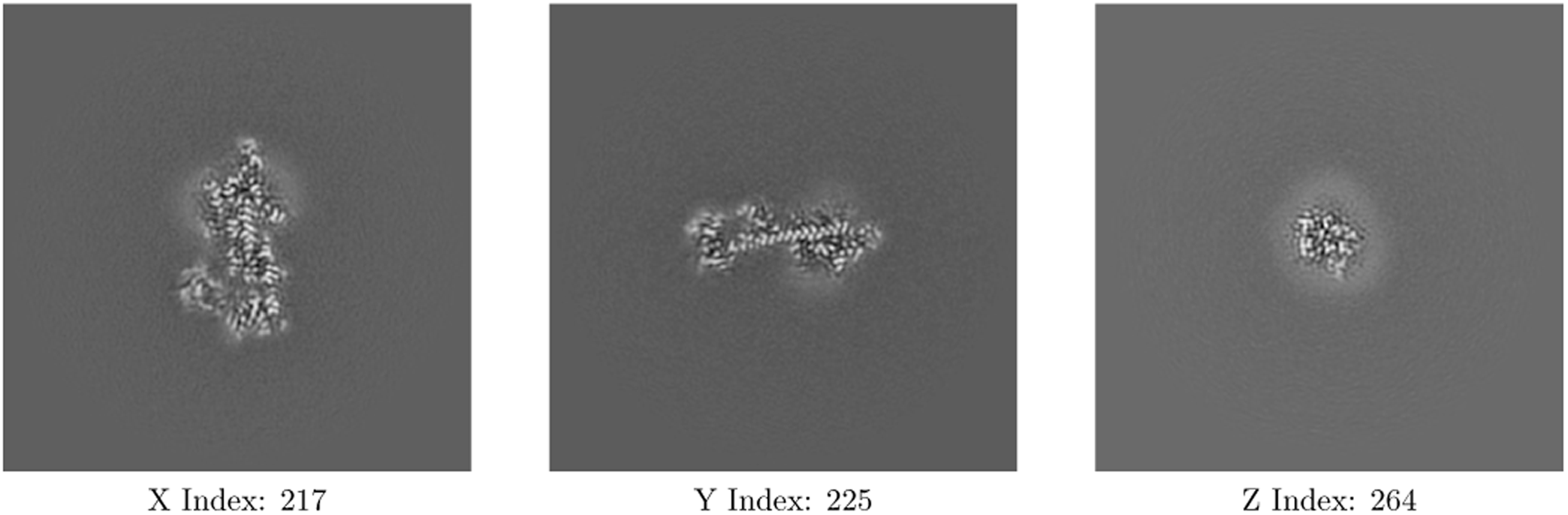

#### 6.3.2 Raw map

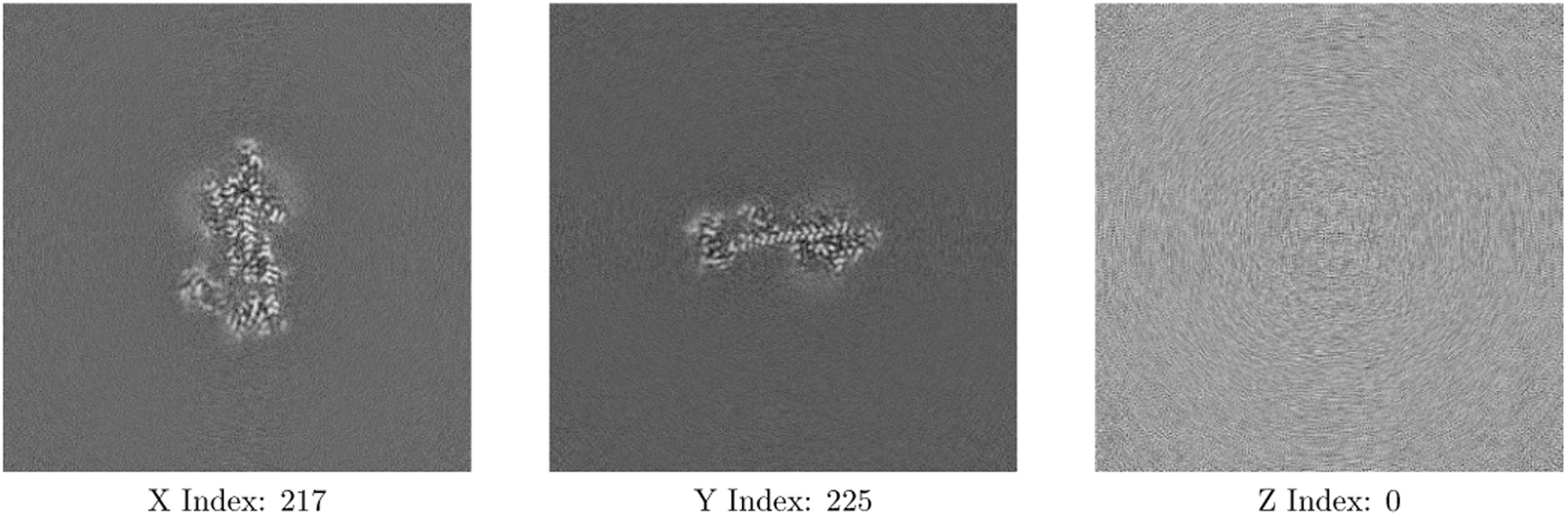

The images above show the largest variance slices of the map in three orthogonal directions.

### 6.4 Orthogonal standard-deviation projections (False-color)

#### 6.4.1 Primary map

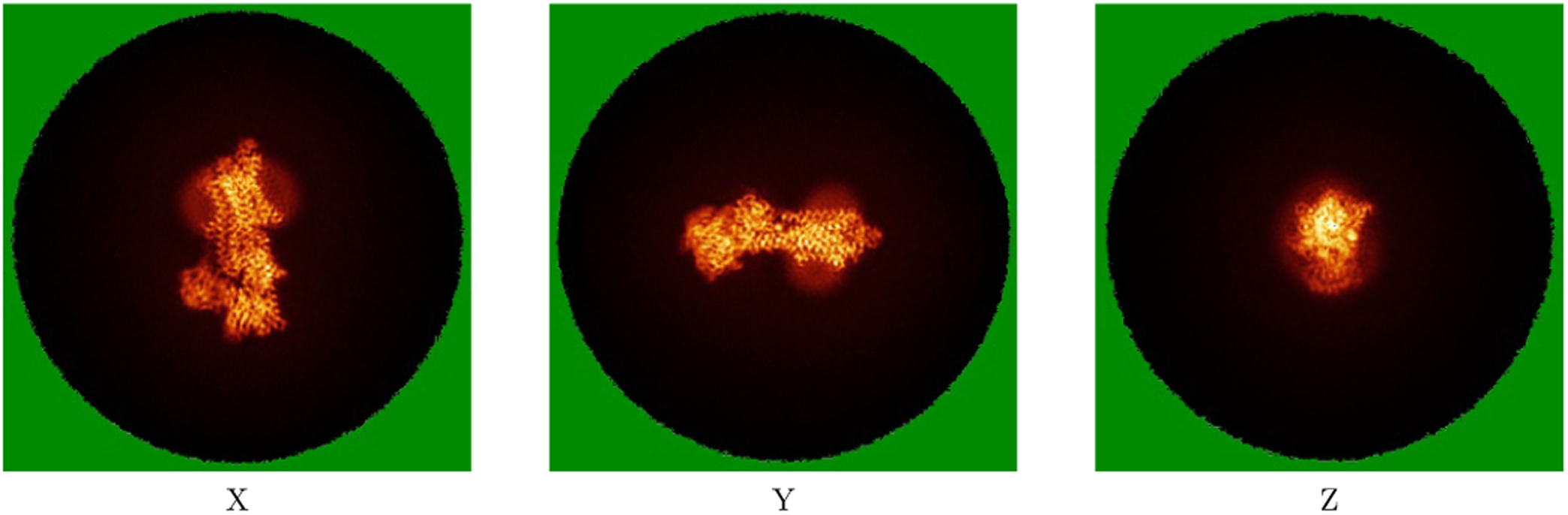

#### 6.4.2 Raw map

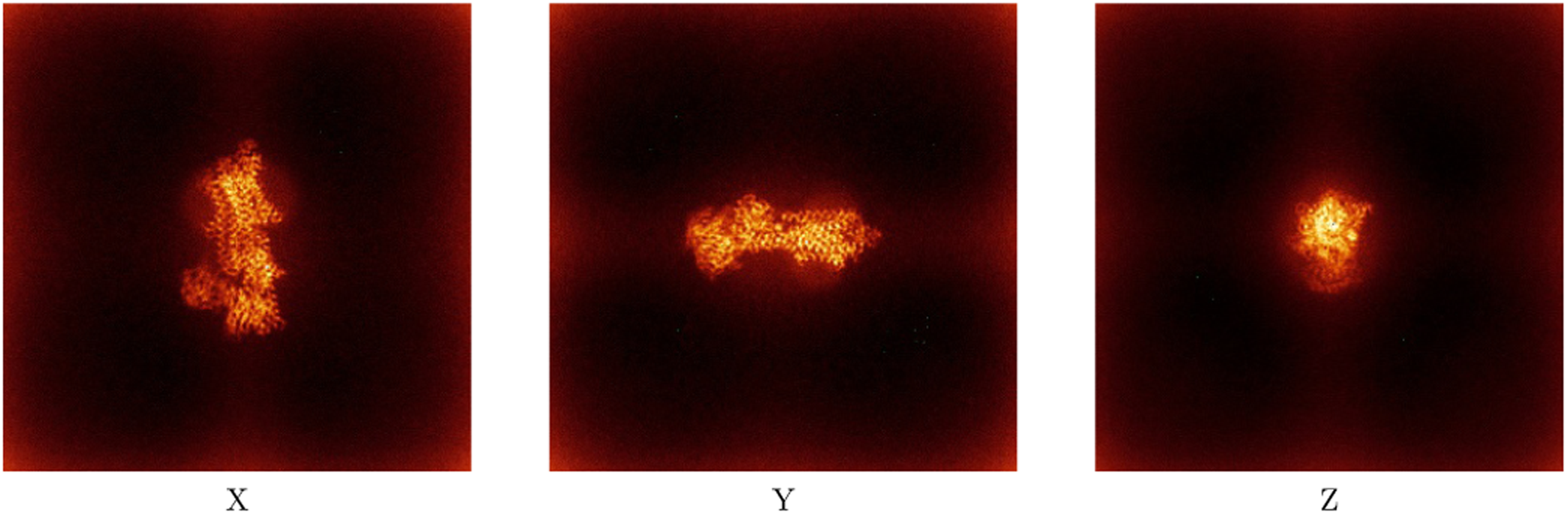

### 6.5 Orthogonal surface views

#### 6.5.1 Primary map

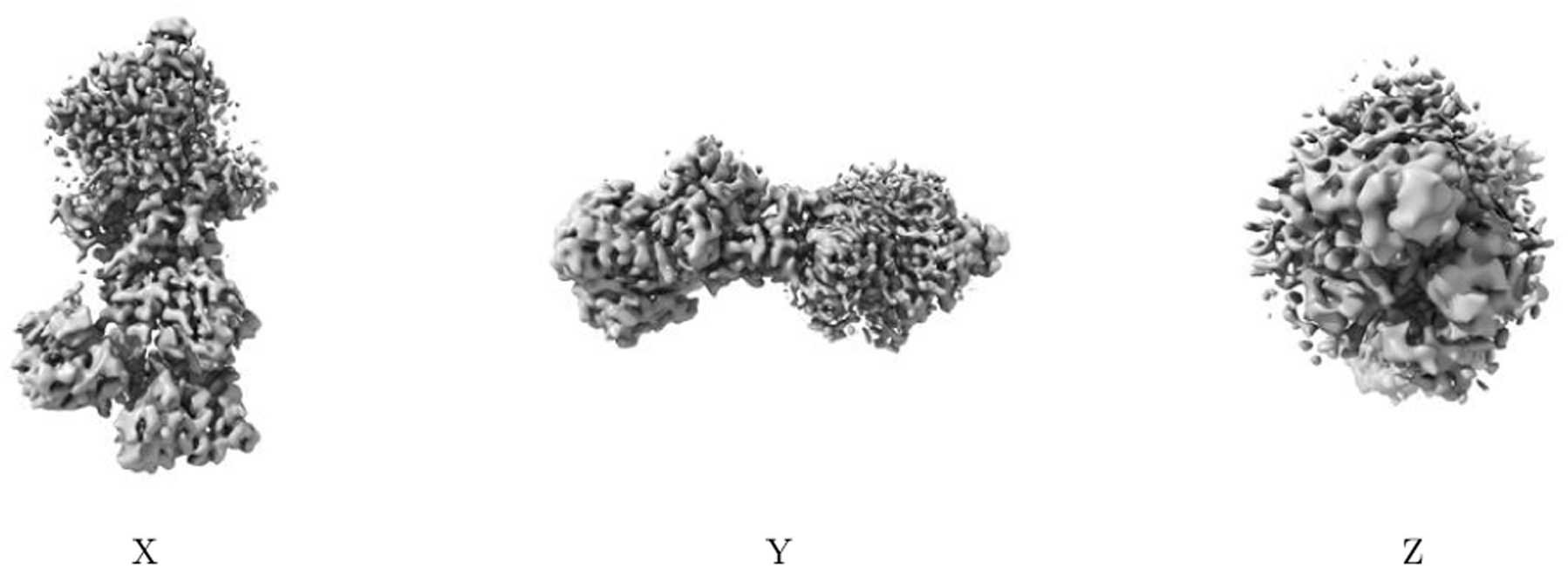

#### 6.5.2 Raw map

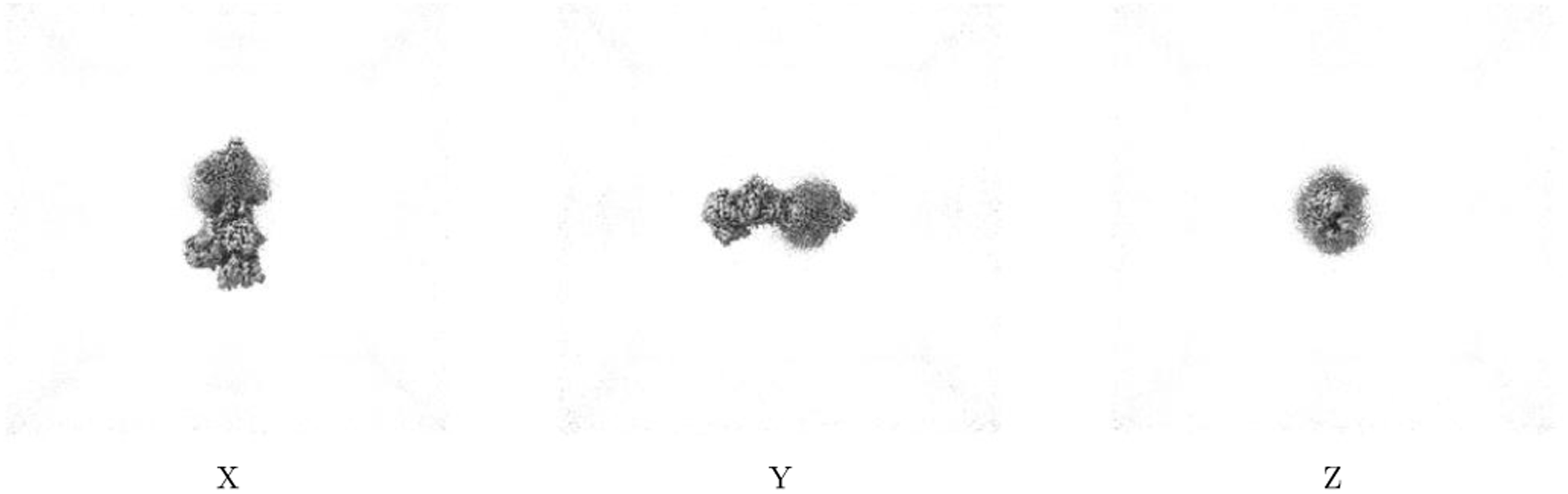

### 6.6 Mask visualisation

This section was not generated. No masks/segmentation were deposited.

## 7 Map analysis

This section contains the results of statistical analysis of the map.

### 7.1 Map-value distribution

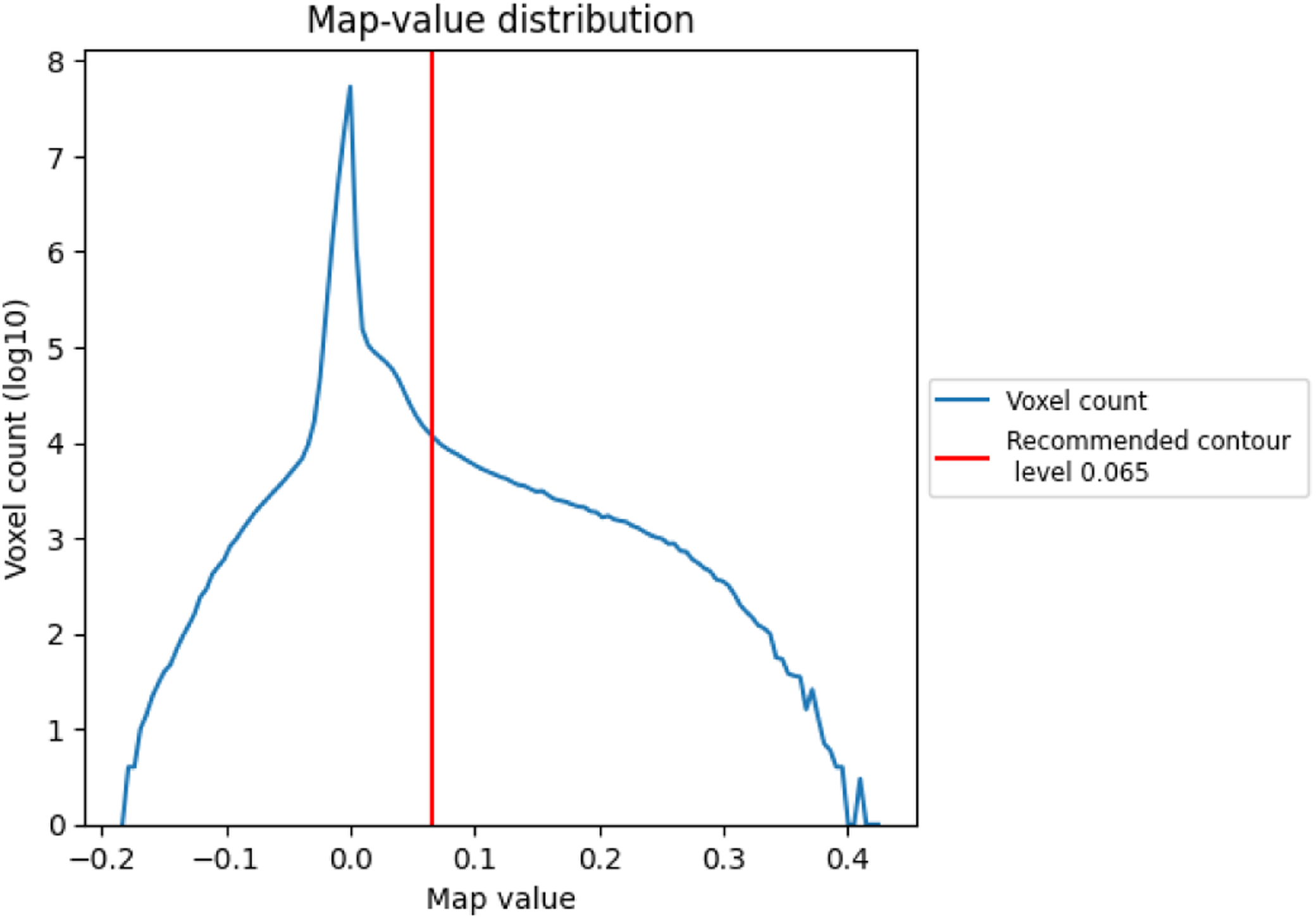

### 7.2 Volume estimate

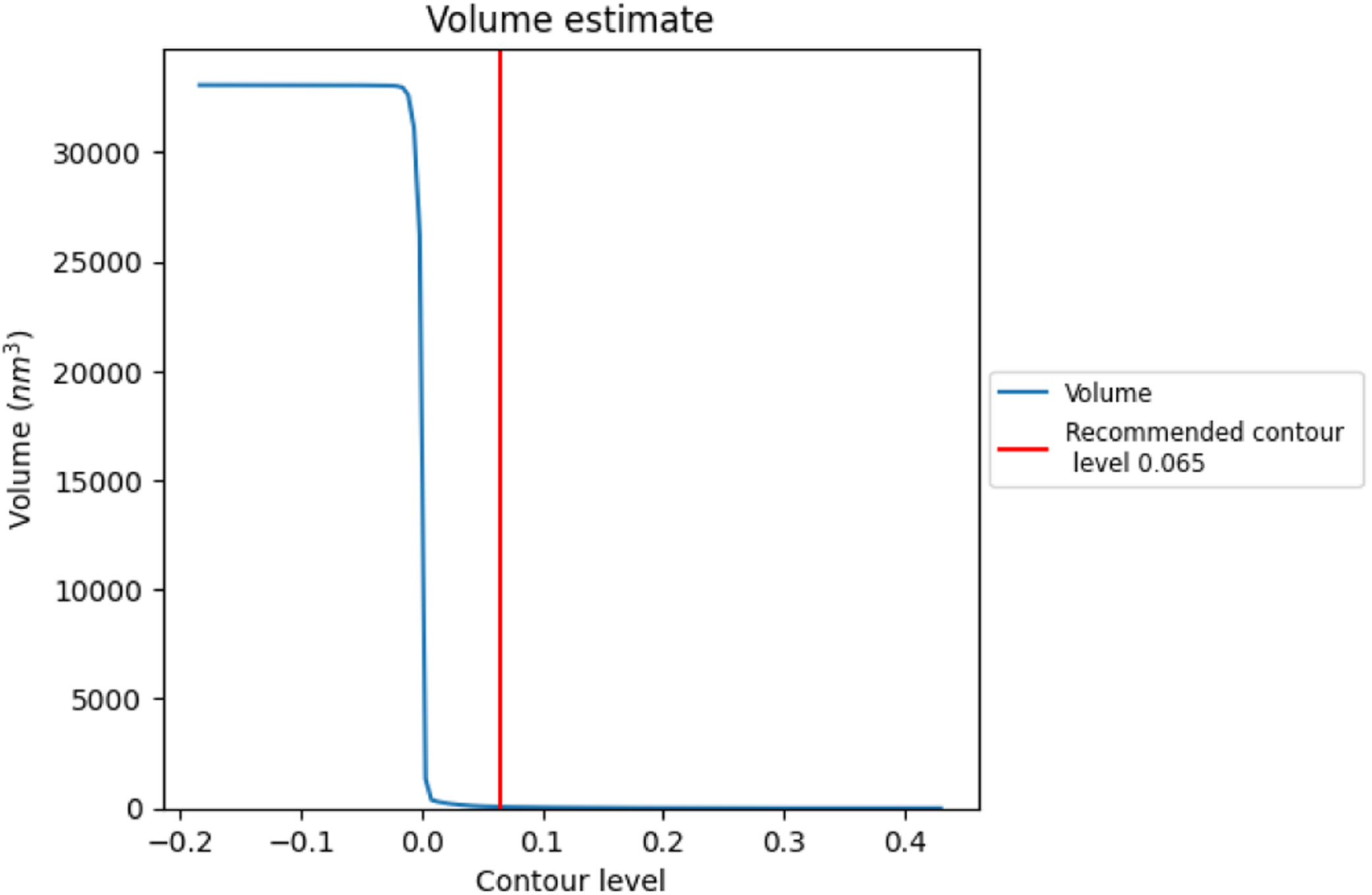

The volume at the recommended contour level is 64 nm^3^; this corresponds to an approximate mass of 58 kDa.

### 7.3 Rotationally averaged power spectrum

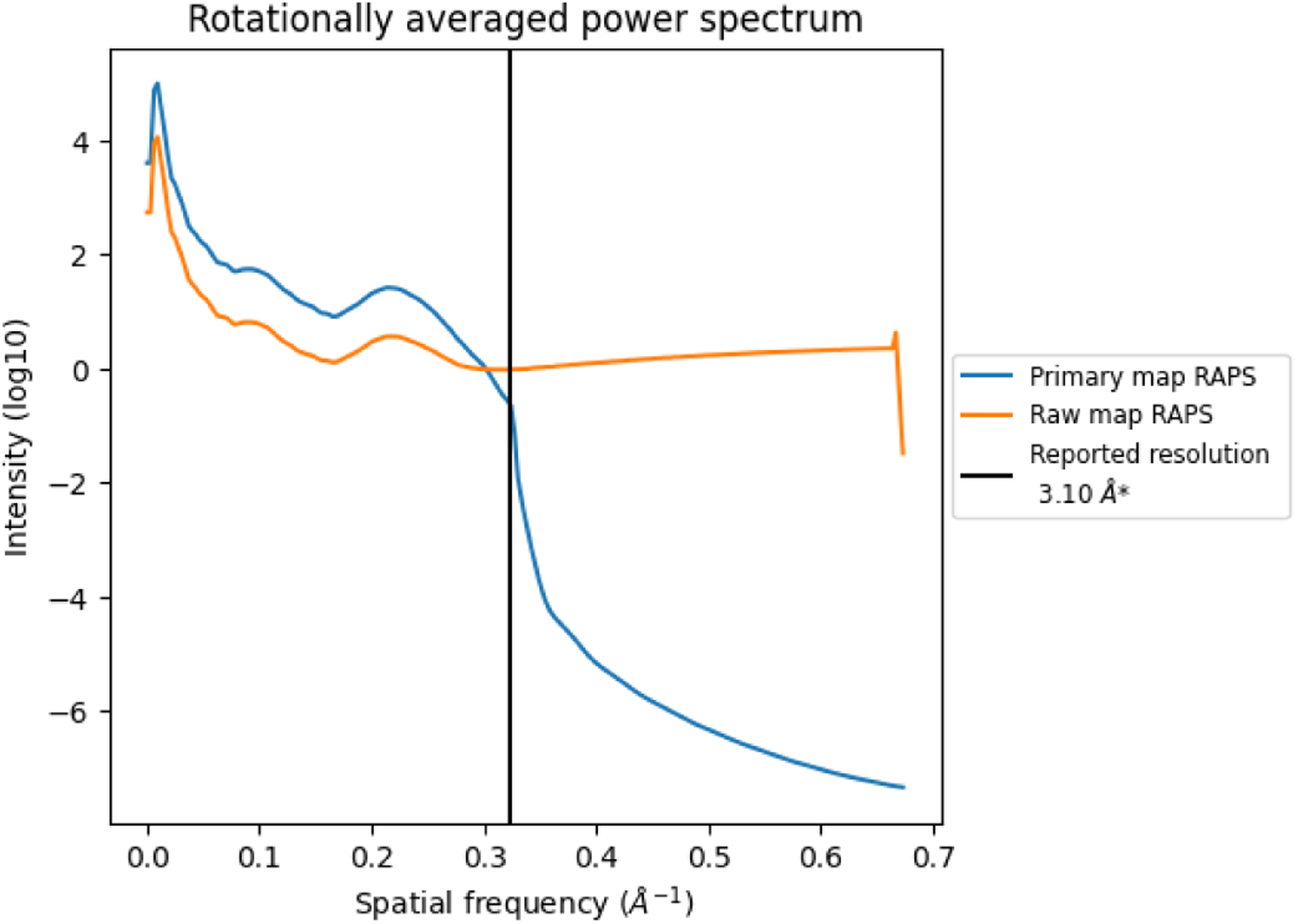

*Reported resolution corresponds to spatial frequency of 0.323 Å*^−^*^1^

## 8 Fourier-Shell correlation

### 8.1 FSC

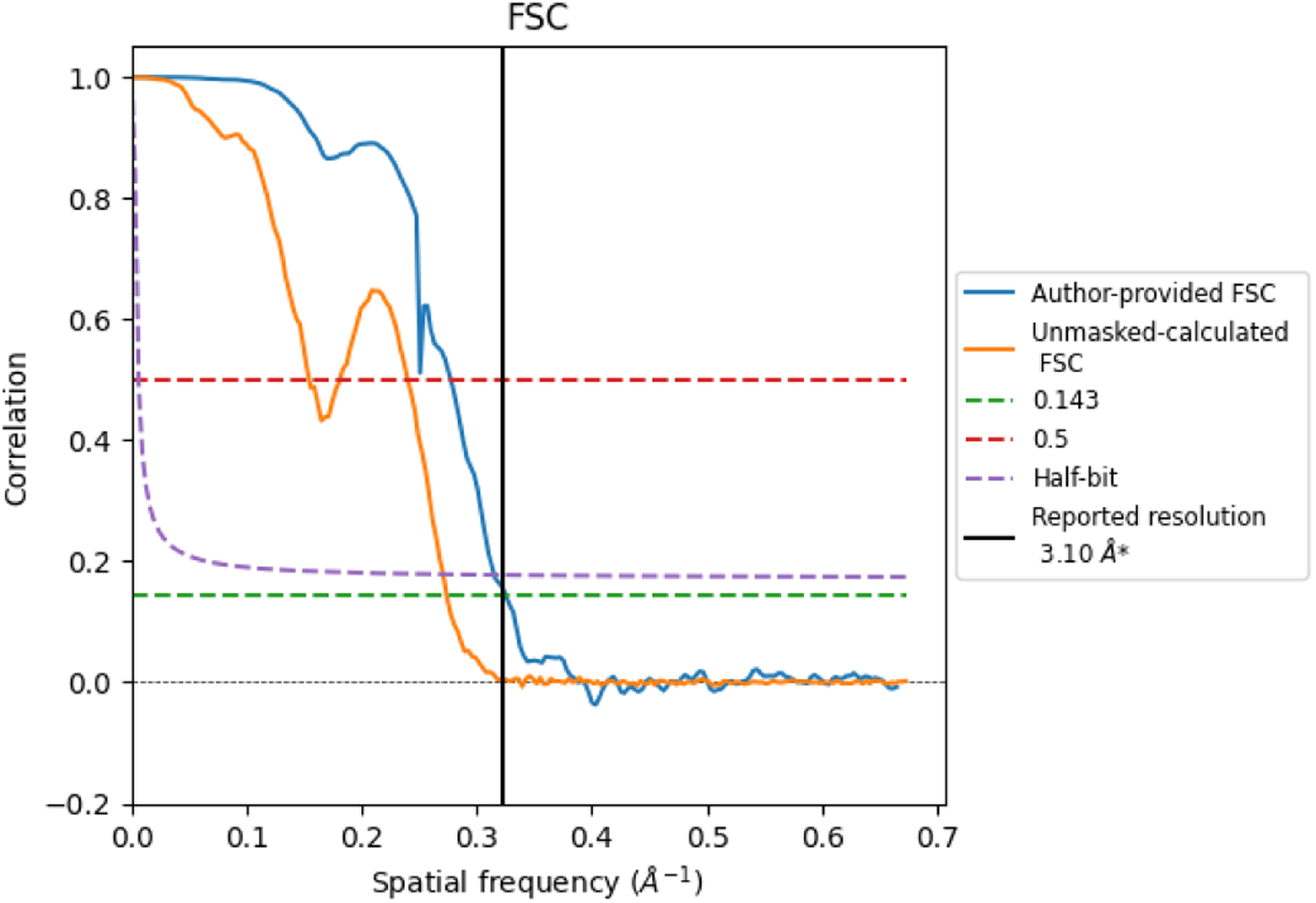

*Reported resolution corresponds to spatial frequency of 0.323 Å^-1^

### 8.2 Resolution estimates

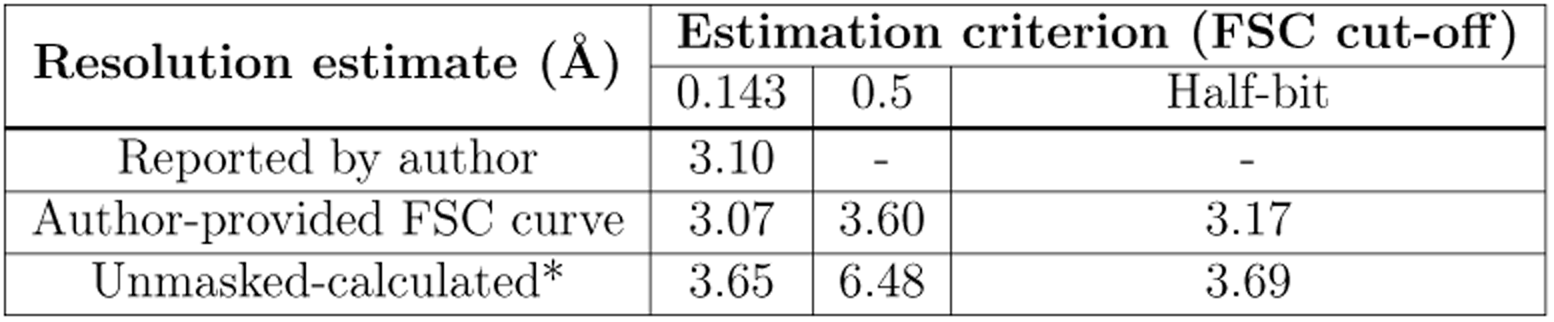

*Resolution estimate based on FSC curve calculated by comparison of deposited half-maps. The value from deposited half-maps intersecting FSC 0.143 CUT-OFF 3.65 differs from the reported value 3.1 by more than 10 %

## 9 Map-model fit

This section contains information regarding the fit between EMDB map EMD-73820 and PDB model 9Z5O. Per-residue inclusion information can be found in section 3 on page 4.

### 9.1 Map-model overlay

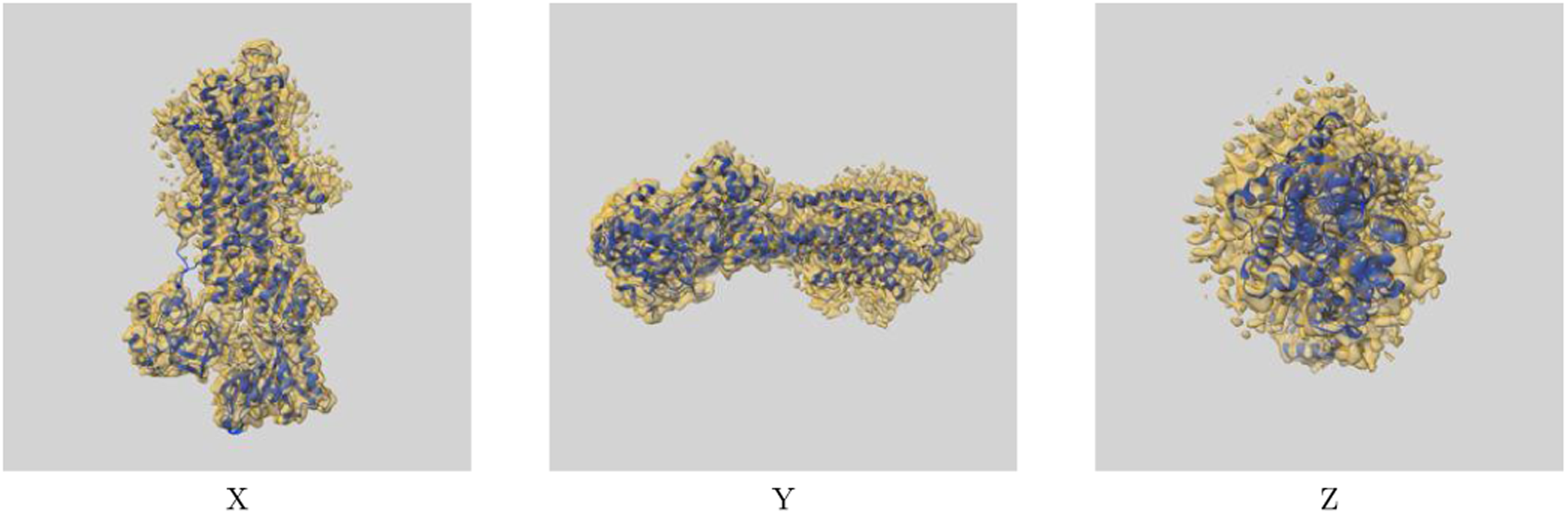

### 9.2 Q-score mapped to coordinate model

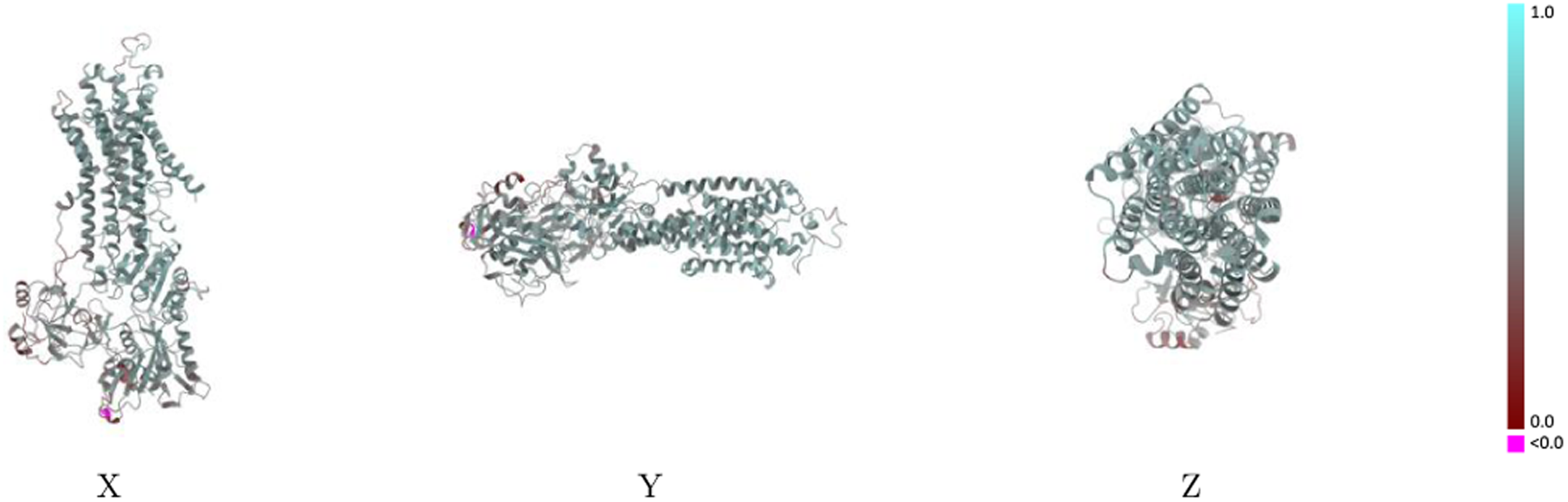

### 9.3 Atom inclusion mapped to coordinate model

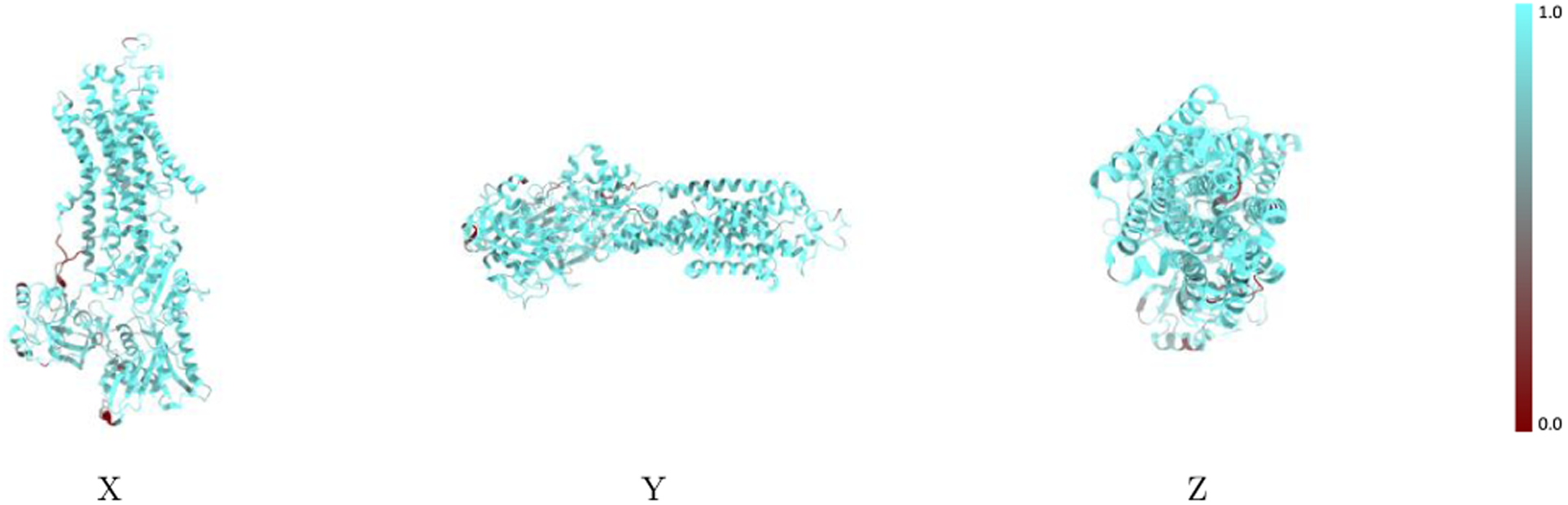

### 9.4 Atom inclusion

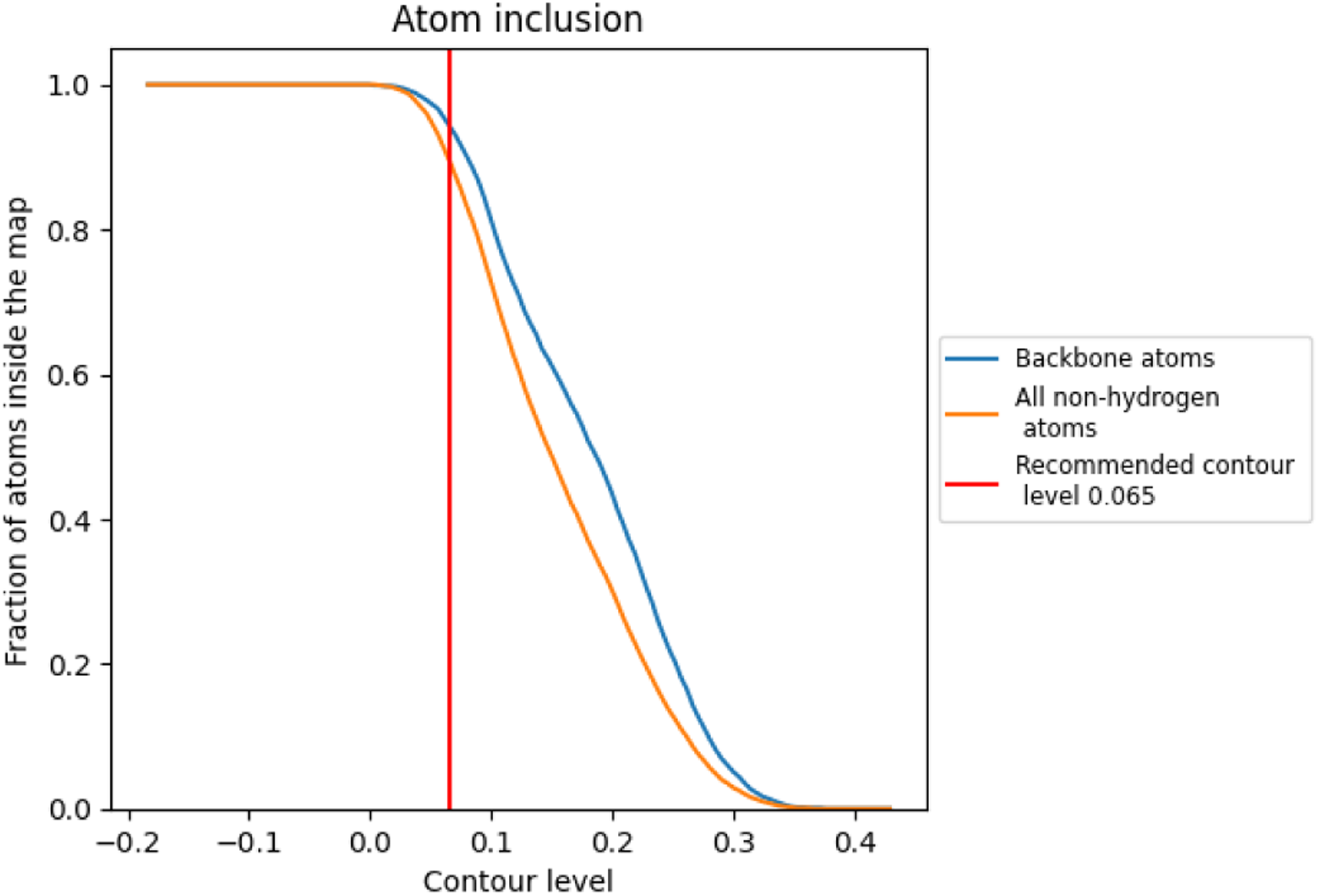

At the recommended contour level, 94% of all backbone atoms, 90% of all non-hydrogen atoms, are inside the map.

### 9.5 Map-model fit summary

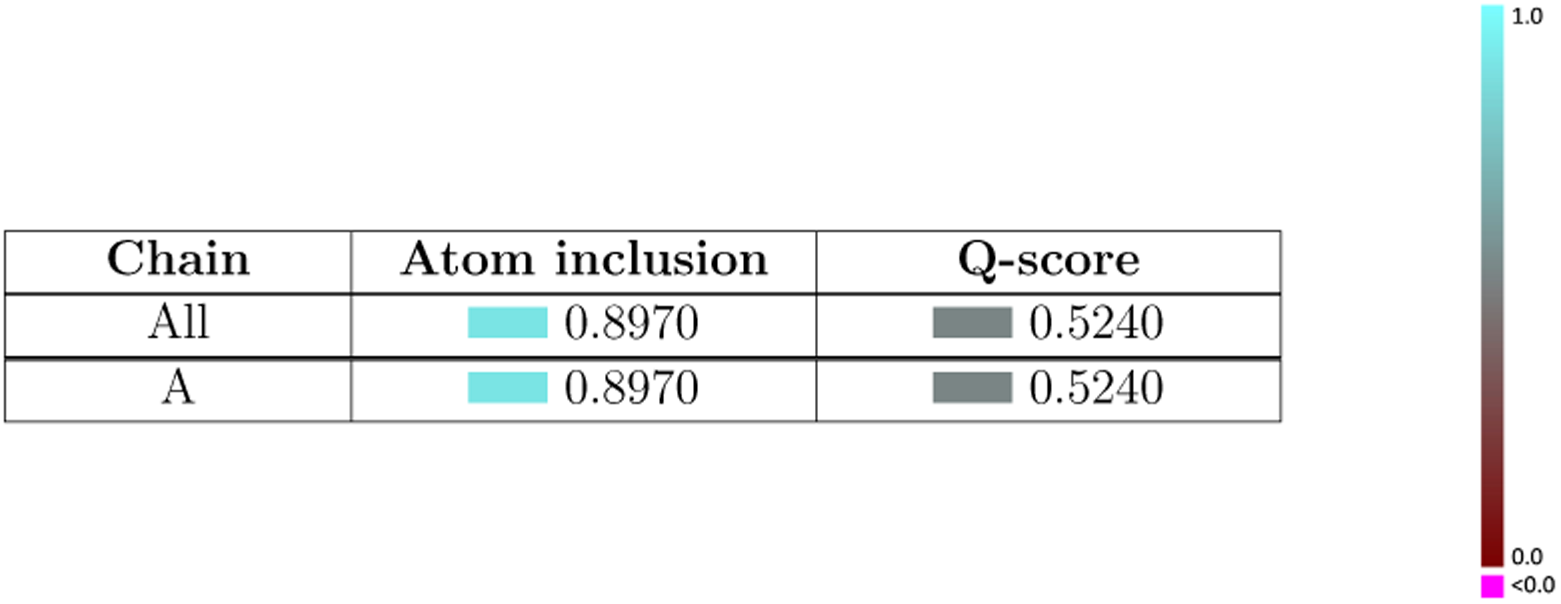

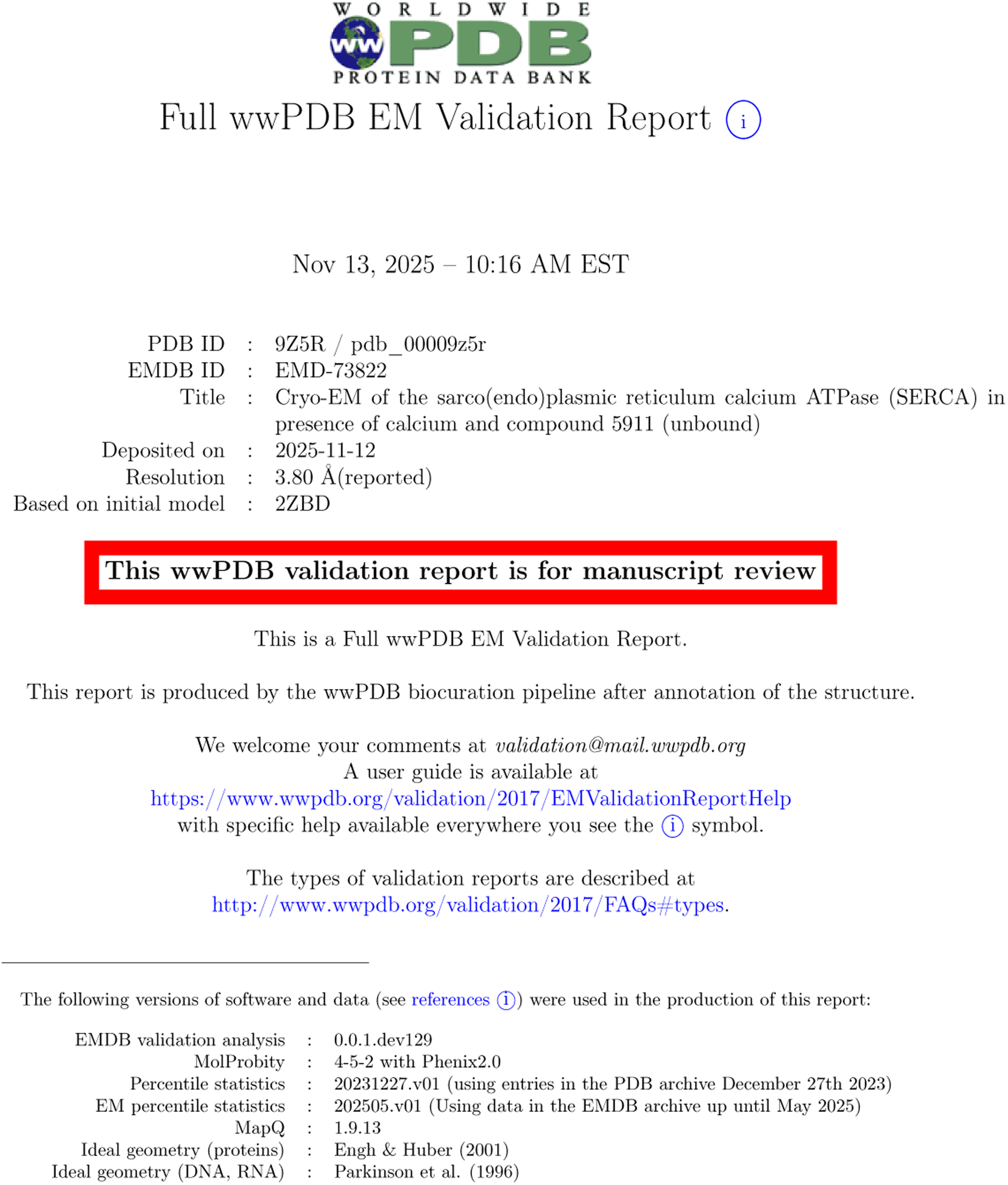

## 1 Overall quality at a glance

The following experimental techniques were used to determine the structure:*ELECTRON MICROSCOPY*

The reported resolution of this entry is 3.80 Å.

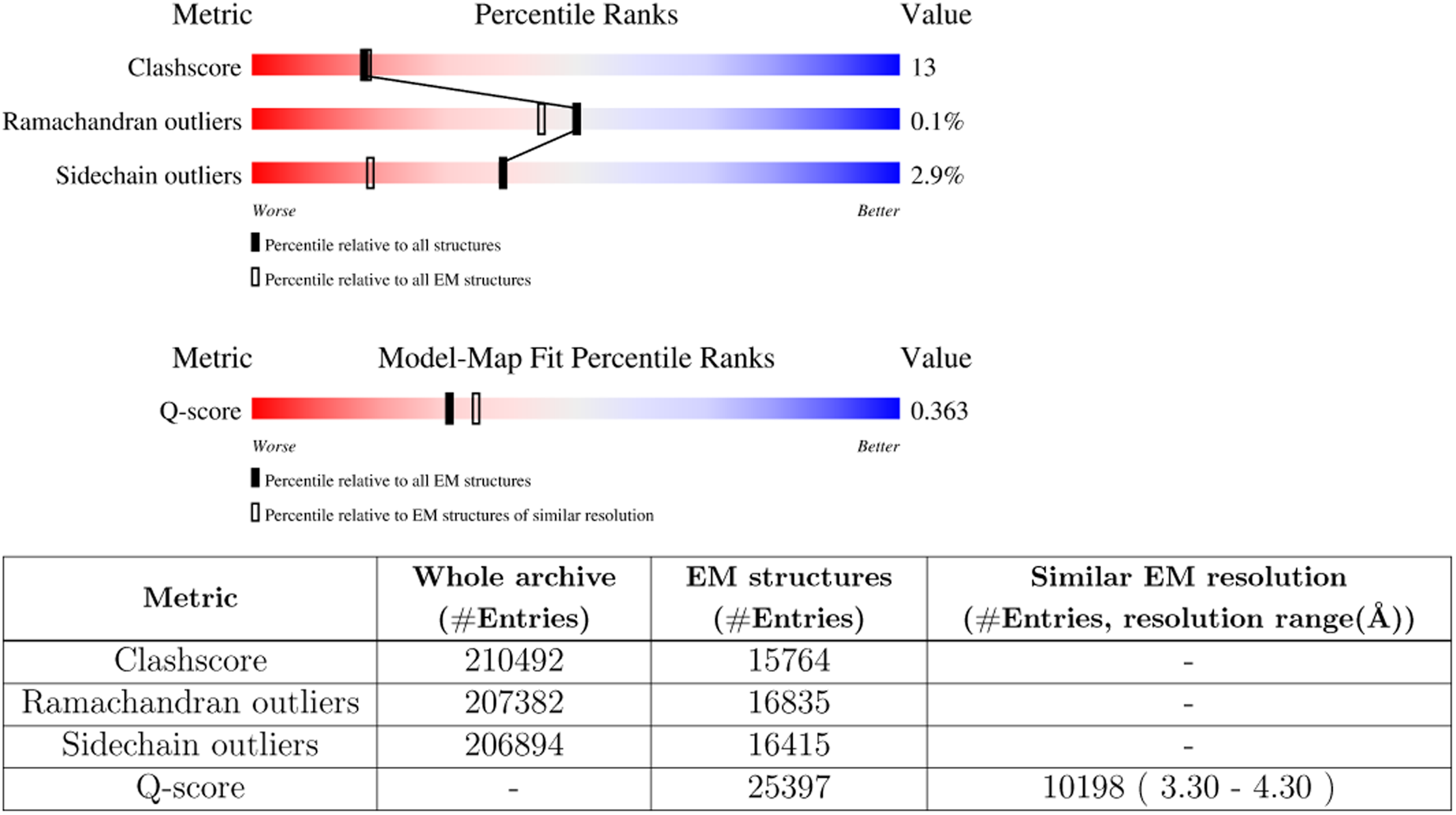

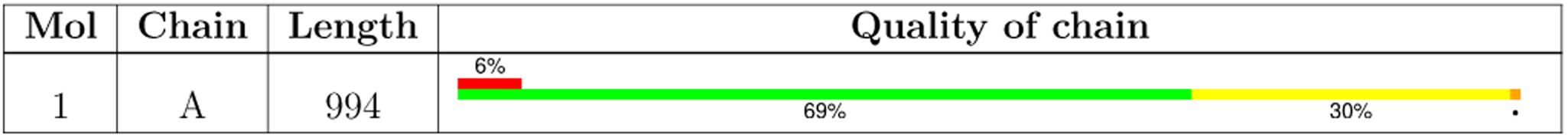

## 2 Entry composition

There are 2 unique types of molecules in this entry. The entry contains 7668 atoms, of which 0 are hydrogens and 0 are deuteriums.

- Molecule 1 is a protein called Sarcoplasmic/endoplasmic reticulum calcium ATPase 1.

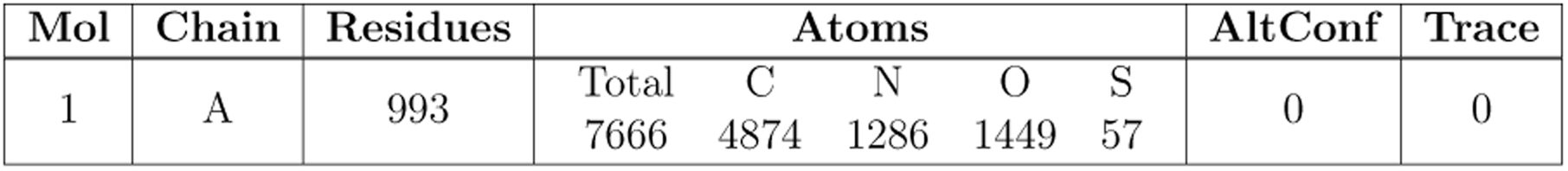

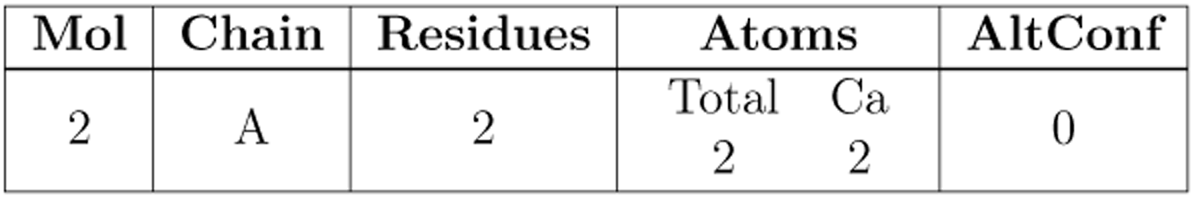

## 3 Residue-property plots

- Molecule 1: Sarcoplasmic/endoplasmic reticulum calcium ATPase 1

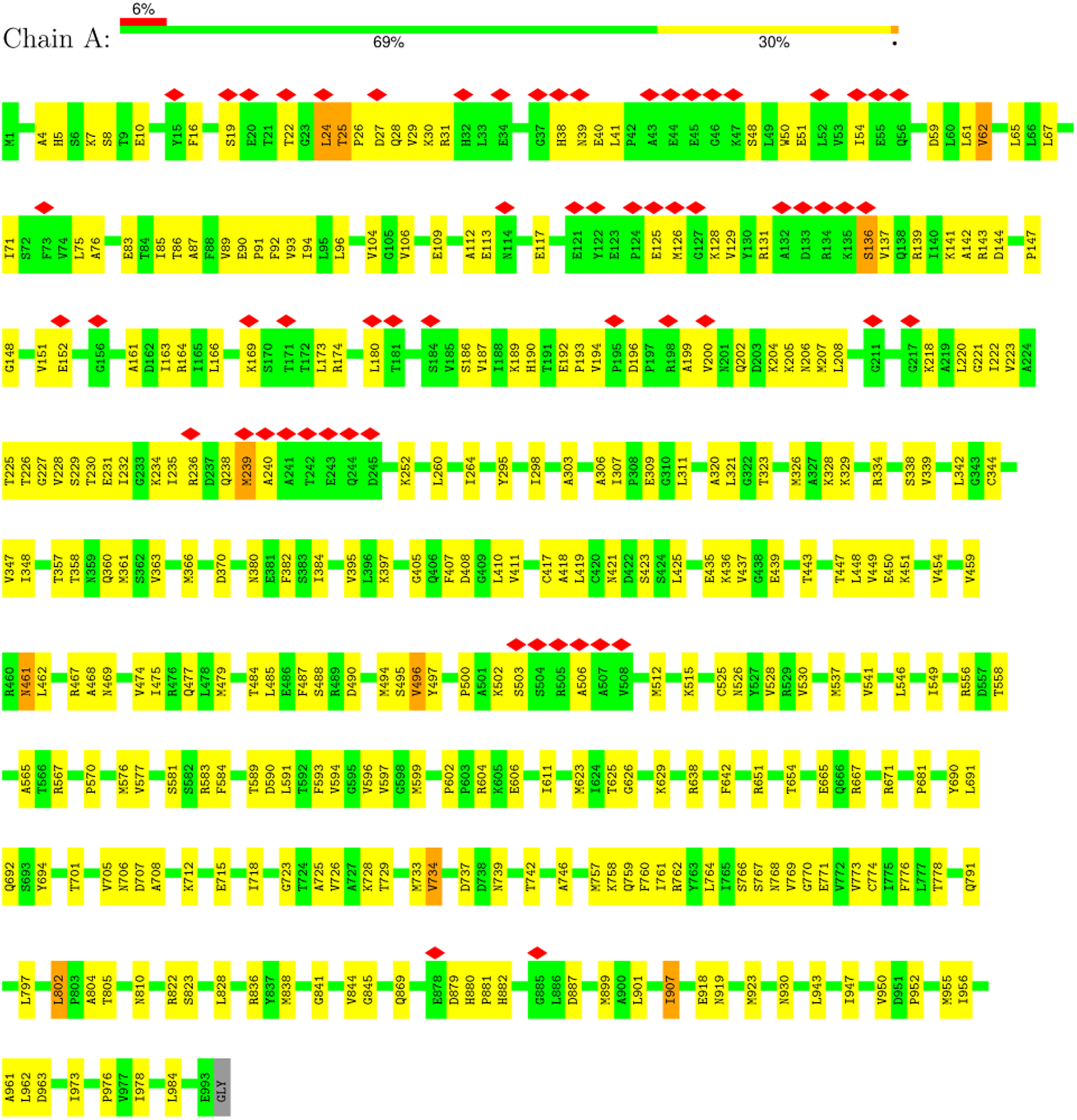

## 4 Experimental information

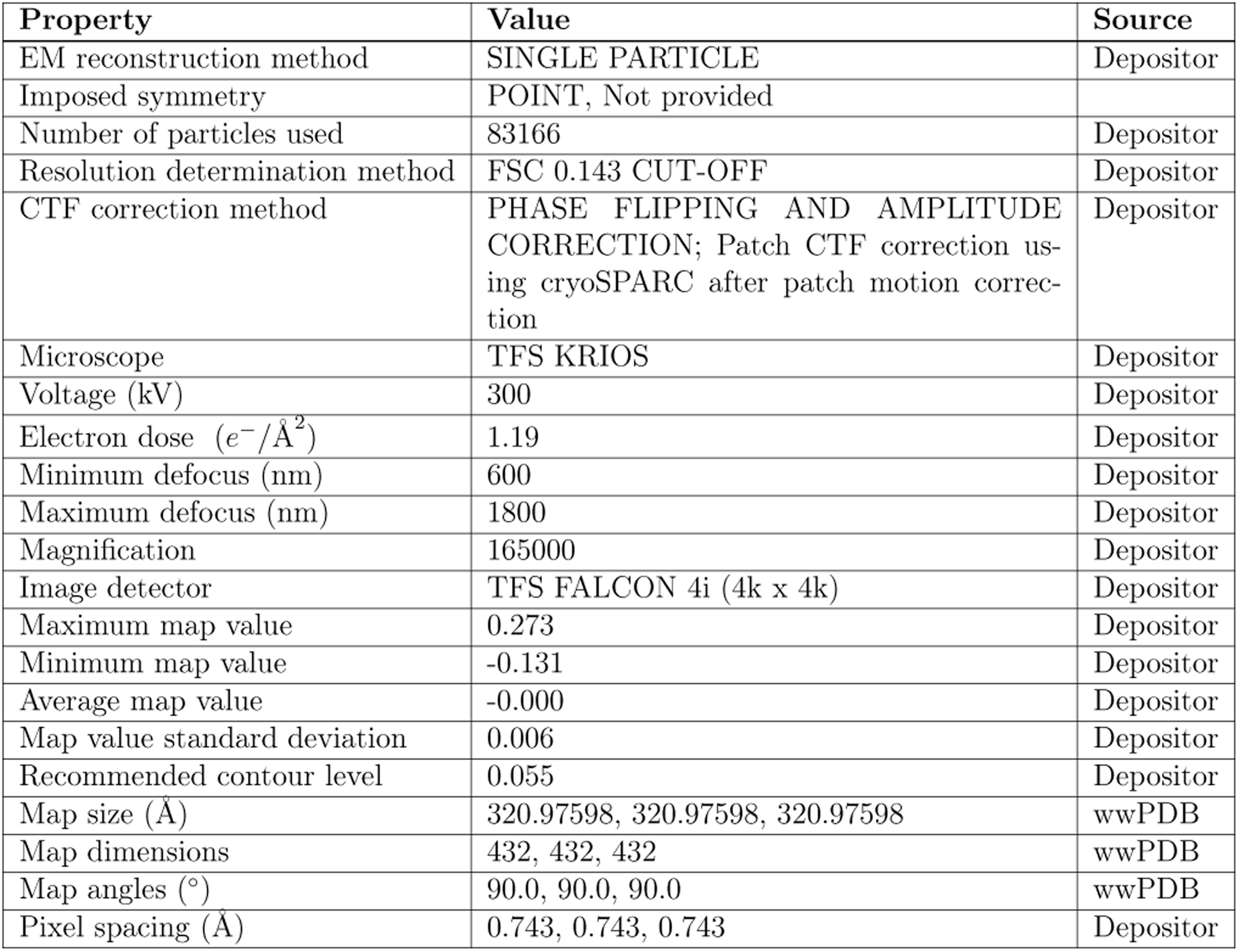

## 5 Model quality

### 5.1 Standard geometry

Bond lengths and bond angles in the following residue types are not validated in this section: CA

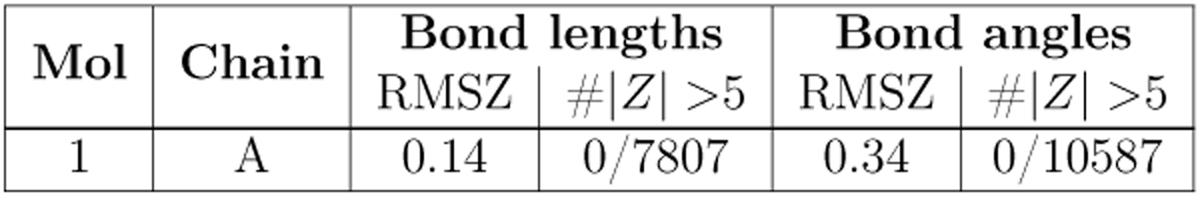

There are no bond length outliers.

There are no bond angle outliers.

There are no chirality outliers.

There are no planarity outliers.

### 5.2 Too-close contacts

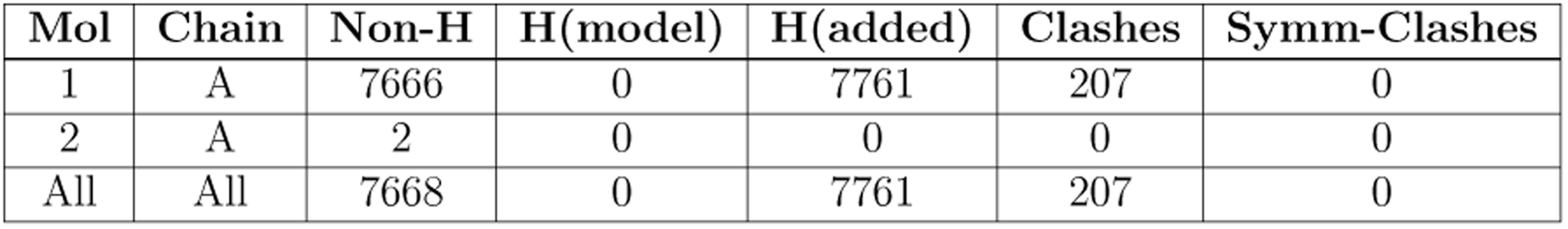

The all-atom clashscore is defined as the number of clashes found per 1000 atoms (including hydrogen atoms). The all-atom clashscore for this structure is 13.

All (207) close contacts within the same asymmetric unit are listed below, sorted by their clash magnitude.

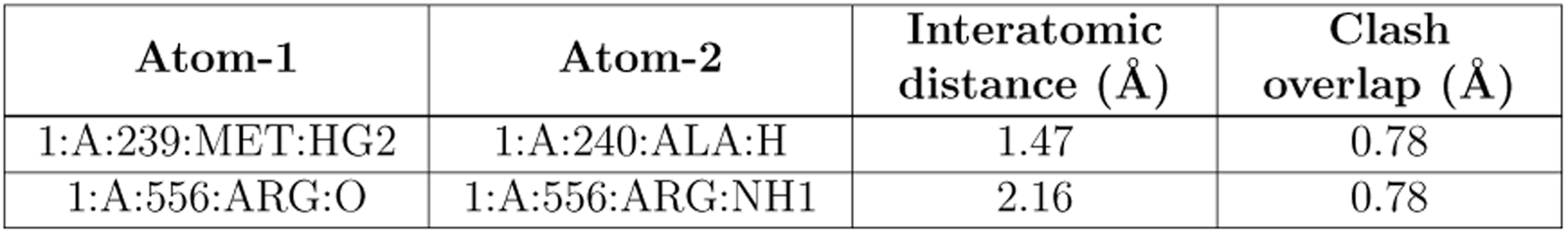
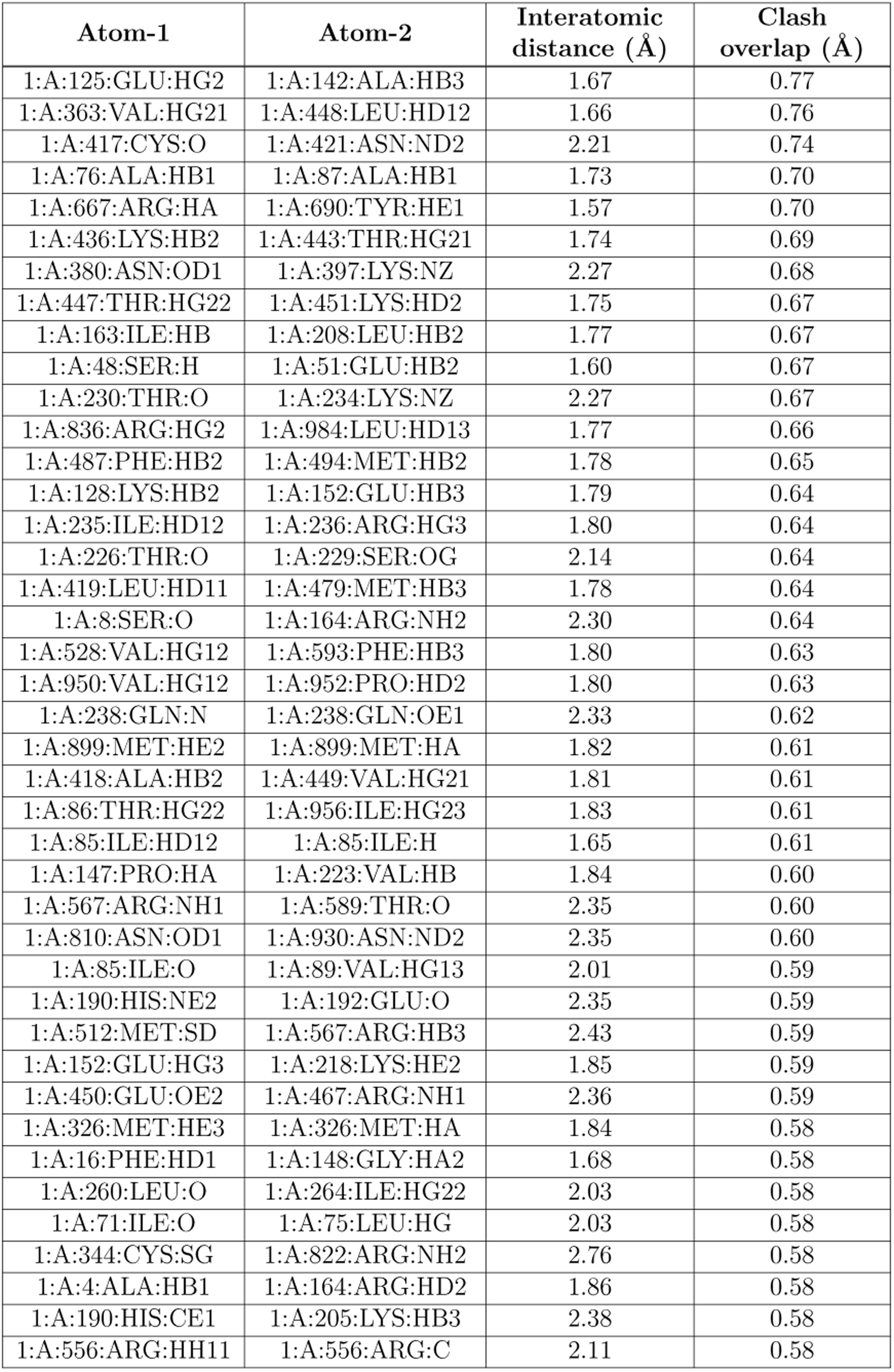
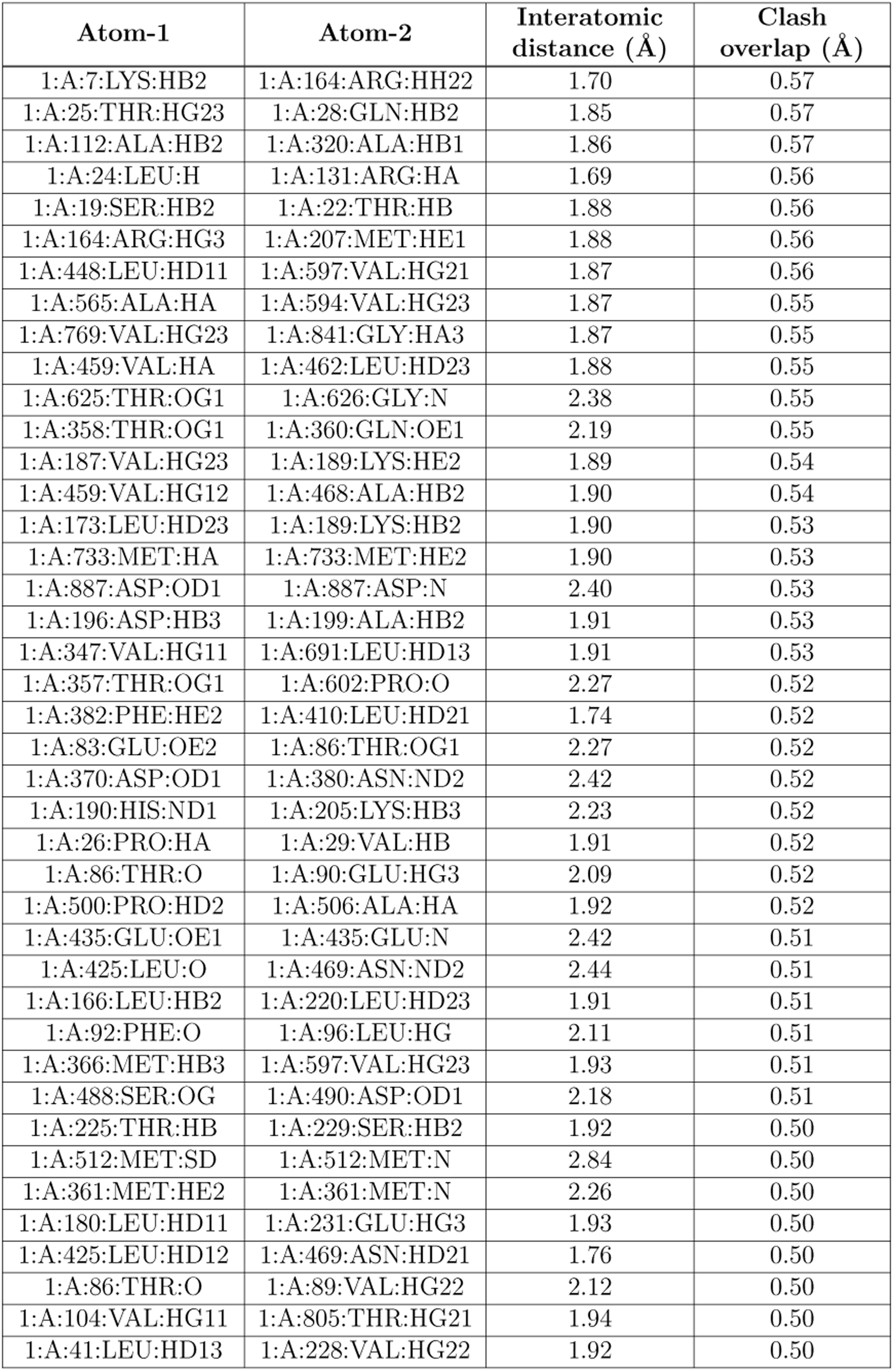
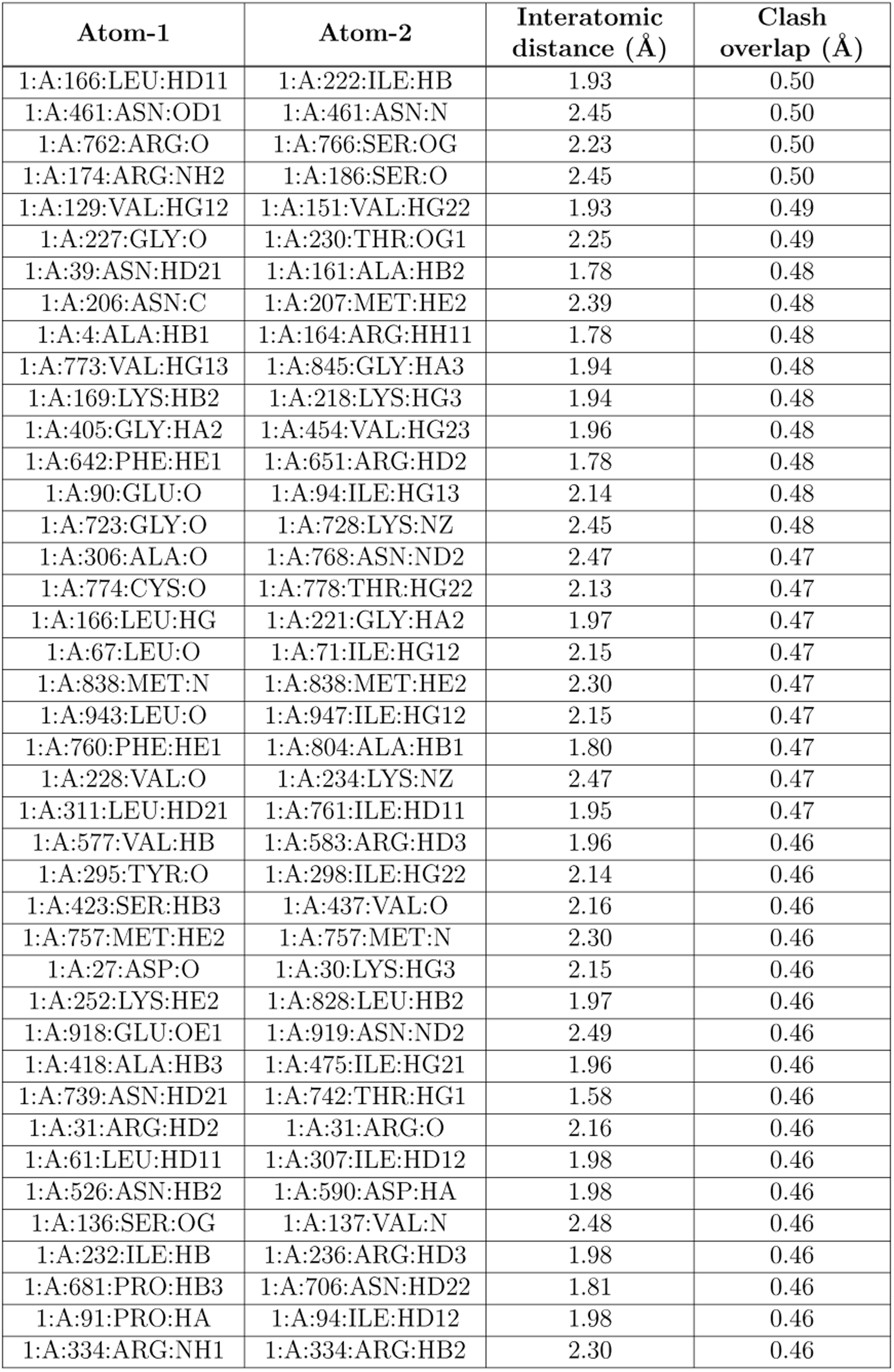
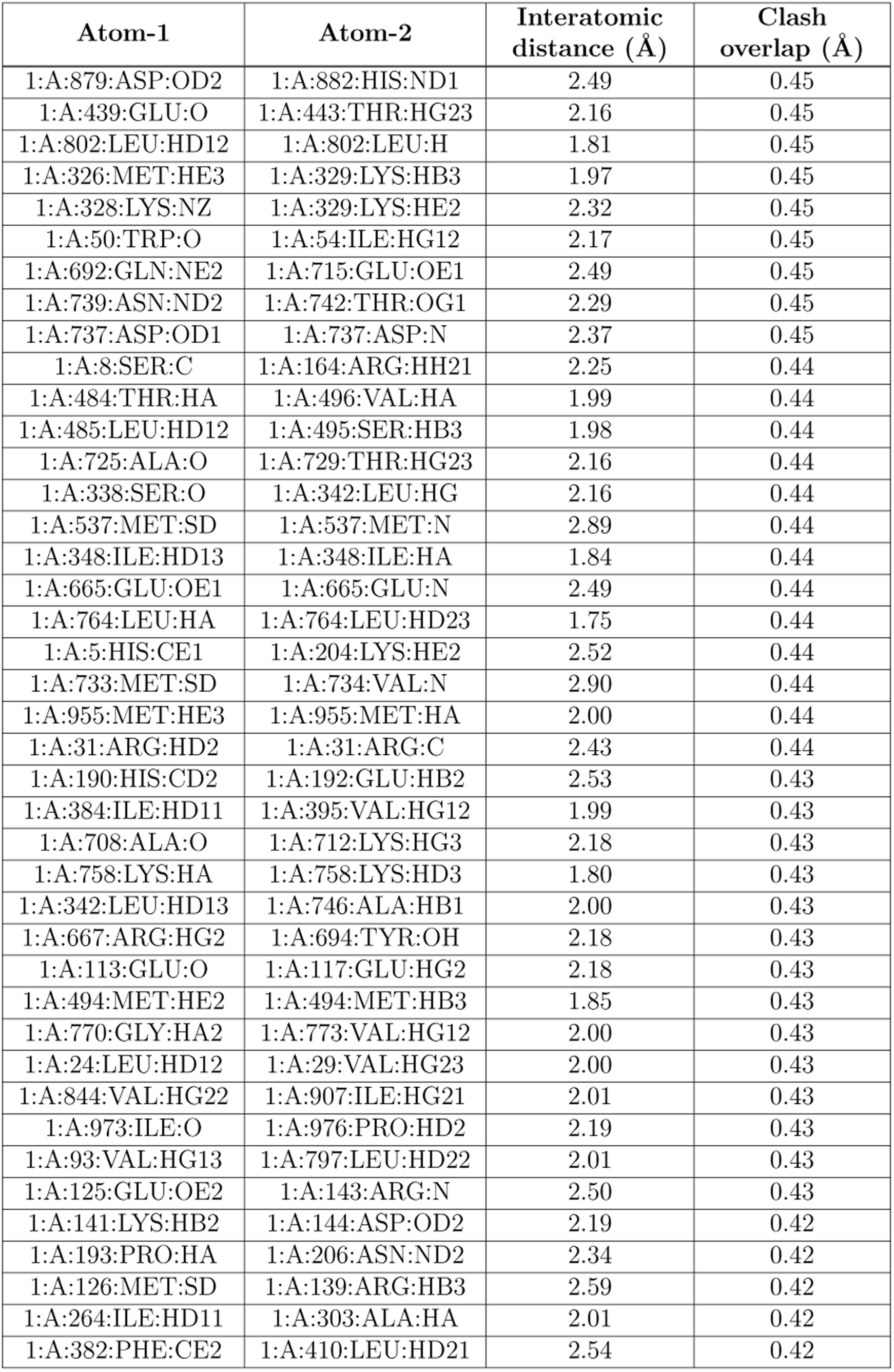
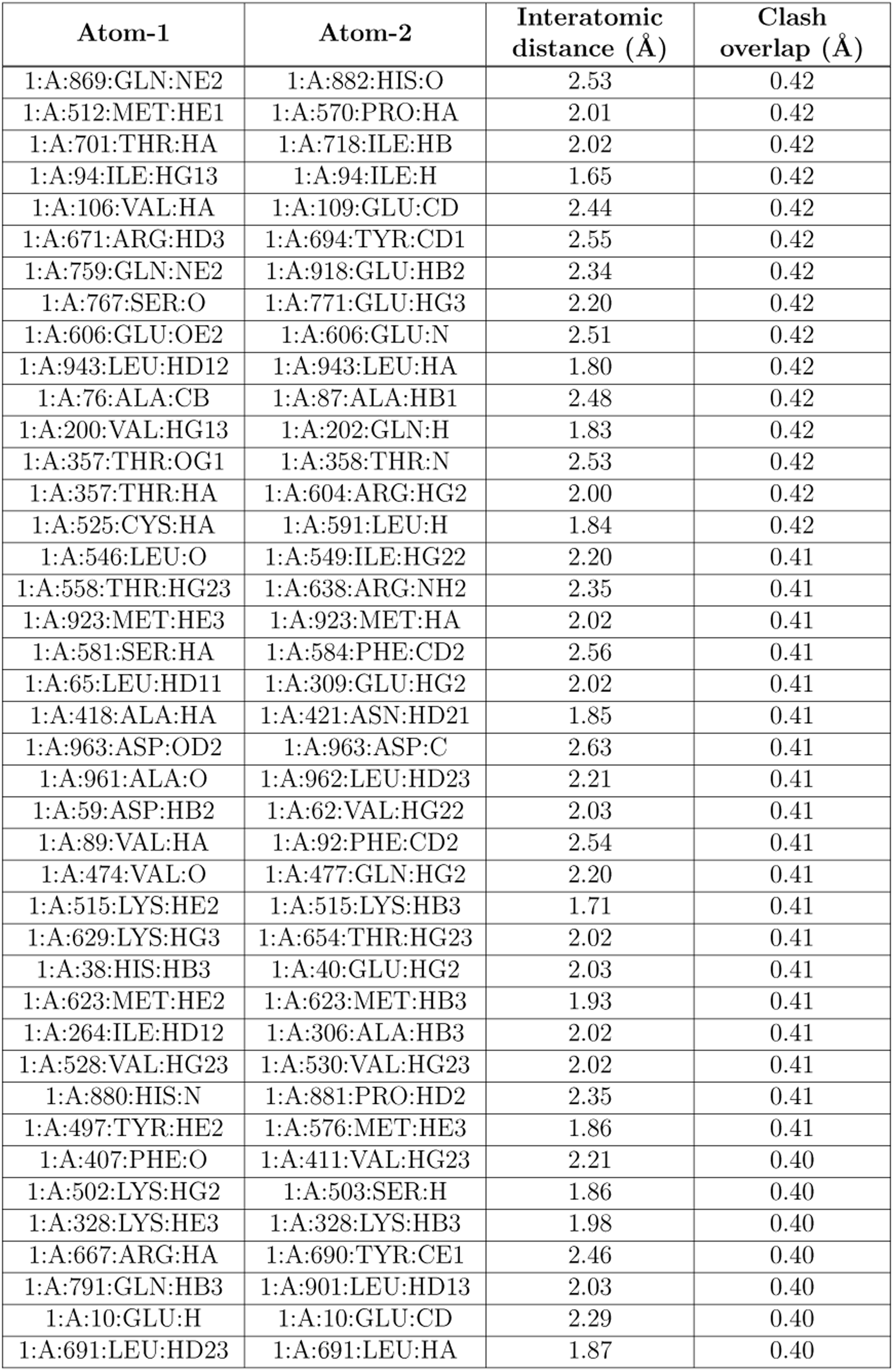

There are no symmetry-related clashes.

### 5.3 Torsion angles

#### 5.3.1 Protein backbone

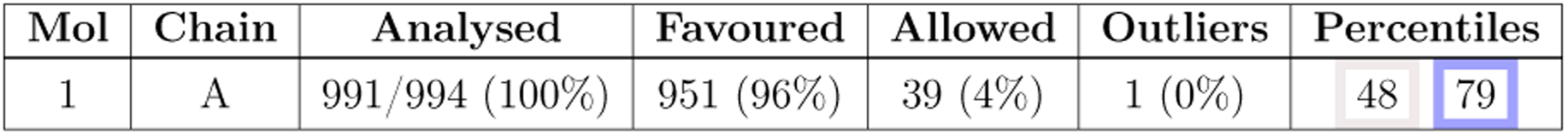

All (1) Ramachandran outliers are listed below:

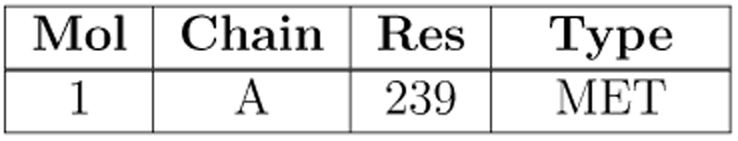

#### 5.3.2 Protein sidechains

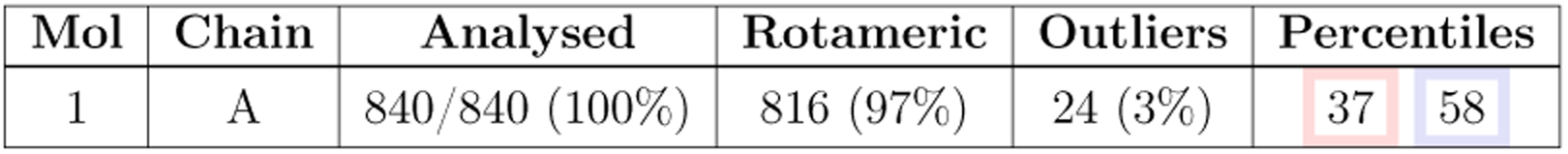

All (24) residues with a non-rotameric sidechain are listed below:

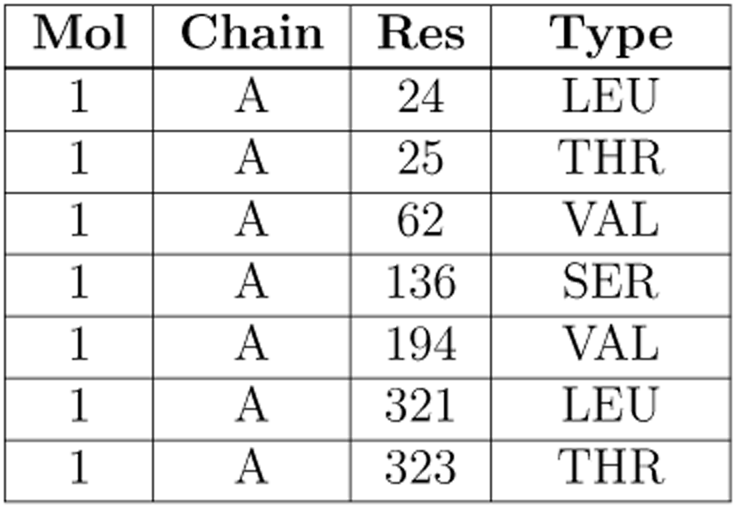
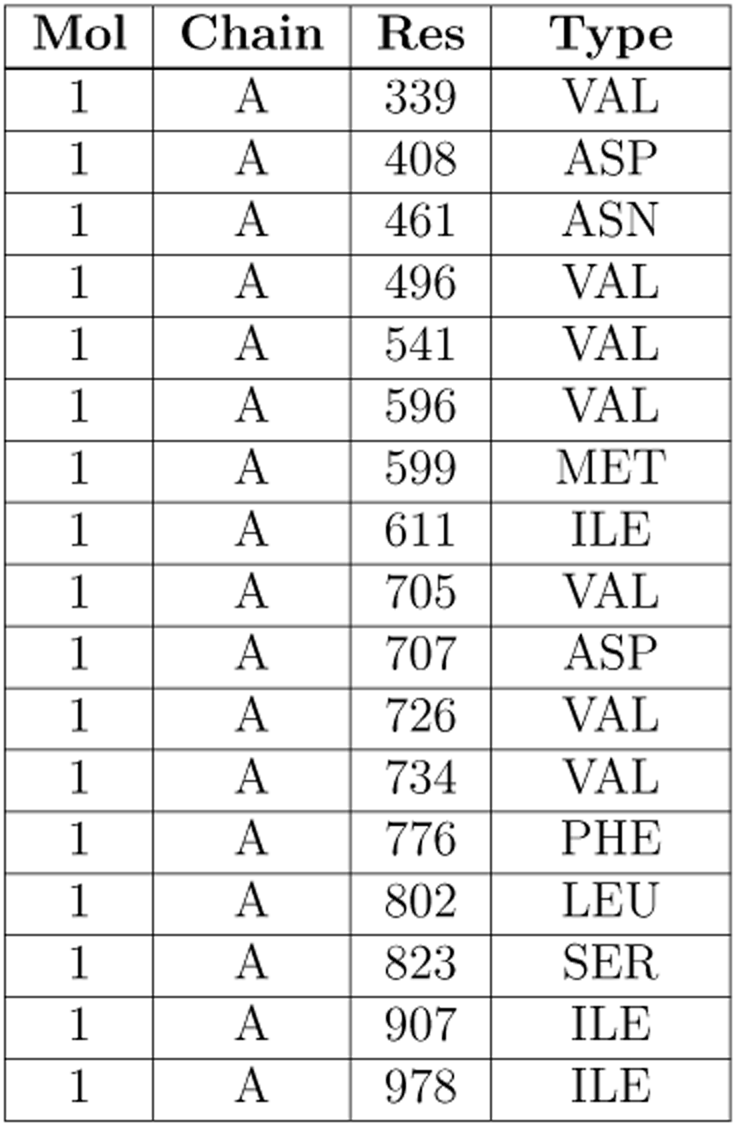

Sometimes sidechains can be flipped to improve hydrogen bonding and reduce clashes. All (15) such sidechains are listed below:

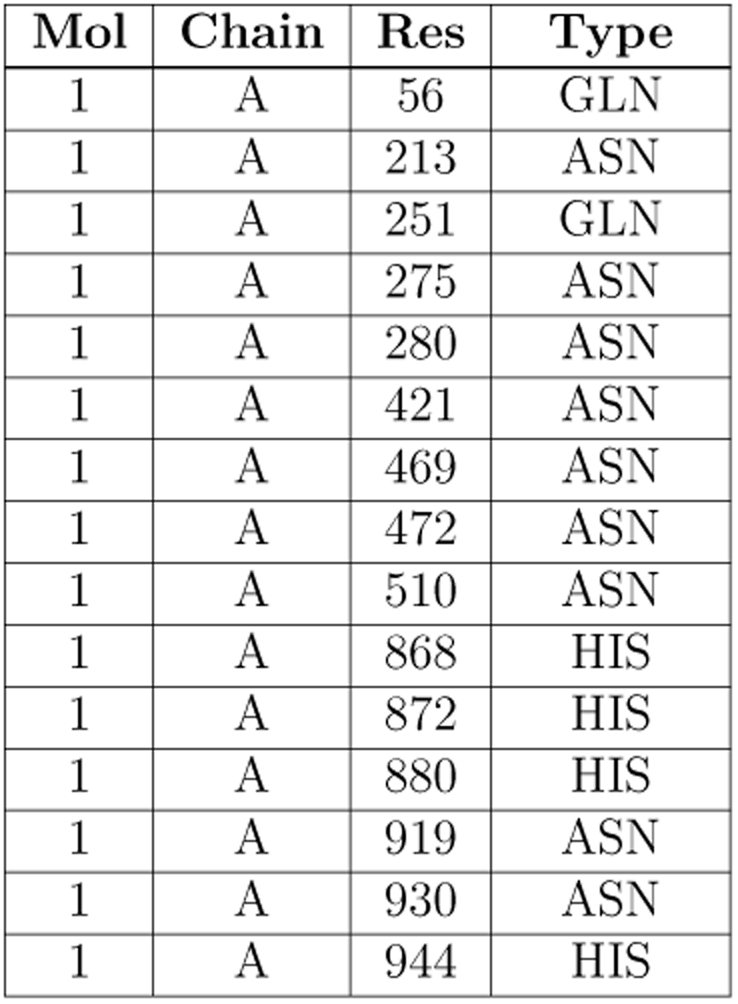

#### 5.3.3 RNA

There are no RNA molecules in this entry.

### 5.4 Non-standard residues in protein, DNA, RNA chains

There are no non-standard protein/DNA/RNA residues in this entry.

### 5.5 Carbohydrates

There are no oligosaccharides in this entry.

### 5.6 Ligand geometry

Of 2 ligands modelled in this entry, 2 are monoatomic - leaving 0 for Mogul analysis. There are no bond length outliers.

There are no bond angle outliers. There are no chirality outliers.

There are no torsion outliers. There are no ring outliers.

No monomer is involved in short contacts.

### 5.7 Other polymers

There are no such residues in this entry.

### 5.8 Polymer linkage issues

There are no chain breaks in this entry.

## 6 Map visualisation

This section contains visualisations of the EMDB entry EMD-73822. These allow visual inspection of the internal detail of the map and identification of artifacts.

### 6.1 Orthogonal projections

#### 6.1.1 Primary map

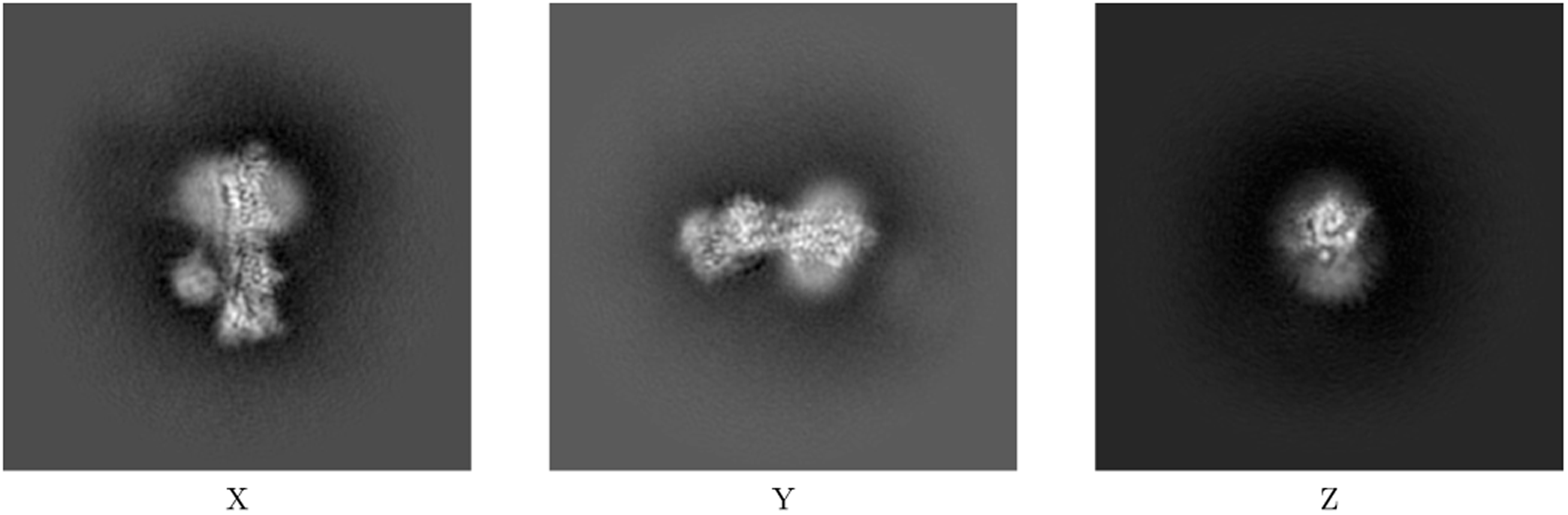

#### 6.1.2 Raw map

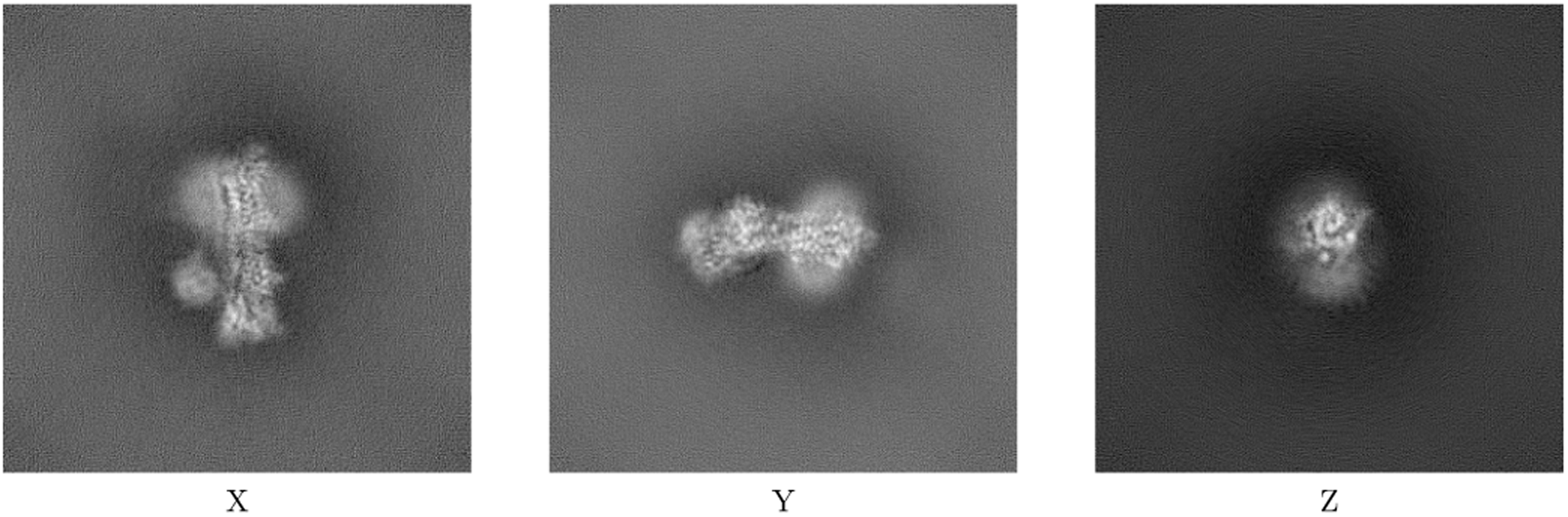

The images above show the map projected in three orthogonal directions.

### 6.2 Central slices

#### 6.2.1 Primary map

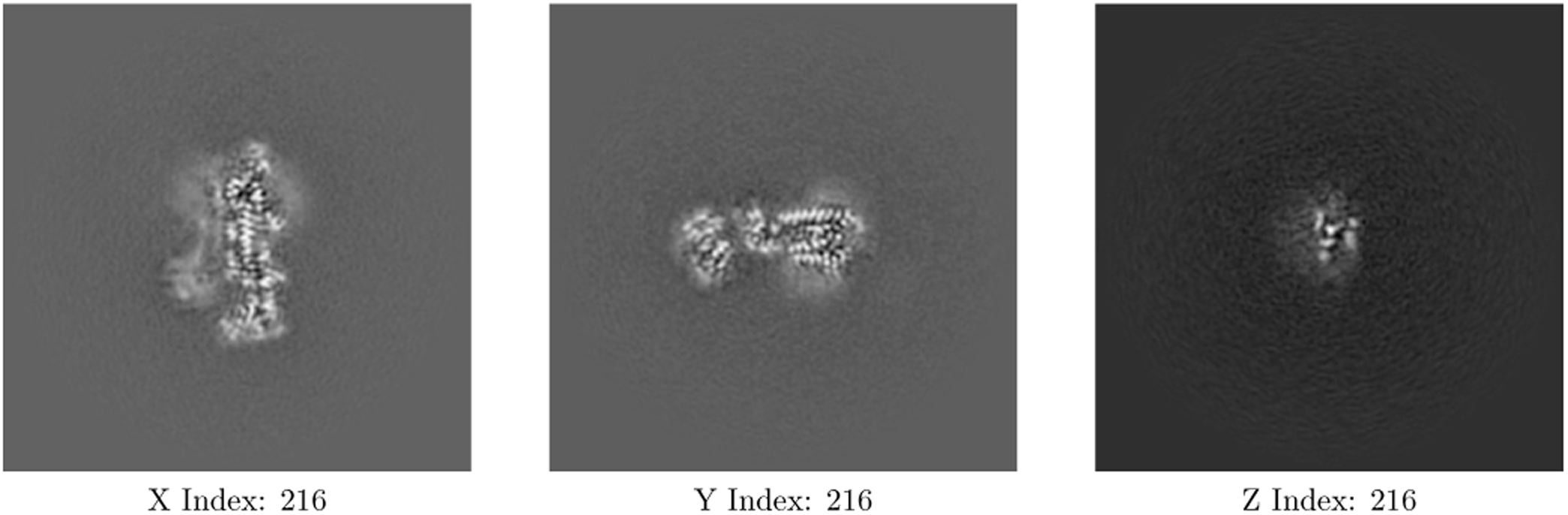

#### 6.2.2 Raw map

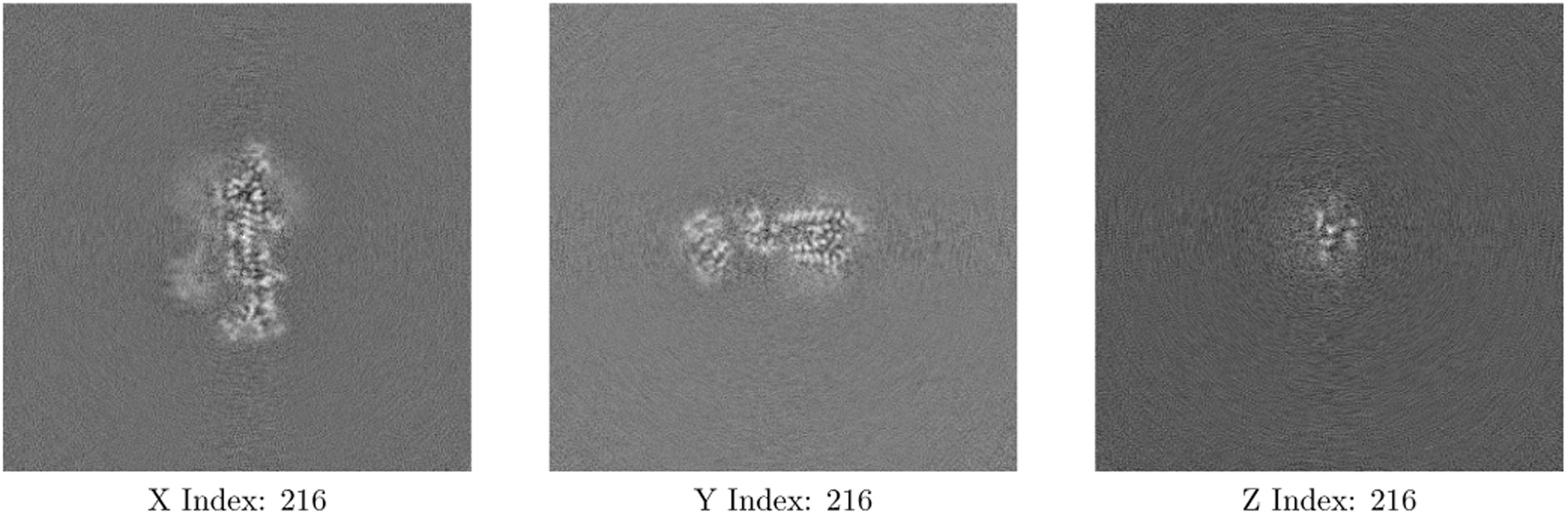

The images above show central slices of the map in three orthogonal directions.

### 6.3 Largest variance slices

#### 6.3.1 Primary map

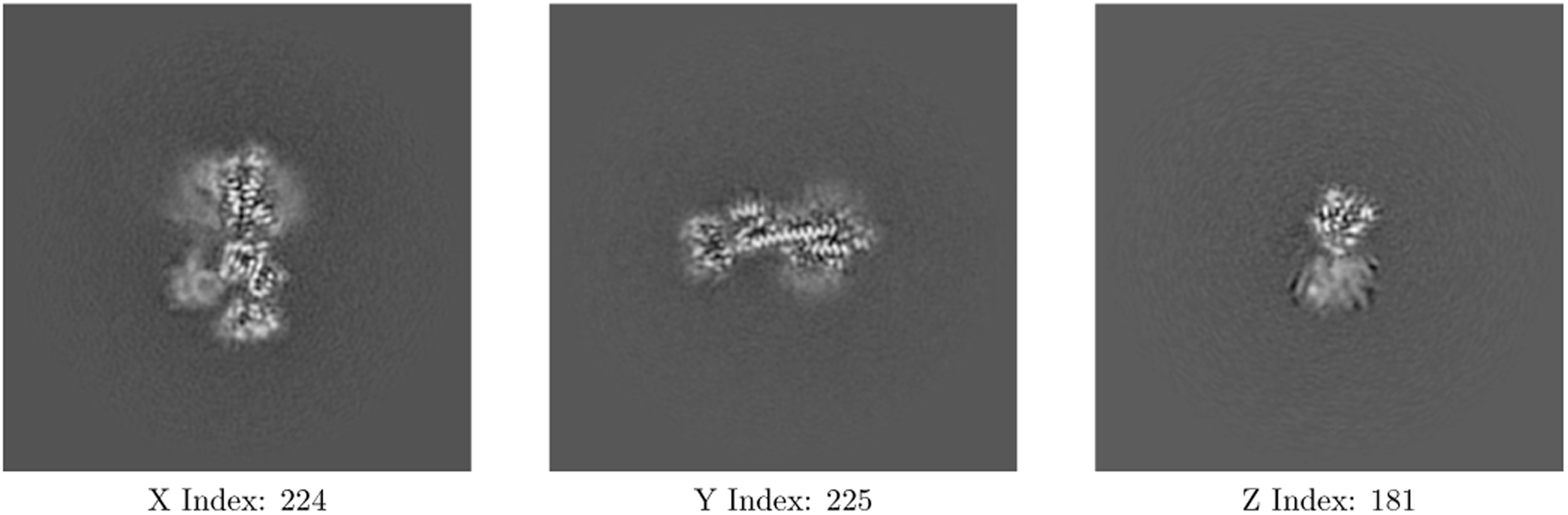

#### 6.3.2 Raw map

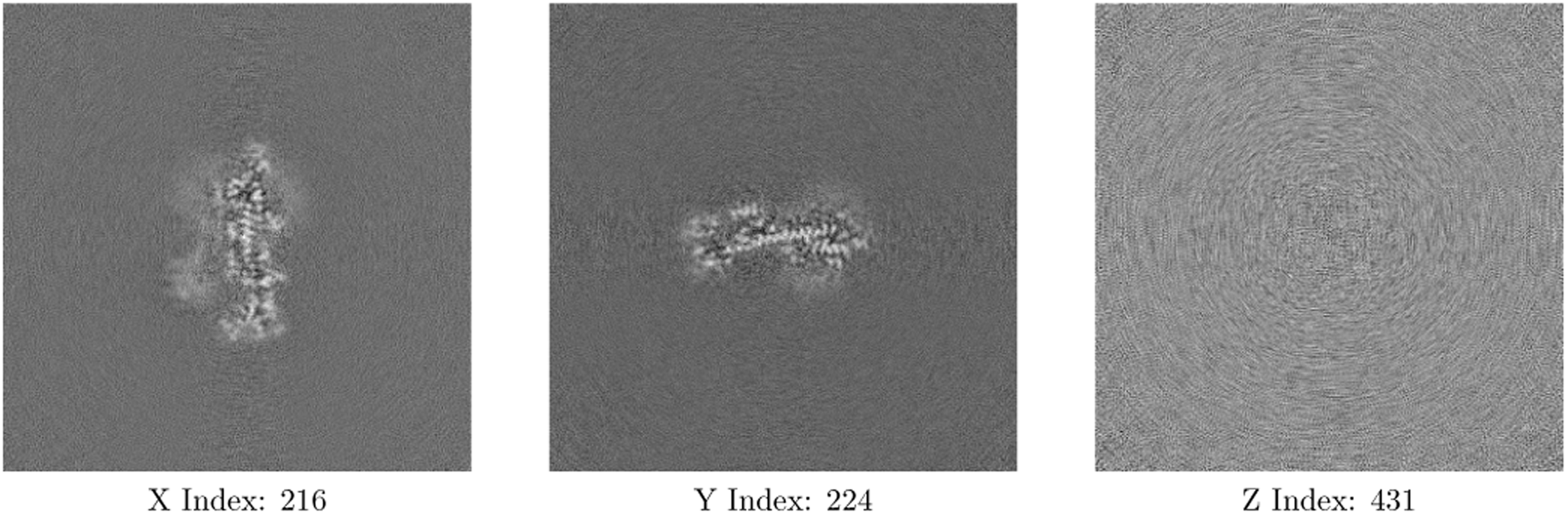

The images above show the largest variance slices of the map in three orthogonal directions.

### 6.4 Orthogonal standard-deviation projections (False-color)

#### 6.4.1 Primary map

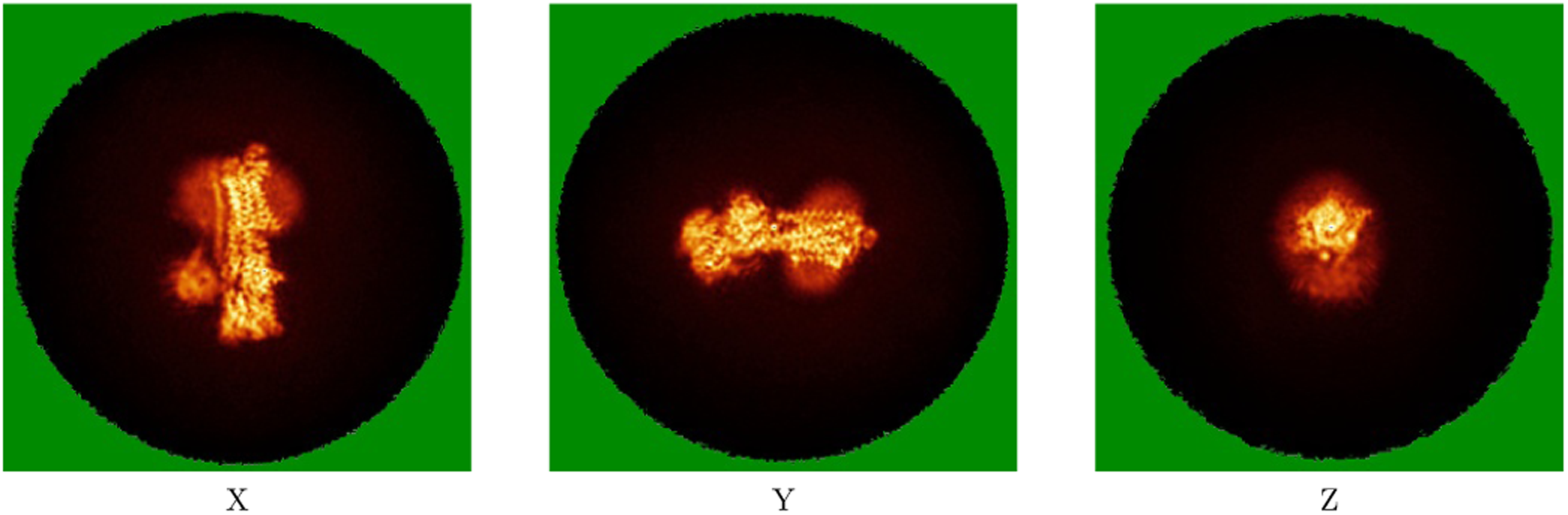

#### 6.4.2 Raw map

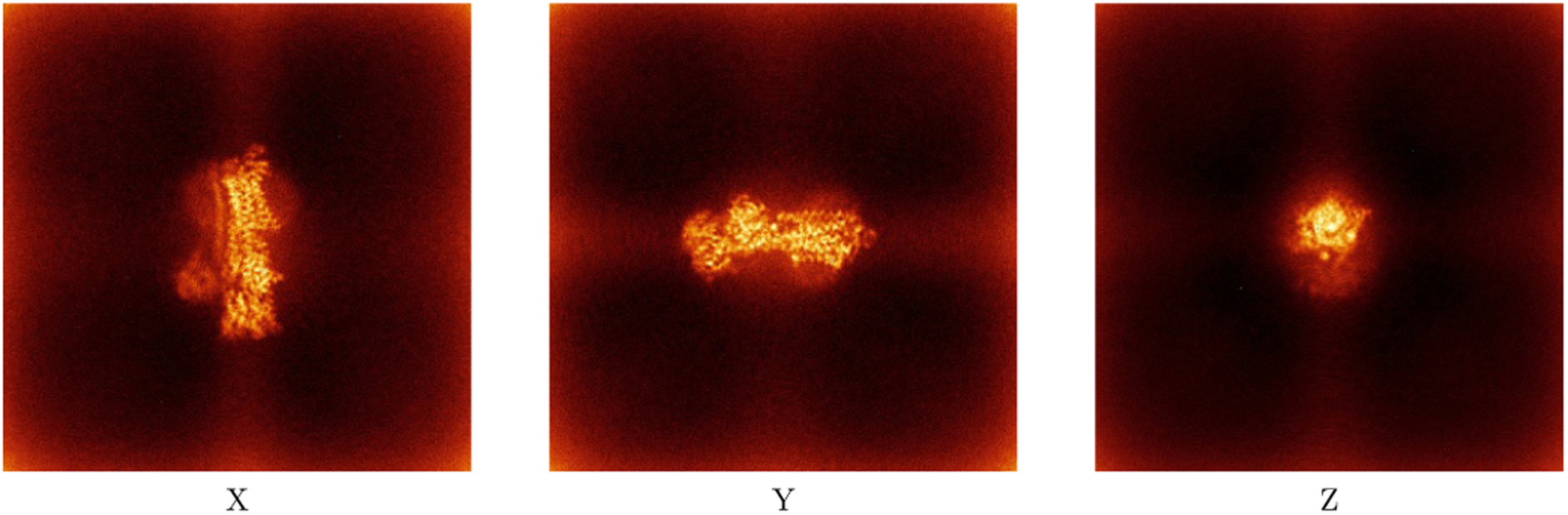

### 6.5 Orthogonal surface views

#### 6.5.1 Primary map

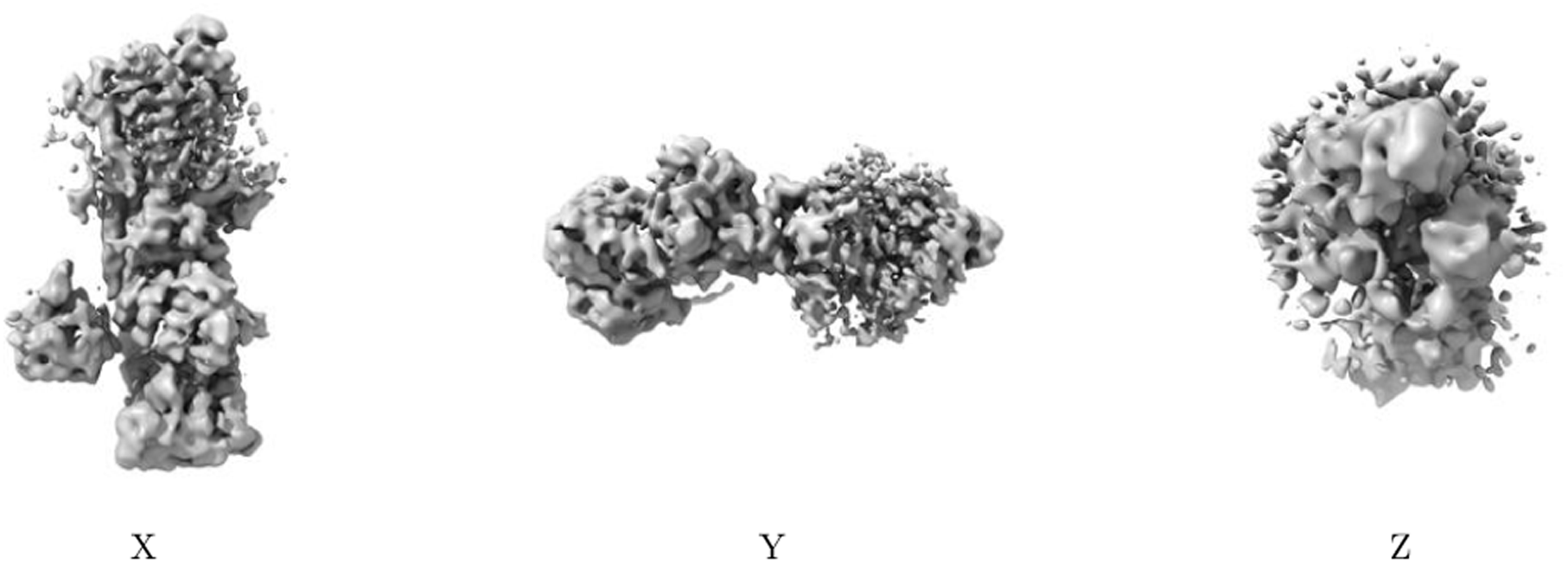

The images above show the 3D surface view of the map at the recommended contour level 0.055. These images, in conjunction with the slice images, may facilitate assessment of whether an appropriate contour level has been provided.

#### 6.5.2 Raw map

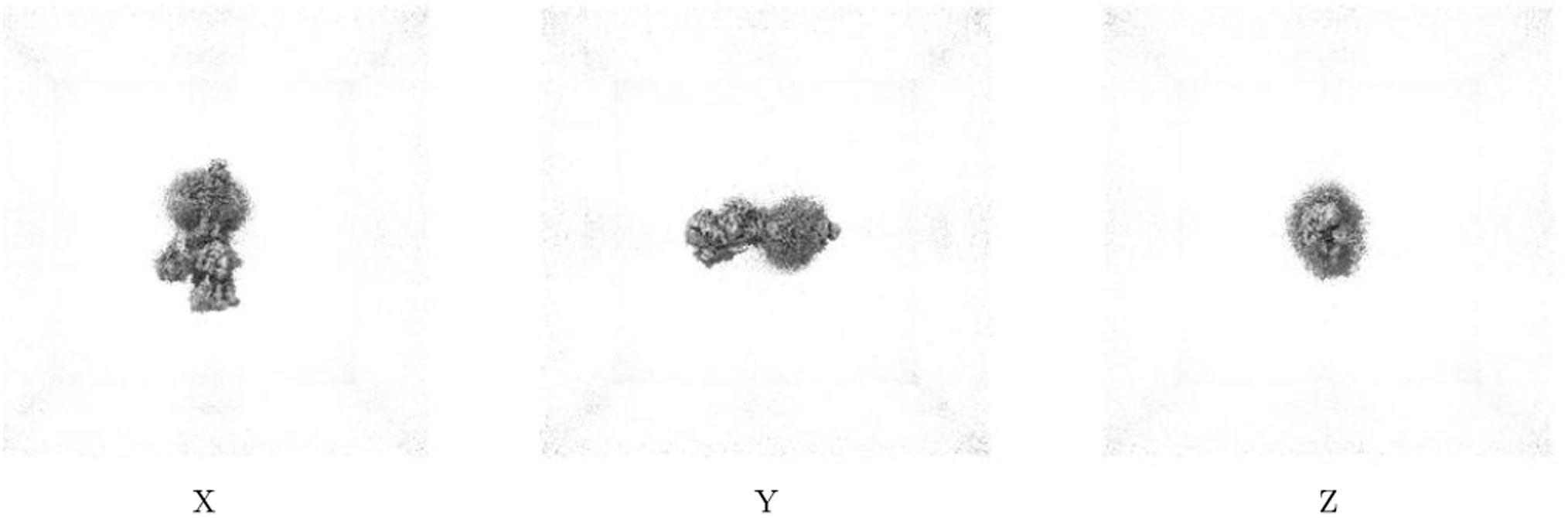

### 6.6 Mask visualisation

This section was not generated. No masks/segmentation were deposited.

## 7 Map analysis

This section contains the results of statistical analysis of the map.

### 7.1 Map-value distribution

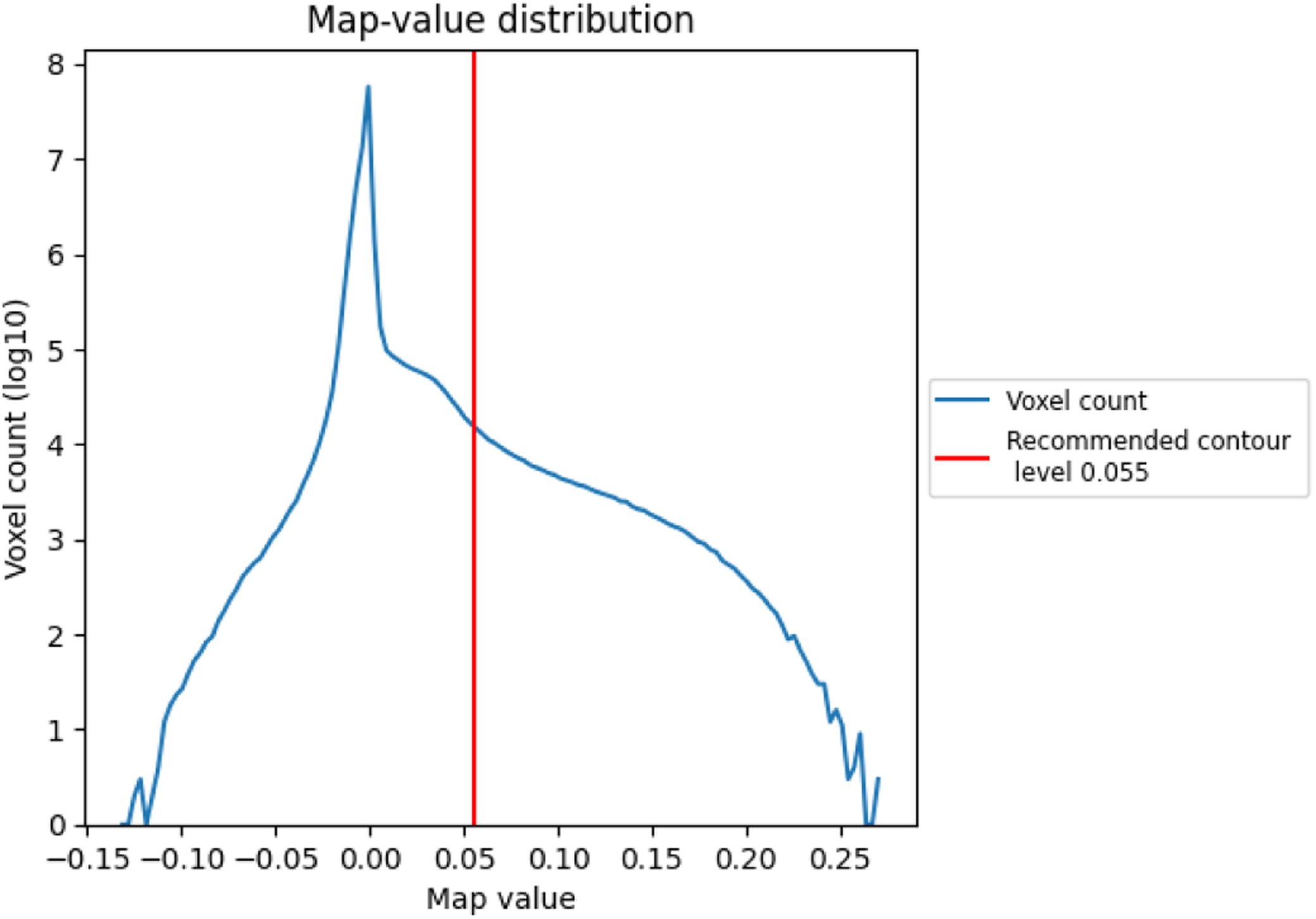

### 7.2 Volume estimate

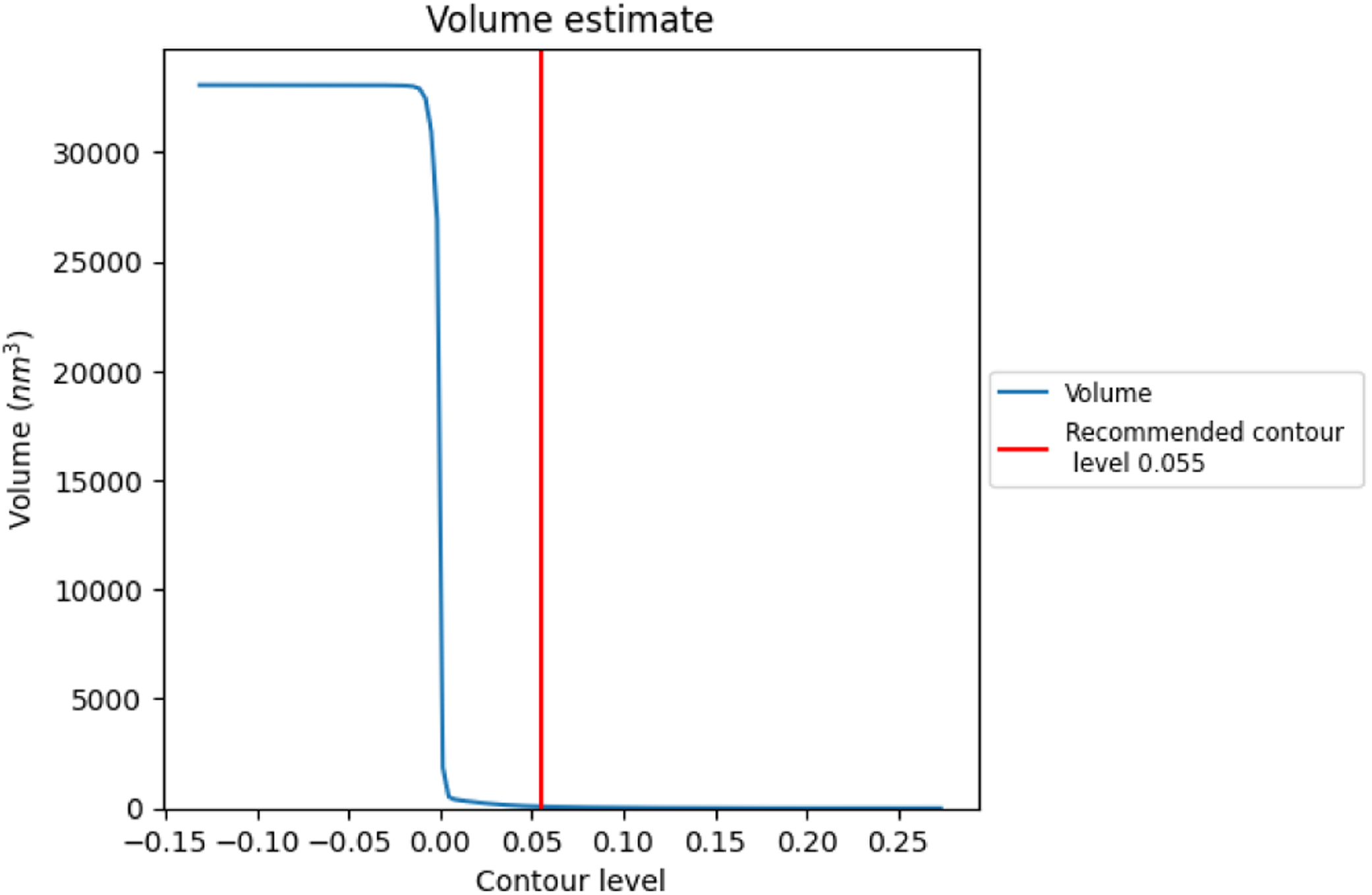

The volume at the recommended contour level is 77 nm^3^; this corresponds to an approximate mass of 70 kDa.

### 7.3 Rotationally averaged power spectrum

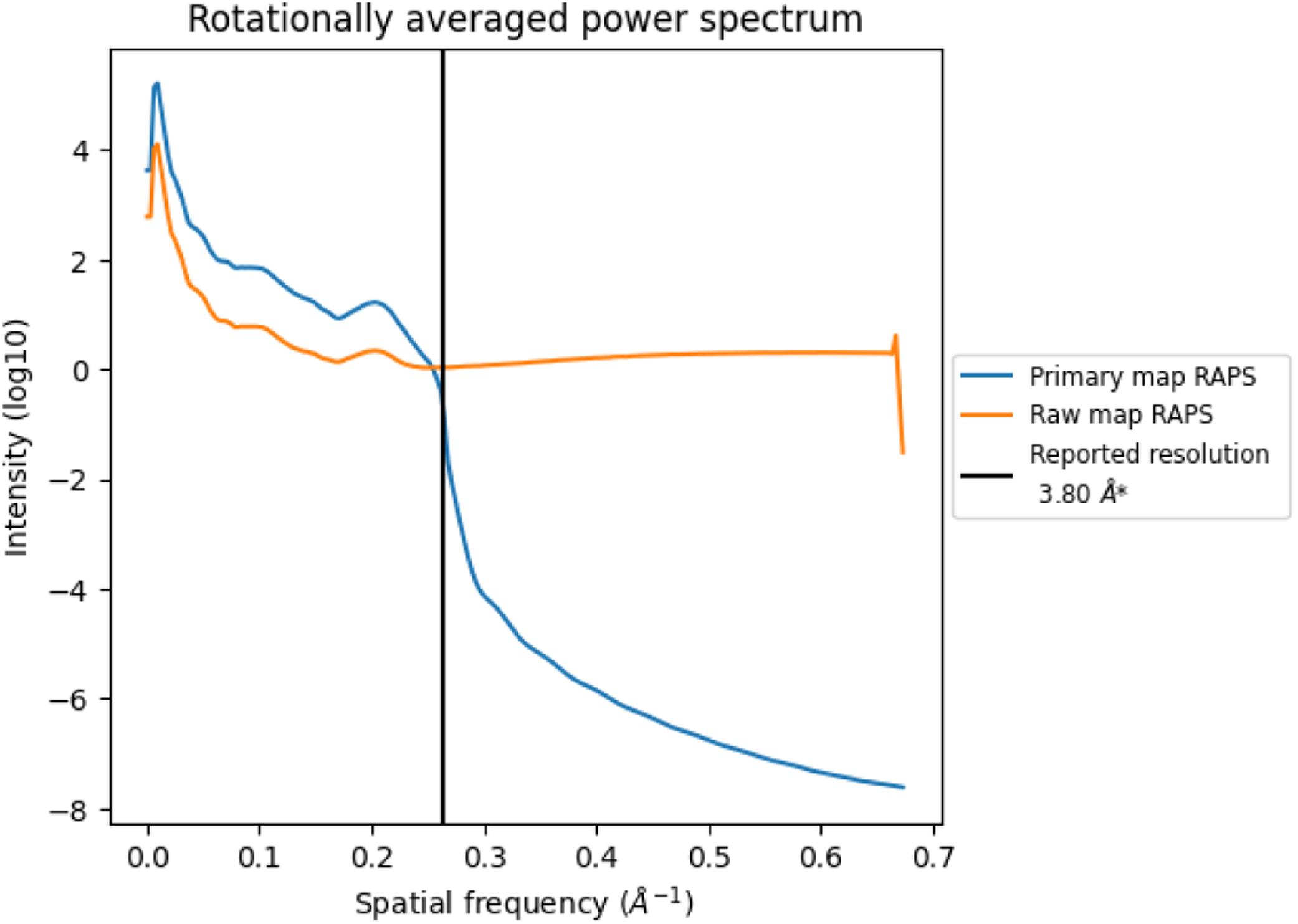

*Reported resolution corresponds to spatial frequency of 0.263 Å*^−^*^1^

## 8 Fourier-Shell correlation

### 8.1 FSC

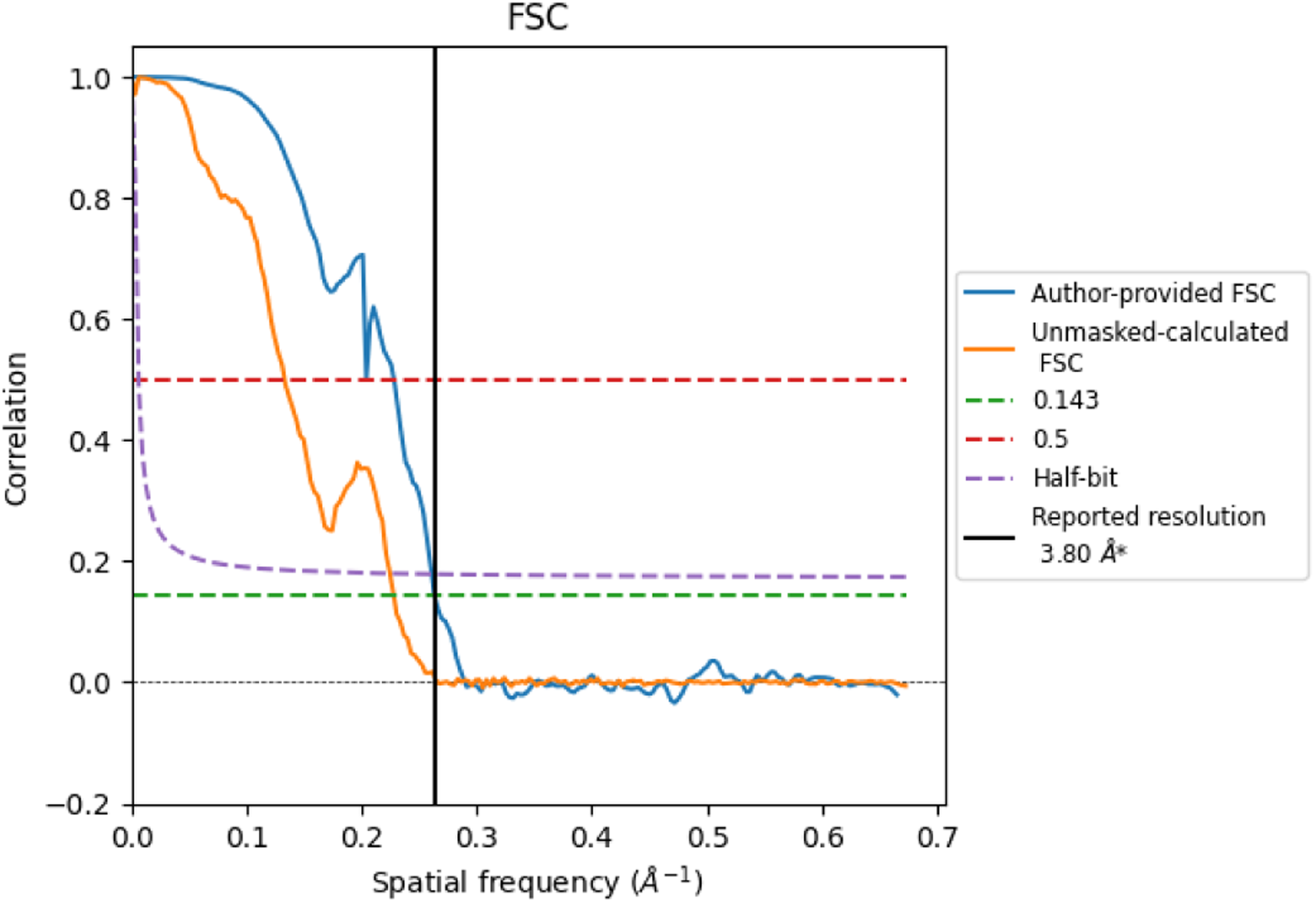

*Reported resolution corresponds to spatial frequency of 0.263 Å^-1^

### 8.2 Resolution estimates

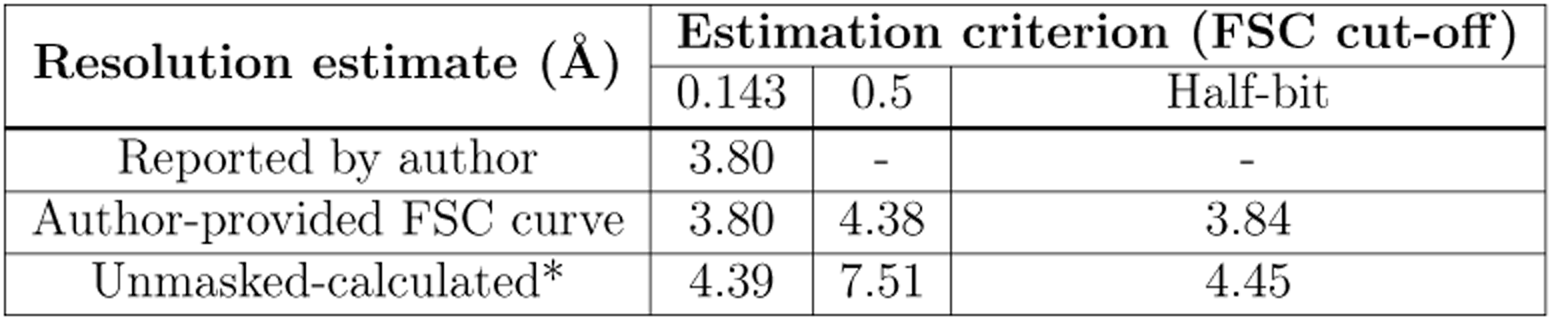

*Resolution estimate based on FSC curve calculated by comparison of deposited half-maps. The value from deposited half-maps intersecting FSC 0.143 CUT-OFF 4.39 differs from the reported value 3.8 by more than 10 %

## 9 Map-model fit

This section contains information regarding the fit between EMDB map EMD-73822 and PDB model 9Z5R. Per-residue inclusion information can be found in section 3 on page 4.

### 9.1 Map-model overlay

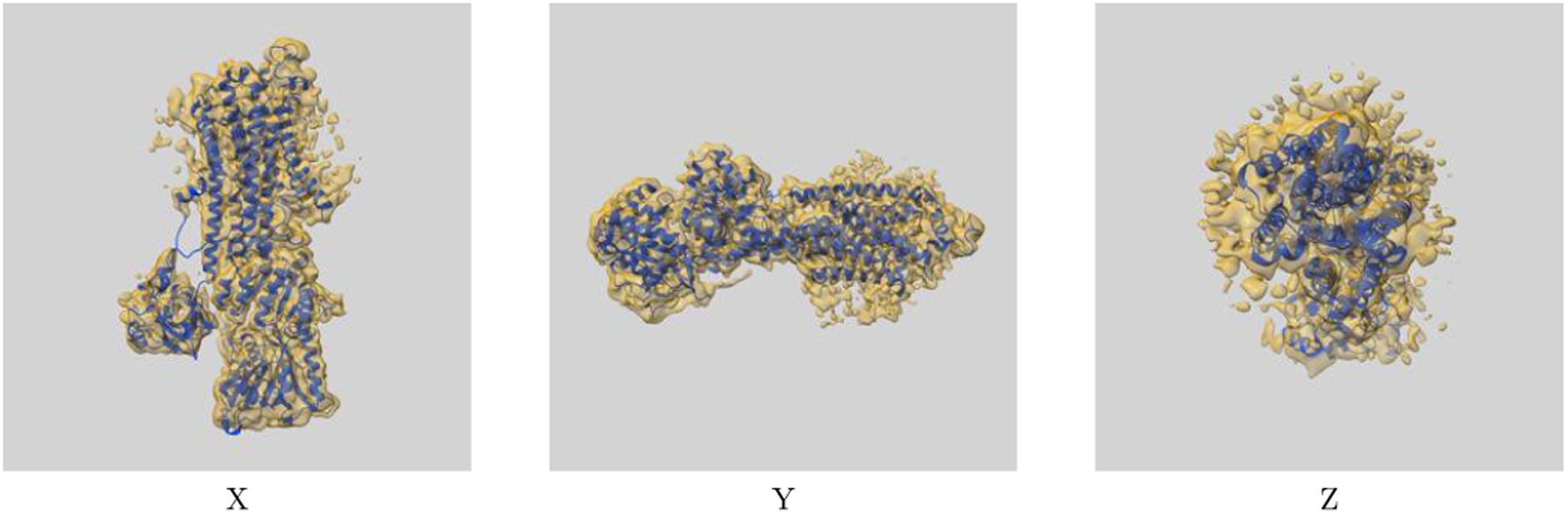

The images above show the 3D surface view of the map at the recommended contour level 0.055 at 50% transparency in yellow overlaid with a ribbon representation of the model coloured in blue. These images allow for the visual assessment of the quality of fit between the atomic model and the map.

### 9.2 Q-score mapped to coordinate model

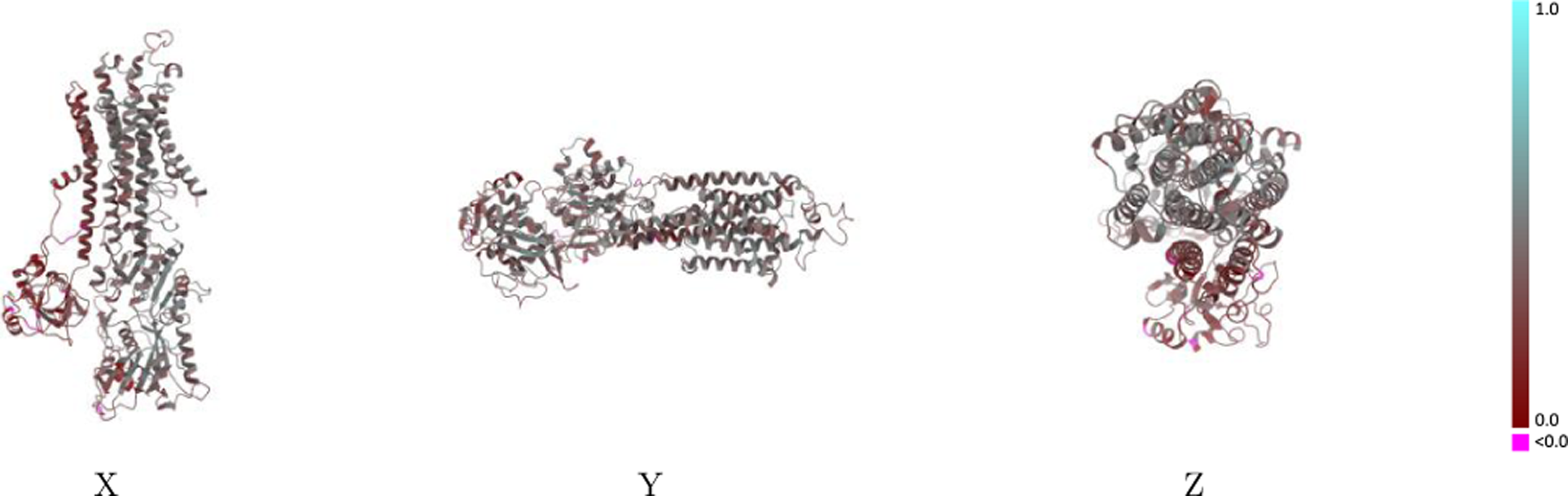

### 9.3 Atom inclusion mapped to coordinate model

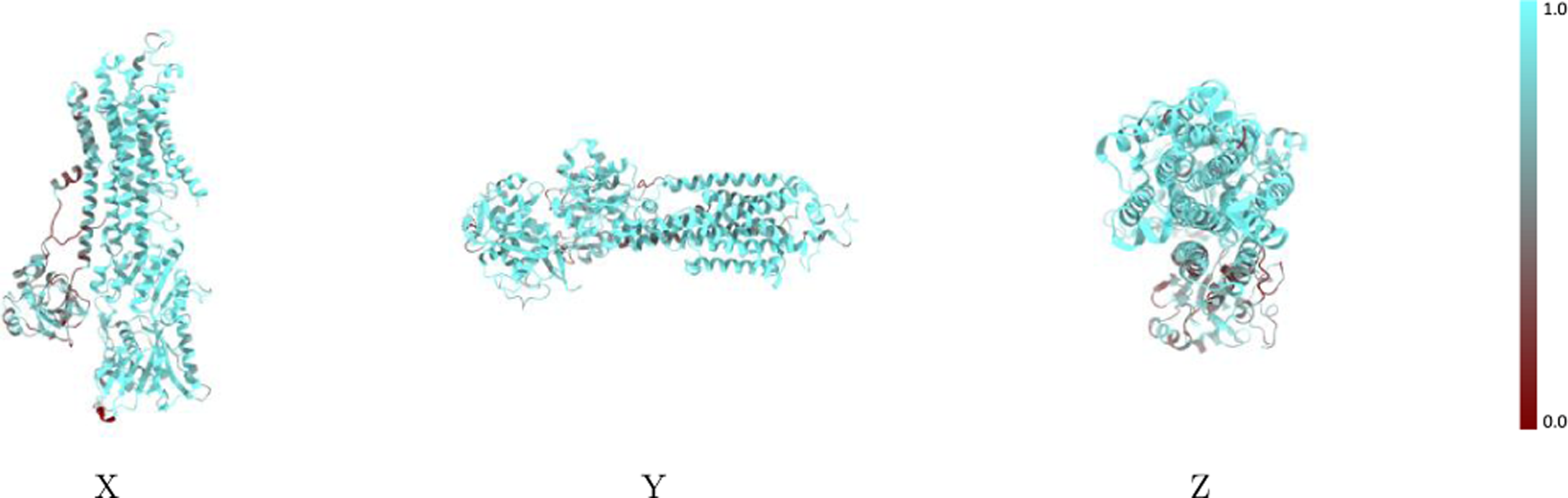

The images above show the model with each residue coloured according to its atom inclusion. This shows to what extent they are inside the map at the recommended contour level (0.055).

### 9.4 Atom inclusion

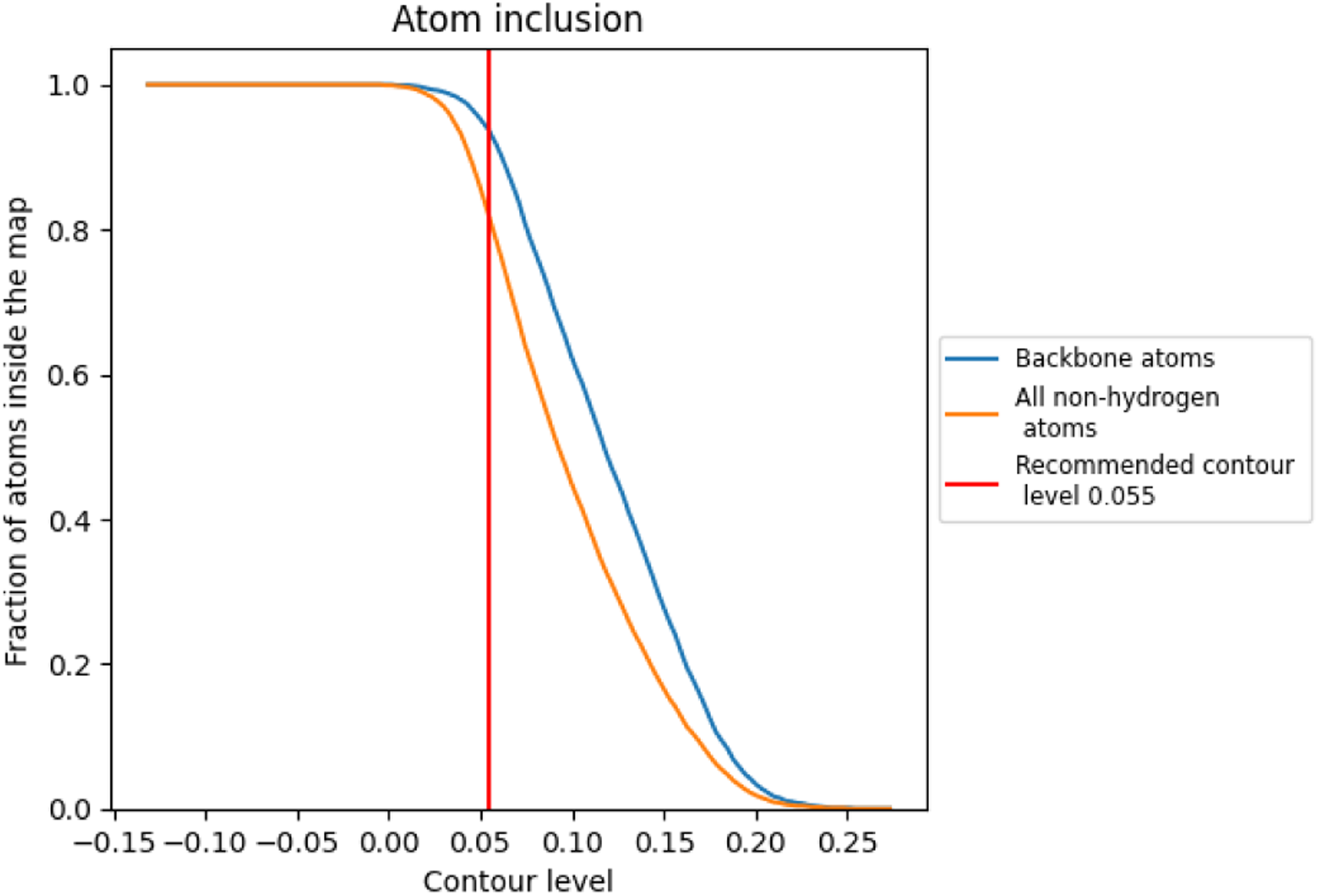

At the recommended contour level, 94% of all backbone atoms, 82% of all non-hydrogen atoms, are inside the map.

### 9.5 Map-model fit summary

The table lists the average atom inclusion at the recommended contour level (0.055) and Q-score for the entire model and for each chain.

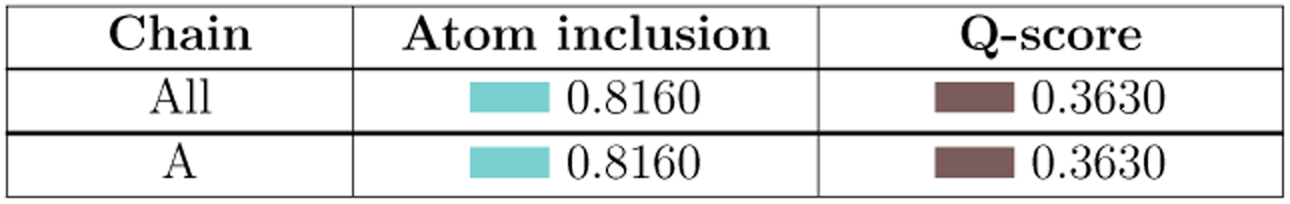

